# Comparison of conservation strategies for California Channel Island Oak (*Quercus tomentella*) using climate suitability predicted from genomic data

**DOI:** 10.1101/2024.05.23.592743

**Authors:** Alayna Mead, Sorel Fitz-Gibbon, John Knapp, Victoria Sork

## Abstract

Management strategies, such as assisted gene flow, could increase resilience to climate change in tree populations. Knowledge of evolutionary history and genetic structure of species is needed to assess the risks and benefits of different strategies. *Quercus tomentella*, or Island Oak, is a rare oak restricted to six Channel Islands in California, USA, and Baja California, Mexico. Previous work has shown that Island Oaks on each island are genetically differentiated, but it is unclear whether assisted gene flow could enable populations to tolerate future climates. We performed whole-genome sequencing on Island Oak individuals and *Q. chrysolepis*, a closely related species that hybridizes with Island Oak (127 total), to characterize genetic structure and introgression across its range and assess the relationship between genomic variation and climate. We introduce and assess three potential management strategies with different trade-offs between conserving historic genetic structure and enabling populations to survive changing climates: the *status quo* approach; ecosystem preservation approach, which conserves the trees and their associated biodiversity; and species preservation approach, which conserves the species. We compare the impact of these approaches on predicted maladaptation to climate using Gradient Forest. We also introduce a climate suitability index to identify optimal pairs of seed sources and planting sites for approaches involving assisted gene flow. We found one island (Santa Rosa) that could benefit from the ecosystem preservation approach and also serve as a species preservation site. Overall, we find that both the ecosystem and species preservation approaches will do better than the *status quo* approach. If preserving Island Oak ecosystems is the goal, assisted migration into multiple sites could produce adapted populations. If the goal is to preserve a species, the Santa Rosa population would be suitable. This case study both illustrates viable conservation strategies for Island Oak and introduces a framework for tree conservation.

## Introduction

Climate change may be proceeding too quickly for many populations to adapt to new conditions, particularly long-lived and foundational species such as forest trees, threatening not only species but the entire ecosystem (Aitken et al., 2008). However, species often contain abundant genetic variation that may enable them to adapt to novel climates, which could be a tool for management and conservation. Populations are frequently adapted to local environments that vary across the species range (Leimu & Fischer, 2008; Sexton et al., 2014; Sork, 2017; Wang & Bradburd, 2014) and this heterogeneity can enable adaptation to novel climates. In natural populations, gene flow can introduce genetic variation into a population, including alleles that are adaptive under particular environments (Savolainen et al., 2007). If these alleles are adaptive in the new environment and selection is strong enough, they will increase in frequency, enabling the population to be better adapted to environmental conditions. If adaptation is too slow to match the pace of climate change, as may be the case for species with long generation times, one potential management strategy is to intentionally introduce genetic variation that is likely to be adaptive under future conditions. This strategy, assisted gene flow, involves transferring genetic material (such as seeds or individuals) among populations within a species range (Aitken & Bemmels, 2016; Aitken & Whitlock, 2013; Browne et al., 2019).

The viability, risks, and benefits of assisted gene flow are likely to vary among species. First, assisted gene flow for climate change mitigation requires standing genetic variation that is likely to be adaptive under future climates in some parts of the species range. Additionally, if populations within a species have low levels of gene flow and are genetically diverged, introducing new alleles may cause outbreeding depression or disrupt existing local adaptation (Grummer et al., 2022). Genomic data can be useful in evaluating these requirements. A species’ genome-wide genetic structure can be used to understand historic patterns of gene flow among populations and evaluate whether introducing non-local alleles would lead to maladaptation, while putatively adaptive alleles can be used to predict whether, and where, future climate conditions match historic climates of each population. Using genotype-environment associations to detect putatively adaptive genetic variation, it is possible to identify the climate variables that are most important in explaining genetic variation, determine whether there is overlap between the historic and future climate for important climate variables, and estimate the gap between the historic climate that a population has evolved in and the future climate at its current location or throughout the species range (Fitzpatrick & Keller, 2015; Gougherty et al., 2021; Lachmuth et al., 2024; Rellstab, 2021; Rellstab et al., 2015; Sork et al., 2013). Traditionally, the goal of assisted gene flow is to enable the persistence of existing populations by introducing genetic variation likely to be pre-adapted to future climates. However, another benefit is the preservation of genetic diversity present within a population that may otherwise go extinct. Using genomic data to identify optimal matches between populations and future climate conditions, it is possible to make specific seed-sourcing recommendations (Shryock et al., 2021; Yu et al., 2022) and compare the predicted effect that different management strategies may have on the match between adaptive genomic variation and future climate.

Trees have life history characteristics that make them good candidates for the use of assisted gene flow to enable population response to climate change: they often experience high gene flow followed by strong selection on seedlings, enabling local adaptation (Alberto et al., 2013). High levels of gene flow are facilitated by their predominantly outcrossing mating system and long-distance pollen dispersal by wind (Petit & Hampe, 2006). Despite the resulting lack of strong genetic structure, local adaptation is common within tree populations, likely due to the high genetic diversity of a large panmictic species combined with high seedling mortality introducing strong selection against non-locally adapted alleles (Savolainen et al., 2007, 2013; Sork, 2016, 2017; Sork et al., 2013). Strong selection pressure against deleterious alleles, whether introduced through natural or human-assisted gene flow, should reduce their introduction into the gene pool. Landscape genomic studies generally support these characteristics of trees, finding that genetic differences among populations increase with geographic distance (a pattern of isolation by distance), but that environmental variables also play a role in determining genetic variation across the range (Fitzpatrick & Keller, 2015; Gugger et al., 2021; Jia et al., 2020; Martins et al., 2018).

Many species that are the focus of conservation efforts have restricted ranges and/or fragmented populations, and it is unclear whether assisted gene flow is a viable strategy for such species, or whether they should be conserved using other methods. Here we focus on Island Oak *(Quercus tomentella)*, a relictual island tree species existing on only six islands, as a case study in using genomics to compare specific assisted gene flow scenarios. In comparison to widespread tree species, rare species occurring on islands may be comprised of disjunct populations with limited gene flow (Di Santo et al., 2022; Gugger et al., 2018). If gene flow among islands is restricted, populations may be too deeply diverged for successful assisted gene flow. If the species does not occur across a range of climates and contain sufficient adaptive genetic variation to those climates, standing genetic variation that will be beneficial under future climates may not be available to introduce into other populations. Studies across a wide range of taxa have found that island species have genetic or phenotypic variation that is associated with climate or habitat variation, suggesting local adaptation (Cheek et al., 2022; Gamboa et al., 2022; Langin et al., 2015). If island species contain sufficient variation in climate-adaptive alleles across populations, assisted gene flow could be a viable strategy for their conservation.

Here we present different strategies for conserving Island Oak under future climate conditions, comparing the outcomes with the goals of preserving Island Oak to retain its associated ecosystem or to simply preserve the species, and provide a case study for translating genomic data into predictions of risk. We implemented a landscape genomics analysis to identify associations between genetic variation and climate and identify putatively adaptive SNPs. We used the genomic offset, calculated from the genomic-informed difference between a population’s historic climate and the future climate at its local site or other sites where it could be planted (Capblancq et al., 2020; Fitzpatrick & Keller, 2015; Gougherty et al., 2021; Lachmuth et al., 2024), as a proxy for maladaptation. We also developed the climate suitability index, a novel comparison metric derived from the genomic offset that measures the similarity of climates, as scaled by their importance in contributing to genetic variation. We compared genomic offset and climate suitability for Island Oak populations under three conservation strategies:

1. ***Status quo* approach**: The simplest approach, and the one currently used, is to protect existing populations and augment oak groves by planting acorns from the same island or watershed. This approach maintains the historic genetic population structure, but may not enable populations to survive climate change.
2. **Ecosystem preservation approach**: To maintain existing oak groves and re-establish previous ones so that Island Oak ecosystems and their associated biodiversity are sustained, restoration projects could use assisted gene flow to create a viable oak population under future climates by introducing genotypes that are likely to be best-adapted to future climate conditions at a planting location.
3. **Species preservation approach**: When a species is threatened throughout its native range, it is sometimes necessary to focus on strategies that will preserve the species, either *in situ* or *ex situ*. *In situ* sites could include existing oak groves, and/or they could be sites on the islands where they are not present, but have historically existed. Alternatively, species preservation could be accomplished through *ex situ* planting sites on the mainland (Rosenberger et al., 2022; Westwood et al., 2021). As *ex situ* preservation typically cultivates trees in botanical gardens and arboreta, targeting sites with optimal climates is less important than for wild populations, so we focus here on *in situ* preservation of Island Oaks in their native range.

In this study, we analyzed whole-genome sequencing data collected from the entire species range of Island Oak to assess the best strategy for preserving the species and its associated ecosystems under future climate change scenarios on the California Islands (Figure 1 B, C). Our goals were: 1) Characterize the processes structuring genetic variation in *Q. tomentella*, such as gene flow, hybridization, geography, and climate. We documented the genetic structure of *Q. tomentella* along with the closely related species *Q. chrysolepis* and their hybrids to describe the historic patterns of divergence and gene flow. 2) Compare the effect of three different management strategies, including two assisted gene flow strategies addressing different management goals, on the predicted maladaptation of Island Oak under future climates. While a combination of ecological and restoration management strategies will likely be needed to conserve this species, here we focus on the potential contribution of assisted gene flow to their success.

**Figure 1.**
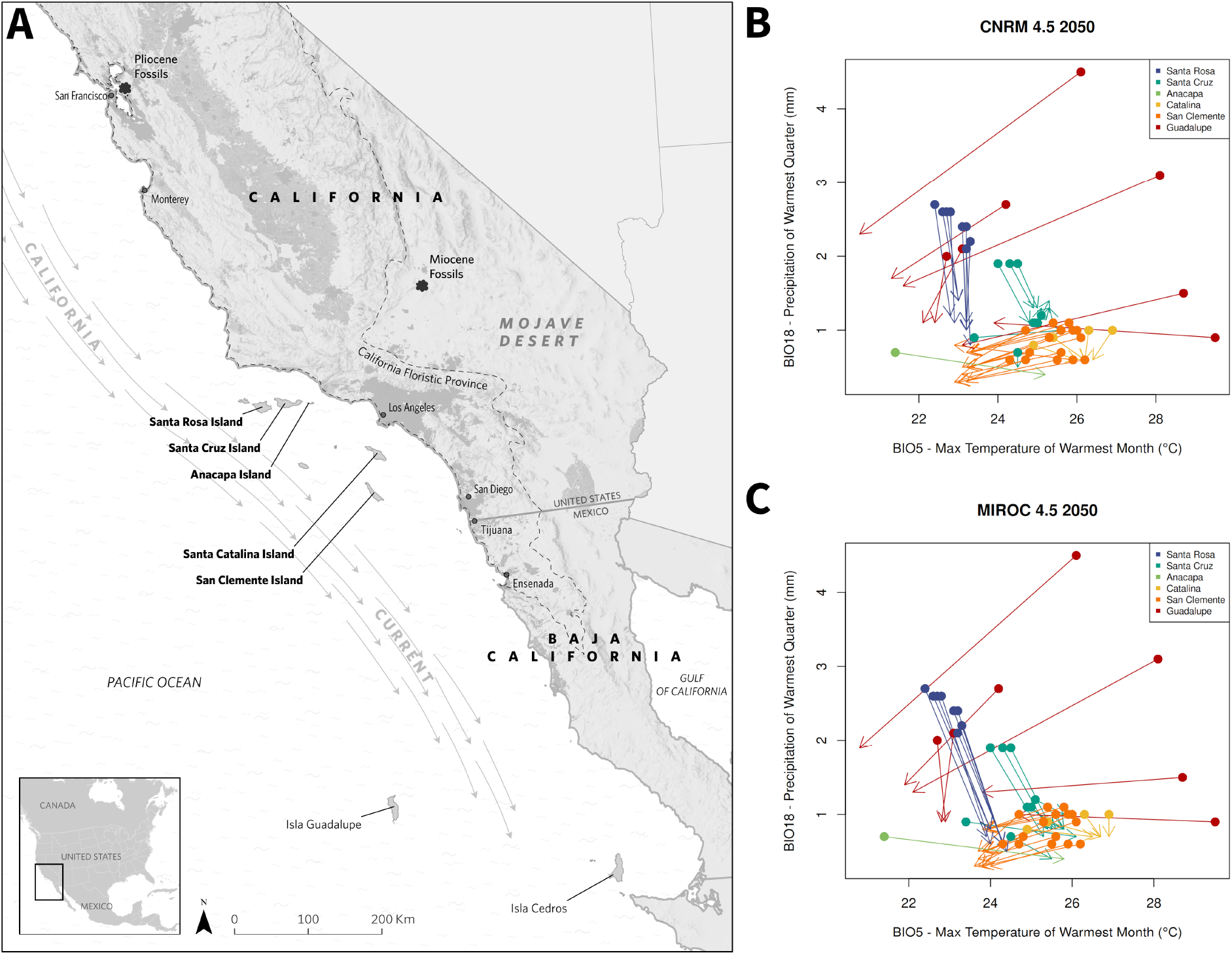
A) Map showing location of the six islands sampled, relation to the mainland, and direction of the California Current. Locations of fossil sites of *Q. declinata*, the fossil equivalent of *Q. tomentella*, from the Miocene (Axelrod, 1939) and Pliocene (Axelrod, 1944) are marked on mainland California. Isla Cedros to the southeast is shown where *Q. cedroensis* occurs. Dark grey shading indicates developed land. Map credit: Irina Koroleva, The Nature Conservancy. B and C) Predicted shifts in climate (from the periods 1970-2000 to 2041-2060) at each sampled location based on max temperature of the warmest month (BIO5) and precipitation of the warmest quarter (BIO12) for two climate models, CNRM (B) and MIROC (C) under Representative Concentration Pathway (RCP) 4.5, in which emissions peak in 2040 then decline. Closed circles indicate historic climate for a given location (1970-2000), and arrowheads indicate the projected future climate (2041-2060) for the same location.

## Methods

### Study species

*Quercus tomentella* (section *Protobalanus)*, or Island Oak, is a rare relictual island tree found on six of the California Islands off the western coast of the United States (the Channel Islands) and Mexico (Isla Guadalupe) (Figure 1A). This species is categorized as endangered by the IUCN (Beckman & Jerome, 2017), and its threats include browsing from non-native herbivores, erosion, changes in hydrology, and climate change (Beckman et al., 2019; Beckman & Jerome, 2017); and on Santa Rosa Island, the loss of a native disperser, the Island Scrub-jay (Delaney & Wayne, 2005; Morrison et al., 2011). Island Oak forms groves that co-occur with island endemic plant species and varieties, forming unique ecosystems that support animal species including the Island Scrub-jay (Pesendorfer et al., 2018; Sawyer et al., 2009). Based on microsatellite markers, *Q. tomentella* is genetically differentiated across islands and has lower genetic diversity than its more widespread relative *Q. chrysolepis* (Ashley et al., 2018). Additionally, it can propagate clonally, which may contribute to its lower genetic diversity. However, to understand how selection has shaped adaptive genetic variation within and across islands, we used whole genome sequence data, which will often reveal different spatial patterns than neutral markers such as microsatellites (Fitzpatrick & Keller, 2015; Gugger et al., 2021; Martins et al., 2018) and provide a better understanding of adaptive potential. While Island Oak currently has a restricted and fragmented range, it was once widespread throughout mainland California during the Miocene and Pliocene (Figure 1A), then became restricted to the coast, and eventually the California Islands as the mainland climate became less temperate (Axelrod, 1967; Muller, 1965). During glacial periods with lower sea levels, the Northern Channel Islands were connected as a single island, called Santa Rosae, as recently as 11,000 years ago (Kennett et al., 2008; Reeder-Myers et al., 2015). During this time, there may have been more gene flow among Santa Rosae Island populations. It is also possible that gene flow occurred through translocation by Indigenous peoples, as acorns are an important food source, although no direct evidence exists (Rick et al., 2019). Hybridization may be a source of genetic variation: *Q. tomentella* hybridizes with *Q. chrysolepis*, a widespread species on mainland California that is also present on five of the islands (Ashley et al., 2018), and genomic and fossil data support a history of co-occurrence and ancient introgression between the two species (Axelrod, 1944b, 1944a; Ortego et al., 2018).

### Collections

Leaf samples from *Q. tomentella* individuals were collected across the six islands encompassing the species range. Because *Q. tomentella* hybridizes with *Q. chrysolepis*, we also collected co-occurring *Q. chrysolepis* and putative hybrid individuals from the islands, and *Q. chrysolepis* individuals from across northern and southern mainland California (southernmost in San Bernardino county, northernmost in Siskiyou county, Table S1). These two species are distinct from other oak species present on the islands, but hybrid or introgressed individuals can have intermediate characteristics. Collectors attempted to distinguish between the two species but field classifications of intermediate individuals may have varied by collector. We ultimately defined species using the genomic data. We attempted to avoid collecting multiple samples from the same clone by collecting from only one stem within the same grove or cluster of stems, with the exception of Anacapa Island where only one grove of trees exists. We primarily collected mature trees, but some saplings were included when this was not possible. Leaf samples were either dried on silica or were placed on ice, then were frozen at -80° C. In total, based on species identifications in the field, we sequenced and analyzed data for 107 *Q. tomentella* individuals, 17 *Q. chrysolepis* individuals (8 from mainland California and 9 from the islands), and 3 putative hybrids. On average we collected 20 individuals per island (minimum 5, maximum 30).

### DNA extraction and sequencing

Leaves were hand-ground in liquid nitrogen using a mortar and pestle, and approximately 50 mg of tissue was used for DNA extraction. DNA was extracted from leaves using a modified version of the Qiagen DNeasy Plant Mini Kit protocol. First, to remove polyphenols, a prewash step was performed twice. One mL of prewash buffer was added to ground leaf tissue, ground in a bead mill for 20 seconds, centrifuged for 10 minutes at 10,000 RPM, and the supernatant was discarded. Prewash buffer consisted of (per sample) 100 ul Tris, 100 ul EDTA, 200 ul 5 M NaCl, 600 ul molecular grade water, and 0.01 g PVP. Following the prewash, the Qiagen protocol was followed. For the silica-dried leaf samples (from Guadalupe and San Clemente Islands) we were unable to extract sufficient amounts of DNA, so extraction was performed by the California Conservation Genomics Project mini-core using the Macherey-Nagel NucleoMag Plant kit, with the following modifications: PVP and Proteinase K were added during the digest, and an additional ethanol wash was added before elution. Extracted DNA was sent to UC Davis DNA Technologies and Expression Analysis Cores for library preparation using a custom SeqWell kit which used half the standard volume of reagents compared to a standard kit, and was shown by SeqWell to work well for our samples. Whole-genome sequencing was performed on a NovaSeq 6000 using 150 bp, paired-end sequencing.

### Filtering and variant calling

Adapters were trimmed from raw reads using Trim Galore, and reads with a length less than 20 bp were removed. Reads were not trimmed based on quality scores during this step. Reads were aligned to the annotated *Q. lobata* genome (Valley Oak Genome 3.2, Sork et al., 2022), rather than the Island Oak genome, because it is equally distant to both *Q. tomentella* and *Q. chrysolepis* and is annotated. Aligning to a different species (85.8% genetic similarity, Mead et al., 2024) may exclude some loci that are highly diverged between the two species, but our goal was to characterize general spatial patterns of genetic variation rather than identify all potential candidate genes. Alignment used BWA-MEM, with ‘markShorterSplits’ and ‘readGroupHeaderLine’ options enabled. Duplicate reads were marked and removed using GATK MarkDuplicates. Variants were called using GATK HaplotypeCaller with the ‘emit-ref-confidence’ option set to ‘GVCF’. Variants were hard-filtered using GATK VariantFiltration, with SNPs and indels filtered separately. For SNPs, we removed variants with quality by depth (QD) <2, quality (QUAL) <30, mapping quality (MQ) < 40, phred-scaled strand bias (FS) >60, symmetric odds ratio strand bias (SOR) >3, mapping quality rank sum (MQRankSum) < -12.5, and read position rank sum (ReadPosRankSum) < -8. We removed indels with QD<2, FS>200, QUAL<30, and ReadPosRankSum< -20. Repetitive regions of the genome were removed using vcftools (Danecek et al., 2011) based on the reference genome.

From this set of high-quality variants, we selected biallelic SNPs with high coverage across all samples for further analysis. Using bcftools (version 1.15.1, Danecek et al., 2021), we selected only biallelic SNPs, and removed SNPs with a mean depth across all samples <5, set individual genotypes with depth <5 to missing, removed SNPs with a minor allele frequency <0.01, and removed SNPs that were missing in >10% of individuals. The resulting filtered VCF file was converted to BED file format using PLINK (version 1.90b6.26, Chang et al., 2015), and variants in linkage disequilibrium (LD) were pruned using a window size of 50 variants, a window shift value of 10 variants, and an R^2^ threshold of 0.1, which identifies variant pairs with a correlation >0.1 within the given window and prunes them until no correlated pairs remain. Unless otherwise noted, analyses were run on this filtered and LD-pruned dataset of 585,298 SNPs. For analyses requiring no missing data, SNPs were imputed by assigning missing individuals the most common SNP (total missingness in the filtered dataset was 5%).

Climate variables were extracted for each locality from WorldClim (version 2.1, historic climate data for 1970-2000) at 30-second resolution. Island Oaks are present on five Channel Islands off the coast of California, USA and on Guadalupe Island, Mexico (six islands in total). Of the five Channel Islands with Island Oaks present, the Northern Channel Islands (Santa Rosa, Santa Cruz, and Anacapa) are cooler and receive more precipitation than the Southern Channel Islands (Santa Catalina and San Clemente, Figure 1B and C). Precipitation seasonality varies across an east/west gradient in the Northern Channel Islands, with the western part of the range experiencing less precipitation seasonality and more summer rainfall. All experience a Mediterranean climate, with warm, dry summers. Guadalupe Island, off the coast of Baja California, Mexico, has a steep elevation gradient, warmer winter temperatures, and less precipitation, but has less yearly variation in temperature and precipitation, and receives more precipitation during the dry season than the Channel Islands. We also extracted future predictions of climate at our sample sites using downscaled CMIP5 climate models from WorldClim averaged over 2041-2060 (hereafter referred to as 2050 climate) under two RCP scenarios: 4.5 (with emissions peaking around 2040) and 8.5 (rising emissions through the 21^st^century, IPCC 2014). We choose two climate models with different outcomes for Southern California, CNRM-CM5 (a warmer/wetter scenario) and MIROC-ESM (a hotter/drier scenario) (Underwood et al., 2019).

### Genetic Structure

Divergence among island populations was calculated using Weir and Cockerham’s F_ST_ for the full set of filtered, but not LD-pruned, SNPs using fast-wcfst (Fontenot, 2024, commit 338682f0b1). We converted the variance returned from fast-wcfst to standard deviation and calculated F_ST_ values for one standard deviation above and below the average as a measure of variance among loci. To understand the effect of introgression on genetic structure, we also calculated F_ST_ among ancestry groups within islands, designating each individual as *Q. chrysolepis*, Channel Islands *Q. tomentella*, or Guadalupe *Q. tomentella* if ancestry at K=3 was ≥ 0.9 for either of the three groups, and otherwise designating it as a hybrid. To perform a principal components analysis (PCA) on the imputed SNP set, we used the R package vegan (version 2.6-2, Oksanen et al., 2019). Because Guadalupe Island trees were divergent from the other populations (see results), we also ran a PCA with those samples excluded to better visualize the genetic structure among the remaining islands. ADMIXTURE (version 1.3.0, Alexander et al. 2009) was used to estimate the ancestry of individuals and characterize genetic structure across the species range. For the entire dataset, we ran ADMIXTURE for K values 1-10, where K is the number of hypothetical ancestral populations. We also ran ADMIXTURE on subsets of the data to investigate fine-scale genetic structure for the following sets: all samples except Guadalupe Island (K=1-10) and individuals from Santa Cruz and Santa Rosa islands alone (K=1-5), testing lower K values because cross-validation errors for the full dataset increased with higher K values (see results and Figure S2). Because Guadalupe Island populations appear deeply diverged from the other Island Oak populations and are unlikely to experience contemporary gene flow, we ran further analyses of climate-associated genetic variation separately for the five northernmost islands (hereafter, Channel Islands) and for Guadalupe Island. Results from ADMIXTURE and PCA (see below) suggest a history of introgression between the two species on the islands rather than occasional hybridization events producing F_1_ offspring. Because introgression can be a source of genetic variation for natural selection (Suarez-Gonzalez et al., 2018), we decided to include the hybrid individuals and those that morphologically appeared to be *Q. chrysolepis* from the islands in further analyses of putatively adaptive genetic variation.

We evaluated the effect of geography and climate on genetic differentiation among individuals. First, we tested whether individuals showed a pattern of isolation by distance by calculating the genetic distance between each pair of individuals as the proportion of variable SNPs not identical by state (calculated using PLINK with the --distance option), and testing for a relationship with geographic distance. To assess the effects of geography and climate on genetic variation in the Channel Islands populations, we used a partial redundancy analysis (RDA) in the vegan package, which partitioned the genetic variance into proportions that could be statistically explained by climate, by geographical location (via latitude and longitude), by each of these factors alone while controlling for the other, and by their joint influence. We tested the significance of each explanatory factor using the `anova.cca` function with 99 permutations.

An RDA was also used to identify candidate SNPs that are associated with climate variables. Because our goal was to characterize broad spatial patterns of climate-associated SNPs rather than identify specific climate-adaptive genes, we used the LD-pruned set of genes. Candidate SNPs identified here are not necessarily involved in climate adaptation, but may be linked to such regions. We reduced the climate and environmental variables to a subset with correlations ≤0.7, following Dormann et al. (2013): BIO5 (Max Temperature of Warmest Month), BIO6 (Min Temperature of Coldest Month), BIO15 (Precipitation Seasonality), BIO18 (Precipitation of Warmest Quarter), BIO19 (Precipitation of Coldest Quarter), and elevation. A preliminary analysis found that candidate SNPs were strongly influenced by the five individuals from Anacapa, which are from one small grove and had high genetic similarity to each other. Many of the resulting candidate SNPs were associated with the maxiumum temperature of the warmest month, which is lowest at the Anacapa site. In further analysis with Gradient Forest (see below), including the Anacapa samples appeared to result in overfitting of the model, with unsampled regions that had similar climate to Anacapa clustering separately from all other regions. Since these trees are relatively isolated, it is difficult to determine whether this association is the result of natural selection on temperature or due to inbreeding within this population, so we selected the candidate SNPs using an RDA on the four larger Channel Islands (Santa Rosa, Santa Cruz, Catalina, and San Clemente), all of which had samples collected in multiple regions within the island. We replicated the analysis to identify candidate SNPs on Guadalupe Island, which is genetically diverged from the other islands but has relatively high genetic diversity and climatic variation, potentially allowing for local adaptation within the island. Following Forester et al. (2018), we identified candidate SNPs as those that were strongly associated with the multivariate climate space. We considered candidate SNPs to be those that were outliers (more than 4 standard deviations) on the first three RDA axes. We did not correct for population structure because genetic differentiation among islands was relatively low (see Results, Table 1) and the partial RDA showed a lesser influence of geography on genetic variation after controlling for climate (see Results, Table 2), and because correcting for population structure in an RDA results in reduced power and increased false positive rates in systems with low population structure (Forester et al., 2018).

**Table 1.** Pairwise *F*_ST_ (Weir and Cockerham’s) across all island samples and mainland *Q. chrysolepis* samples. Parentheses indicate one standard deviation above and below the mean *F*_ST_ value across all loci evaluated. Negative values do not have biological meaning different from a value of zero (Willing et al., 2012).

|  | Mainland | Santa Rosa Island | Santa Cruz Island | Anacapa Island | Catalina Island | San Clemente Island | Guadalupe Island | Avg | Avg, excluding mainland |
| --- | --- | --- | --- | --- | --- | --- | --- | --- | --- |
| **Mainland** | – | 0.063<br>(-0.064, 0.19) | 0.038<br>(-0.062, 0.138) | 0.065<br>(-0.098, 0.227) | 0.024<br>(-0.055, 0.104) | 0.024<br>(-0.058, 0.107) | 0.053<br>(-0.056, 0.163) | 0.045 | – |
| **Santa Rosa Island** | 0.063<br>(-0.064, 0.19) | – | 0.015<br>(-0.026, 0.057) | 0.036<br>(-0.095, 0.167) | 0.029<br>(-0.035, 0.093) | 0.021<br>(-0.033, 0.075) | 0.059<br>(-0.048, 0.166) | 0.037 | 0.032 |
| **Santa Cruz Island** | 0.038<br>(-0.062, 0.138) | 0.015<br>(-0.026, 0.057) | – | 0.014<br>(-0.094, 0.121) | 0.016<br>(-0.03, 0.061) | 0.01<br>(-0.029, 0.049) | 0.045<br>(-0.046, 0.136) | 0.023 | 0.02 |
| **Anacapa Island** | 0.065<br>(-0.098, 0.227) | 0.036<br>(-0.095, 0.167) | 0.014<br>(-0.094, 0.121) | – | 0.03<br>(-0.093, 0.152) | 0.021<br>(-0.09, 0.132) | 0.075<br>(-0.098, 0.247) | 0.04 | 0.035 |
| **Catalina Island** | 0.024<br>(-0.055, 0.104) | 0.029<br>(-0.035, 0.093) | 0.016<br>(-0.03, 0.061) | 0.03<br>(-0.093, 0.152) | – | 0.009<br>(-0.032, 0.051) | 0.043<br>(-0.042, 0.127) | 0.025 | 0.025 |
| **San Clemente Island** | 0.024<br>(-0.058, 0.107) | 0.021<br>(-0.033, 0.075) | 0.01<br>(-0.029, 0.049) | 0.021<br>(-0.09, 0.132) | 0.009<br>(-0.032, 0.051) | – | 0.041<br>(-0.043, 0.125) | 0.021 | 0.02 |
| **Guadalupe Island** | 0.053<br>(-0.056, 0.163) | 0.059<br>(-0.048, 0.166) | 0.045<br>(-0.046, 0.136) | 0.075<br>(-0.098, 0.247) | 0.043<br>(-0.042, 0.127) | 0.041<br>(-0.043, 0.125) | – | 0.053 | 0.052 |

**Table 2.** Partitioning of genetic variance in Channel Island samples among climate alone (controlling for geography) and geography alone (latitude and longitude, while controlling for climate), as well as the joint influence of climate and geography that is confounded and cannot be disentangled into separate effects. First column includes the total amount of variation explained, and second column is as a proportion of variance explained by the full model (0.038). *P*-values were calculated from 99 permutations, except for the joint influence, which is calculated by subtracting the variance explained by climate and geography alone from the variance explained by the full model, and is not testable using permutations.

| Model | Adjusted R<sup>2</sup> | Proportion of explainable variance | *P* |
| --- | --- | --- | --- |
| full model (climate + geography) | 0.038 | 1 | 0.01 ** |
| Climate alone | 0.023 | 0.616 | 0.01 ** |
| Geography alone | 0.009 | 0.225 | 0.01 ** |
| Joint influence of climate and geography | 0.006 | 0.159 |  |

We annotated candidate SNPs using SnpEff (version 5.2c, Cingolani et al., 2012), which can predict the coding effect of variants. First we built a custom database using the *Q. lobata* genome, then used SnpSift to filter results to just the candidate genes. We also used the bedtools closest tool (version 2.31.0, Quinlan & Hall, 2010) and the annotation file for the *Q. lobata* genome to extract the ten sequence ontology features (e.g. gene, mRNA, exon) closest to each SNP and their upstream or downstream distance relative to the SNP (-D option). We then calculated the number of SNPs occurring within a gene or mRNA transcript (with distance = 0).

### Predicting climate suitability

Gradient Forest (GF, version 0.1-32, Ellis et al., 2012) was used to identify nonlinear relationships between SNPs and environmental factors, and to predict the spatial change in allelic composition, or genomic turnover, across the range of Island Oak. We ran GF on three sets of the imputed SNP data: 1) all SNPs to predict background genomic turnover for the Channel Islands, 2) candidate SNPs identified from the redundancy analysis for the Channel Islands to predict putatively adaptive turnover for the Channel Islands, and 3) candidate SNPs identified from Guadalupe Island to predict genomic turnover within Guadalupe Island and across the entire species range. Geographic location cannot be included as a predictor in GF analysis, so principal coordinates of neighbor matrix (PCNM) axes were calculated as a proxy for geography across a wide range of scales (Borcard & Legendre, 2002) using the `pcnm` function in the vegan package (version 2.6-4, Oksanen et al., 2019). To calculate the PCNM axes, we set the truncation distance separately for datasets because they encompassed different areas. For the two analyses on the Channel Islands, a truncation distance of 150 km was used in calculating PCNM axes to maintain connections between populations on the Northern and Southern Channel Islands and avoid statistical artifacts resulting from our clustered sampling design. As recommended by Fitzpatrick & Keller (2015), we included the first half of the axes with positive eigenvectors (denoting a positive spatial correlation) in the analysis, in this case 5 PCNM axes. For Guadalupe, we used a truncation distance of 430 km to account for the larger geographic area and retained 3 PCNM axes. We also included the same climate variables that were used in the redundancy analysis as explanatory variables. The `gradientForest` command was run with the following parameters: number of trees = 500, correlation threshold = 0.5, and max level = log_2_(0.368 × number of samples/2), following Fitzpatrick & Keller (2015).

We predicted the genetic composition and turnover across the range using the relationships between climate and allele frequency (in this case, a value of 0, 1, or 2 indicating the number of copies of a variant present in an individual tree) to scale climate variables by their genetic importance in contributing to genetic variation and predict genomic composition at non-sampled locations based on climates. The first three predicted axes were mapped to RGB color values which were combined for each cell and mapped, as in Fitzpatrick & Keller (2015). We did this color assignment for the three sets of SNPs. For the Channel Islands, we also mapped the difference in genetic turnover between the genome-wide SNP dataset and the candidate SNPs using the Procrustes residuals (Fitzpatrick & Keller, 2015; Peres-Neto & Jackson, 2001).

Using the Gradient Forest predictions of genomic turnover, we calculated the difference between historic and future climates using the genomic-scaled climate variables, or the genomic offset. Like similar studies, this method makes the assumption that populations are locally adapted to their historic climate conditions. Genomic offset was calculated for different pairings of seed sources (or populations) adapted to its historic climate, and planting sites experiencing future climate, with climate variables defined for each 30-second by 30-second grid cell from Bioclim. Because offset is predicted from gridded climate data, these “populations” are not defined biologically, but are groups of trees within the same climate grid cell that are likely to have similar adaptive genetic variation based on our gradient forest model. We calculated the genomic offset for each population under our three conservation scenarios: 1) the *status quo* approach to conservation, in which populations are maintained at their present locations, in which offset is calculated for historic and future climates at their current locality; 2) ecosystem preservation, in which non-local, pre-adapted genotypes are transplanted into a site, and offset is calculated between the future climate at one of our sample sites and the historic climate of all sampled populations; and 3) species preservation, in which genotypes are conserved by transplanting them to optimal sites, and offset is calculated for the historic climate of each population and all future climates on the islands where they could be planted. These are similar to the local, reverse, and forward offset (respectively) calculated in Gougherty et al. (2021), but here we explicitly test different conservation scenarios for each population. For scenario 3, we calculated offset on two sets of locations: only the sites where trees were sampled in this study, and all possible sites on the islands.

We also calculated a relative climate suitability metric by dividing genomic offset for each possible pair of seed sources and planting sites by the maximum offset calculated in the model, so that a value of 1 represents climate that is identical to climate of origin, and 0 represents the least suitable site. To visualize the predicted optimal scenarios for assisted gene flow, we selected the pairs of seed source and planting sites with the greatest climate suitability for each site (ecosystem preservation) and population (species preservation).

## Results

### Genetic Structure

*F*_ST_ values among island pairs and mainland *Q. chrysolepis* ranged from near 0 to 0.075 (Table 1), with lower values among the groups containing genetically *Q. chrysolepis* or hybrid individuals (Figure S1). Within the island populations, the population from Guadalupe Island was the most divergent from other populations (average pairwise *F*_ST_ = 0.052), followed by Anacapa and Santa Rosa (average pairwise *F*_ST_ = 0.035 and 0.032, respectively). In fact, the average divergence between Guadalupe Island populations and the Channel Islands populations was similar to that of the divergence between mainland *Q. chrysolepis* and the island populations primarily consisting of *Q. tomentella.* PCA and ADMIXTURE plots also supported divergence of Guadalupe populations from other *Q. tomentella* samples in California (Figures 2 and 3).

**Figure 2.**
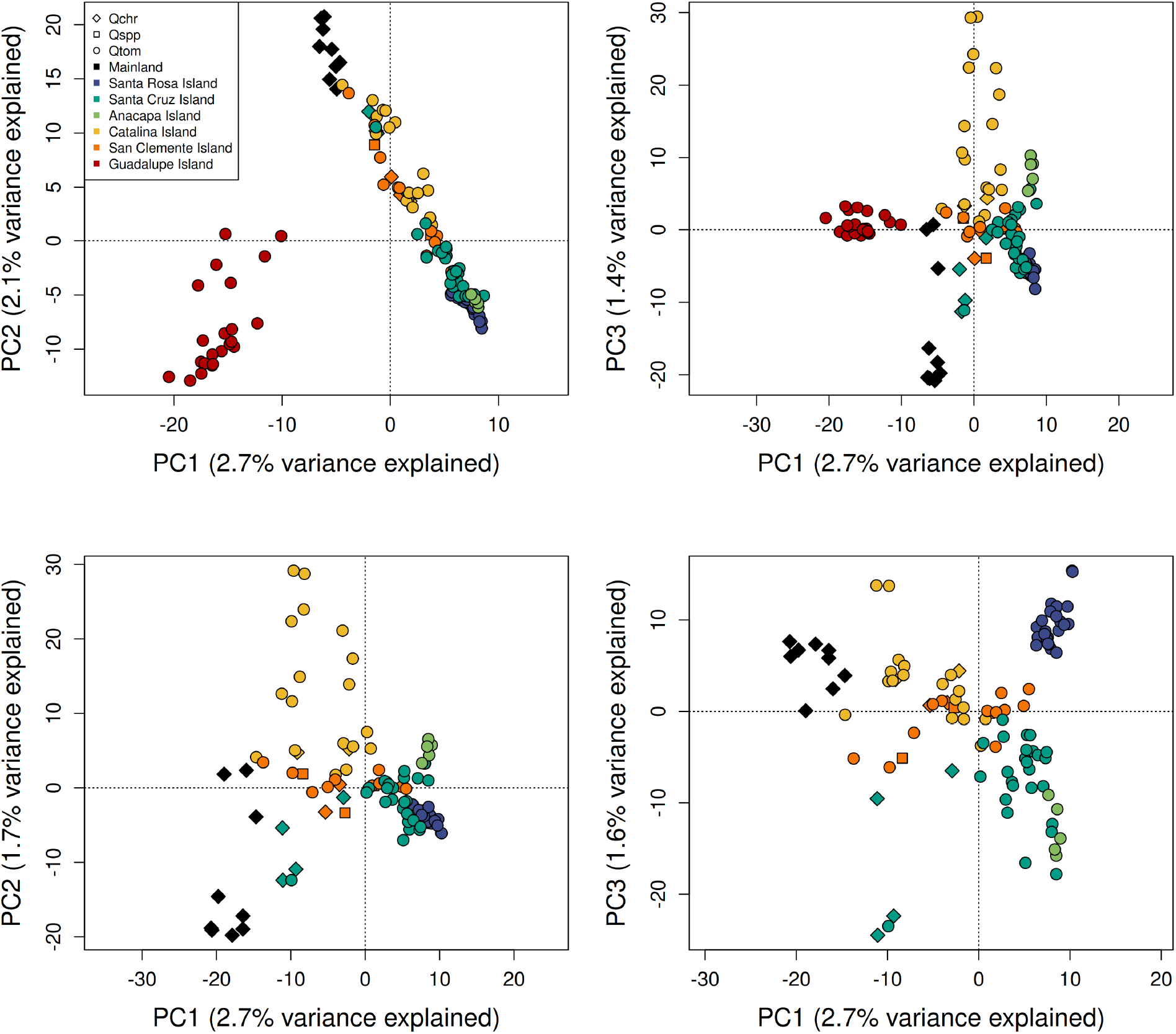
PCAs of genetic variation across SNPs for axes 1-3 for all samples (top) and with Guadalupe Island excluded (bottom). Color of points indicates their collection location, and shape indicates the species as identified in the field.

**Figure 3.**
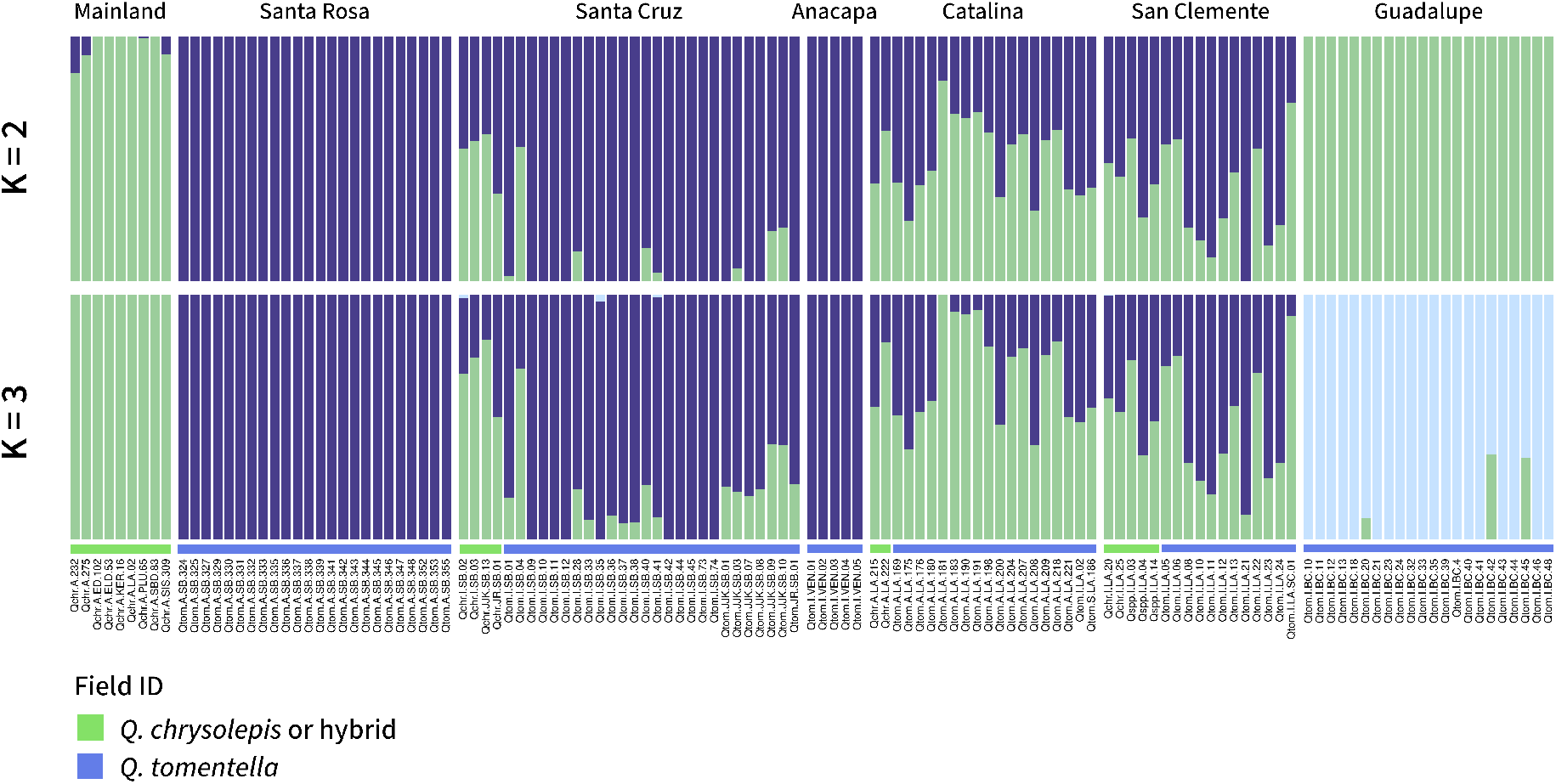
ADMIXTURE results when the number of ancestral groups (K) was set to values 2 and 3. K=1 (a single ancestral population) had the lowest cros-validation error (Figure S2), but here we show larger K values to visualize hierarchical structure. Each color indicates a group, and each bar shows the ancestry proportions of each group for an individual tree. Colored bars at the bottom indicate the field species ID. Individuals with admixed ancestry were often, but not always, identified as *Q. chrysolepis* or hybrids. See Figure S3 for plots for K = 4 and 5.

We found evidence of widespread introgression between the two species. Mainland *Q. chrysolepis* samples clustered separately from the island individuals in the PCA, but island individuals that were identified in the field as *Q. chrysolepis* or possible hybrids primarily cluster within the island samples rather than with mainland *Q. chrysolepis* (Figure 2). ADMIXTURE plots show that *Q. chrysolepis* or hybrid individuals on the Northern Channel Islands have mixed ancestry, with portions from mainland *Q. chrysolepis* and from the northern *Q. tomentella* groups (Figure 3). Trees from the Southern Channel Islands had more *Q. chrysolepis* ancestry than those from the Northern Channel Islands (Figure 3), with nearly all individuals having some level of admixture, even those morphologically consistent with *Q. tomentella*.

ADMIXTURE results for the full dataset, for the dataset with Guadalupe sample excluded, and for the dataset with only the Northern Channel Islands (Santa Cruz and Santa Rosa) all had the lowest cross-validation error for K=1 (Figure S2), meaning that in the best model, all individuals come from a single ancestral population (although for the set of all samples, CV error only increased slightly for K=2 and 3). We plotted admixture proportions for multiple K values to visualize hierarchical genetic structure (Meirmans, 2015) (Figure 3). The plot for K = 2 separates mainland *Q. chrysolepis* and Guadalupe Island as one ancestry group and the Northern Channel Islands as another, with admixture between the two groups occurring in hybrid individuals and in the Southern Channel Islands. For K = 3, Guadalupe Island forms a separate ancestry group. Across multiple K values, trees from the Southern Channel Islands were more admixed, with ancestry proportions from both mainland *Q. chrysolepis* and the Northern Channel Island populations (Figure 3, Figure S3). Running ADMIXTURE on a subset dataset including only two of the Northern Channel Islands, Santa Rosa and Santa Cruz, revealed fine-scale genetic structure. For example, one tree on the southwestern ridge of Santa Cruz shared some ancestry with Santa Rosa samples, suggesting that gene flow has occurred from Santa Rosa to Santa Cruz populations (Figure S3 and S4), although this result should be interpreted with caution as it includes only one individual. We found a weak effect of isolation-by-distance, with the most closely related individuals occurring on the same island, but only slight increases of genetic distance with geographic distance for inter-island pairs of trees (logistic model P <2.2e-16, adjusted R^2^ = 0.3421) (Figure S5).

### Climate-associated genetic variation

Multiple redundancy analysis models were tested in order to partition genetic variance into proportions explained by climate, geography, their joint influence, and each factor alone (controlling for the effects of the other factor). All models significantly explained genetic variation (*P* = 0.01 for all), indicating that both geography and climate shape genetic variation (Table 2). However, climate explained a greater proportion of variance than geography: in the full model, geography and climate together explained 3.8% of the genetic variation; with climate alone explaining 2.3%, geography alone explaining 0.9%, and their joint influence explaining 0.6%. As a proportion of total explainable variance, climate alone explained 61.6%, geography alone explained 22.5%, and their joint influence explained 15.9%.

The redundancy analysis identified 560 candidate climate-associated SNPs that were outliers in the Channel Islands dataset and 346 in the Guadalupe dataset (Figure S6). For each candidate SNP, we determined which climate variable was most strongly correlated. For the Channel Islands, there were 174 SNPs correlated most strongly with precipitation of the coldest quarter, 155 with elevation, 88 with maximum temperature of the warmest month, 73 with precipitation of the warmest quarter, 47 with precipitation seasonality, and 23 with minimum temperature of the coldest month. For Guadalupe, 204 SNPs were most strongly correlated with minimum temperature of the coldest month, 129 with maximum temperature of the warmest month, 9 with elevation, 2 with precipitation seasonality, and 2 with precipitation of the coldest quarter. For the Channel Islands, 61% (343) of candidate genes were located within a gene and 59% (331) were within an mRNA. SnpEff can identify multiple effects for each SNP, such as multiple nearby genes. SnpEff identified 13% of SNPs (73) as having at least one moderate effect (non-disruptive but potentially affecting protein function, such as a missense variant), 10% (59) as having a low effect (unlikely to change protein function, such as a synonymous variant), all 562 SNPs having a “modifier” impact (upstream or downstream of a gene), and no SNPs having a “high” impact (a major change to a protein-coding regions potentially causing loss of function, such as an added stop codon). For the candidate SNPs on Guadalupe, 49% (169) were located within a gene and 45% (155) were located within an mRNA, a slightly lower proportion than on the Channel Islands. Three candidate SNPs out of 348 (8.6%) had a high impact variant, 9.8% (34) had a moderate impact, 8.3% (29) had a low impact, and all SNPs had a modifier impact. Few of the Channel Islands candidate SNPs were private to one island (5% of reference and 10% of alternate alleles), supporting our findings that gene flow likely occurs among the five islands, resulting in relatively low population structure. The RDA loadings, climate correlations, and annotations for each candidate SNP are reported in Supplementary Table S2.

When Gradient Forest was used to analyze genome-wide SNPs in the Channel Islands samples, the five PCNM axes describing geographic distance had greater importance values than the climate variables (Figure S7), suggesting a spatial influence on genetic variation. However, for the candidate SNPs, important variables included both PCNM axes and environmental variables (with PCNM3, minimum temperature of the coldest month, and elevation having the highest R^2^ weighted importance values for the Channel Islands; and maximum temperature of the warmest month, precipitation of the coldest quarter, and elevation having the highest values for Guadalupe) (Figure S5). As expected, overall importance values were also higher when considering only the candidate SNPs, indicating stronger associations. The relationship between allelic variation and climate, or genomic turnover, was mapped to visualize predicted spatial patterns of climate-associated genetic variation for the Channel Islands (Figure 4) and for Guadalupe (Figure 5). The Southern Channel Islands were more similar to each other, and the Northern Channel Islands showed variation across an east-west gradient associated with precipitation seasonality (BIO15) and precipitation of the warmest quarter (BIO18). When predicting turnover using candidate SNPs identified from Guadalupe, there was more turnover within Guadalupe than across the Channel Islands (Figure 5), consistent with the genetic differentiation of Island Oaks on Guadalupe from the other islands, as well as putatively adaptive genetic differentiation within the island.

**Figure 4.**
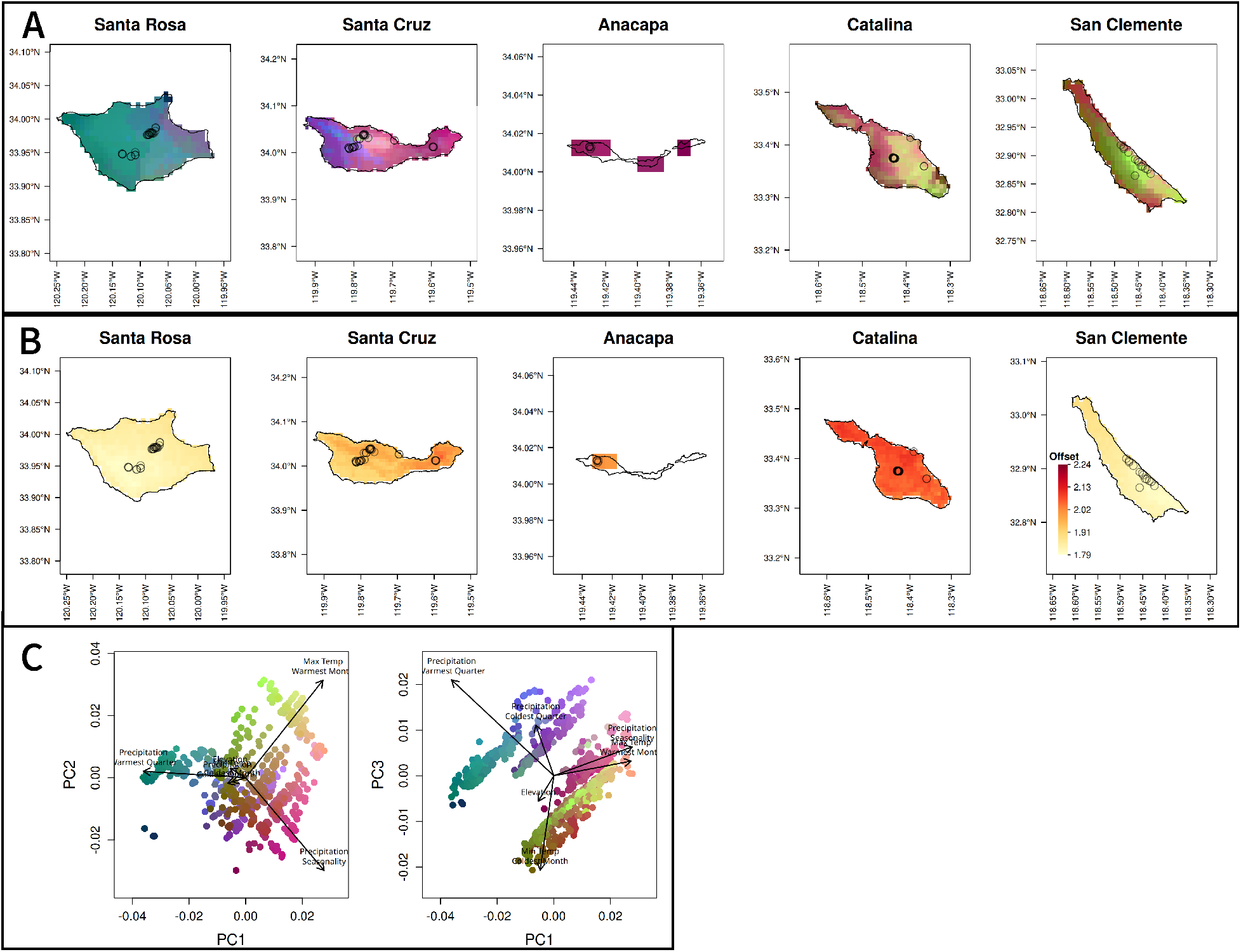
A) Maps of genetic turnover predicted by gradient forest for candidate SNPs on each of the Channel Islands. Climate variables are scaled by their importance in predicting genetic variation in the gradient forest model and their relationship with allelic turnover as shown in Figure S8, then the genomic composition of a cell is predicted based on its climate. The first three PCs of the genomic composition are mapped to RGB values, so regions with similar colors are expected to have similar genomic composition. Points indicate sample locations. B) Map of the genetic offset, or predicted difference in the current genetic composition and that which would be ideal under future climate conditions. Darker red indicates a greater likelihood of future maladaptation. Results are shown for the CNRM 4.5 scenario for the year 2050 (offsets under other models and scenarios are in Figure S13). C) PCA of scaled climate values for each cell, with colors matching each grid cell in A.

**Figure 5.**
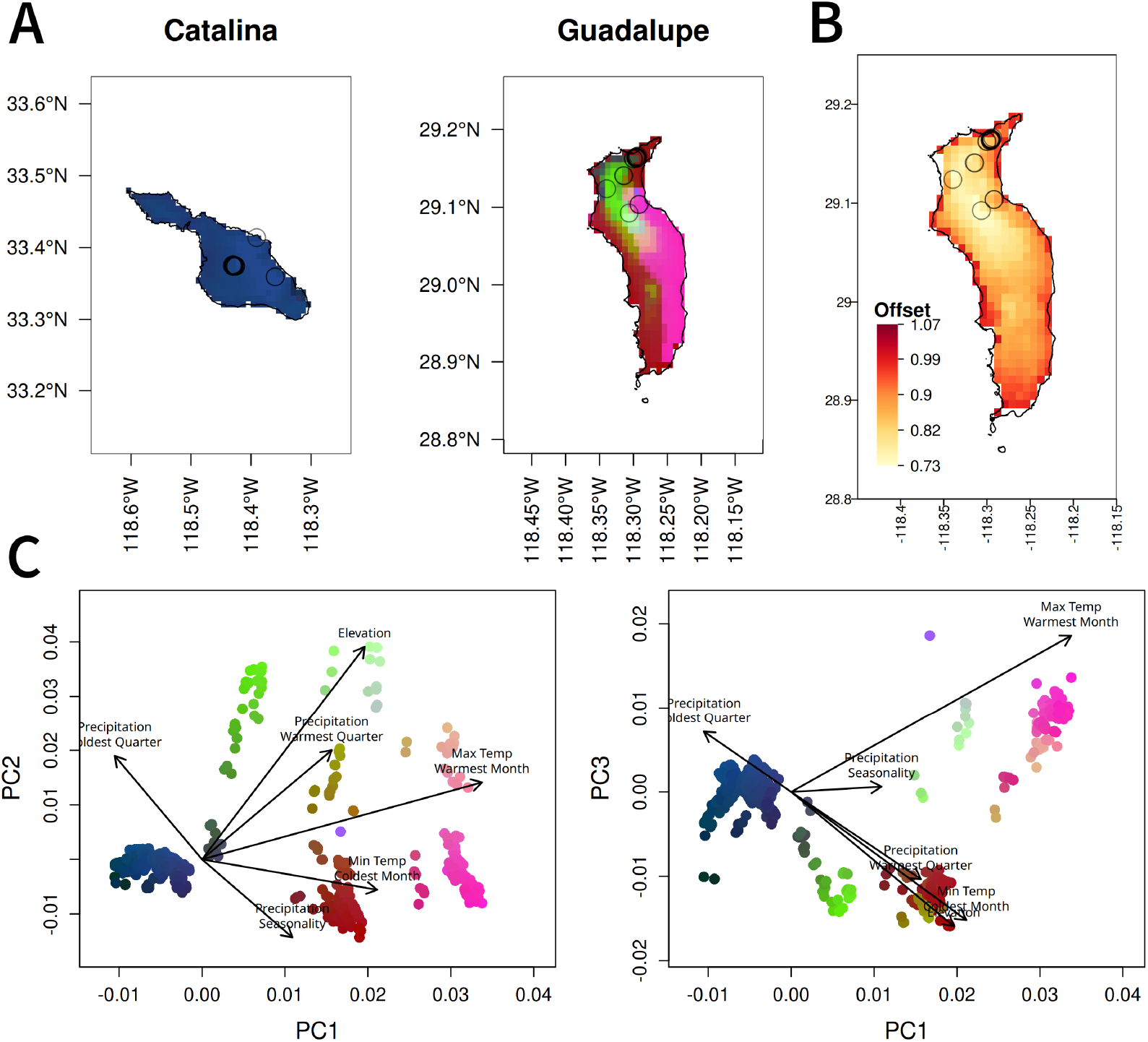
A) Maps of genetic turnover predicted by gradient forest for candidate SNPs on Guadalupe Island. Catalina Island is included as a comparison, but all other Channel Islands were similar, clustering in the dark blue/purple regions with lower BIO5 (max temperature of the warmest month). Climate variables are scaled by their importance in predicting genetic variation in the gradient forest model, then the genomic composition of a cell is predicted based on its climate. The first three PCs of the genomic composition are mapped to RGB values, so regions with similar colors are expected to have similar genomic composition. Points indicate sample locations. B) Map of the genetic offset, or predicted difference in the current genetic composition and that which would be ideal under future climate conditions. Darker red indicates a greater likelihood of future maladaptation. Results are shown for the CNRM 4.5 scenario for the year 2050; offsets under other models and scenarios are shown in Figure S14. Colors only indicate offset within Guadalupe and are not comparable to offset in other islands in Figure 4. C) PCA of scaled climate values for each cell, with colors matching each grid cell in A.

#### Genomic offsets and conservation strategies

Catalina Island has the greatest genomic offset under 2050 climate (CNRM model, RCP 4.5), followed by Anacapa and eastern Santa Cruz (Figure 4). These results were consistent across the two climate models and the two RCP scenarios tested (Figure S13). Within Guadalupe, populations closer to the coast have a higher offset (Figure 5). Offset values calculated for the two datasets (Channel Islands and Guadalupe) are not comparable to each other, so we cannot determine whether the risk for Guadalupe is higher or lower than for the Channel Islands.

Both conservation strategies implementing assisted gene flow reduced the genomic offset compared to the *status quo* (Figure 6). For most sampled sites, the genomic offset was reduced most with the species preservation approach compared to the ecosystem preservation approach, meaning that transplanting populations to a location with optimal climate reduced the chance of maladaptation for that population more than introducing non-local genotypes into the population. Offsets were slightly reduced when all parts of the islands were included as possible planting sites, rather than only the sites with sampled oak populations (Figure 6), indicating that some future optimal sites may be outside of the range sampled in this study. Optimal planting sites were similar for all populations and included sites that will be coolest under future climates: Santa Rosa followed by San Clemente (although under CNRM 8.5, San Clemente has higher suitability than Santa Rosa – Supplemental Table S1). Because Santa Rosa populations are already located in the part of the species range that will have future climate conditions that are most similar to current conditions for the species, they benefit more from the ecosystem preservation approach than the species preservation approach, unlike other populations.

**Figure 6.**
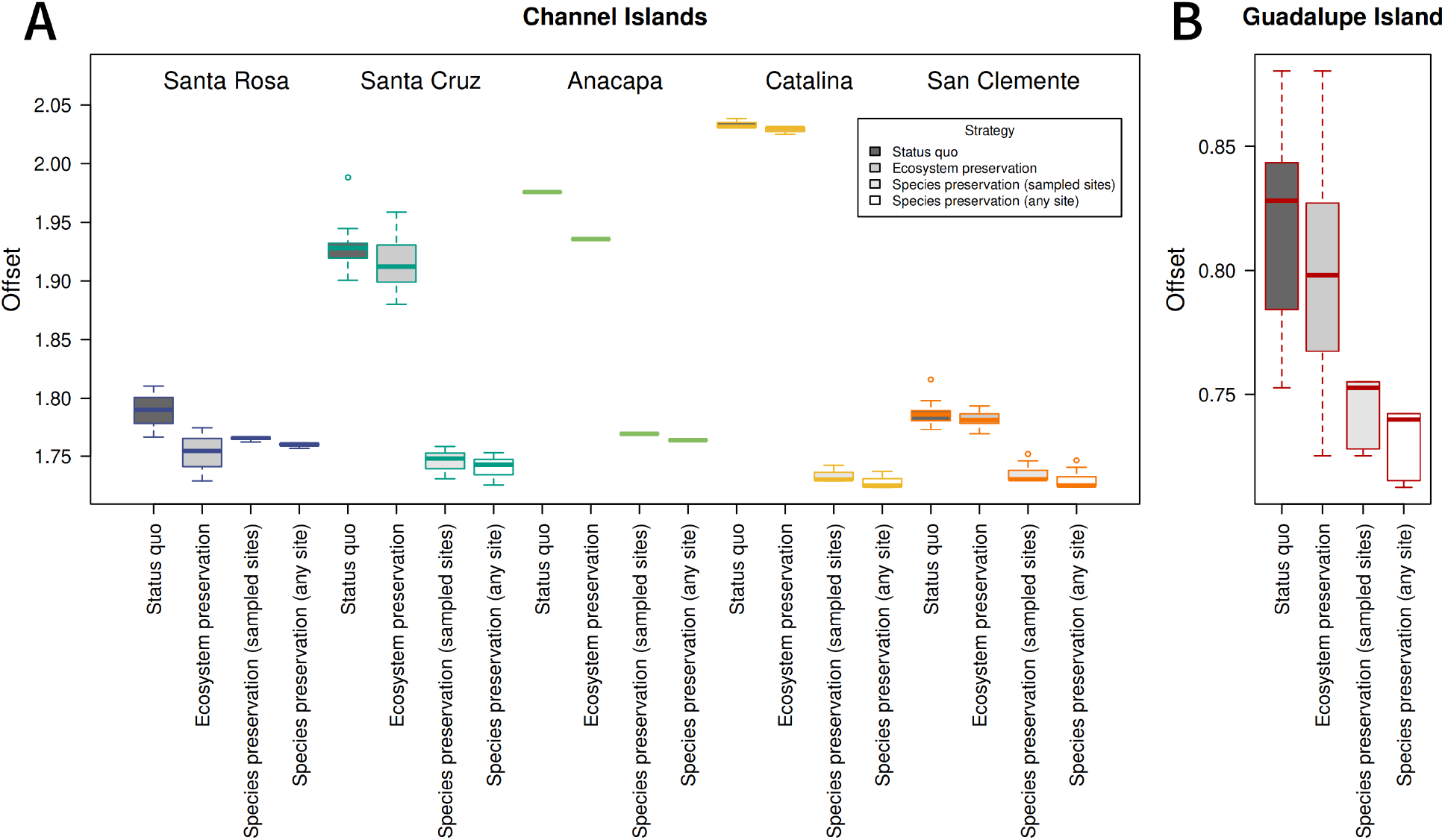
Comparison of genomic offsets under four management scenarios for the Channel Islands (A) and Guadalupe Island (B). Higher offsets represent greater maladaptation to future climate. Offsets were calculated for each grid cell that included sampled trees, referred to as “populations,” and are sorted by island. The *status quo* strategy indicates offsets for each population at its current site. In the ecosystem preservation strategy, the offset is given for a scenario in which the site is planted with the non-local population predicted to be best-adapted to its 2050 conditions. In the species preservation strategy, the offset is given for the population on the island if it were transplanted to another site with the most optimal 2050 conditions, either when including the sampled sites or all sites on the islands. For A and B, offset values are calculated separately, with transplantation only within the Channel Islands or Guadalupe Island, and the offsets are not comparable between the two plots.

## Discussion

In this study, we use the combined evidence from landscape genomics and genome-wide genetic structure to compare the potential benefits of several conservation strategies for Island Oak and present our results as a case study in using genomic data to inform the conservation of rare species. Our genomic analyses reveal great potential for using seed sources to manage, restore, and expand Island Oak ecosystems and to preserve the species, despite the fact that currently all localities are predicted to be maladapted under future climates. These findings point to the importance of using assisted gene flow for populations on many of the islands to maximize their potential for survival under future climate environments.

### Genetic structure

Information about a species’ genetic structure can be used to inform seed sourcing decisions, balancing the goals of climate adaptation and preserving historic genetic structure and gene flow patterns. Our results show genetic differentiation across islands, but *F*_ST_ values are relatively low for an island species (Gugger et al., 2018; Villa-Machío et al., 2020, but see Di Santo et al., 2022), indicating weak population structure among islands (Table 1). The ADMIXTURE cross-validation error was minimized with one ancestral population, supporting the finding of low population structure. Guadalupe Island populations, which are at a geographic distance from the Channel Islands populations that make pollen transfer extremely unlikely, are also the most genetically divergent from other island populations (average pairwise *F*_ST_ = 0.052, Table 1). On the Channel Islands, Santa Rosa populations appear to be the most genetically distinct, while other islands are less differentiated from each other and show signs of admixture (Table 1, Figure 2 and 3), similar to the pattern found previously in Song Sparrows on the Northern Channel Islands (Gamboa et al., 2022). The patterns of genetic structure found here may result from asymmetric gene flow shaped by typical wind patterns during spring flowering (Figure 1A). In the Channel Islands, winds are primarily from the northwest, so trees on Santa Rosa Island are unlikely to receive pollen from other islands, except under rare spring Santa Ana wind conditions, which normally occur in Fall when oaks are not producing pollen. However, Santa Cruz and Anacapa may receive pollen from Santa Rosa, and San Clemente and Catalina may receive pollen from all three northern islands, explaining the increased admixture observed in these individuals (Figure 3 and S3). These results suggest that the Northern Channel Islands are unlikely to receive alleles from the warmer Southern islands, reducing the pool of available alleles that could be adaptive under warmer future climates. Similar to our results, previously analyzed microsatellite data found isolation of Guadalupe and greater admixture in the Catalina and San Clemente populations (Ashley et al., 2018), but our ADMIXTURE results show more distinct ancestries between Santa Rosa and the islands to its east, possibly because the whole-genome dataset captured additional genetic variation resulting from adaptation to differing climates across islands. Patterns of differentiation among islands were also similar to our results, but are not directly comparable because different markers and differentiation statistics were used (Ashley et al., 2018).

Introgression with *Q. chrysolepis* appears to be widespread, except on Santa Rosa and Anacapa, and is likely to be another important factor shaping genetic structure in Island Oak. Admixture between the two species is particularly evident in the Southern Channel Islands, but also present within some individuals in the Northern Channel Islands. Island trees that were identified as *Q. chrysolepis* based on field morphology clustered more closely with *Q. tomentella* individuals than with the mainland *Q. chrysolepis* individuals, suggesting a long history of introgression between the two species on these islands (Figure 2). Our genetic results suggest that *Q. chrysolepis* may not be present on Santa Rosa or Anacapa, where they have been previously reported (Rosatti & Tucker, 2014). The isolated Guadalupe Island population, however, appears to be somewhat genetically similar to the mainland *Q. chrysolepis;* having similar values along PC1 (Figure 2) and forming the same ancestry group for K=2 (Figure 3).

Previous work has hinted at a complex evolutionary history in these two species. Ortego et al. (2018) suggested that *Q. chrysolepis* is a non-monophyletic species that split into a northern and southern lineage, following which *Q. tomentella* split from the southern lineage of *Q. chrysolepis* in the late Pliocene or early Pleistocene, and that the two species hybridized when they were sympatric on the mainland. Colonization of the Channel Islands may have occurred when California shifted toward a more Mediterranean climate and *Q. tomentella* became restricted to the coast (Axelrod, 1967; Muller, 1965). Additionally, it is possible that the Guadalupe population has experienced introgression with *Q. cedrocensis*, which we were unable to sample but which is present on Cedros Island to the southeast of Guadalupe Island (Figure 1A), as well as on the mainland in southern San Diego County and northern Baja California. In the future, demographic modeling that includes the Guadalupe Island populations could be used to clarify the timing of divergence between Guadalupe and the other islands, and the timing of introgression with *Q. chrysolepis* or other oak species in section *Protobalanus.* Hybridization is sometimes considered a conservation concern, but here it seems to be a part of a long evolutionary history of introgression between the two species, as is common in oaks (Kremer & Hipp, 2020; O’Donnell et al., 2021), and could be the source of novel genetic variation and ongoing selection. Experiments could compare the fitness of individuals with different levels of hybrid ancestry in different conditions to test whether introgression could be adaptive in some environments.

### Climate-associated genetic variation

To test whether genetic variance in Island Oak populations was better explained by climate (consistent with local adaptation) or geography (consistent with neutral differentiation), we used a redundancy analysis to partition genetic variation into proportion explained by climate and geography separately. Climate accounted for the majority (62%) of the explainable genetic variation among the Channel Island populations when controlling for the effects of geography (Table 2). Thus, natural selection is likely affecting the evolution of these island populations more than neutral differences resulting from genetic drift and limited gene flow within or among islands. There is a significant effect of geography as well, although overall differentiation among islands is relatively low (Table 1) and a pattern of isolation by distance primarily shows that the most genetically similar individuals occur on the same island, and that genetic distance increases only weakly with distance among islands (Figure S5). Climate variables that were important in the redundancy analysis and Gradient Forest analysis varied, suggesting that elevation, temperature, and precipitation variables are important in local adaptation, without a strong influence of any one factor, unlike some tree species for which genomic turnover is most associated with precipitation variables (Gugger et al., 2018; Martins et al., 2018). The moderating impact of ocean temperatures and summer fog providing additional moisture through fog drip (Fischer et al., 2009, 2016; Williams et al., 2008; Woolsey et al., 2018) on the islands may make these variables indistinguishable.

In addition to the effect that climate has in explaining genetic variation across islands, climate-associated genomic turnover indicates genetic variation within islands consistent with local adaptation to climate gradients. On Santa Cruz Island, climate-associated genetic variation differed across an east/west gradient (Figure 4A). The full set of SNPs and the climate-associated candidate SNPs produced different predictions for the rate of genomic change across climate gradients (genomic turnover) on eastern Santa Cruz Island (Figure S11), suggesting that gene flow occurs between eastern and western Santa Cruz populations, but that selection results in differentiation of climate-associated alleles between the regions. Similar patterns of genetic differentiation on Santa Cruz have been found previously in Island Scrub-jays (Cheek et al., 2022; Langin et al., 2015) and in lizards (Trumbo et al., 2021). Local adaptation could occur as a result of the precipitation gradient in the Northern Channel Islands, in which eastern regions have less summer precipitation and greater precipitation seasonality. Taken together, the larger explanatory effect of climate on genetic differentiation and the patterns of putatively adaptive genetic variation across the landscape suggest that local adaptation may be possible at small scales and in the presence of gene flow, even within relatively small island populations, as found previously (Cheek et al., 2022; Gamboa et al., 2022; Gugger et al., 2018; Hamilton et al., 2017; Langin et al., 2015). In contrast, populations on Catalina and San Clemente show less turnover of adaptive alleles within islands. Candidate SNPs from Guadalupe showed very little variation or turnover on the Channel Islands (Figure 5A), suggesting that local adaptation among the Guadalupe populations results from variation in SNPs that are less variable across the Channel Islands. While reciprocal transplants would be needed to definitively identify local adaptation, either among or within islands, the pattern of genomic turnover within both Guadalupe and Santa Cruz islands suggests fine-scale local adaptation, consistent with predictions of local adaptation for forest trees (Sork, 2016). These results indicate that the assumption of local adaptation used in landscape genomics is reasonable and that assisted gene flow strategies using existing genetic variation are viable for this system.

### Genetic offset and maladaptation

Genetic offset is a measure of vulnerability to climate change, based on associations between genomic structure and future climates. Our results predict that, without intervention, populations on Catalina will be most maladapted to future climates in comparison to the other Channel Islands populations, followed by eastern Santa Cruz populations (Figure 4C). If gene flow naturally occurs from the Northern to Southern Channel Islands, as suggested by neutral genetic structure, both Catalina and San Clemente may currently receive northern alleles that are maladaptive at hotter sites and could become more maladaptive as climate change progresses. Catalina is projected to experience a greater increase in temperature than San Clemente (Figure 1B), resulting in a larger genomic offset. Within Guadalupe, populations at lower elevations closer to the coast are predicted to have a higher genomic offset. While all populations are likely to experience some degree of maladaptation to future climates, those with a greater genomic offset could be prioritized in conservation efforts. We cannot compare offsets among Guadalupe and the other islands, so this analysis is most useful in identifying localities within the island that are most at risk. Lotterhos (2024) cautions against interpreting genomic offset as a metric of maladaptation when calculated from genotype-environment associations because it is possible that future climates could be more beneficial than historic climates. However, given the absence of Island Oak from warmer and drier microclimates on the islands and its status as a paleoendemic, we believe future climates are unlikely to be beneficial for this species. Common garden or growth chamber experiments comparing the response of Island Oaks to warmer or cooler temperatures should be performed in future research to test this assumption.

### Evaluation of conservation strategies

For current and future restoration projects taking place on several islands, landscape genomic findings can inform decisions about what seed sources should be used to maximize success for plantings. However, different conservation strategies may be preferred depending on management priorities and policies, and conservation decisions must take those values into account. If the priority is to maintain “natural” patterns of genetic variation, seeds should only be introduced among populations that have historically experienced gene flow. However, to conserve existing Island Oak ecosystems and the species that depend on them, it may be necessary to introduce alleles that are putatively “pre-adapted” to future climates through assisted gene flow (Aitken & Bemmels, 2016; Aitken & Whitlock, 2013). By investigating genetic structure and determining climate suitability for all possible seed sources and planting sites, it is possible to determine the risks and benefits of different strategies. Here, we compare the predicted genomic offset and climate suitability under three different management strategies: (1) conservation of local oak populations without assisted gene flow (*status quo*) which may include the use of local acorns in restoration projects, (2) ecosystem preservation – conserving existing Island Oak ecosystems by introducing non-local populations that are likely to have the greatest fitness under future conditions, or (3) species preservation – conserving the species and its genetic variation in at least part of its native range by planting populations at sites with optimal future climate.

Since all populations are predicted to be maladapted to future conditions (Figure 4C and 5B), maintaining historic patterns of gene flow may not effectively conserve populations. Because future climates are projected to be far outside the current climate space occupied by island oak (Figure S12), there may not be existing genetic variation within the species that will be well-adapted to future climates. As a result, the risk of maladaptation is reduced most by preserving populations at sites that will have climate most similar to that of current Island Oak populations, as in the species preservation approach (Figure 6). Using the species preservation approach to prioritize the preservation of genetic diversity, the optimal planting sites for all populations are those with the coolest projected future climate – Santa Rosa Island, followed by San Clemente Island (Figure 7). These locations could be used to preserve genotypes from across the species range. However, this approach would focus on conserving Island Oak in only a portion of its native range, potentially neglecting Island Oak ecosystems in regions most affected by climate change. While the ecosystem preservation approach does not reduce genomic offset as much as the species preservation approach, it is an improvement over the *status quo* (Figure 6), preserving existing oak groves by introducing genotypes that are likely to be best-adapted to their future conditions. The ideal outcome would conserve all existing island oak ecosystems; for this reason, we suggest prioritizing ecosystem-preservation, while also conserving genetic diversity as a bet-hedge to prevent extinction and loss of genetic variation.

**Figure 7.**
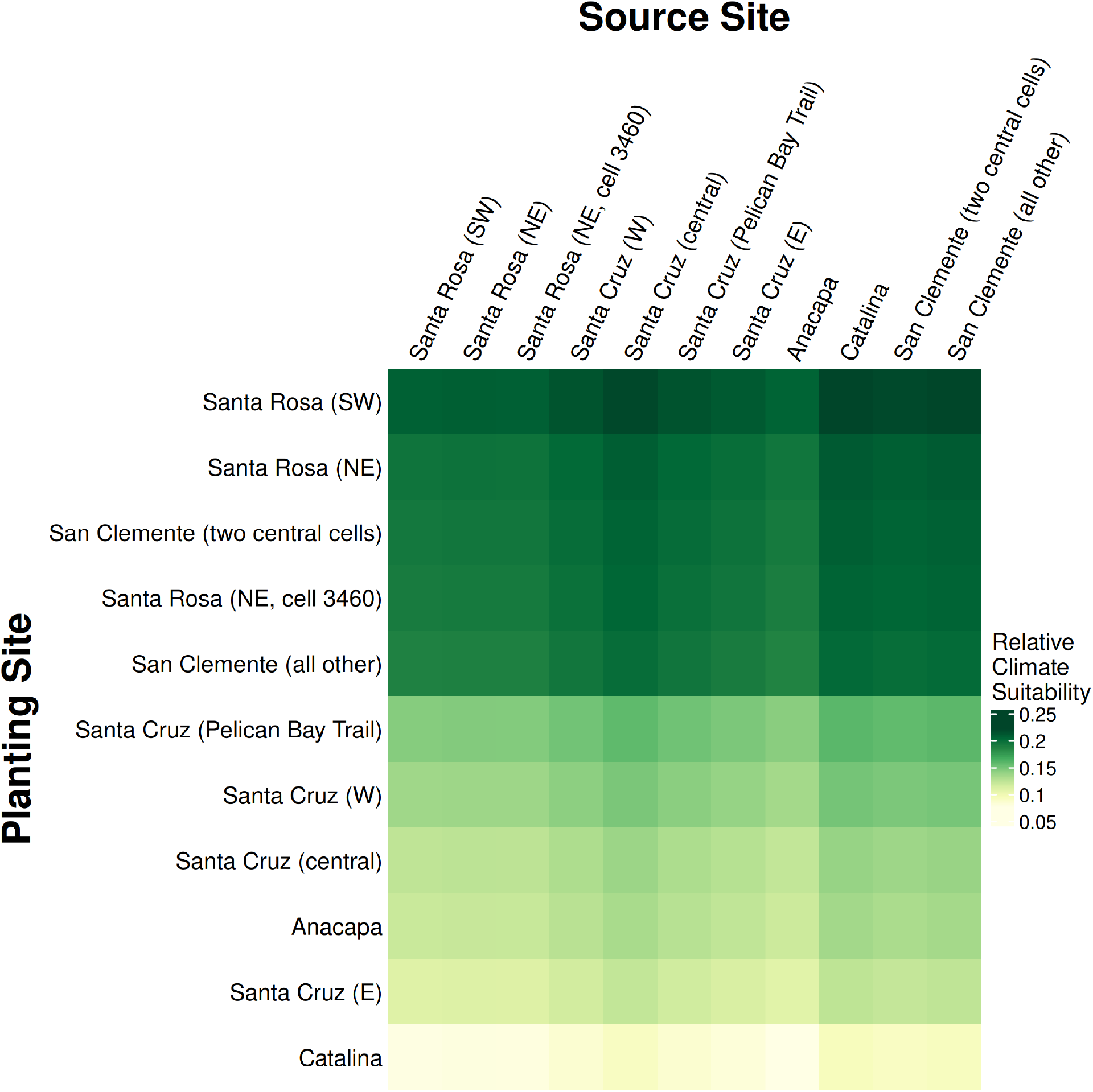
Climate suitability in 2050 when matching a sampled source site (columns) to a planting site (rows). Suitability was calculated for each grid cell where trees were collected for this study, and nearby grid cells with similar values were averaged for clarity. Planting sites (rows) are sorted by average suitability, with the optimal sites at the top. Columns are sorted geographically (roughly north to south and west to east). Under the ecosystem preservation approach, optimal source populations for restoring a given site can be selected by looking at climate suitability within a row. Under the species preservation approach, optimal planting sites for a given population can be selected by looking at climate suitability within a column.

For ecosystem-preservation to be successful, there must be overlap between the climate space of the species’ historic and future climates. Our finding that species preservation resulted in the lowest offset may differ across species by range size and the magnitude of climatic change predicted within the range, with ecosystem-preservation being more successful in species with wider climatic ranges (particularly latitudinal). If, like Island Oak, the species range will experience climates not matched by any historic climates in the species range, assisted gene flow would still result in a mismatch between climate and genomic variation – although the mismatch can likely be reduced in comparison to taking no action.

Climate suitability can be used to compare possible seed sources for a given planting site while taking into account other goals, such as conserving the species’ genetic structure. For example, the ideal seed sources for Santa Rosa’s future climate are populations on Catalina and San Clemente, the warmer Southern Channel Islands, which have experienced more admixture with *Q. chrysolepis*. While assisted gene flow may introduce beneficial alleles, it may also disrupt the natural genetic structure of populations on Santa Rosa island, which appear to be relatively isolated and have no evidence of admixture with *Q. chrysolepis*. However, the climate suitability of nearby central Santa Cruz Island populations is nearly as high as those for the Southern Channel Islands (Figure 7). As the two islands were connected as recently as 11,000 years ago (Kennett et al., 2008; Reeder-Myers et al., 2015) and have less extensive admixture with *Q. chrysolepis*, using central Santa Cruz as a seed source for Santa Rosa could represent a more moderate assisted gene flow approach that could still introduce beneficial alleles. Using climate suitability, managers can identify a subset of optimal seed sources, then select those that best maintain historical evolutionary structure for planting.

For Guadalupe Island, the optimal seed source was a region with high BIO6 (minimum temperature of the coldest month) and the optimal planting site was the westernmost part of the island sampled (Figure 5, Table S1). In preserving Guadalupe Island genotypes, predicted maladaptation was minimized when planting within the island, even when the cooler Channel Islands were included as possible planting sites (Table S1). The Guadalupe Island Oaks appear to be differentiated from the Channel Islands at both neutral and putatively adaptive SNPs, and are unlikely to experience contemporary gene flow from these populations. Because of their divergence, it is possible that hybridization between Guadalupe and Channel Islands populations could result in inbreeding depression. Therefore, the Guadalupe island oaks are probably best managed as a separate lineage at present. However, including Guadalupe populations in future studies, including common garden experiments, could help determine whether they contain unique genetic variation could be beneficial under hotter climates when introduced to the Channel Islands.

The ecosystem- and species-preservation approaches could be complementary and used in combination with other conservation strategies. Because Santa Rosa Island will have cooler future climates than other islands it could be an ideal planting site for other populations. Moreover, because the native Santa Rosa individuals are likely adapted to a cooler, more moderate climate, they could benefit from the introduction of genetic variation that is likely to be beneficial under climate change. In addition to maintaining genetic diversity within the native range, *ex situ* collections in botanic gardens and arboreta can be used to conserve the genetic diversity of species with seeds that cannot be stored through traditional seed banking, such as oaks (Rosenberger et al., 2022; Westwood et al., 2021). These living collections could be used as tissue or seed sources for restoration, or for replanting when a natural population is lost in events such as wildfire, landslide, or pathogen outbreaks. Lastly, assisted gene flow is not likely to be a viable strategy for populations that are predicted to be most maladapted to future temperatures and are also from the warmest part of the species range, such as Catalina Island. For such populations, we propose that supplementary ecological studies should be undertaken to identify seed sources (for example, specific maternal trees) that may be better adapted to future climate conditions. For species that are seriously threatened by future climates, additional options can be considered. An alternative ecosystem preservation approach would be to combine genomic resources with common gardens to identify warm-adapted genotypes with high fitness under future environments by associating fitness with genomic variation, using genome-informed breeding values (see Browne et al., 2019). If it is important to preserve the species *per se*, seedlings could be planted into e*x situ* locations, even outside of the current species range, such as botanical gardens with suitable climate environments (Rosenberger et al., 2022; Westwood et al., 2021).

### Limitations

Landscape genomics tools can be a powerful tool for identifying strategies to conserve species under changing climates; however, our recommendations depend on the assumption that populations are locally adapted to modern climate conditions. Because *Q. tomentella* is a paleoendemic that experienced range contraction and likely survives on the California Islands because of their more moderate climates, it is possible that populations are already maladapted to present climate conditions, as found for California populations of valley oak (Browne et al., 2019), meaning we underestimate maladaptation here. Furthermore, studies that have tested whether genomic offset can predict growth in a common garden have found mixed results (Fitzpatrick et al., 2021; Lind et al., 2023) so genetic and climate data may not be sufficient to predict optimal climates. Additionally, our predictions define optimal climate as the mean values over 30 years for a 30-second square grid cell. In reality, trees experience climatic variation across seasons, years, and microclimates (such as those created by the north-facing canyons that Island Oaks often inhabit) which are not represented by this climate data. Common garden experiments could validate our predictions of ideal population-climate matches prior to implementing assisted gene flow. For example, populations from across the species range could be planted at a restoration site and monitored over 5-10 years to determine which have the highest fitness at the site, and could be removed before they reach reproductive age if introducing non-local alleles is a concern. Santa Rosa Island would be an ideal location for such an experiment, as it is the optimal planting site for all populations, and local populations could potentially benefit from the introduction of warmer-adapted populations. Additional common gardens on warmer islands, such as Catalina, or under controlled growth chambers at different temperature regimes, could test which individuals will be maladapted to warmer future climate conditions. Nonetheless, our findings indicate that island populations are likely to be maladapted to future climates and that assisted gene flow can reduce the predicted maladaptation, so genomic-informed seed sourcing should be considered as a possible management strategy to conserve and restore Island Oak ecosystems.

### Conclusions

Assisted gene flow could reduce maladaptation of Island Oak to future climates relative to the restoring populations with local seeds. Conserving the genetic diversity of populations on islands that are projected to have less dramatic climate shifts, in this case, Santa Rosa and San Clemente, would conserve the species in parts of its native range. In addition, introducing genotypes tolerant to warmer climates into populations could help preserve existing, but vulnerable, island oak ecosystems. These two assisted gene flow strategies could be combined together: for example, by establishing restorations at ideal sites that will conserve range-wide genetic diversity while also introducing beneficial alleles to existing oak populations that would otherwise not be well-adapted to future climates. This study identified one locality of major concern (Catalina Island), which may require further ecological and genomic studies to identify seed sources within the island that may be better adapted to future climate conditions. This case study of Island Oak illustrates the distinction between merely preserving the species and preserving the ecosystem it shapes, which are important considerations for all conservation studies.

## Data Availability

Sequencing data will be uploaded to NCBI upon manuscript acceptance. Locality and climate data for collected samples are available as Table S1.

Analysis scripts will be made available on Github and archived at Zenodo upon manuscript acceptance. R markdown output files are included as supplementary information.

## Supporting information

Supplementary Rmarkdown Outputs

## Notes

### Competing Interest Statement

The authors have declared no competing interest.

### Summary of Updates

Additional analysis of candidate genes (with results in Table S2), Table 1 and Figure 3 revised, four additional supplementary figures, edits for clarity

## References

Aitken, S. N., & Bemmels, J. B. (2016). Time to get moving: Assisted gene flow of forest trees. Evolutionary Applications, 9(1), 271–290. 10.1111/eva.12293

Aitken, S. N., & Whitlock, M. C. (2013). Assisted gene flow to facilitate local adaptation to climate change. Annu. Rev. Ecol. Evol. Syst, 44, 367–388. 10.1146/annurev-ecolsys-110512-135747

Aitken, S. N., Yeaman, S., Holliday, J. A., Wang, T., & Curtis-McLane, S. (2008). Adaptation, migration or extirpation: Climate change outcomes for tree populations. Evolutionary Applications, 1(1), 95–111. 10.1111/j.1752-4571.2007.00013.x

Alberto, F. J., Aitken, S. N., Alía, R., González-Martínez, S. C., Hänninen, H., Kremer, A., Lefèvre, F., Lenormand, T., Yeaman, S., Whetten, R., & Savolainen, O. (2013). Potential for evolutionary responses to climate change—Evidence from tree populations. Global Change Biology, 19(6), 1645–1661. 10.1111/gcb.12181

Alexander, D. H., Novembre, J., & Lange, K. (2009). Fast model-based estimation of ancestry in unrelated individuals. Genome Research, 19(9), 1655–1664. 10.1101/gr.094052.109

Ashley, M. V., Backs, J. R., Kindsvater, L., & Abraham, S. T. (2018). Genetic variation and structure in an endemic island oak, *Quercus tomentella*, and mainland canyon oak, *Quercus chrysolepis*. International Journal of Plant Sciences, 179(2), 151–161. 10.1086/696023

Axelrod, D. I. (1939). A miocene flora from the western border of the Mohave desert. The Carnegie Institution of Washington. https://catalog.hathitrust.org/Record/001639456

Axelrod, D. I. (1944). The Sonoma Flora. In R. W. Chaney (Ed.), Pliocene floras of California and Oregon (pp. 167–218). The Carnegie Institution of Washington. https://catalog.hathitrust.org/Record/001639473

Axelrod, D. I. (1967). Geologic history of the Californian insular flora. 1st Symposium on the Biology of the California Islands, 267–314. https://repository.library.csuci.edu/bitstream/handle/10139/784/Axelrod_Geologic_Histor y_California_Insular_Flora∼.pdf?sequence=5

Beckman, E., & Jerome, D. (2017). Quercus tomentella. IUCN Red List of Threatened Species. https://www.iucnredlist.org/species/30959/2799049

Beckman, E., Meyer, A., Denvir, A., Gill, D., Man, G., Pivorunas, D., Shaw, K., & Westwood, M. (2019). Conservation Gap Analysis of Native U.S. Oaks. The Morton Arboretum.

Borcard, D., & Legendre, P. (2002). All-scale spatial analysis of ecological data by means of principal coordinates of neighbour matrices. Ecological Modelling, 153(1), 51–68. 10.1016/S0304-3800(01)00501-4

Browne, L., Wright, J. W., Fitz-Gibbon, S., Gugger, P. F., & Sork, V. L. (2019). Adaptational lag to temperature in valley oak (*Quercus lobata*) can be mitigated by genome-informed assisted gene flow. Proceedings of the National Academy of Sciences, 116(50), 25179– 25185. 10.1073/pnas.1908771116

Capblancq, T., Fitzpatrick, M. C., Bay, R. A., Exposito-Alonso, M., & Keller, S. R. (2020). Genomic Prediction of (Mal)Adaptation Across Current and Future Climatic Landscapes. Annual Review of Ecology, Evolution, and Systematics, 51(1), 245–269. 10.1146/annurev-ecolsys-020720-042553

Chang, C. C., Chow, C. C., Tellier, L. C., Vattikuti, S., Purcell, S. M., & Lee, J. J. (2015). Second-generation PLINK: Rising to the challenge of larger and richer datasets. GigaScience, 4(1), s13742–015-0047–0048. 10.1186/s13742-015-0047-8

Cheek, R. G., Forester, B. R., Salerno, P. E., Trumbo, D. R., Langin, K. M., Chen, N., Scott Sillett, T., Morrison, S. A., Ghalambor, C. K., & Chris Funk, W. (2022). Habitat-linked genetic variation supports microgeographic adaptive divergence in an island-endemic bird species. Molecular Ecology, 31(10), 2830–2846. 10.1111/mec.16438

Cingolani, P., Platts, A., Wang, L. L., Coon, M., Nguyen, T., Wang, L., Land, S. J., Lu, X., & Ruden, D. M. (2012). A program for annotating and predicting the effects of single nucleotide polymorphisms, SnpEff: SNPs in the genome of Drosophila melanogaster strain w1118; iso-2; iso-3. Fly, 6(2), 80–92. 10.4161/fly.19695

Danecek, P., Auton, A., Abecasis, G., Albers, C. A., Banks, E., DePristo, M. A., Handsaker, R. E., Lunter, G., Marth, G. T., Sherry, S. T., McVean, G., Durbin, R., & 1000 Genomes Project Analysis Group. (2011). The variant call format and VCFtools. Bioinformatics, 27(15), 2156–2158. 10.1093/bioinformatics/btr330

Danecek, P., Bonfield, J. K., Liddle, J., Marshall, J., Ohan, V., Pollard, M. O., Whitwham, A., Keane, T., McCarthy, S. A., Davies, R. M., & Li, H. (2021). Twelve years of SAMtools and BCFtools. GigaScience, 10(2), giab008. 10.1093/gigascience/giab008

Delaney, K. S., & Wayne, R. K. (2005). Adaptive Units for Conservation: Population Distinction and Historic Extinctions in the Island Scrub-Jay. Conservation Biology, 19(2), 523–533. 10.1111/j.1523-1739.2005.00424.x

Di Santo, L. N., Hoban, S., Parchman, T. L., Wright, J. W., & Hamilton, J. A. (2022). Reduced representation sequencing to understand the evolutionary history of Torrey pine (*Pinus torreyana* parry) with implications for rare species conservation. Molecular Ecology, 31(18), 4622–4639. 10.1111/mec.16615

Dormann, C. F., Elith, J., Bacher, S., Buchmann, C., Carl, G., Carré, G., Marquéz, J. R. G., Gruber, B., Lafourcade, B., Leitão, P. J., Münkemüller, T., McClean, C., Osborne, P. E., Reineking, B., Schröder, B., Skidmore, A. K., Zurell, D., & Lautenbach, S. (2013). Collinearity: A review of methods to deal with it and a simulation study evaluating their performance. Ecography, 36(1), 27–46. 10.1111/j.1600-0587.2012.07348.x

Ellis, N., Smith, S. J., & Pitcher, C. R. (2012). Gradient Forests: Calculating importance gradients on physical predictors. Ecology, 93, 156–168.

Fischer, D. T., Still, C. J., Ebert, C. M., Baguskas, S. A., & Park Williams, A. (2016). Fog drip maintains dry season ecological function in a California coastal pine forest. Ecosphere, 7(6), e01364. 10.1002/ecs2.1364

Fischer, D. T., Still, C. J., & Williams, A. P. (2009). Significance of summer fog and overcast for drought stress and ecological functioning of coastal California endemic plant species. Journal of Biogeography, 36(4), 783–799. 10.1111/j.1365-2699.2008.02025.x

Fitzpatrick, M. C., Chhatre, V. E., Soolanayakanahally, R. Y., & Keller, S. R. (2021). Experimental support for genomic prediction of climate maladaptation using the machine learning approach Gradient Forests. Molecular Ecology Resources, 21(8), 2749–2765. 10.1111/1755-0998.13374

Fitzpatrick, M. C., & Keller, S. R. (2015). Ecological genomics meets community-level modelling of biodiversity: Mapping the genomic landscape of current and future environmental adaptation. Ecology Letters, 18(1), 1–16. 10.1111/ele.12376

Fontenot, A. (2024). Fast-wcfst (Version Commit 338682f0b1). https://codeberg.org/fontenot/fast-wcfst

Forester, B. R., Lasky, J. R., Wagner, H. H., & Urban, D. L. (2018). Comparing methods for detecting multilocus adaptation with multivariate genotype–environment associations. Molecular Ecology, 27(9), 2215–2233. 10.1111/mec.14584

Gamboa, M. P., Ghalambor, C. K., Scott Sillett, T., Morrison, S. A., & Chris Funk, W. (2022). Adaptive divergence in bill morphology and other thermoregulatory traits is facilitated by restricted gene flow in song sparrows on the California Channel Islands. Molecular Ecology, 31(2), 603–619. 10.1111/mec.16253

Gougherty, A. V., Keller, S. R., & Fitzpatrick, M. C. (2021). Maladaptation, migration and extirpation fuel climate change risk in a forest tree species. Nature Climate Change, 11(2), 166–171. 10.1038/s41558-020-00968-6

Grummer, J. A., Booker, T. R., Matthey-Doret, R., Nietlisbach, P., Thomaz, A. T., & Whitlock, M. C. (2022). The immediate costs and long-term benefits of assisted gene flow in large populations. Conservation Biology, 36(4), e13911. 10.1111/cobi.13911

Gugger, P. F., Fitz-Gibbon, S. T., Albarrán-Lara, A., Wright, J. W., & Sork, V. L. (2021). Landscape genomics of *Quercus lobata* reveals genes involved in local climate adaptation at multiple spatial scales. Molecular Ecology, 30(2), 406–423. 10.1111/mec.15731

Gugger, P. F., Liang, C. T., Sork, V. L., Hodgskiss, P., & Wright, J. W. (2018). Applying landscape genomic tools to forest management and restoration of Hawaiian koa (*Acacia koa*) in a changing environment. Evolutionary Applications, 11(2), 231–242. 10.1111/eva.12534

Hamilton, J. A., Royauté, R., Wright, J. W., Hodgskiss, P., & Ledig, F. T. (2017). Genetic conservation and management of the California endemic, Torrey pine (*Pinus torreyana* Parry): Implications of genetic rescue in a genetically depauperate species. Ecology and Evolution, 7(18), 7370–7381. 10.1002/ece3.3306

IPCC. (2014). Climate Change 2014: Synthesis Report. Contribution of Working Groups I, II and III to the Fifth Assessment Report of the Intergovernmental Panel on Climate Change. https://www.ipcc.ch/report/ar5/syr/

Jia, K.-H., Zhao, W., Maier, P. A., Hu, X.-G., Jin, Y., Zhou, S.-S., Jiao, S.-Q., El-Kassaby, Y. A., Wang, T., Wang, X.-R., & Mao, J.-F. (2020). Landscape genomics predicts climate change-related genetic offset for the widespread *Platycladus orientalis* (Cupressaceae). Evolutionary Applications, 13(4), 665–676. 10.1111/eva.12891

Kennett, D. J., Kennett, J. P., West, G. J., Erlandson, J. M., Johnson, J. R., Hendy, I. L., West, A., Culleton, B. J., Jones, T. L., & Stafford, T. W. (2008). Wildfire and abrupt ecosystem disruption on California’s Northern Channel Islands at the Ållerød–Younger Dryas boundary (13.0–12.9ka). Quaternary Science Reviews, 27(27), 2530–2545. 10.1016/j.quascirev.2008.09.006

Kremer, A., & Hipp, A. L. (2020). Oaks: An evolutionary success story. New Phytologist, 226(4), 987–1011. 10.1111/nph.16274

Lachmuth, S., Capblancq, T., Prakash, A., Keller, S. R., & Fitzpatrick, M. C. (2024). Novel genomic offset metrics integrate local adaptation into habitat suitability forecasts and inform assisted migration. Ecological Monographs, 94(1), e1593. 10.1002/ecm.1593

Langin, K. M., Sillett, T. S., Funk, W. C., Morrison, S. A., Desrosiers, M. A., & Ghalambor, C. K. (2015). Islands within an island: Repeated adaptive divergence in a single population. Evolution, 69(3), 653–665. 10.1111/evo.12610

Leimu, R., & Fischer, M. (2008). A meta-analysis of local adaptation in plants. PLOS ONE, 3(12), e4010. 10.1371/journal.pone.0004010

Lind, B. M., Candido-Ribeiro, R., Singh, P., Lu, M., Vidakovic, D. O., Booker, T. R., Whitlock, M. C., Yeaman, S., Isabel, N., & Aitken, S. N. (2023). How useful is genomic data for predicting maladaptation to future climate? BioRxiv. 10.1101/2023.02.10.528022

Lotterhos, K. E. (2024). Interpretation issues with “genomic vulnerability” arise from conceptual issues in local adaptation and maladaptation. Evolution Letters, qrae004. 10.1093/evlett/qrae004

Martins, K., Gugger, P. F., Llanderal-Mendoza, J., González-Rodríguez, A., Fitz-Gibbon, S. T., Zhao, J.-L., Rodríguez-Correa, H., Oyama, K., & Sork, V. L. (2018). Landscape genomics provides evidence of climate-associated genetic variation in Mexican populations of *Quercus rugosa*. Evolutionary Applications, 11(10), 1842–1858. 10.1111/eva.12684

Mead, A., Fitz-Gibbon, S. T., Escalona, M., Beraut, E., Sacco, S., Marimuthu, M. P. A., Nguyen, O., & Sork, V. L. (2024). The genome assembly of Island Oak (Quercus tomentella), a relictual island tree species. *Journal of Heredity*, esae002. 10.1093/jhered/esae002

Meirmans, P. G. (2015). Seven common mistakes in population genetics and how to avoid them. Molecular Ecology, 24(13), 3223–3231. 10.1111/mec.13243

Morrison, S. A., Sillett, T. S., Ghalambor, C. K., Fitzpatrick, J. W., Graber, D. M., Bakker, V. J., Bowman, R., Collins, C. T., Collins, P. W., Delaney, K. S., Doak, D. F., Koenig, W. D., Laughrin, L., Lieberman, A. A., Marzluff, J. M., Reynolds, M. D., Scott, J. M., Stallcup, J. A., Vickers, W., & Boyce, W. M. (2011). Proactive Conservation Management of an Island-endemic Bird Species in the Face of Global Change. BioScience, 61(12), 1013– 1021. 10.1525/bio.2011.61.12.11

Muller, C. H. (1965). Relictual origins of insular endemics in *Quercus*. 1st Symposium on the Biology of the California Islands.

O’Donnell, S. T., Fitz-Gibbon, S. T., & Sork, V. L. (2021). Ancient Introgression Between Distantly Related White Oaks (Quercus sect. Quercus) Shows Evidence of Climate-Associated Asymmetric Gene Exchange. Journal of Heredity, 112(7), 663–670. 10.1093/jhered/esab053

Oksanen, J., Blanchet, F. G., Friendly, M., Kindt, R., Legendre, P., McGlinn, D., Minchin, P. R., O’Hara, R. B., Simpson, G. L., Solymos, P., Stevens, M. H. H., Szoecs, E., & Wagner, H. (2019). vegan: Community Ecology Package (Version 2.5-5). https://CRAN.R-project.org/package=vegan

Ortego, J., Gugger, P. F., & Sork, V. L. (2018). Genomic data reveal cryptic lineage diversification and introgression in Californian golden cup oaks (section *Protobalanus*). New Phytologist, 218(2), 804–818. 10.1111/nph.14951

Peres-Neto, P. R., & Jackson, D. A. (2001). How well do multivariate data sets match? The advantages of a Procrustean superimposition approach over the Mantel test. Oecologia, 129(2), 169–178. 10.1007/s004420100720

Pesendorfer, M. B., Baker, C. M., Stringer, M., McDonald-Madden, E., Bode, M., McEachern, A. K., Morrison, S. A., & Sillett, T. S. (2018). Oak habitat recovery on California’s largest islands: Scenarios for the role of corvid seed dispersal. Journal of Applied Ecology, 55(3), 1185–1194. 10.1111/1365-2664.13041

Petit, R. J., & Hampe, A. (2006). Some evolutionary consequences of being a tree. Annual Review of Ecology, Evolution, and Systematics, 37, 187–214.

Quinlan, A. R., & Hall, I. M. (2010). BEDTools: A flexible suite of utilities for comparing genomic features. Bioinformatics, 26(6), 841–842. 10.1093/bioinformatics/btq033

Reeder-Myers, L., Erlandson, J. M., Muhs, D. R., & Rick, T. C. (2015). Sea level, paleogeography, and archeology on California’s Northern Channel islands. Quaternary Research, 83(2), 263–272. 10.1016/j.yqres.2015.01.002

Rellstab, C. (2021). Genomics helps to predict maladaptation to climate change. Nature Climate Change, 11(2), 85–86. 10.1038/s41558-020-00964-w

Rellstab, C., Gugerli, F., Eckert, A. J., Hancock, A. M., & Holderegger, R. (2015). A practical guide to environmental association analysis in landscape genomics. Molecular Ecology, 24(17), 4348–4370. 10.1111/mec.13322

Rick, T. C., Hofman, C. A., & Reeder-Myers, L. A. (2019). Why Translocate? Evaluating the Evidence and Reasons for Ancient Human Introductions of Wildlife to California’s Islands. In K. M. Gill, M. Fauvelle, & J. M. Erlandson (Eds.), An Archaeology of Abundance: Reevaluating the Marginality of California’s Islands (pp. 248–272). University Press of Florida.

Rosatti, T. J., & Tucker, J. M. (2014). Quercus chrysolepis. In Jepson Flora Project. https://ucjeps.berkeley.edu/eflora/eflora_display.php?tid=40565

Rosenberger, K., Schumacher, E., Brown, A., & Hoban, S. (2022). Species-tailored sampling guidelines remain an efficient method to conserve genetic diversity ex situ: A study on threatened oaks. Biological Conservation, 275, 109755. 10.1016/j.biocon.2022.109755

Savolainen, O., Lascoux, M., & Merilä, J. (2013). Ecological genomics of local adaptation. Nature Reviews Genetics, 14(11), 807–820. 10.1038/nrg3522

Savolainen, O., Pyhäjärvi, T., & Knürr, T. (2007). Gene Flow and Local Adaptation in Trees. Annual Review of Ecology, Evolution, and Systematics, 38(1), 595–619. 10.1146/annurev.ecolsys.38.091206.095646

Sawyer, J. O., Keeler-Wolf, T., & Evens, J. M. (2009). CNPS Alliance: Quercus tomentella— Lyonothamnus floribundus. In A Manual of California Vegetation, Second Edition (p. 1300). California Native Plant Society. https://vegetation.cnps.org/alliance/523

Sexton, J. P., Hangartner, S. B., & Hoffmann, A. A. (2014). Genetic Isolation by Environment or Distance: Which Pattern of Gene Flow Is Most Common? Evolution, 68(1), 1–15. 10.1111/evo.12258

Shryock, D. F., Washburn, L. K., DeFalco, L. A., & Esque, T. C. (2021). Harnessing landscape genomics to identify future climate resilient genotypes in a desert annual. Molecular Ecology, 30(3), 698–717. 10.1111/mec.15672

Sork, V. L. (2016). Gene flow and natural selection shape spatial patterns of genes in tree populations: Implications for evolutionary processes and applications. Evolutionary Applications, 9(1), 291–310. 10.1111/eva.12316

Sork, V. L. (2017). Genomic studies of local adaptation in natural plant populations. Journal of Heredity, 109(1), 3–15. 10.1093/jhered/esx091

Sork, V. L., Aitken, S. N., Dyer, R. J., Eckert, A. J., Legendre, P., & Neale, D. B. (2013). Putting the landscape into the genomics of trees: Approaches for understanding local adaptation and population responses to changing climate. Tree Genetics and Genomes, 9(4), 901– 911. 10.1007/s11295-013-0596-x

Sork, V. L., Cokus, S. J., Fitz-Gibbon, S. T., Zimin, A. V., Puiu, D., Garcia, J. A., Gugger, P. F., Henriquez, C. L., Zhen, Y., Lohmueller, K. E., Pellegrini, M., & Salzberg, S. L. (2022). High-quality genome and methylomes illustrate features underlying evolutionary success of oaks. Nature Communications, 13(1), 1. 10.1038/s41467-022-29584-y

Suarez-Gonzalez, A., Lexer, C., & Cronk, Q. C. B. (2018). Adaptive introgression: A plant perspective. Biology Letters, 14(3), 20170688. 10.1098/rsbl.2017.0688

Trumbo, D. R., Funk, W. C., Pauly, G. B., & Robertson, J. M. (2021). Conservation genetics of an island-endemic lizard: Low Ne and the critical role of intermediate temperatures for genetic connectivity. Conservation Genetics, 22(5), 783–797. 10.1007/s10592-021-01362-1

Underwood, E. C., Hollander, A. D., Safford, H. D., Kim, J. B., Srivastava, L., & Drapek, R. J. (2019). The impacts of climate change on ecosystem services in southern California. Ecosystem Services, 39, 101008. 10.1016/j.ecoser.2019.101008

Villa-Machío, I., Fernández de Castro, A. G., Fuertes-Aguilar, J., & Nieto Feliner, G. (2020). Colonization history of the Canary Islands endemic Lavatera acerifolia, (Malvaceae) unveiled with genotyping-by-sequencing data and niche modelling. Journal of Biogeography, 47(4), 993–1005. 10.1111/jbi.13808

Wang, I. J., & Bradburd, G. S. (2014). Isolation by environment. Molecular Ecology, 23(23), 5649–5662. 10.1111/mec.12938

Westwood, M., Cavender, N., Meyer, A., & Smith, P. (2021). Botanic garden solutions to the plant extinction crisis. PLANTS, PEOPLE, PLANET, 3(1), 22–32. 10.1002/ppp3.10134

Williams, A. P., Still, C. J., Fischer, D. T., & Leavitt, S. W. (2008). The influence of summertime fog and overcast clouds on the growth of a coastal Californian pine: A tree-ring study. Oecologia, 156(3), 601–611. 10.1007/s00442-008-1025-y

Willing, E.-M., Dreyer, C., & Oosterhout, C. van. (2012). Estimates of Genetic Differentiation Measured by FST Do Not Necessarily Require Large Sample Sizes When Using Many SNP Markers. PLOS ONE, 7(8), e42649. 10.1371/journal.pone.0042649

Woolsey, J., Hanna, C., Mceachern, K., Anderson, S., & Hartman, B. D. (2018). Regeneration and Expansion of Quercus tomentella (Island Oak) Groves on Santa Rosa Island. Western North American Naturalist, 78(4), 758–767. 10.3398/064.078.0415

Yu, Y., Aitken, S. N., Rieseberg, L. H., & Wang, T. (2022). Using landscape genomics to delineate seed and breeding zones for lodgepole pine. New Phytologist, 235(4), 1653– 1664. 10.1111/nph.18223

