## Supplementary Figures for "Comparison of conservation strategies for California Channel Island Oak (*Quercus tomentella*) using climate suitability predicted from genomic data"

Supplementary Material

| | Mainland_Qchr | Santa Rosa Island_Qtom | Santa Cruz Island_Qtom | Santa Cruz Island_hybrid | Anacapa Island_Qtom | Catalina Island_hybrid | Catalina Island_Qchr | San Clemente Island_hybrid | San Clemente Island_Qchr | Guadalupe Island_Guadalup | Guadalupe Island_hybrid |
|---|---|---|---|---|---|---|---|---|---|---|---|
| Mainland_Qchr | NA | 0.063 | 0.05 | 0.028 | 0.065 | 0.027 | 0.014 | 0.026 | -0.099 | 0.056 | -0.019 |
| Santa Rosa Island_Qtom | 0.063 | NA | 0.017 | 0.025 | 0.036 | 0.029 | 0.059 | 0.021 | -0.055 | 0.061 | 0.034 |
| Santa Cruz Island_Qtom | 0.05 | 0.017 | NA | 0.012 | 0.022 | 0.02 | 0.043 | 0.013 | -0.061 | 0.057 | 0.021 |
| Santa Cruz Island_hybrid | 0.028 | 0.025 | 0.012 | NA | 0.024 | 0.013 | 0.017 | 0.009 | -0.102 | 0.047 | -0.015 |
| Anacapa Island_Qtom | 0.065 | 0.036 | 0.022 | 0.024 | NA | 0.03 | 0.078 | 0.021 | 0.063 | 0.079 | 0.094 |
| Catalina Island_hybrid | 0.027 | 0.029 | 0.02 | 0.013 | 0.03 | NA | 0.011 | 0.01 | -0.1 | 0.047 | -0.012 |
| Catalina Island_Qchr | 0.014 | 0.059 | 0.043 | 0.017 | 0.078 | 0.011 | NA | 0.013 | -0.037 | 0.065 | 0.016 |
| San Clemente Island_hybrid | 0.026 | 0.021 | 0.013 | 0.009 | 0.021 | 0.01 | 0.013 | NA | -0.112 | 0.044 | -0.02 |
| San Clemente Island_Qchr | -0.099 | -0.055 | -0.061 | -0.102 | 0.063 | -0.1 | -0.037 | -0.112 | NA | -0.044 | -0.045 |
| uadalupe Island_Guadalupe | 0.056 | 0.061 | 0.057 | 0.047 | 0.079 | 0.047 | 0.065 | 0.044 | -0.044 | NA | -0.014 |
| Guadalupe Island_hybrid | -0.019 | 0.034 | 0.021 | -0.015 | 0.094 | -0.012 | 0.016 | -0.02 | -0.045 | -0.014 | NA |

**Figure S1.** Pairwise $F_{ST}$ values for individuals grouped by island and ancestry. Ancestry for individuals is defined based on based on K=3 assignments, categorized as Channel Islands *Q. tomentella* ("Qtom"), *Q. chrysolepis* ("Qchr"), or Guadalupe Island Oak ("Guadalupe") if ancestry proportions for these categories were ≥0.9, and otherwise categorized as hybrid with mixed ancestry.

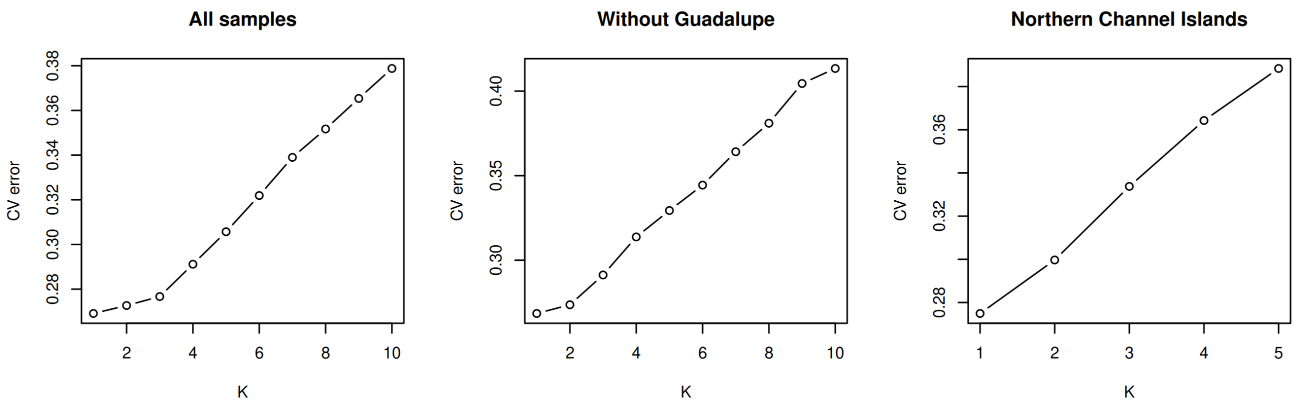


**Figure S2**. Cross-validation errors of ADMIXTURE at different K values, run on datasets including all samples, excluding Guadalupe, and including only the Northern Channel Islands (Santa Rosa and Santa Cruz).

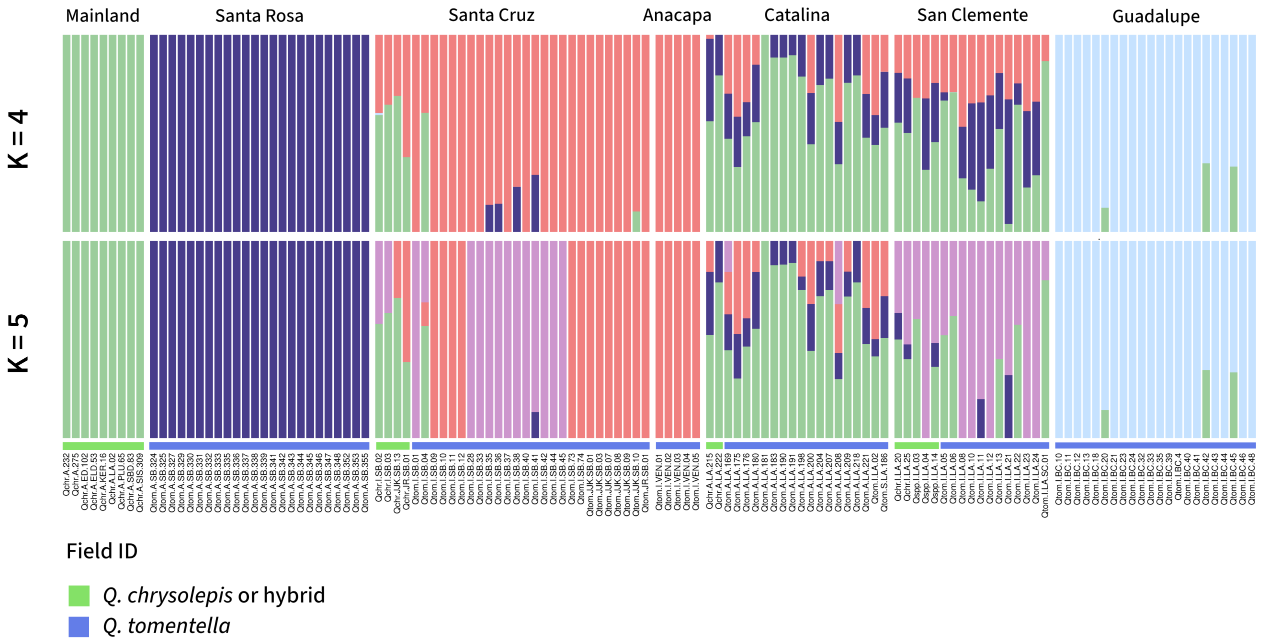


**Figure S3**. ADMIXTURE results when the number of ancestral groups (K) was set to 4 and 5, which had higher cross-validation errors than lower K values (Figure S1) but may describe more recent splits in hierarchical structure. Each color indicates a group, and each bar shows the ancestry proportions of each group for an individual tree. Colored bars at the bottom indicate the field species ID. Individuals with admixed ancestry were often, but not always, identified as *Q. chrysolepis* or hybrids.

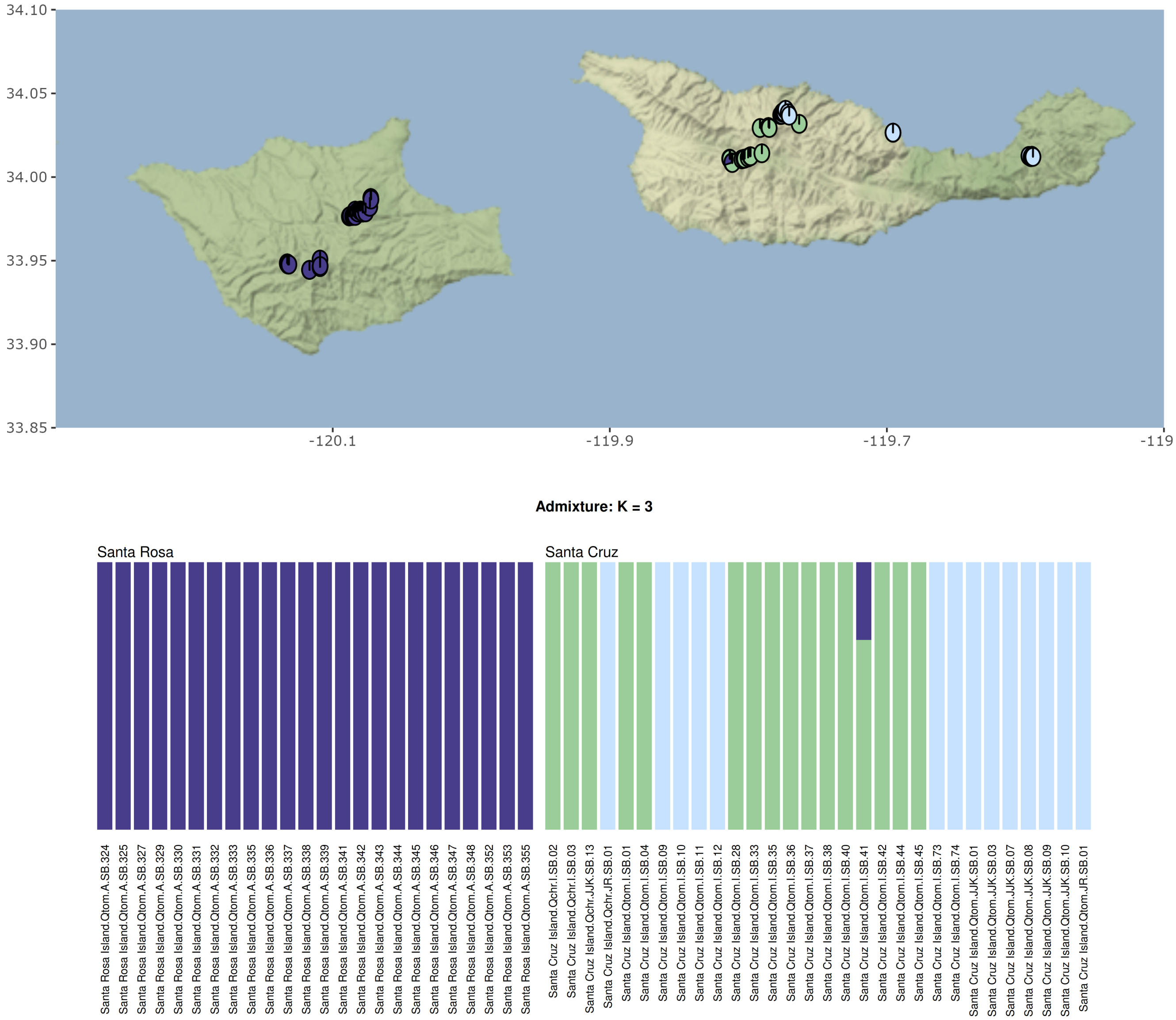


**Figure S4.** Results of ADMIXTURE analysis with K=3, run on a data subset including only Santa Rosa and Santa Cruz island individuals, revealed one individual on the Western end of Santa Cruz island with partial ancestry matching that of all Santa Rosa Island trees, suggesting gene flow from Santa Rosa to Santa Cruz Island.

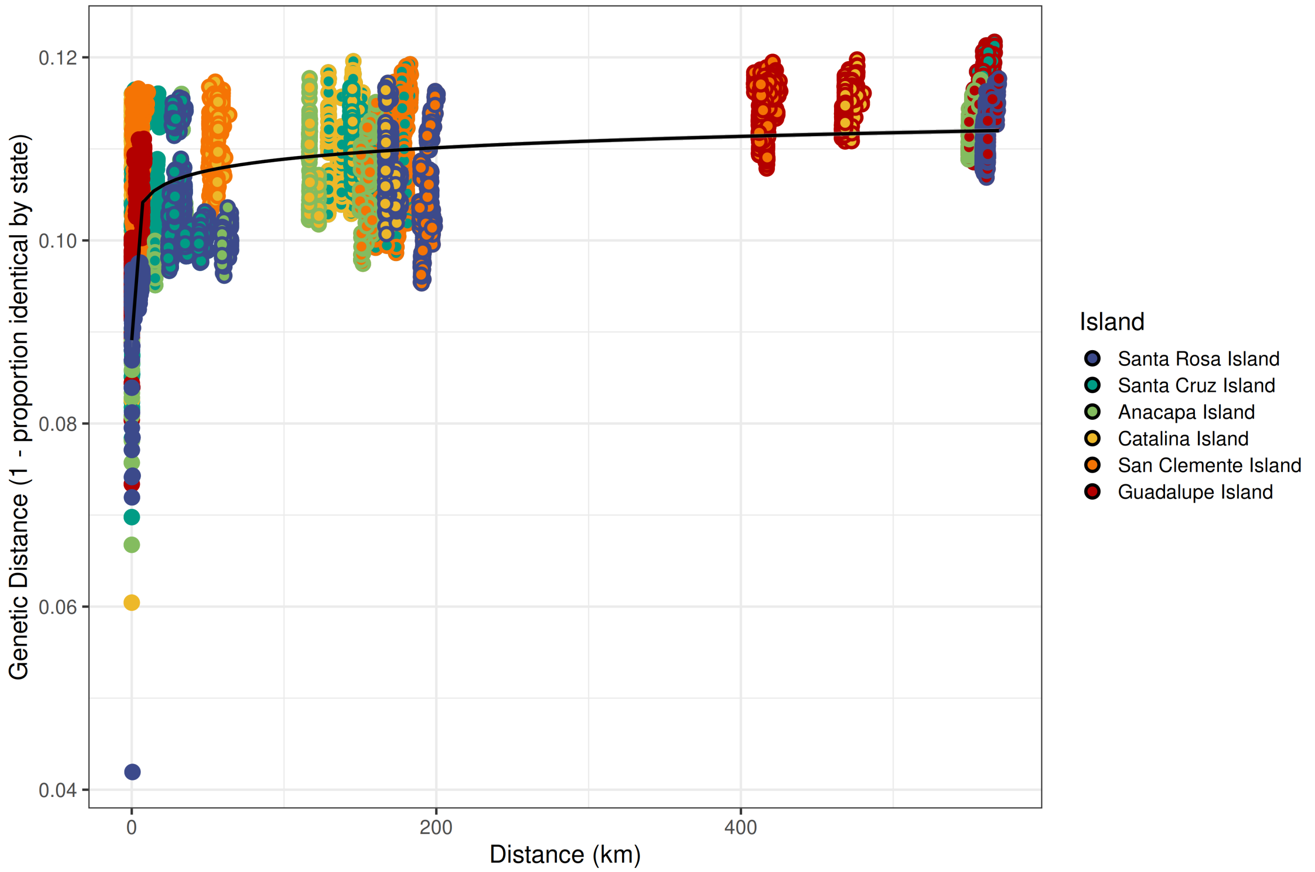


**Figure S5.** Test of isolation-by-distance. Each point indicates the geographic distance and genetic distance of a pair of sampled individuals, with the two colors indicating the island of each individual in the pair (solid points are pairs from the same island). Line shows a best-fit logistic curve produced using the lm function in R (P <2.2e-16, adjusted $R^2$ = 0.3421).

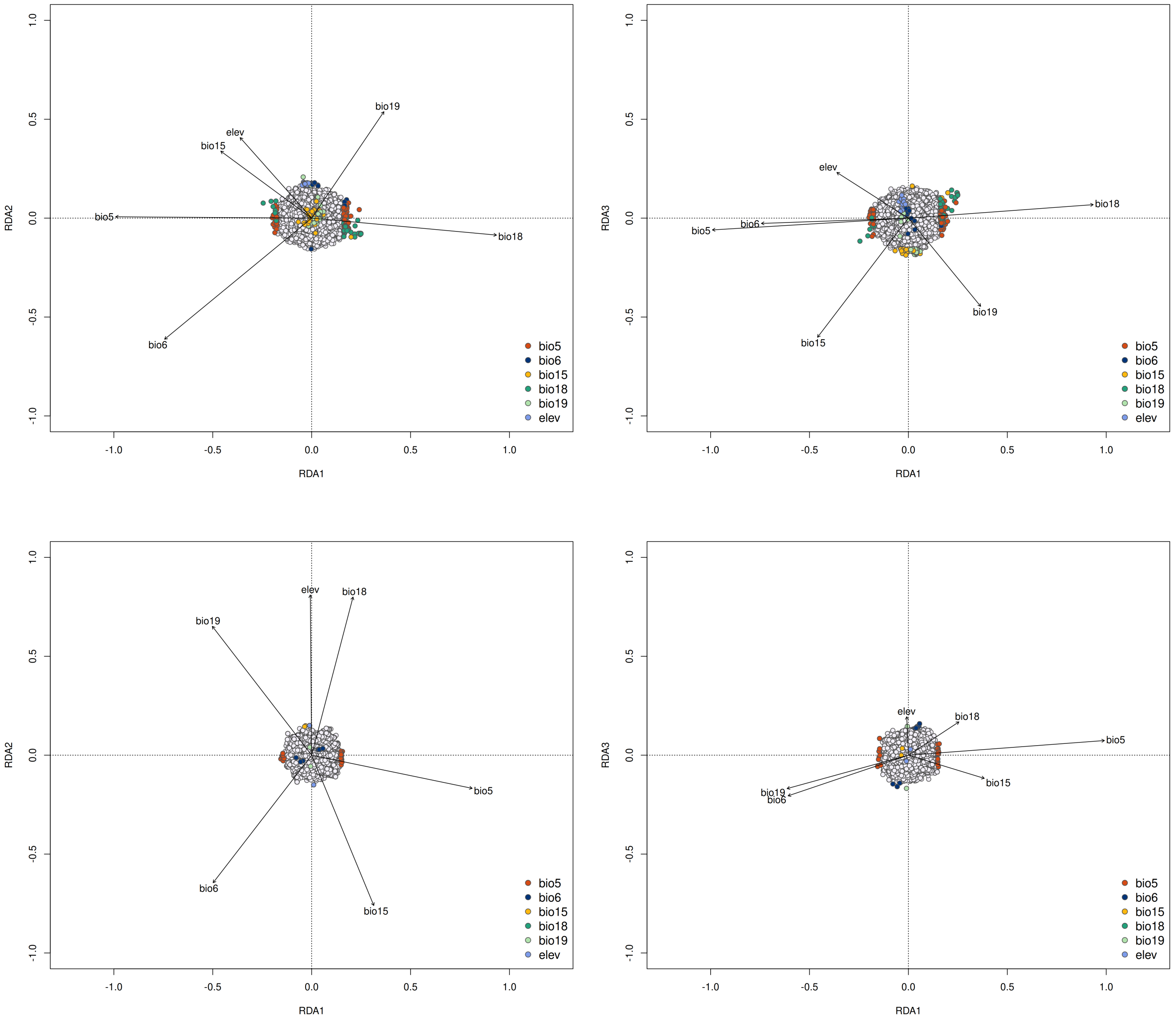


**Figure S6**. Candidate SNPs identified from the redundancy analysis on the four large Channel Islands (Santa Rosa, Santa Cruz, San Clemente, and Catalina, top) and on the Guadalupe Island individuals (bottom). Each point indicates a SNP, with colored points indicating candidate SNPs that are outliers in their climate loadings (≥4 standard deviations), indicating they are strongly associated with climate. Colors indicate the climate variable they have the greatest correlation with. Bioclimatic variables are as follows: BIO5 = Maximum Temperature of Warmest Month (°C), BIO6 = Minimum Temperature of Coldest Month (°C), BIO15 = Precipitation Seasonality (Coefficient of Variation), BIO18 = Precipitation of Warmest Quarter (mm), BIO19 = Precipitation of Coldest Quarter (mm).

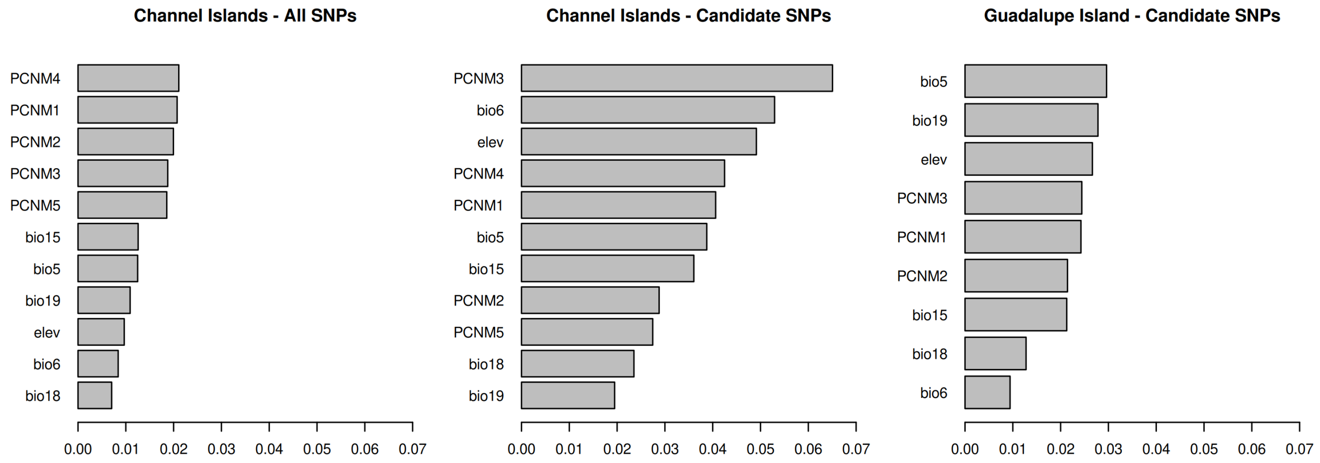


**Figure S7**. $R^2$ weighted importance values of the climate (bioclim) and geographic (PCNM) variables for each dataset. Importance values are averaged across all SNPs for each variable. Bioclimatic variables are as follows: BIO5 = Maximum Temperature of Warmest Month (°C), BIO6 = Minimum Temperature of Coldest Month (°C), BIO15 = Precipitation Seasonality (Coefficient of Variation), BIO18 = Precipitation of Warmest Quarter (mm), BIO19 = Precipitation of Coldest Quarter (mm).

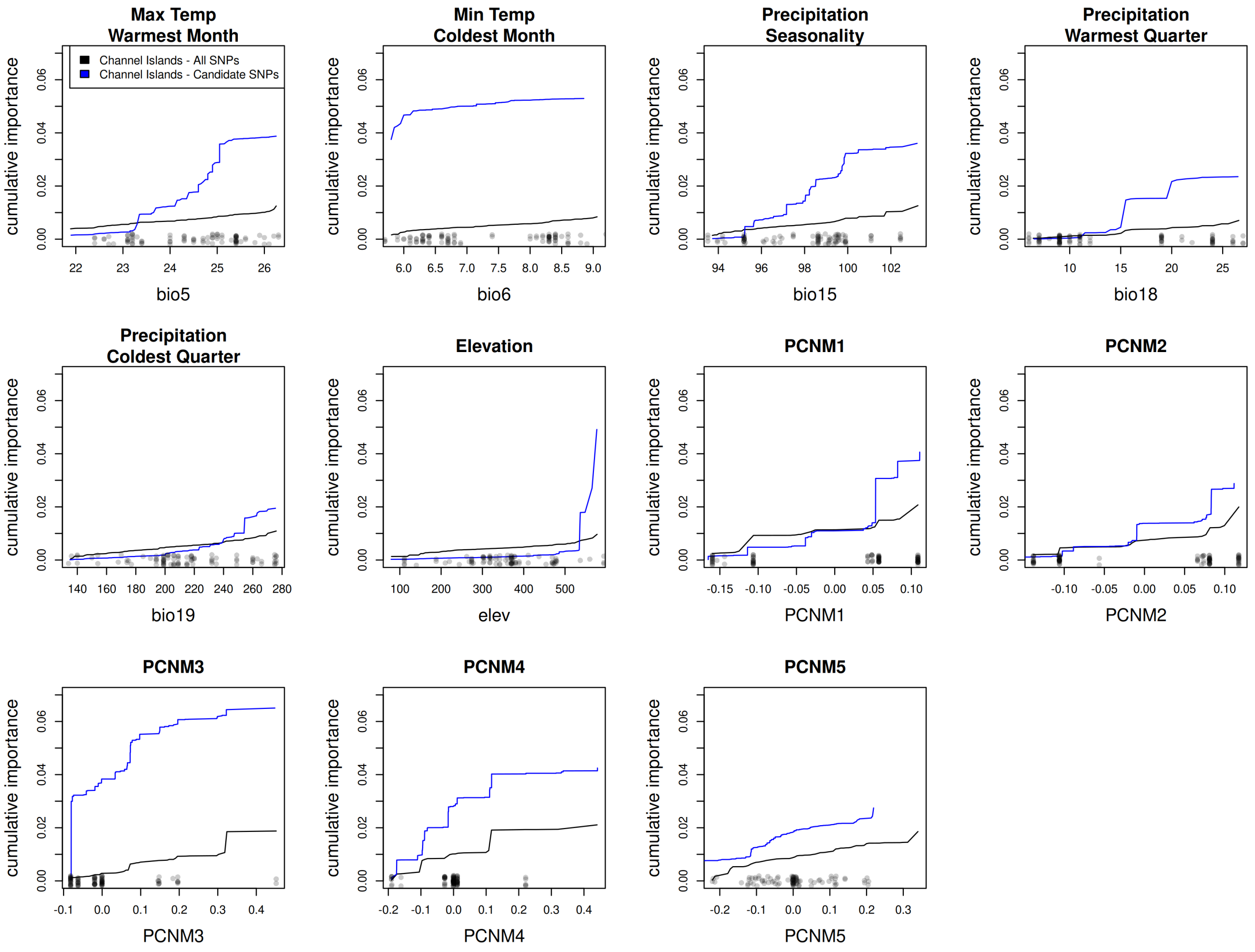


**Figure S8**. Cumulative importance of climatic and geographic variables in explaining allelic turnover in the California Channel Islands dataset, for all SNPs (black) and for candidate climate-associated SNPs (blue). Steep vertical slopes indicate greater differences in allele frequency for those values of the environmental gradient. Points at bottom of plot indicate actual values for sampled localities and are jittered vertically. Bioclimatic variables are as follows: BIO5 = Maximum Temperature of Warmest Month (°C), BIO6 = Minimum Temperature of Coldest Month (°C), BIO15 = Precipitation Seasonality (Coefficient of Variation), BIO18 = Precipitation of Warmest Quarter (mm), BIO19 = Precipitation of Coldest Quarter (mm).

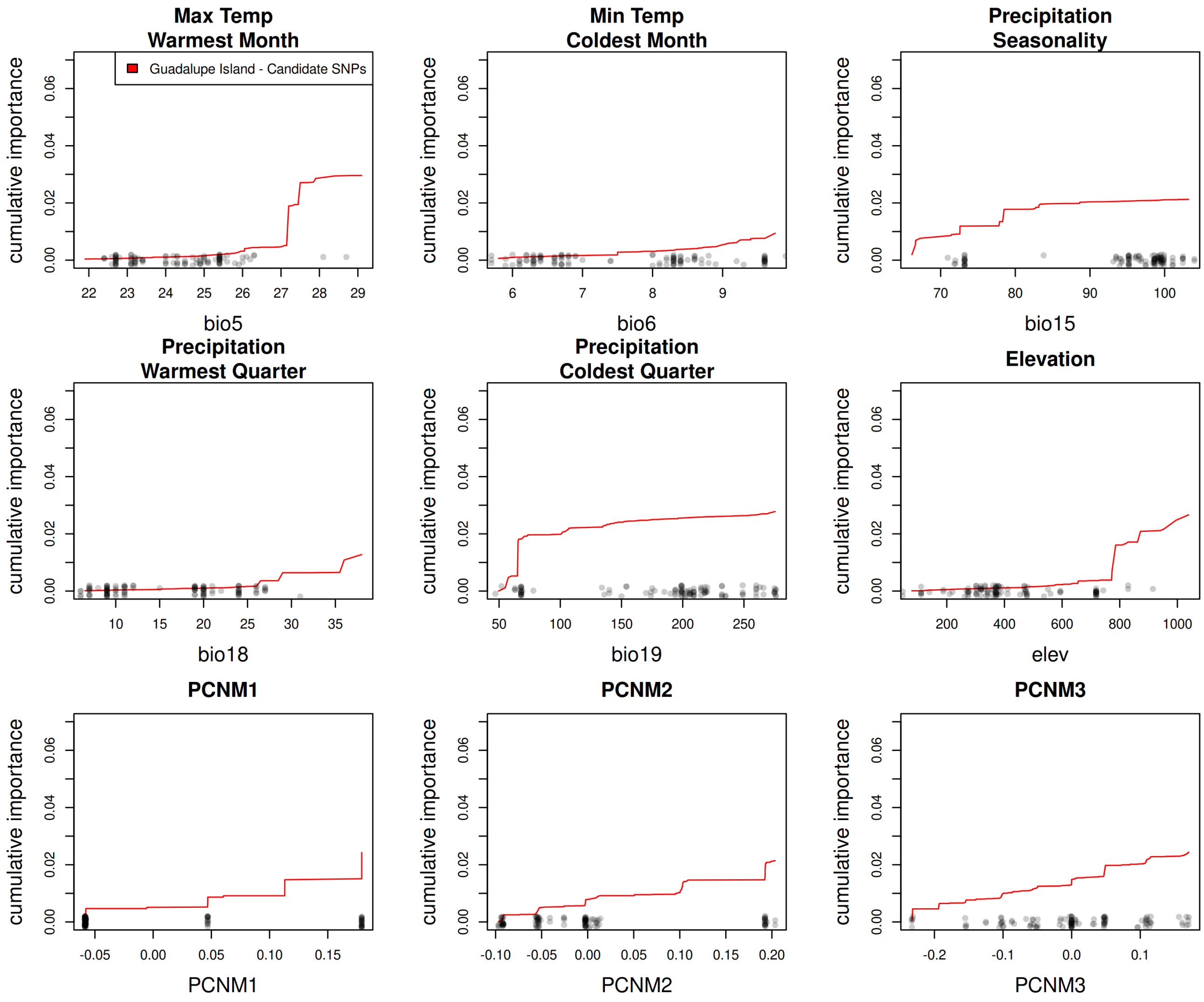


**Figure S9**. Cumulative importance of climatic and geographic variables in explaining allelic turnover in the candidate climate-associated SNPs for Guadalupe Island. Steep vertical slopes indicate greater differences in allele frequency for those values of the environmental gradient. Points at bottom of plot indicate actual values for sampled localities and are jittered vertically. Bioclimatic variables are as follows: BIO5 = Maximum Temperature of Warmest Month (°C), BIO6 = Minimum Temperature of Coldest Month (°C), BIO15 = Precipitation Seasonality (Coefficient of Variation), BIO18 = Precipitation of Warmest Quarter (mm), BIO19 = Precipitation of Coldest Quarter (mm).

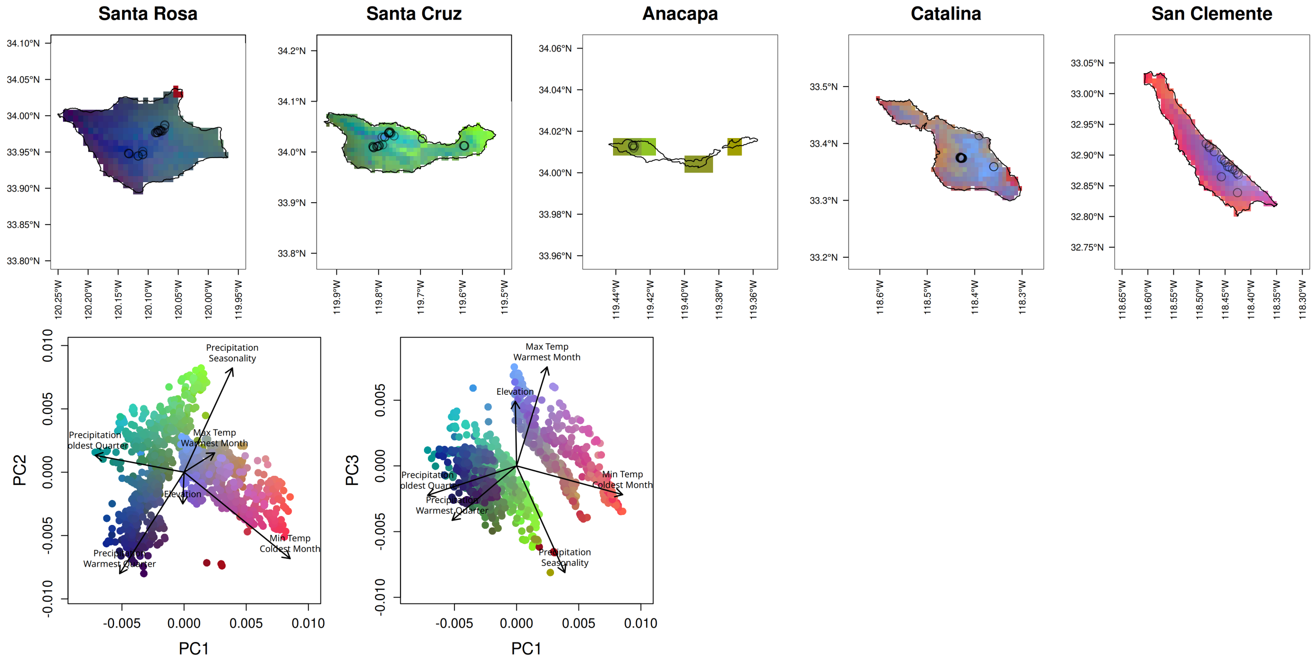


**Figure S10**. (Top) Maps of genetic turnover predicted by gradient forest for the set of all SNPs on each of the Channel Islands. Climate variables are scaled by their importance in predicting genetic variation in the gradient forest model and their relationship with allelic turnover as shown in Figure S2, then the genomic composition of a cell is predicted based on its climate. The first three PCs of the genomic composition are mapped to RGB values, so regions with similar colors are expected to have similar genomic composition. Points indicate sample locations. (Bottom) PCA of scaled climate values for each cell, with colors matching each grid cell in maps.

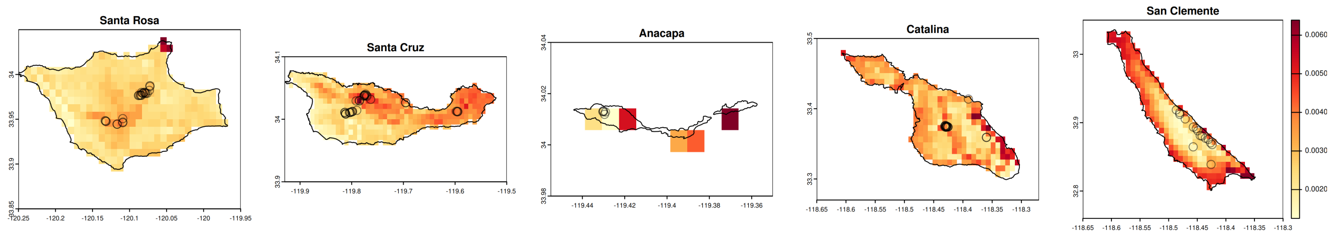


**Figure S11**. Difference in genomic turnover between the set of all SNPs (Figure S4) and the candidate climate-associated SNPs (Figure 4), calculated using Procrustes residuals. Darker red indicates the two datasets make different predictions.

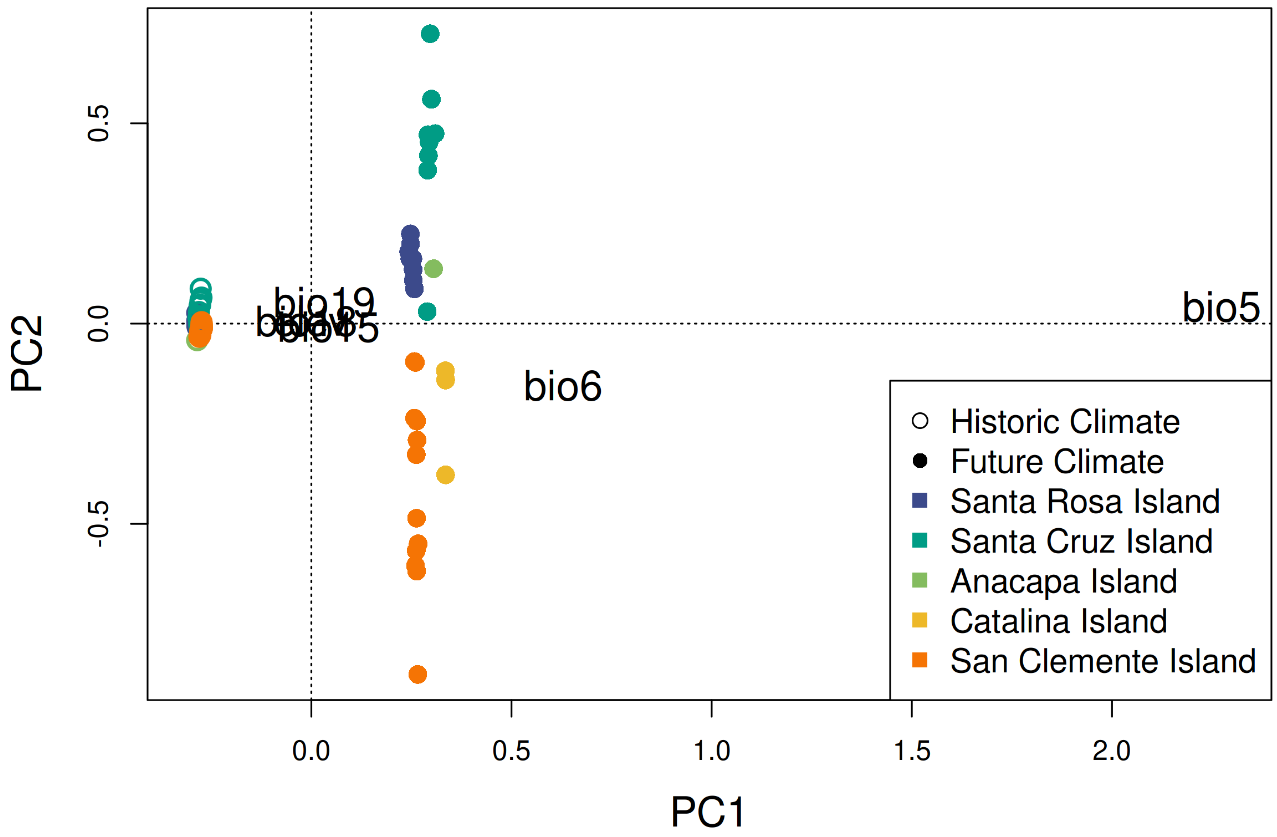


**Figure S12**. PCA of past and future climate variables, transformed by gradient forest based on their explanatory power for genomic data. The difference in these values are used to calculate the genomic offset for a site. Future climates are far outside the range of historic climates for Island Oak in California.

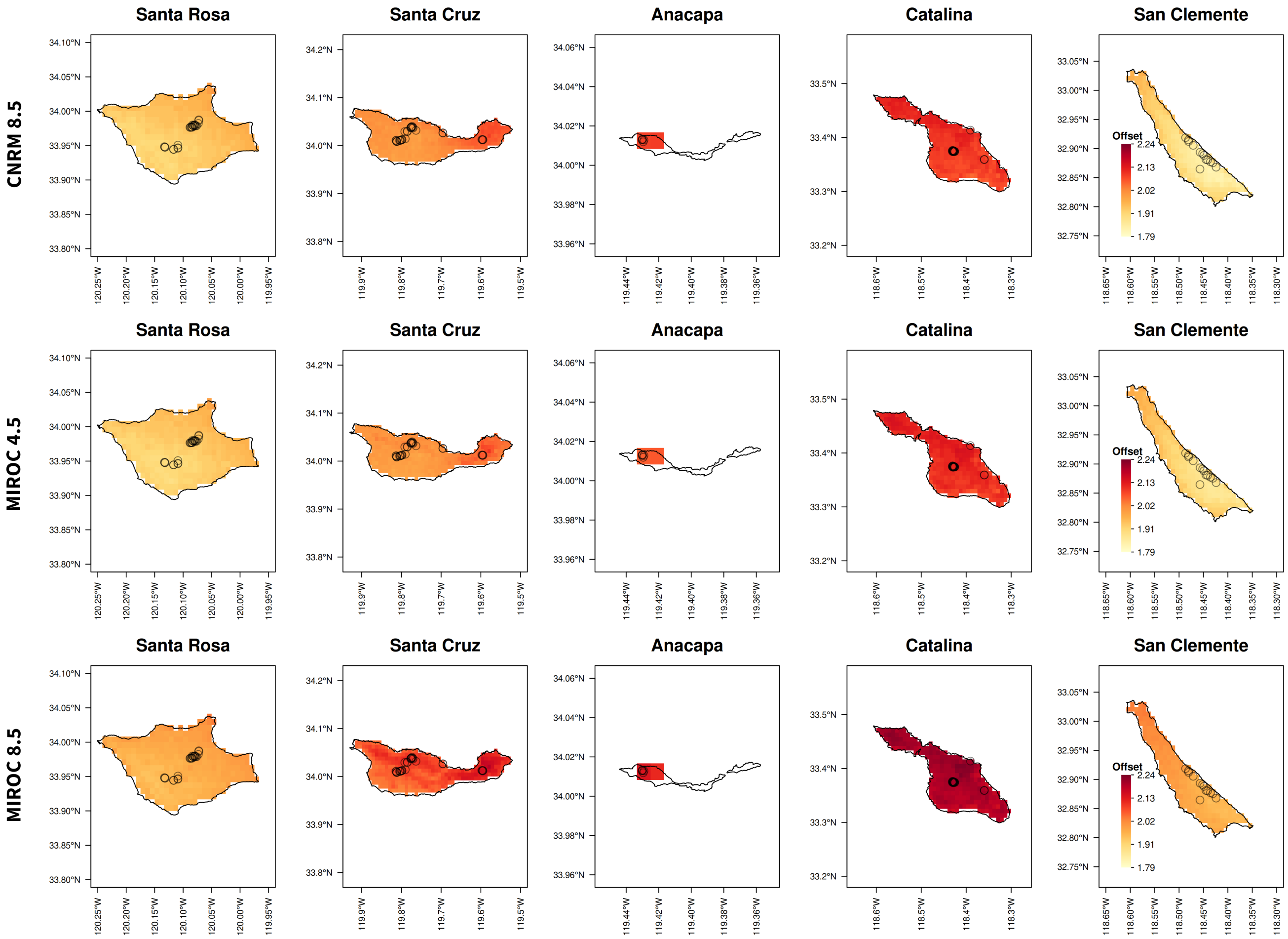


**Figure S13**. Map of the genomic offset for the Channel Islands under three additional climate scenarios (CNRM 8.5, MIROC 4.5, and MIROC 8.5) as in Figure 4B. Genomic offset is the predicted difference in the current genetic composition and that which would be ideal under future climate conditions. Darker red indicates a greater likelihood of future maladaptation. Points indicate sample locations.

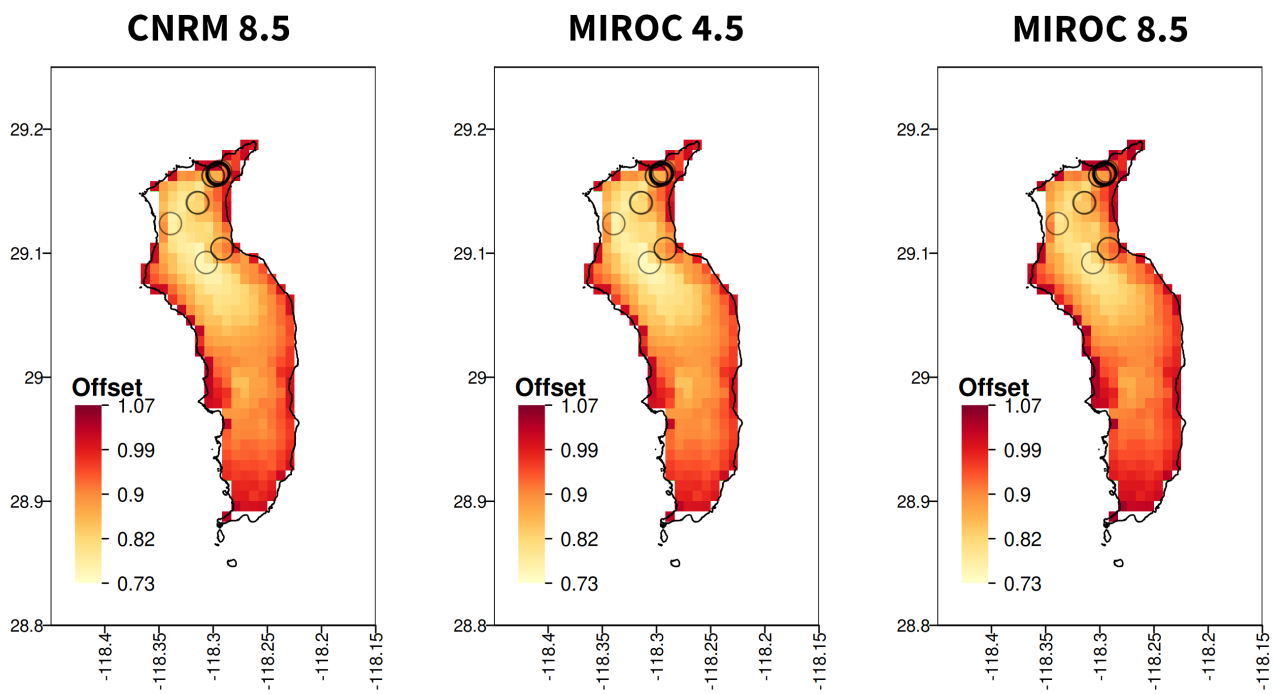


**Figure S14**. Map of the genomic offset for Guadalupe Island under three additional climate scenarios (CNRM 8.5, MIROC 4.5, and MIROC 8.5) as in Figure 5B. Genomic offset is the predicted difference in the current genetic composition and that which would be ideal under future climate conditions. Darker red indicates a greater likelihood of future maladaptation. Points indicate sample locations.
