## Supplementary Rmarkdown Outputs for "Comparison of conservation strategies for California Channel Island Oak (*Quercus tomentella*) using climate suitability predicted from genomic data": gradient_forest_candidate_SNPs_Channel_islands.html


### Gradient Forest - Candidate SNPs

###### Alayna Mead

#### 2024-05-14

- 1 Setup
  - 1.1 Libraries
  - 1.2 Input and setup data
  - 1.3 Calculate MEM variables
- 2 Run Gradient Forest
- 3 Setup for mapping
  - 3.1 Load climate rasters
  - 3.2 Mapping functions
- 4 Mapping
  - 4.1 Genomic turnover
  - 4.2 Climate PCA
  - 4.3 Genomic Offsets
- 5 Offset calculations
  - 5.1 Offsets for specific populations,
    without movement
  - 5.2 Offsets with assisted gene
    flow
- 6 Climate suitability

### 1 Setup

#### 1.1 Libraries

```
library(gradientForest)
library(psych) # for pairs.panels()
library(vegan) # for pcnm()
```

```
## Loading required package: permute
```

```
## Loading required package: lattice
```

```
## This is vegan 2.6-4
```

```
library(RColorBrewer) # colors
library(sf) # spatial functions
```

```
## Linking to GEOS 3.12.0, GDAL 3.8.4, PROJ 9.3.1; sf_use_s2() is TRUE
```

```
library(terra) # spatial functions
```

```
## terra 1.7.55
```

```
## 
## Attaching package: 'terra'
```

```
## The following objects are masked from 'package:psych':
## 
##     describe, distance, rescale
```

```
library(ComplexHeatmap)
```

```
## Loading required package: grid
```

```
## 
## Attaching package: 'grid'
```

```
## The following object is masked from 'package:terra':
## 
##     depth
```

```
## ========================================
## ComplexHeatmap version 2.16.0
## Bioconductor page: http://bioconductor.org/packages/ComplexHeatmap/
## Github page: https://github.com/jokergoo/ComplexHeatmap
## Documentation: http://jokergoo.github.io/ComplexHeatmap-reference
## 
## If you use it in published research, please cite either one:
## - Gu, Z. Complex Heatmap Visualization. iMeta 2022.
## - Gu, Z. Complex heatmaps reveal patterns and correlations in multidimensional 
##     genomic data. Bioinformatics 2016.
## 
## 
## The new InteractiveComplexHeatmap package can directly export static 
## complex heatmaps into an interactive Shiny app with zero effort. Have a try!
## 
## This message can be suppressed by:
##   suppressPackageStartupMessages(library(ComplexHeatmap))
## ========================================
```

```
## 
## Attaching package: 'ComplexHeatmap'
```

```
## The following object is masked from 'package:terra':
## 
##     draw
```

```
sessionInfo()
```

```
## R version 4.3.3 (2024-02-29)
## Platform: x86_64-pc-linux-gnu (64-bit)
## Running under: Arch Linux
## 
## Matrix products: default
## BLAS:   /usr/lib/libblas.so.3.12.0 
## LAPACK: /usr/lib/liblapack.so.3.12.0
## 
## locale:
##  [1] LC_CTYPE=en_US.UTF-8       LC_NUMERIC=C              
##  [3] LC_TIME=en_US.UTF-8        LC_COLLATE=en_US.UTF-8    
##  [5] LC_MONETARY=en_US.UTF-8    LC_MESSAGES=en_US.UTF-8   
##  [7] LC_PAPER=en_US.UTF-8       LC_NAME=C                 
##  [9] LC_ADDRESS=C               LC_TELEPHONE=C            
## [11] LC_MEASUREMENT=en_US.UTF-8 LC_IDENTIFICATION=C       
## 
## time zone: America/New_York
## tzcode source: system (glibc)
## 
## attached base packages:
## [1] grid      stats     graphics  grDevices utils     datasets  methods  
## [8] base     
## 
## other attached packages:
## [1] ComplexHeatmap_2.16.0 terra_1.7-55          sf_1.0-14            
## [4] RColorBrewer_1.1-3    vegan_2.6-4           lattice_0.22-5       
## [7] permute_0.9-7         psych_2.3.6           gradientForest_0.1-36
## 
## loaded via a namespace (and not attached):
##  [1] shape_1.4.6          circlize_0.4.15      rjson_0.2.21        
##  [4] xfun_0.39            bslib_0.4.2          GlobalOptions_0.1.2 
##  [7] vctrs_0.6.2          tools_4.3.3          generics_0.1.3      
## [10] stats4_4.3.3         parallel_4.3.3       tibble_3.2.1        
## [13] proxy_0.4-27         fansi_1.0.4          cluster_2.1.6       
## [16] pkgconfig_2.0.3      Matrix_1.6-5         KernSmooth_2.23-22  
## [19] S4Vectors_0.38.2     lifecycle_1.0.3      compiler_4.3.3      
## [22] mnormt_2.1.1         codetools_0.2-19     clue_0.3-65         
## [25] htmltools_0.5.5      class_7.3-22         sass_0.4.5          
## [28] yaml_2.3.7           pillar_1.9.0         crayon_1.5.2        
## [31] jquerylib_0.1.4      MASS_7.3-60.0.1      classInt_0.4-9      
## [34] cachem_1.0.7         iterators_1.0.14     foreach_1.5.2       
## [37] nlme_3.1-164         tidyselect_1.2.0     digest_0.6.31       
## [40] dplyr_1.1.2          splines_4.3.3        fastmap_1.1.1       
## [43] colorspace_2.1-0     cli_3.6.1            magrittr_2.0.3      
## [46] utf8_1.2.3           e1071_1.7-13         rmarkdown_2.21      
## [49] matrixStats_1.2.0    png_0.1-8            GetoptLong_1.0.5    
## [52] evaluate_0.20        knitr_1.42           IRanges_2.34.1      
## [55] doParallel_1.0.17    mgcv_1.9-1           rlang_1.1.0         
## [58] Rcpp_1.0.10          glue_1.6.2           DBI_1.1.3           
## [61] BiocGenerics_0.46.0  extendedForest_1.6.1 rstudioapi_0.14     
## [64] jsonlite_1.8.4       R6_2.5.1             units_0.8-3
```

#### 1.2 Input and setup data

`clim` is a large dataframe with all climate variables
(bioclim, basin characterization model, future projections) for each
sample collected

`snps` is a dataframe of imputed SNPs (rows are samples,
columns are SNPs, values are 0, 1, or 2 for the number of copies of the
variant). SNPs are filtered, LD-pruned, and biallelic, missing data was
imputed in RDA script because gradient forest doesn’t allow NAs.

`snps.cand` is the imputed snps subset to candidate SNPs
produced from the RDA.

```
# climate data
load('data/clean/climate_locality_data_BCM_and_bioclim.rda')
clim <- as.data.frame(clim.all)
rm(clim.all)
str(clim)
```

```
## 'data.frame':    295 obs. of  149 variables:
##  $ original_row_order                : int  1 2 3 4 5 6 7 8 9 10 ...
##  $ ID_field                          : chr  "QUTO-I-SB-01" "QUCH-I-SB-02" "QUCH-I-SB-03" "QUTO-I-SB-04" ...
##  $ ID_seq                            : chr  "Qtom.I.SB.01" "Qchr.I.SB.02" "Qchr.I.SB.03" "Qtom.I.SB.04" ...
##  $ species                           : chr  "Quercus tomentella" "Quercus chrysolepis" "Quercus chrysolepis" "Quercus tomentella" ...
##  $ collection_date                   : chr  "2020-11-19" "2020-11-19" "2020-11-19" "2020-11-19" ...
##  $ collector                         : chr  "John Knapp" "John Knapp" "John Knapp" "John Knapp" ...
##  $ batch_ID                          : int  21 21 21 21 21 21 21 21 21 21 ...
##  $ island                            : Factor w/ 7 levels "Mainland","Santa Rosa Island",..: 3 3 3 3 3 3 3 3 3 3 ...
##  $ locality                          : chr  "Christy Pines" "west of Diablo Pk." "west of Diablo Pk." "north slope northewest of Diablo Pk." ...
##  $ elev_ft                           : int  1432 2077 2392 2369 1963 NA 1317 1352 1187 NA ...
##  $ elev_m                            : num  NA NA NA NA NA NA NA NA NA NA ...
##  $ habitat                           : chr  "island woodland / pine forest" "QUCH woodland" "QUCH woodland" "island woodland / oak woodland" ...
##  $ other_species                     : chr  "" "" "" "" ...
##  $ tree_description                  : chr  "single tree, ~20 ft tall" "multi-trunked tree, ~17 ft tall" "multi-trunked tree, 15 ft tall" "50 ft large single trunk tree" ...
##  $ other_notes                       : chr  "" "" "" "possible hybrid with QUCH" ...
##  $ ID_vcf                            : chr  "Qtom.I.SB.01" "Qchr.I.SB.02" "Qchr.I.SB.03" "Qtom.I.SB.04" ...
##  $ island_id                         : Factor w/ 306 levels "Mainland.Qchr.A.232",..: 151 145 146 152 66 67 68 69 153 154 ...
##  $ aet2070_2099_ave_CCSM4_rcp85      : num  341 284 290 290 292 ...
##  $ aprpck2070_2099_ave_CCSM4_rcp85   : num  0 0 0 0 0 0 0 0 0 0 ...
##  $ cwd2070_2099_ave_CCSM4_rcp85      : num  902 920 931 931 938 ...
##  $ pet2070_2099_ave_CCSM4_rcp85      : num  1243 1205 1221 1221 1230 ...
##  $ ppt2070_2099_ave_CCSM4_rcp85      : num  607 643 643 643 643 ...
##  $ rch2070_2099_ave_CCSM4_rcp85      : num  99.42 6.28 6.14 6.14 6.01 ...
##  $ run2070_2099_ave_CCSM4_rcp85      : num  167 353 347 347 345 ...
##  $ tmn2070_2099_ave_CCSM4_rcp85      : num  10.5 10.3 10.2 10.2 10.3 ...
##  $ tmx2070_2099_ave_CCSM4_rcp85      : num  22.4 21.4 21.4 21.4 21.6 ...
##  $ aet2070_2099_ave_CNRM_rcp85       : num  367 310 316 316 319 ...
##  $ aprpck2070_2099_ave_CNRM_rcp85    : num  0 0 0 0 0 0 0 0 0 0 ...
##  $ cwd2070_2099_ave_CNRM_rcp85       : num  885 904 914 914 920 ...
##  $ pet2070_2099_ave_CNRM_rcp85       : num  1252 1214 1230 1230 1239 ...
##  $ ppt2070_2099_ave_CNRM_rcp85       : num  797 845 845 845 845 ...
##  $ rch2070_2099_ave_CNRM_rcp85       : num  106.25 7.31 7.07 7.07 7 ...
##  $ run2070_2099_ave_CNRM_rcp85       : num  324 528 522 522 520 ...
##  $ tmn2070_2099_ave_CNRM_rcp85       : num  11 10.7 10.7 10.7 10.7 ...
##  $ tmx2070_2099_ave_CNRM_rcp85       : num  22.8 21.7 21.7 21.7 21.9 ...
##  $ aet2070_2099_ave_Fgoals_rcp85     : num  286 237 242 242 244 ...
##  $ aprpck2070_2099_ave_Fgoals_rcp85  : num  0 0 0 0 0 0 0 0 0 0 ...
##  $ cwd2070_2099_ave_Fgoals_rcp85     : num  974 985 997 997 1004 ...
##  $ pet2070_2099_ave_Fgoals_rcp85     : num  1260 1222 1239 1239 1248 ...
##  $ ppt2070_2099_ave_Fgoals_rcp85     : num  384 407 407 407 407 ...
##  $ rch2070_2099_ave_Fgoals_rcp85     : num  58.01 4.89 4.75 4.75 4.67 ...
##  $ run2070_2099_ave_Fgoals_rcp85     : num  40.5 165.1 160.7 160.7 158.9 ...
##  $ tmn2070_2099_ave_Fgoals_rcp85     : num  11.1 10.9 10.8 10.8 10.8 ...
##  $ tmx2070_2099_ave_Fgoals_rcp85     : num  23.3 22.3 22.2 22.2 22.4 ...
##  $ aet1951_1980_ave_HST_1583635268   : num  336 276 281 281 284 ...
##  $ aprpck1951_1980_ave_HST_1583635303: num  0 0 0 0 0 0 0 0 0 0 ...
##  $ cwd1951_1980_ave_HST_1583635273   : num  810 832 842 842 850 ...
##  $ pet1951_1980_ave_HST_1583635277   : num  1149 1110 1126 1126 1137 ...
##  $ ppt1951_1980_ave_HST_1583635286   : num  553 589 591 591 587 ...
##  $ rch1951_1980_ave_HST_1583635295   : num  89.7 6.36 6.34 6.34 6.24 ...
##  $ run1951_1980_ave_HST_1583635299   : num  122 303 300 300 294 ...
##  $ tmn1951_1980_ave_HST_1583635311   : num  6.95 7.28 7.25 7.25 7.42 ...
##  $ tmx1951_1980_ave_HST_1583635306   : num  19.5 17.9 17.9 17.9 18.2 ...
##  $ aet2070_2099_ave_IPSL_rcp85       : num  306 258 263 263 265 ...
##  $ aprpck2070_2099_ave_IPSL_rcp85    : num  0 0 0 0 0 0 0 0 0 0 ...
##  $ cwd2070_2099_ave_IPSL_rcp85       : num  979 988 1000 1000 1007 ...
##  $ pet2070_2099_ave_IPSL_rcp85       : num  1285 1246 1264 1264 1273 ...
##  $ ppt2070_2099_ave_IPSL_rcp85       : num  577 611 611 611 611 ...
##  $ rch2070_2099_ave_IPSL_rcp85       : num  81.84 5.2 5.2 5.2 5.19 ...
##  $ run2070_2099_ave_IPSL_rcp85       : num  189 348 343 343 341 ...
##  $ tmn2070_2099_ave_IPSL_rcp85       : num  12.7 12.4 12.3 12.3 12.4 ...
##  $ tmx2070_2099_ave_IPSL_rcp85       : num  24.1 23.1 23.1 23.1 23.3 ...
##  $ aet2070_2099_ave_MIROC_rcp45      : num  338 289 293 293 296 ...
##  $ aprpck2070_2099_ave_MIROC_rcp45   : num  0 0 0 0 0 0 0 0 0 0 ...
##  $ cwd2070_2099_ave_MIROC_rcp45      : num  906 916 928 928 935 ...
##  $ pet2070_2099_ave_MIROC_rcp45      : num  1244 1205 1222 1222 1231 ...
##  $ ppt2070_2099_ave_MIROC_rcp45      : num  437 462 462 462 462 ...
##  $ rch2070_2099_ave_MIROC_rcp45      : num  53.46 5.34 5.21 5.21 5.09 ...
##  $ run2070_2099_ave_MIROC_rcp45      : num  46.1 168.1 163.5 163.5 161.3 ...
##  $ tmn2070_2099_ave_MIROC_rcp45      : num  10.4 10.2 10.1 10.1 10.2 ...
##  $ tmx2070_2099_ave_MIROC_rcp45      : num  22.6 21.5 21.5 21.5 21.7 ...
##  $ aet2070_2099_ave_MIROC_rcp85      : num  297 251 255 255 257 ...
##  $ aprpck2070_2099_ave_MIROC_rcp85   : num  0 0 0 0 0 0 0 0 0 0 ...
##  $ cwd2070_2099_ave_MIROC_rcp85      : num  988 995 1008 1008 1016 ...
##  $ pet2070_2099_ave_MIROC_rcp85      : num  1285 1246 1263 1263 1272 ...
##  $ ppt2070_2099_ave_MIROC_rcp85      : num  392 415 415 415 415 ...
##  $ rch2070_2099_ave_MIROC_rcp85      : num  52.41 4 3.96 3.96 3.92 ...
##  $ run2070_2099_ave_MIROC_rcp85      : num  43.1 159.8 156 156 154.3 ...
##  $ tmn2070_2099_ave_MIROC_rcp85      : num  12.2 11.9 11.9 11.9 11.9 ...
##  $ tmx2070_2099_ave_MIROC_rcp85      : num  24.4 23.3 23.3 23.3 23.5 ...
##  $ aet2070_2099_ave_MPI_rcp45        : num  356 297 304 304 306 ...
##  $ aprpck2070_2099_ave_MPI_rcp45     : num  0 0 0 0 0 0 0 0 0 0 ...
##  $ cwd2070_2099_ave_MPI_rcp45        : num  864 884 893 893 900 ...
##  $ pet2070_2099_ave_MPI_rcp45        : num  1219 1181 1197 1197 1206 ...
##  $ ppt2070_2099_ave_MPI_rcp45        : num  617 654 654 654 654 ...
##  $ rch2070_2099_ave_MPI_rcp45        : num  104.63 7.51 7.37 7.37 7.31 ...
##  $ run2070_2099_ave_MPI_rcp45        : num  157 349 343 343 340 ...
##  $ tmn2070_2099_ave_MPI_rcp45        : num  9.49 9.24 9.16 9.16 9.22 ...
##  $ tmx2070_2099_ave_MPI_rcp45        : num  21.4 20.4 20.4 20.4 20.5 ...
##  $ bio1                              : num  14.6 14.2 14.2 14.2 14.3 ...
##  $ bio2                              : num  11.4 12 12 12 11.9 ...
##  $ bio3                              : num  64.6 62 62 62 63.4 ...
##  $ bio4                              : num  247 282 282 282 269 ...
##  $ bio5                              : num  24.3 25.1 25.1 25.1 24.7 ...
##  $ bio6                              : num  6.6 5.7 5.7 5.7 6 ...
##  $ bio7                              : num  17.7 19.4 19.4 19.4 18.7 ...
##  $ bio8                              : num  12.2 11.4 11.4 11.4 11.6 ...
##  $ bio9                              : num  16.6 16.8 16.8 16.8 16.7 ...
##  $ bio10                             : num  17.8 17.9 17.9 17.9 17.8 ...
##   [list output truncated]
```

```
# imputed SNPs
# file made in RDA script
# named 'imp', rename to 'snps'
load('data/clean/Qtom107.Qchr17.Qssp3.20220906.qlob.ef.repeatsOut.renamedChrsVars.biallelicSNPs.meanDP5.genoDP5.MAF0.01.missing0.9.ldPruned.additive_imputed.rda')
snps <- imp
rm(imp)


# get candidate SNPs from RDA
# named 'cand' or 'cand.sort'

# version for candidate SNPs determined without Guadalupe
# load('results/redundancy_analysis/RDA_candidate_climate_SNP_table_noGuad.rda')
# rownames(cand.sort) <- cand.sort$snp
# str(cand.sort)
# cand <- cand.sort

# version with candidate SNPs determined without Guadalupe or Anacapa
load('results/redundancy_analysis/RDA_candidate_climate_SNP_table_noGuadAna.rda')
rownames(cand) <- cand$snp
str(cand)
```

```
## 'data.frame':    560 obs. of  11 variables:
##  $ axis       : num  1 1 1 1 1 1 1 1 1 1 ...
##  $ snp        : chr  "1_388897_C_T_T" "1_2592491_G_T_T" "1_10042008_T_A_A" "1_16403258_G_A_G" ...
##  $ loading    : num  0.0836 0.0893 0.069 -0.0677 0.0635 ...
##  $ bio5       : num  -0.728 -0.75 -0.627 0.612 -0.549 ...
##  $ bio6       : num  -0.418 -0.625 -0.384 0.434 -0.343 ...
##  $ bio15      : num  -0.685 -0.629 -0.317 0.415 -0.567 ...
##  $ bio18      : num  0.76 0.822 0.551 -0.568 0.594 ...
##  $ bio19      : num  0.02654 0.2577 0.14488 -0.00857 0.01968 ...
##  $ elev       : num  -0.248 -0.135 -0.234 0.158 -0.148 ...
##  $ predictor  : chr  "bio18" "bio18" "bio5" "bio5" ...
##  $ correlation: num  0.76 0.822 0.627 0.612 0.594 ...
```

```
snps <-  as.matrix(snps)
snps[1:10, 1:10]
```

```
##                1_39789_A_G_G 1_52176_G_A_A 1_52195_G_A_A 1_52213_C_T_T
## Qchr.A.232                 1             0             0             1
## Qchr.A.275                 1             0             0             0
## Qchr.A.ED.102              1             0             0             0
## Qchr.A.ELD.53              1             0             0             1
## Qchr.A.KER.16              1             0             0             0
## Qchr.A.LA.02               1             0             0             0
## Qchr.A.LA.215              1             0             0             1
## Qchr.A.LA.222              1             0             0             0
## Qchr.A.SIS.309             0             0             0             0
## Qchr.A.PLU.65              1             0             0             0
##                1_52248_T_C_T 1_52331_G_T_T 1_53841_G_A_A 1_53940_T_C_T
## Qchr.A.232                 0             0             0             1
## Qchr.A.275                 0             0             0             1
## Qchr.A.ED.102              0             0             1             0
## Qchr.A.ELD.53              0             0             0             1
## Qchr.A.KER.16              0             1             0             1
## Qchr.A.LA.02               0             0             0             1
## Qchr.A.LA.215              0             0             0             1
## Qchr.A.LA.222              0             0             0             1
## Qchr.A.SIS.309             0             0             0             1
## Qchr.A.PLU.65              0             0             0             1
##                1_53944_A_G_A 1_53966_G_A_A
## Qchr.A.232                 0             0
## Qchr.A.275                 0             0
## Qchr.A.ED.102              0             0
## Qchr.A.ELD.53              0             0
## Qchr.A.KER.16              0             0
## Qchr.A.LA.02               1             0
## Qchr.A.LA.215              0             0
## Qchr.A.LA.222              0             0
## Qchr.A.SIS.309             1             0
## Qchr.A.PLU.65              1             0
```

```
# first get rid of the individuals without coord data
snps <- snps[rownames(snps) %in% clim$ID_vcf,]

# now get rid of the individuals without SNP data
# also reorder to match snp order
clim <- clim[match(rownames(snps), clim$ID_vcf),]


# check that rows are in same order
cbind(rownames(clim), rownames(snps))
```

```
##        [,1]             [,2]            
##   [1,] "Qchr.A.232"     "Qchr.A.232"    
##   [2,] "Qchr.A.275"     "Qchr.A.275"    
##   [3,] "Qchr.A.ED.102"  "Qchr.A.ED.102" 
##   [4,] "Qchr.A.ELD.53"  "Qchr.A.ELD.53" 
##   [5,] "Qchr.A.KER.16"  "Qchr.A.KER.16" 
##   [6,] "Qchr.A.LA.02"   "Qchr.A.LA.02"  
##   [7,] "Qchr.A.LA.215"  "Qchr.A.LA.215" 
##   [8,] "Qchr.A.LA.222"  "Qchr.A.LA.222" 
##   [9,] "Qchr.A.PLU.65"  "Qchr.A.PLU.65" 
##  [10,] "Qchr.A.SBD.83"  "Qchr.A.SBD.83" 
##  [11,] "Qchr.I.LA.20"   "Qchr.I.LA.20"  
##  [12,] "Qchr.I.LA.25"   "Qchr.I.LA.25"  
##  [13,] "Qchr.I.SB.02"   "Qchr.I.SB.02"  
##  [14,] "Qchr.I.SB.03"   "Qchr.I.SB.03"  
##  [15,] "Qchr.JJK.SB.13" "Qchr.JJK.SB.13"
##  [16,] "Qchr.JR.SB.01"  "Qchr.JR.SB.01" 
##  [17,] "Qtom.A.LA.169"  "Qtom.A.LA.169" 
##  [18,] "Qtom.A.LA.175"  "Qtom.A.LA.175" 
##  [19,] "Qtom.A.LA.176"  "Qtom.A.LA.176" 
##  [20,] "Qtom.A.LA.180"  "Qtom.A.LA.180" 
##  [21,] "Qtom.A.LA.181"  "Qtom.A.LA.181" 
##  [22,] "Qtom.A.LA.183"  "Qtom.A.LA.183" 
##  [23,] "Qtom.A.LA.190"  "Qtom.A.LA.190" 
##  [24,] "Qtom.A.LA.191"  "Qtom.A.LA.191" 
##  [25,] "Qtom.A.LA.198"  "Qtom.A.LA.198" 
##  [26,] "Qtom.A.LA.200"  "Qtom.A.LA.200" 
##  [27,] "Qtom.A.LA.204"  "Qtom.A.LA.204" 
##  [28,] "Qtom.A.LA.207"  "Qtom.A.LA.207" 
##  [29,] "Qtom.A.LA.208"  "Qtom.A.LA.208" 
##  [30,] "Qtom.A.LA.209"  "Qtom.A.LA.209" 
##  [31,] "Qtom.A.LA.218"  "Qtom.A.LA.218" 
##  [32,] "Qtom.A.LA.221"  "Qtom.A.LA.221" 
##  [33,] "Qtom.A.SB.324"  "Qtom.A.SB.324" 
##  [34,] "Qtom.A.SB.325"  "Qtom.A.SB.325" 
##  [35,] "Qtom.A.SB.327"  "Qtom.A.SB.327" 
##  [36,] "Qtom.A.SB.329"  "Qtom.A.SB.329" 
##  [37,] "Qtom.A.SB.330"  "Qtom.A.SB.330" 
##  [38,] "Qtom.A.SB.331"  "Qtom.A.SB.331" 
##  [39,] "Qtom.A.SB.332"  "Qtom.A.SB.332" 
##  [40,] "Qtom.A.SB.333"  "Qtom.A.SB.333" 
##  [41,] "Qtom.A.SB.335"  "Qtom.A.SB.335" 
##  [42,] "Qtom.A.SB.336"  "Qtom.A.SB.336" 
##  [43,] "Qtom.A.SB.337"  "Qtom.A.SB.337" 
##  [44,] "Qtom.A.SB.338"  "Qtom.A.SB.338" 
##  [45,] "Qtom.A.SB.339"  "Qtom.A.SB.339" 
##  [46,] "Qtom.A.SB.341"  "Qtom.A.SB.341" 
##  [47,] "Qtom.A.SB.342"  "Qtom.A.SB.342" 
##  [48,] "Qtom.A.SB.343"  "Qtom.A.SB.343" 
##  [49,] "Qtom.A.SB.344"  "Qtom.A.SB.344" 
##  [50,] "Qtom.A.SB.345"  "Qtom.A.SB.345" 
##  [51,] "Qtom.A.SB.346"  "Qtom.A.SB.346" 
##  [52,] "Qtom.A.SB.347"  "Qtom.A.SB.347" 
##  [53,] "Qtom.A.SB.348"  "Qtom.A.SB.348" 
##  [54,] "Qtom.A.SB.352"  "Qtom.A.SB.352" 
##  [55,] "Qtom.A.SB.353"  "Qtom.A.SB.353" 
##  [56,] "Qtom.A.SB.355"  "Qtom.A.SB.355" 
##  [57,] "Qtom.I.BC.10"   "Qtom.I.BC.10"  
##  [58,] "Qtom.I.BC.11"   "Qtom.I.BC.11"  
##  [59,] "Qtom.I.BC.12"   "Qtom.I.BC.12"  
##  [60,] "Qtom.I.BC.13"   "Qtom.I.BC.13"  
##  [61,] "Qtom.I.BC.18"   "Qtom.I.BC.18"  
##  [62,] "Qtom.I.BC.20"   "Qtom.I.BC.20"  
##  [63,] "Qtom.I.BC.21"   "Qtom.I.BC.21"  
##  [64,] "Qtom.I.BC.23"   "Qtom.I.BC.23"  
##  [65,] "Qtom.I.BC.24"   "Qtom.I.BC.24"  
##  [66,] "Qtom.I.BC.32"   "Qtom.I.BC.32"  
##  [67,] "Qtom.I.BC.33"   "Qtom.I.BC.33"  
##  [68,] "Qtom.I.BC.35"   "Qtom.I.BC.35"  
##  [69,] "Qtom.I.BC.39"   "Qtom.I.BC.39"  
##  [70,] "Qtom.I.BC.4"    "Qtom.I.BC.4"   
##  [71,] "Qtom.I.BC.40"   "Qtom.I.BC.40"  
##  [72,] "Qtom.I.BC.41"   "Qtom.I.BC.41"  
##  [73,] "Qtom.I.BC.42"   "Qtom.I.BC.42"  
##  [74,] "Qtom.I.BC.43"   "Qtom.I.BC.43"  
##  [75,] "Qtom.I.BC.44"   "Qtom.I.BC.44"  
##  [76,] "Qtom.I.BC.45"   "Qtom.I.BC.45"  
##  [77,] "Qtom.I.BC.46"   "Qtom.I.BC.46"  
##  [78,] "Qtom.I.BC.48"   "Qtom.I.BC.48"  
##  [79,] "Qtom.I.LA.05"   "Qtom.I.LA.05"  
##  [80,] "Qtom.I.LA.06"   "Qtom.I.LA.06"  
##  [81,] "Qtom.I.LA.08"   "Qtom.I.LA.08"  
##  [82,] "Qtom.I.LA.10"   "Qtom.I.LA.10"  
##  [83,] "Qtom.I.LA.11"   "Qtom.I.LA.11"  
##  [84,] "Qtom.I.LA.12"   "Qtom.I.LA.12"  
##  [85,] "Qtom.I.LA.13"   "Qtom.I.LA.13"  
##  [86,] "Qtom.I.LA.21"   "Qtom.I.LA.21"  
##  [87,] "Qtom.I.LA.22"   "Qtom.I.LA.22"  
##  [88,] "Qtom.I.LA.23"   "Qtom.I.LA.23"  
##  [89,] "Qtom.I.LA.24"   "Qtom.I.LA.24"  
##  [90,] "Qtom.I.SB.01"   "Qtom.I.SB.01"  
##  [91,] "Qtom.I.SB.04"   "Qtom.I.SB.04"  
##  [92,] "Qtom.I.SB.09"   "Qtom.I.SB.09"  
##  [93,] "Qtom.I.SB.10"   "Qtom.I.SB.10"  
##  [94,] "Qtom.I.SB.11"   "Qtom.I.SB.11"  
##  [95,] "Qtom.I.SB.12"   "Qtom.I.SB.12"  
##  [96,] "Qtom.I.SB.28"   "Qtom.I.SB.28"  
##  [97,] "Qtom.I.SB.33"   "Qtom.I.SB.33"  
##  [98,] "Qtom.I.SB.35"   "Qtom.I.SB.35"  
##  [99,] "Qtom.I.SB.36"   "Qtom.I.SB.36"  
## [100,] "Qtom.I.SB.37"   "Qtom.I.SB.37"  
## [101,] "Qtom.I.SB.38"   "Qtom.I.SB.38"  
## [102,] "Qtom.I.SB.40"   "Qtom.I.SB.40"  
## [103,] "Qtom.I.SB.41"   "Qtom.I.SB.41"  
## [104,] "Qtom.I.SB.42"   "Qtom.I.SB.42"  
## [105,] "Qtom.I.SB.44"   "Qtom.I.SB.44"  
## [106,] "Qtom.I.SB.45"   "Qtom.I.SB.45"  
## [107,] "Qtom.I.SB.73"   "Qtom.I.SB.73"  
## [108,] "Qtom.I.SB.74"   "Qtom.I.SB.74"  
## [109,] "Qtom.I.VEN.01"  "Qtom.I.VEN.01" 
## [110,] "Qtom.I.VEN.02"  "Qtom.I.VEN.02" 
## [111,] "Qtom.I.VEN.03"  "Qtom.I.VEN.03" 
## [112,] "Qtom.I.VEN.04"  "Qtom.I.VEN.04" 
## [113,] "Qtom.I.VEN.05"  "Qtom.I.VEN.05" 
## [114,] "Qtom.JJK.SB.01" "Qtom.JJK.SB.01"
## [115,] "Qtom.JJK.SB.03" "Qtom.JJK.SB.03"
## [116,] "Qtom.JJK.SB.07" "Qtom.JJK.SB.07"
## [117,] "Qtom.JJK.SB.08" "Qtom.JJK.SB.08"
## [118,] "Qtom.JJK.SB.09" "Qtom.JJK.SB.09"
## [119,] "Qtom.JJK.SB.10" "Qtom.JJK.SB.10"
## [120,] "Qtom.JR.SB.01"  "Qtom.JR.SB.01" 
## [121,] "Qtom.S.LA.186"  "Qtom.S.LA.186"
```

```
sum(rownames(clim) != rownames(snps))
```

```
## [1] 0
```

```
# rename SNP rownames to the same as clim rownames
rownames(snps) <- rownames(clim[clim$ID_vcf == rownames(snps),])

# subset imputed snps to just the candidate SNPs
snps.cand <- snps[, colnames(snps) %in% cand$snp]
dim(snps.cand) # [1]   93 560
```

```
## [1] 121 560
```

```
# rename columns - gf complains about it starting with a number
colnames(snps.cand) <- make.names(colnames(snps.cand))

# check again
cbind(rownames(clim), rownames(snps))
```

```
##        [,1]             [,2]            
##   [1,] "Qchr.A.232"     "Qchr.A.232"    
##   [2,] "Qchr.A.275"     "Qchr.A.275"    
##   [3,] "Qchr.A.ED.102"  "Qchr.A.ED.102" 
##   [4,] "Qchr.A.ELD.53"  "Qchr.A.ELD.53" 
##   [5,] "Qchr.A.KER.16"  "Qchr.A.KER.16" 
##   [6,] "Qchr.A.LA.02"   "Qchr.A.LA.02"  
##   [7,] "Qchr.A.LA.215"  "Qchr.A.LA.215" 
##   [8,] "Qchr.A.LA.222"  "Qchr.A.LA.222" 
##   [9,] "Qchr.A.PLU.65"  "Qchr.A.PLU.65" 
##  [10,] "Qchr.A.SBD.83"  "Qchr.A.SBD.83" 
##  [11,] "Qchr.I.LA.20"   "Qchr.I.LA.20"  
##  [12,] "Qchr.I.LA.25"   "Qchr.I.LA.25"  
##  [13,] "Qchr.I.SB.02"   "Qchr.I.SB.02"  
##  [14,] "Qchr.I.SB.03"   "Qchr.I.SB.03"  
##  [15,] "Qchr.JJK.SB.13" "Qchr.JJK.SB.13"
##  [16,] "Qchr.JR.SB.01"  "Qchr.JR.SB.01" 
##  [17,] "Qtom.A.LA.169"  "Qtom.A.LA.169" 
##  [18,] "Qtom.A.LA.175"  "Qtom.A.LA.175" 
##  [19,] "Qtom.A.LA.176"  "Qtom.A.LA.176" 
##  [20,] "Qtom.A.LA.180"  "Qtom.A.LA.180" 
##  [21,] "Qtom.A.LA.181"  "Qtom.A.LA.181" 
##  [22,] "Qtom.A.LA.183"  "Qtom.A.LA.183" 
##  [23,] "Qtom.A.LA.190"  "Qtom.A.LA.190" 
##  [24,] "Qtom.A.LA.191"  "Qtom.A.LA.191" 
##  [25,] "Qtom.A.LA.198"  "Qtom.A.LA.198" 
##  [26,] "Qtom.A.LA.200"  "Qtom.A.LA.200" 
##  [27,] "Qtom.A.LA.204"  "Qtom.A.LA.204" 
##  [28,] "Qtom.A.LA.207"  "Qtom.A.LA.207" 
##  [29,] "Qtom.A.LA.208"  "Qtom.A.LA.208" 
##  [30,] "Qtom.A.LA.209"  "Qtom.A.LA.209" 
##  [31,] "Qtom.A.LA.218"  "Qtom.A.LA.218" 
##  [32,] "Qtom.A.LA.221"  "Qtom.A.LA.221" 
##  [33,] "Qtom.A.SB.324"  "Qtom.A.SB.324" 
##  [34,] "Qtom.A.SB.325"  "Qtom.A.SB.325" 
##  [35,] "Qtom.A.SB.327"  "Qtom.A.SB.327" 
##  [36,] "Qtom.A.SB.329"  "Qtom.A.SB.329" 
##  [37,] "Qtom.A.SB.330"  "Qtom.A.SB.330" 
##  [38,] "Qtom.A.SB.331"  "Qtom.A.SB.331" 
##  [39,] "Qtom.A.SB.332"  "Qtom.A.SB.332" 
##  [40,] "Qtom.A.SB.333"  "Qtom.A.SB.333" 
##  [41,] "Qtom.A.SB.335"  "Qtom.A.SB.335" 
##  [42,] "Qtom.A.SB.336"  "Qtom.A.SB.336" 
##  [43,] "Qtom.A.SB.337"  "Qtom.A.SB.337" 
##  [44,] "Qtom.A.SB.338"  "Qtom.A.SB.338" 
##  [45,] "Qtom.A.SB.339"  "Qtom.A.SB.339" 
##  [46,] "Qtom.A.SB.341"  "Qtom.A.SB.341" 
##  [47,] "Qtom.A.SB.342"  "Qtom.A.SB.342" 
##  [48,] "Qtom.A.SB.343"  "Qtom.A.SB.343" 
##  [49,] "Qtom.A.SB.344"  "Qtom.A.SB.344" 
##  [50,] "Qtom.A.SB.345"  "Qtom.A.SB.345" 
##  [51,] "Qtom.A.SB.346"  "Qtom.A.SB.346" 
##  [52,] "Qtom.A.SB.347"  "Qtom.A.SB.347" 
##  [53,] "Qtom.A.SB.348"  "Qtom.A.SB.348" 
##  [54,] "Qtom.A.SB.352"  "Qtom.A.SB.352" 
##  [55,] "Qtom.A.SB.353"  "Qtom.A.SB.353" 
##  [56,] "Qtom.A.SB.355"  "Qtom.A.SB.355" 
##  [57,] "Qtom.I.BC.10"   "Qtom.I.BC.10"  
##  [58,] "Qtom.I.BC.11"   "Qtom.I.BC.11"  
##  [59,] "Qtom.I.BC.12"   "Qtom.I.BC.12"  
##  [60,] "Qtom.I.BC.13"   "Qtom.I.BC.13"  
##  [61,] "Qtom.I.BC.18"   "Qtom.I.BC.18"  
##  [62,] "Qtom.I.BC.20"   "Qtom.I.BC.20"  
##  [63,] "Qtom.I.BC.21"   "Qtom.I.BC.21"  
##  [64,] "Qtom.I.BC.23"   "Qtom.I.BC.23"  
##  [65,] "Qtom.I.BC.24"   "Qtom.I.BC.24"  
##  [66,] "Qtom.I.BC.32"   "Qtom.I.BC.32"  
##  [67,] "Qtom.I.BC.33"   "Qtom.I.BC.33"  
##  [68,] "Qtom.I.BC.35"   "Qtom.I.BC.35"  
##  [69,] "Qtom.I.BC.39"   "Qtom.I.BC.39"  
##  [70,] "Qtom.I.BC.4"    "Qtom.I.BC.4"   
##  [71,] "Qtom.I.BC.40"   "Qtom.I.BC.40"  
##  [72,] "Qtom.I.BC.41"   "Qtom.I.BC.41"  
##  [73,] "Qtom.I.BC.42"   "Qtom.I.BC.42"  
##  [74,] "Qtom.I.BC.43"   "Qtom.I.BC.43"  
##  [75,] "Qtom.I.BC.44"   "Qtom.I.BC.44"  
##  [76,] "Qtom.I.BC.45"   "Qtom.I.BC.45"  
##  [77,] "Qtom.I.BC.46"   "Qtom.I.BC.46"  
##  [78,] "Qtom.I.BC.48"   "Qtom.I.BC.48"  
##  [79,] "Qtom.I.LA.05"   "Qtom.I.LA.05"  
##  [80,] "Qtom.I.LA.06"   "Qtom.I.LA.06"  
##  [81,] "Qtom.I.LA.08"   "Qtom.I.LA.08"  
##  [82,] "Qtom.I.LA.10"   "Qtom.I.LA.10"  
##  [83,] "Qtom.I.LA.11"   "Qtom.I.LA.11"  
##  [84,] "Qtom.I.LA.12"   "Qtom.I.LA.12"  
##  [85,] "Qtom.I.LA.13"   "Qtom.I.LA.13"  
##  [86,] "Qtom.I.LA.21"   "Qtom.I.LA.21"  
##  [87,] "Qtom.I.LA.22"   "Qtom.I.LA.22"  
##  [88,] "Qtom.I.LA.23"   "Qtom.I.LA.23"  
##  [89,] "Qtom.I.LA.24"   "Qtom.I.LA.24"  
##  [90,] "Qtom.I.SB.01"   "Qtom.I.SB.01"  
##  [91,] "Qtom.I.SB.04"   "Qtom.I.SB.04"  
##  [92,] "Qtom.I.SB.09"   "Qtom.I.SB.09"  
##  [93,] "Qtom.I.SB.10"   "Qtom.I.SB.10"  
##  [94,] "Qtom.I.SB.11"   "Qtom.I.SB.11"  
##  [95,] "Qtom.I.SB.12"   "Qtom.I.SB.12"  
##  [96,] "Qtom.I.SB.28"   "Qtom.I.SB.28"  
##  [97,] "Qtom.I.SB.33"   "Qtom.I.SB.33"  
##  [98,] "Qtom.I.SB.35"   "Qtom.I.SB.35"  
##  [99,] "Qtom.I.SB.36"   "Qtom.I.SB.36"  
## [100,] "Qtom.I.SB.37"   "Qtom.I.SB.37"  
## [101,] "Qtom.I.SB.38"   "Qtom.I.SB.38"  
## [102,] "Qtom.I.SB.40"   "Qtom.I.SB.40"  
## [103,] "Qtom.I.SB.41"   "Qtom.I.SB.41"  
## [104,] "Qtom.I.SB.42"   "Qtom.I.SB.42"  
## [105,] "Qtom.I.SB.44"   "Qtom.I.SB.44"  
## [106,] "Qtom.I.SB.45"   "Qtom.I.SB.45"  
## [107,] "Qtom.I.SB.73"   "Qtom.I.SB.73"  
## [108,] "Qtom.I.SB.74"   "Qtom.I.SB.74"  
## [109,] "Qtom.I.VEN.01"  "Qtom.I.VEN.01" 
## [110,] "Qtom.I.VEN.02"  "Qtom.I.VEN.02" 
## [111,] "Qtom.I.VEN.03"  "Qtom.I.VEN.03" 
## [112,] "Qtom.I.VEN.04"  "Qtom.I.VEN.04" 
## [113,] "Qtom.I.VEN.05"  "Qtom.I.VEN.05" 
## [114,] "Qtom.JJK.SB.01" "Qtom.JJK.SB.01"
## [115,] "Qtom.JJK.SB.03" "Qtom.JJK.SB.03"
## [116,] "Qtom.JJK.SB.07" "Qtom.JJK.SB.07"
## [117,] "Qtom.JJK.SB.08" "Qtom.JJK.SB.08"
## [118,] "Qtom.JJK.SB.09" "Qtom.JJK.SB.09"
## [119,] "Qtom.JJK.SB.10" "Qtom.JJK.SB.10"
## [120,] "Qtom.JR.SB.01"  "Qtom.JR.SB.01" 
## [121,] "Qtom.S.LA.186"  "Qtom.S.LA.186"
```

```
sum(rownames(clim) != rownames(snps))
```

```
## [1] 0
```

```
# remove Guadalupe and Mainland Qchr samples

keep <- clim$island %in% c("Santa Rosa Island", "Santa Cruz Island", "Anacapa Island", "Catalina Island", "San Clemente Island")
clim <- clim[keep,]
snps <- snps[keep,]
snps.cand <- snps.cand[keep,]

# check that rows are in same order
cbind(rownames(clim), rownames(snps))
```

```
##       [,1]             [,2]            
##  [1,] "Qchr.A.LA.215"  "Qchr.A.LA.215" 
##  [2,] "Qchr.A.LA.222"  "Qchr.A.LA.222" 
##  [3,] "Qchr.I.LA.20"   "Qchr.I.LA.20"  
##  [4,] "Qchr.I.LA.25"   "Qchr.I.LA.25"  
##  [5,] "Qchr.I.SB.02"   "Qchr.I.SB.02"  
##  [6,] "Qchr.I.SB.03"   "Qchr.I.SB.03"  
##  [7,] "Qchr.JJK.SB.13" "Qchr.JJK.SB.13"
##  [8,] "Qchr.JR.SB.01"  "Qchr.JR.SB.01" 
##  [9,] "Qtom.A.LA.169"  "Qtom.A.LA.169" 
## [10,] "Qtom.A.LA.175"  "Qtom.A.LA.175" 
## [11,] "Qtom.A.LA.176"  "Qtom.A.LA.176" 
## [12,] "Qtom.A.LA.180"  "Qtom.A.LA.180" 
## [13,] "Qtom.A.LA.181"  "Qtom.A.LA.181" 
## [14,] "Qtom.A.LA.183"  "Qtom.A.LA.183" 
## [15,] "Qtom.A.LA.190"  "Qtom.A.LA.190" 
## [16,] "Qtom.A.LA.191"  "Qtom.A.LA.191" 
## [17,] "Qtom.A.LA.198"  "Qtom.A.LA.198" 
## [18,] "Qtom.A.LA.200"  "Qtom.A.LA.200" 
## [19,] "Qtom.A.LA.204"  "Qtom.A.LA.204" 
## [20,] "Qtom.A.LA.207"  "Qtom.A.LA.207" 
## [21,] "Qtom.A.LA.208"  "Qtom.A.LA.208" 
## [22,] "Qtom.A.LA.209"  "Qtom.A.LA.209" 
## [23,] "Qtom.A.LA.218"  "Qtom.A.LA.218" 
## [24,] "Qtom.A.LA.221"  "Qtom.A.LA.221" 
## [25,] "Qtom.A.SB.324"  "Qtom.A.SB.324" 
## [26,] "Qtom.A.SB.325"  "Qtom.A.SB.325" 
## [27,] "Qtom.A.SB.327"  "Qtom.A.SB.327" 
## [28,] "Qtom.A.SB.329"  "Qtom.A.SB.329" 
## [29,] "Qtom.A.SB.330"  "Qtom.A.SB.330" 
## [30,] "Qtom.A.SB.331"  "Qtom.A.SB.331" 
## [31,] "Qtom.A.SB.332"  "Qtom.A.SB.332" 
## [32,] "Qtom.A.SB.333"  "Qtom.A.SB.333" 
## [33,] "Qtom.A.SB.335"  "Qtom.A.SB.335" 
## [34,] "Qtom.A.SB.336"  "Qtom.A.SB.336" 
## [35,] "Qtom.A.SB.337"  "Qtom.A.SB.337" 
## [36,] "Qtom.A.SB.338"  "Qtom.A.SB.338" 
## [37,] "Qtom.A.SB.339"  "Qtom.A.SB.339" 
## [38,] "Qtom.A.SB.341"  "Qtom.A.SB.341" 
## [39,] "Qtom.A.SB.342"  "Qtom.A.SB.342" 
## [40,] "Qtom.A.SB.343"  "Qtom.A.SB.343" 
## [41,] "Qtom.A.SB.344"  "Qtom.A.SB.344" 
## [42,] "Qtom.A.SB.345"  "Qtom.A.SB.345" 
## [43,] "Qtom.A.SB.346"  "Qtom.A.SB.346" 
## [44,] "Qtom.A.SB.347"  "Qtom.A.SB.347" 
## [45,] "Qtom.A.SB.348"  "Qtom.A.SB.348" 
## [46,] "Qtom.A.SB.352"  "Qtom.A.SB.352" 
## [47,] "Qtom.A.SB.353"  "Qtom.A.SB.353" 
## [48,] "Qtom.A.SB.355"  "Qtom.A.SB.355" 
## [49,] "Qtom.I.LA.05"   "Qtom.I.LA.05"  
## [50,] "Qtom.I.LA.06"   "Qtom.I.LA.06"  
## [51,] "Qtom.I.LA.08"   "Qtom.I.LA.08"  
## [52,] "Qtom.I.LA.10"   "Qtom.I.LA.10"  
## [53,] "Qtom.I.LA.11"   "Qtom.I.LA.11"  
## [54,] "Qtom.I.LA.12"   "Qtom.I.LA.12"  
## [55,] "Qtom.I.LA.13"   "Qtom.I.LA.13"  
## [56,] "Qtom.I.LA.21"   "Qtom.I.LA.21"  
## [57,] "Qtom.I.LA.22"   "Qtom.I.LA.22"  
## [58,] "Qtom.I.LA.23"   "Qtom.I.LA.23"  
## [59,] "Qtom.I.LA.24"   "Qtom.I.LA.24"  
## [60,] "Qtom.I.SB.01"   "Qtom.I.SB.01"  
## [61,] "Qtom.I.SB.04"   "Qtom.I.SB.04"  
## [62,] "Qtom.I.SB.09"   "Qtom.I.SB.09"  
## [63,] "Qtom.I.SB.10"   "Qtom.I.SB.10"  
## [64,] "Qtom.I.SB.11"   "Qtom.I.SB.11"  
## [65,] "Qtom.I.SB.12"   "Qtom.I.SB.12"  
## [66,] "Qtom.I.SB.28"   "Qtom.I.SB.28"  
## [67,] "Qtom.I.SB.33"   "Qtom.I.SB.33"  
## [68,] "Qtom.I.SB.35"   "Qtom.I.SB.35"  
## [69,] "Qtom.I.SB.36"   "Qtom.I.SB.36"  
## [70,] "Qtom.I.SB.37"   "Qtom.I.SB.37"  
## [71,] "Qtom.I.SB.38"   "Qtom.I.SB.38"  
## [72,] "Qtom.I.SB.40"   "Qtom.I.SB.40"  
## [73,] "Qtom.I.SB.41"   "Qtom.I.SB.41"  
## [74,] "Qtom.I.SB.42"   "Qtom.I.SB.42"  
## [75,] "Qtom.I.SB.44"   "Qtom.I.SB.44"  
## [76,] "Qtom.I.SB.45"   "Qtom.I.SB.45"  
## [77,] "Qtom.I.SB.73"   "Qtom.I.SB.73"  
## [78,] "Qtom.I.SB.74"   "Qtom.I.SB.74"  
## [79,] "Qtom.I.VEN.01"  "Qtom.I.VEN.01" 
## [80,] "Qtom.I.VEN.02"  "Qtom.I.VEN.02" 
## [81,] "Qtom.I.VEN.03"  "Qtom.I.VEN.03" 
## [82,] "Qtom.I.VEN.04"  "Qtom.I.VEN.04" 
## [83,] "Qtom.I.VEN.05"  "Qtom.I.VEN.05" 
## [84,] "Qtom.JJK.SB.01" "Qtom.JJK.SB.01"
## [85,] "Qtom.JJK.SB.03" "Qtom.JJK.SB.03"
## [86,] "Qtom.JJK.SB.07" "Qtom.JJK.SB.07"
## [87,] "Qtom.JJK.SB.08" "Qtom.JJK.SB.08"
## [88,] "Qtom.JJK.SB.09" "Qtom.JJK.SB.09"
## [89,] "Qtom.JJK.SB.10" "Qtom.JJK.SB.10"
## [90,] "Qtom.JR.SB.01"  "Qtom.JR.SB.01" 
## [91,] "Qtom.S.LA.186"  "Qtom.S.LA.186"
```

```
sum(rownames(clim) != rownames(snps))
```

```
## [1] 0
```

```
cbind(rownames(snps.cand), rownames(snps))
```

```
##       [,1]             [,2]            
##  [1,] "Qchr.A.LA.215"  "Qchr.A.LA.215" 
##  [2,] "Qchr.A.LA.222"  "Qchr.A.LA.222" 
##  [3,] "Qchr.I.LA.20"   "Qchr.I.LA.20"  
##  [4,] "Qchr.I.LA.25"   "Qchr.I.LA.25"  
##  [5,] "Qchr.I.SB.02"   "Qchr.I.SB.02"  
##  [6,] "Qchr.I.SB.03"   "Qchr.I.SB.03"  
##  [7,] "Qchr.JJK.SB.13" "Qchr.JJK.SB.13"
##  [8,] "Qchr.JR.SB.01"  "Qchr.JR.SB.01" 
##  [9,] "Qtom.A.LA.169"  "Qtom.A.LA.169" 
## [10,] "Qtom.A.LA.175"  "Qtom.A.LA.175" 
## [11,] "Qtom.A.LA.176"  "Qtom.A.LA.176" 
## [12,] "Qtom.A.LA.180"  "Qtom.A.LA.180" 
## [13,] "Qtom.A.LA.181"  "Qtom.A.LA.181" 
## [14,] "Qtom.A.LA.183"  "Qtom.A.LA.183" 
## [15,] "Qtom.A.LA.190"  "Qtom.A.LA.190" 
## [16,] "Qtom.A.LA.191"  "Qtom.A.LA.191" 
## [17,] "Qtom.A.LA.198"  "Qtom.A.LA.198" 
## [18,] "Qtom.A.LA.200"  "Qtom.A.LA.200" 
## [19,] "Qtom.A.LA.204"  "Qtom.A.LA.204" 
## [20,] "Qtom.A.LA.207"  "Qtom.A.LA.207" 
## [21,] "Qtom.A.LA.208"  "Qtom.A.LA.208" 
## [22,] "Qtom.A.LA.209"  "Qtom.A.LA.209" 
## [23,] "Qtom.A.LA.218"  "Qtom.A.LA.218" 
## [24,] "Qtom.A.LA.221"  "Qtom.A.LA.221" 
## [25,] "Qtom.A.SB.324"  "Qtom.A.SB.324" 
## [26,] "Qtom.A.SB.325"  "Qtom.A.SB.325" 
## [27,] "Qtom.A.SB.327"  "Qtom.A.SB.327" 
## [28,] "Qtom.A.SB.329"  "Qtom.A.SB.329" 
## [29,] "Qtom.A.SB.330"  "Qtom.A.SB.330" 
## [30,] "Qtom.A.SB.331"  "Qtom.A.SB.331" 
## [31,] "Qtom.A.SB.332"  "Qtom.A.SB.332" 
## [32,] "Qtom.A.SB.333"  "Qtom.A.SB.333" 
## [33,] "Qtom.A.SB.335"  "Qtom.A.SB.335" 
## [34,] "Qtom.A.SB.336"  "Qtom.A.SB.336" 
## [35,] "Qtom.A.SB.337"  "Qtom.A.SB.337" 
## [36,] "Qtom.A.SB.338"  "Qtom.A.SB.338" 
## [37,] "Qtom.A.SB.339"  "Qtom.A.SB.339" 
## [38,] "Qtom.A.SB.341"  "Qtom.A.SB.341" 
## [39,] "Qtom.A.SB.342"  "Qtom.A.SB.342" 
## [40,] "Qtom.A.SB.343"  "Qtom.A.SB.343" 
## [41,] "Qtom.A.SB.344"  "Qtom.A.SB.344" 
## [42,] "Qtom.A.SB.345"  "Qtom.A.SB.345" 
## [43,] "Qtom.A.SB.346"  "Qtom.A.SB.346" 
## [44,] "Qtom.A.SB.347"  "Qtom.A.SB.347" 
## [45,] "Qtom.A.SB.348"  "Qtom.A.SB.348" 
## [46,] "Qtom.A.SB.352"  "Qtom.A.SB.352" 
## [47,] "Qtom.A.SB.353"  "Qtom.A.SB.353" 
## [48,] "Qtom.A.SB.355"  "Qtom.A.SB.355" 
## [49,] "Qtom.I.LA.05"   "Qtom.I.LA.05"  
## [50,] "Qtom.I.LA.06"   "Qtom.I.LA.06"  
## [51,] "Qtom.I.LA.08"   "Qtom.I.LA.08"  
## [52,] "Qtom.I.LA.10"   "Qtom.I.LA.10"  
## [53,] "Qtom.I.LA.11"   "Qtom.I.LA.11"  
## [54,] "Qtom.I.LA.12"   "Qtom.I.LA.12"  
## [55,] "Qtom.I.LA.13"   "Qtom.I.LA.13"  
## [56,] "Qtom.I.LA.21"   "Qtom.I.LA.21"  
## [57,] "Qtom.I.LA.22"   "Qtom.I.LA.22"  
## [58,] "Qtom.I.LA.23"   "Qtom.I.LA.23"  
## [59,] "Qtom.I.LA.24"   "Qtom.I.LA.24"  
## [60,] "Qtom.I.SB.01"   "Qtom.I.SB.01"  
## [61,] "Qtom.I.SB.04"   "Qtom.I.SB.04"  
## [62,] "Qtom.I.SB.09"   "Qtom.I.SB.09"  
## [63,] "Qtom.I.SB.10"   "Qtom.I.SB.10"  
## [64,] "Qtom.I.SB.11"   "Qtom.I.SB.11"  
## [65,] "Qtom.I.SB.12"   "Qtom.I.SB.12"  
## [66,] "Qtom.I.SB.28"   "Qtom.I.SB.28"  
## [67,] "Qtom.I.SB.33"   "Qtom.I.SB.33"  
## [68,] "Qtom.I.SB.35"   "Qtom.I.SB.35"  
## [69,] "Qtom.I.SB.36"   "Qtom.I.SB.36"  
## [70,] "Qtom.I.SB.37"   "Qtom.I.SB.37"  
## [71,] "Qtom.I.SB.38"   "Qtom.I.SB.38"  
## [72,] "Qtom.I.SB.40"   "Qtom.I.SB.40"  
## [73,] "Qtom.I.SB.41"   "Qtom.I.SB.41"  
## [74,] "Qtom.I.SB.42"   "Qtom.I.SB.42"  
## [75,] "Qtom.I.SB.44"   "Qtom.I.SB.44"  
## [76,] "Qtom.I.SB.45"   "Qtom.I.SB.45"  
## [77,] "Qtom.I.SB.73"   "Qtom.I.SB.73"  
## [78,] "Qtom.I.SB.74"   "Qtom.I.SB.74"  
## [79,] "Qtom.I.VEN.01"  "Qtom.I.VEN.01" 
## [80,] "Qtom.I.VEN.02"  "Qtom.I.VEN.02" 
## [81,] "Qtom.I.VEN.03"  "Qtom.I.VEN.03" 
## [82,] "Qtom.I.VEN.04"  "Qtom.I.VEN.04" 
## [83,] "Qtom.I.VEN.05"  "Qtom.I.VEN.05" 
## [84,] "Qtom.JJK.SB.01" "Qtom.JJK.SB.01"
## [85,] "Qtom.JJK.SB.03" "Qtom.JJK.SB.03"
## [86,] "Qtom.JJK.SB.07" "Qtom.JJK.SB.07"
## [87,] "Qtom.JJK.SB.08" "Qtom.JJK.SB.08"
## [88,] "Qtom.JJK.SB.09" "Qtom.JJK.SB.09"
## [89,] "Qtom.JJK.SB.10" "Qtom.JJK.SB.10"
## [90,] "Qtom.JR.SB.01"  "Qtom.JR.SB.01" 
## [91,] "Qtom.S.LA.186"  "Qtom.S.LA.186"
```

```
sum(rownames(snps.cand) != rownames(snps))
```

```
## [1] 0
```

Here `clim` gets subset to variables of interest

```
# use bioclim vars, which have better coverage of San Clemente
bclim <- clim[,c("bio1", "bio2", "bio3", "bio4", "bio5", "bio6", "bio7", "bio8", "bio9", "bio10", "bio11", "bio12", "bio13", "bio14", "bio15", "bio16", "bio17", "bio18", "bio19", "elev")]

# using the same variables used in the RDA
vars <- c('bio5', 'bio6', 'bio15', 'bio18', 'bio19', 'elev')

heatmap(abs(cor(bclim[,vars])), scale = 'none')
```

```
pairs.panels(bclim[,vars], scale = T)
```

```
bclim <- bclim[,vars]
```

```
# Other papers' methods are a bit unclear about whether they used distances or the actual lat/lon values. But the function requires a distance matrix.

# load distance matrix, named 'dist_df', rename to 'dist'
# made in 'calculate_distance_matrix.Rmd'
load('data/clean/geo_distance_matrix.rda')
dist <- dist_df
rm(dist_df)
heatmap(as.matrix(dist), Rowv = NA, Colv = NA, scale = 'none')
```

```
# subset to include only the individuals being used here
dist <- dist[rownames(dist) %in% clim$ID_vcf, colnames(dist) %in% clim$ID_vcf]
dist <- dist[clim$ID_vcf, clim$ID_vcf]

heatmap(as.matrix(dist), Rowv = NA, Colv = NA, scale = 'none')
```

```
# check order
cbind(rownames(dist), clim$ID_vcf)
```

```
##       [,1]             [,2]            
##  [1,] "Qchr.A.LA.215"  "Qchr.A.LA.215" 
##  [2,] "Qchr.A.LA.222"  "Qchr.A.LA.222" 
##  [3,] "Qchr.I.LA.20"   "Qchr.I.LA.20"  
##  [4,] "Qchr.I.LA.25"   "Qchr.I.LA.25"  
##  [5,] "Qchr.I.SB.02"   "Qchr.I.SB.02"  
##  [6,] "Qchr.I.SB.03"   "Qchr.I.SB.03"  
##  [7,] "Qchr.JJK.SB.13" "Qchr.JJK.SB.13"
##  [8,] "Qchr.JR.SB.01"  "Qchr.JR.SB.01" 
##  [9,] "Qtom.A.LA.169"  "Qtom.A.LA.169" 
## [10,] "Qtom.A.LA.175"  "Qtom.A.LA.175" 
## [11,] "Qtom.A.LA.176"  "Qtom.A.LA.176" 
## [12,] "Qtom.A.LA.180"  "Qtom.A.LA.180" 
## [13,] "Qtom.A.LA.181"  "Qtom.A.LA.181" 
## [14,] "Qtom.A.LA.183"  "Qtom.A.LA.183" 
## [15,] "Qtom.A.LA.190"  "Qtom.A.LA.190" 
## [16,] "Qtom.A.LA.191"  "Qtom.A.LA.191" 
## [17,] "Qtom.A.LA.198"  "Qtom.A.LA.198" 
## [18,] "Qtom.A.LA.200"  "Qtom.A.LA.200" 
## [19,] "Qtom.A.LA.204"  "Qtom.A.LA.204" 
## [20,] "Qtom.A.LA.207"  "Qtom.A.LA.207" 
## [21,] "Qtom.A.LA.208"  "Qtom.A.LA.208" 
## [22,] "Qtom.A.LA.209"  "Qtom.A.LA.209" 
## [23,] "Qtom.A.LA.218"  "Qtom.A.LA.218" 
## [24,] "Qtom.A.LA.221"  "Qtom.A.LA.221" 
## [25,] "Qtom.A.SB.324"  "Qtom.A.SB.324" 
## [26,] "Qtom.A.SB.325"  "Qtom.A.SB.325" 
## [27,] "Qtom.A.SB.327"  "Qtom.A.SB.327" 
## [28,] "Qtom.A.SB.329"  "Qtom.A.SB.329" 
## [29,] "Qtom.A.SB.330"  "Qtom.A.SB.330" 
## [30,] "Qtom.A.SB.331"  "Qtom.A.SB.331" 
## [31,] "Qtom.A.SB.332"  "Qtom.A.SB.332" 
## [32,] "Qtom.A.SB.333"  "Qtom.A.SB.333" 
## [33,] "Qtom.A.SB.335"  "Qtom.A.SB.335" 
## [34,] "Qtom.A.SB.336"  "Qtom.A.SB.336" 
## [35,] "Qtom.A.SB.337"  "Qtom.A.SB.337" 
## [36,] "Qtom.A.SB.338"  "Qtom.A.SB.338" 
## [37,] "Qtom.A.SB.339"  "Qtom.A.SB.339" 
## [38,] "Qtom.A.SB.341"  "Qtom.A.SB.341" 
## [39,] "Qtom.A.SB.342"  "Qtom.A.SB.342" 
## [40,] "Qtom.A.SB.343"  "Qtom.A.SB.343" 
## [41,] "Qtom.A.SB.344"  "Qtom.A.SB.344" 
## [42,] "Qtom.A.SB.345"  "Qtom.A.SB.345" 
## [43,] "Qtom.A.SB.346"  "Qtom.A.SB.346" 
## [44,] "Qtom.A.SB.347"  "Qtom.A.SB.347" 
## [45,] "Qtom.A.SB.348"  "Qtom.A.SB.348" 
## [46,] "Qtom.A.SB.352"  "Qtom.A.SB.352" 
## [47,] "Qtom.A.SB.353"  "Qtom.A.SB.353" 
## [48,] "Qtom.A.SB.355"  "Qtom.A.SB.355" 
## [49,] "Qtom.I.LA.05"   "Qtom.I.LA.05"  
## [50,] "Qtom.I.LA.06"   "Qtom.I.LA.06"  
## [51,] "Qtom.I.LA.08"   "Qtom.I.LA.08"  
## [52,] "Qtom.I.LA.10"   "Qtom.I.LA.10"  
## [53,] "Qtom.I.LA.11"   "Qtom.I.LA.11"  
## [54,] "Qtom.I.LA.12"   "Qtom.I.LA.12"  
## [55,] "Qtom.I.LA.13"   "Qtom.I.LA.13"  
## [56,] "Qtom.I.LA.21"   "Qtom.I.LA.21"  
## [57,] "Qtom.I.LA.22"   "Qtom.I.LA.22"  
## [58,] "Qtom.I.LA.23"   "Qtom.I.LA.23"  
## [59,] "Qtom.I.LA.24"   "Qtom.I.LA.24"  
## [60,] "Qtom.I.SB.01"   "Qtom.I.SB.01"  
## [61,] "Qtom.I.SB.04"   "Qtom.I.SB.04"  
## [62,] "Qtom.I.SB.09"   "Qtom.I.SB.09"  
## [63,] "Qtom.I.SB.10"   "Qtom.I.SB.10"  
## [64,] "Qtom.I.SB.11"   "Qtom.I.SB.11"  
## [65,] "Qtom.I.SB.12"   "Qtom.I.SB.12"  
## [66,] "Qtom.I.SB.28"   "Qtom.I.SB.28"  
## [67,] "Qtom.I.SB.33"   "Qtom.I.SB.33"  
## [68,] "Qtom.I.SB.35"   "Qtom.I.SB.35"  
## [69,] "Qtom.I.SB.36"   "Qtom.I.SB.36"  
## [70,] "Qtom.I.SB.37"   "Qtom.I.SB.37"  
## [71,] "Qtom.I.SB.38"   "Qtom.I.SB.38"  
## [72,] "Qtom.I.SB.40"   "Qtom.I.SB.40"  
## [73,] "Qtom.I.SB.41"   "Qtom.I.SB.41"  
## [74,] "Qtom.I.SB.42"   "Qtom.I.SB.42"  
## [75,] "Qtom.I.SB.44"   "Qtom.I.SB.44"  
## [76,] "Qtom.I.SB.45"   "Qtom.I.SB.45"  
## [77,] "Qtom.I.SB.73"   "Qtom.I.SB.73"  
## [78,] "Qtom.I.SB.74"   "Qtom.I.SB.74"  
## [79,] "Qtom.I.VEN.01"  "Qtom.I.VEN.01" 
## [80,] "Qtom.I.VEN.02"  "Qtom.I.VEN.02" 
## [81,] "Qtom.I.VEN.03"  "Qtom.I.VEN.03" 
## [82,] "Qtom.I.VEN.04"  "Qtom.I.VEN.04" 
## [83,] "Qtom.I.VEN.05"  "Qtom.I.VEN.05" 
## [84,] "Qtom.JJK.SB.01" "Qtom.JJK.SB.01"
## [85,] "Qtom.JJK.SB.03" "Qtom.JJK.SB.03"
## [86,] "Qtom.JJK.SB.07" "Qtom.JJK.SB.07"
## [87,] "Qtom.JJK.SB.08" "Qtom.JJK.SB.08"
## [88,] "Qtom.JJK.SB.09" "Qtom.JJK.SB.09"
## [89,] "Qtom.JJK.SB.10" "Qtom.JJK.SB.10"
## [90,] "Qtom.JR.SB.01"  "Qtom.JR.SB.01" 
## [91,] "Qtom.S.LA.186"  "Qtom.S.LA.186"
```

```
sum(rownames(dist) != clim$ID_vcf)
```

```
## [1] 0
```

```
# rename to match
rownames(dist) <- rownames(clim)
sum(rownames(dist) != rownames(clim))
```

```
## [1] 0
```

```
heatmap(as.matrix(dist), Rowv = NA, Colv = NA, scale = 'none')

# convert to km
dist <- dist/1000
```

#### 1.3 Calculate MEM variables

```
# calculate MEM/PCMN variables (used as measures of distance in gradient forest)

# test pcnm with default params
mem <- pcnm(dist)
mem
```

```
## $vectors
##                      PCNM1        PCNM2       PCNM3       PCNM4         PCNM5
## Qchr.A.LA.215  -0.14251161 -0.018059393  0.14239216 -0.05173643 -4.214916e-04
## Qchr.A.LA.222  -0.14251133 -0.018056812  0.14246436 -0.05176154 -4.240382e-04
## Qchr.I.LA.20   -0.14230681 -0.028307399 -0.18312638  0.06191715 -2.513762e-03
## Qchr.I.LA.25   -0.14213194 -0.029402015 -0.21761228  0.07393269  1.567352e-03
## Qchr.I.SB.02    0.07742797 -0.009114040 -0.01742144 -0.05183681 -1.489870e-01
## Qchr.I.SB.03    0.07742346 -0.008893868 -0.01876871 -0.05568224 -1.438874e-01
## Qchr.JJK.SB.13  0.07740500 -0.008162865 -0.02328759 -0.06858405 -1.450610e-01
## Qchr.JR.SB.01   0.07741210 -0.008542671 -0.02108259 -0.06230439 -2.026678e-01
## Qtom.A.LA.169  -0.14239604 -0.017150926  0.17158158 -0.06195092  3.626812e-03
## Qtom.A.LA.175  -0.14252288 -0.018359372  0.13433471 -0.04894149  8.098752e-04
## Qtom.A.LA.176  -0.14252344 -0.018363120  0.13421116 -0.04889820  7.873179e-04
## Qtom.A.LA.180  -0.14251255 -0.018067749  0.14206247 -0.05162026 -5.279459e-04
## Qtom.A.LA.181  -0.14251245 -0.018066818  0.14209315 -0.05163101 -5.233273e-04
## Qtom.A.LA.183  -0.14251076 -0.018051411  0.14258199 -0.05180195 -4.689880e-04
## Qtom.A.LA.190  -0.14251045 -0.018048586  0.14267651 -0.05183507 -4.532342e-04
## Qtom.A.LA.191  -0.14251068 -0.018050691  0.14260668 -0.05181061 -4.642580e-04
## Qtom.A.LA.198  -0.14251044 -0.018048502  0.14269038 -0.05184007 -4.396806e-04
## Qtom.A.LA.200  -0.14251024 -0.018046726  0.14271785 -0.05184929 -4.674702e-04
## Qtom.A.LA.204  -0.14250740 -0.018021223  0.14348988 -0.05211877 -4.205222e-04
## Qtom.A.LA.207  -0.14083853  0.691960245  0.01369803  0.06042168 -5.611915e-04
## Qtom.A.LA.208  -0.14250989 -0.018043930  0.14275663 -0.05186218 -5.155655e-04
## Qtom.A.LA.209  -0.14250828 -0.018028957  0.14328497 -0.05204766 -4.006351e-04
## Qtom.A.LA.218  -0.14251170 -0.018060184  0.14235864 -0.05172459 -4.344511e-04
## Qtom.A.LA.221  -0.14251252 -0.018067659  0.14203272 -0.05160942 -5.656843e-04
## Qtom.A.SB.324   0.07713776 -0.020763127  0.05502893  0.15534065  1.685544e-01
## Qtom.A.SB.325   0.07713921 -0.020741844  0.05489989  0.15497163  1.691599e-01
## Qtom.A.SB.327   0.07713688 -0.020781542  0.05512716  0.15561988  1.624415e-01
## Qtom.A.SB.329   0.07717013 -0.020247595  0.05195581  0.14655897  2.052858e-01
## Qtom.A.SB.330   0.07717013 -0.020247567  0.05195591  0.14655928  2.053969e-01
## Qtom.A.SB.331   0.07718808 -0.019982189  0.05030340  0.14182827  1.950316e-01
## Qtom.A.SB.332   0.07718734 -0.019988644  0.05035434  0.14197546  1.997634e-01
## Qtom.A.SB.333   0.07719028 -0.019977615  0.05020696  0.14154373  1.665369e-01
## Qtom.A.SB.335   0.07726695 -0.018800789  0.04255423  0.11959988 -1.226211e-02
## Qtom.A.SB.336   0.07726294 -0.018876485  0.04300538  0.12088872 -1.783715e-02
## Qtom.A.SB.337   0.07726325 -0.018869203  0.04297188  0.12079422 -1.321619e-02
## Qtom.A.SB.338   0.07725993 -0.018932898  0.04333880  0.12184090 -2.314171e-02
## Qtom.A.SB.339   0.07726054 -0.018922140  0.04327109  0.12164705 -2.383059e-02
## Qtom.A.SB.341   0.07725692 -0.018985349  0.04367691  0.12280993 -1.647268e-02
## Qtom.A.SB.342   0.07725590 -0.019004675  0.04378750  0.12312532 -1.978814e-02
## Qtom.A.SB.343   0.07725341 -0.019051420  0.04405783  0.12389664 -2.662710e-02
## Qtom.A.SB.344   0.07725257 -0.019058449  0.04415794  0.12418967 -3.122255e-03
## Qtom.A.SB.345   0.07725057 -0.019095056  0.04437646  0.12481401 -5.664521e-03
## Qtom.A.SB.346   0.07724909 -0.019120653  0.04453688  0.12527326 -4.296082e-03
## Qtom.A.SB.347   0.07724586 -0.019178482  0.04488570  0.12627034 -6.812654e-03
## Qtom.A.SB.348   0.07724451 -0.019201253  0.04503050  0.12668513 -4.726529e-03
## Qtom.A.SB.352   0.07727324 -0.018690083  0.04182139  0.11749753 -3.421319e-02
## Qtom.A.SB.353   0.07727557 -0.018651837  0.04151057  0.11659975 -6.551879e-02
## Qtom.A.SB.355   0.07727549 -0.018654072  0.04150774  0.11658974 -7.233028e-02
## Qtom.I.LA.05   -0.14218339 -0.029113777 -0.20793038  0.07055120  1.063610e-03
## Qtom.I.LA.06   -0.14217667 -0.029150274 -0.20899026  0.07091903  1.329181e-03
## Qtom.I.LA.08   -0.14219478 -0.029048378 -0.20596182  0.06986680  6.867055e-04
## Qtom.I.LA.10   -0.14231707 -0.028223684 -0.18058342  0.06103210 -2.856616e-03
## Qtom.I.LA.11   -0.14227894 -0.028508973 -0.18935364  0.06408562 -1.728475e-03
## Qtom.I.LA.12   -0.14224084 -0.028765908 -0.19725753  0.06683707 -6.213162e-04
## Qtom.I.LA.13   -0.14219696 -0.029035359 -0.20561454  0.06974675  5.609874e-04
## Qtom.I.LA.21   -0.14229803 -0.028365923 -0.18499169  0.06256746 -2.335691e-03
## Qtom.I.LA.22   -0.14222265 -0.028881313 -0.20078088  0.06806304 -6.429852e-05
## Qtom.I.LA.23   -0.14223377 -0.028812737 -0.19864185  0.06731810 -3.497008e-04
## Qtom.I.LA.24   -0.14214700 -0.029314289 -0.21409989  0.07269799  2.098736e-03
## Qtom.I.SB.01    0.07743950 -0.009529523 -0.01454605 -0.04359127 -2.235013e-02
## Qtom.I.SB.04    0.07742362 -0.008910369 -0.01867814 -0.05542488 -1.485094e-01
## Qtom.I.SB.09    0.07713858 -0.002612343 -0.05697386 -0.16462804  1.288372e-01
## Qtom.I.SB.10    0.07713507 -0.002562511 -0.05728358 -0.16551147  1.285272e-01
## Qtom.I.SB.11    0.07714194 -0.002660137 -0.05669383 -0.16383136  1.221415e-01
## Qtom.I.SB.12    0.07713681 -0.002587180 -0.05714328 -0.16511286  1.233295e-01
## Qtom.I.SB.28    0.07743131 -0.009104629 -0.01720408 -0.05118511 -3.646958e-02
## Qtom.I.SB.33    0.07743679 -0.009382703 -0.01547126 -0.04623539 -3.002326e-02
## Qtom.I.SB.35    0.07743696 -0.009391663 -0.01541446 -0.04607302 -2.941780e-02
## Qtom.I.SB.36    0.07743795 -0.009443730 -0.01508420 -0.04512897 -2.581970e-02
## Qtom.I.SB.37    0.07743983 -0.009549079 -0.01442554 -0.04324716 -2.244059e-02
## Qtom.I.SB.38    0.07744065 -0.009597064 -0.01412626 -0.04239222 -2.119801e-02
## Qtom.I.SB.40    0.07744446 -0.009833325 -0.01264160 -0.03814949 -1.045189e-02
## Qtom.I.SB.41    0.07744458 -0.009842937 -0.01258644 -0.03799245 -1.216419e-02
## Qtom.I.SB.42    0.07744457 -0.009842624 -0.01259089 -0.03800545 -1.319611e-02
## Qtom.I.SB.44    0.07744498 -0.009885099 -0.01236738 -0.03737165 -2.905003e-02
## Qtom.I.SB.45    0.07744505 -0.009891873 -0.01233085 -0.03726794 -3.121505e-02
## Qtom.I.SB.73    0.07732607 -0.005908152 -0.03700471 -0.10771111 -5.338843e-02
## Qtom.I.SB.74    0.07732604 -0.005907934 -0.03700786 -0.10772030 -5.412615e-02
## Qtom.I.VEN.01   0.07662951  0.002921444 -0.09107851 -0.26181511  2.375939e-01
## Qtom.I.VEN.02   0.07662894  0.002926313 -0.09110460 -0.26188889  2.393276e-01
## Qtom.I.VEN.03   0.07662552  0.002956233 -0.09128599 -0.26240511  2.412865e-01
## Qtom.I.VEN.04   0.07663020  0.002915077 -0.09103310 -0.26168498  2.399836e-01
## Qtom.I.VEN.05   0.07342745  0.702381623 -0.02832824  0.02379929 -3.485856e-07
## Qtom.JJK.SB.01  0.07741402 -0.008599908 -0.02070780 -0.06123207 -1.940574e-01
## Qtom.JJK.SB.03  0.07741318 -0.008567355 -0.02091032 -0.06181042 -1.946329e-01
## Qtom.JJK.SB.07  0.07741112 -0.008506335 -0.02130938 -0.06295214 -2.036085e-01
## Qtom.JJK.SB.08  0.07740963 -0.008474984 -0.02153091 -0.06358760 -2.149142e-01
## Qtom.JJK.SB.09  0.07740917 -0.008419498 -0.02183319 -0.06444623 -1.984567e-01
## Qtom.JJK.SB.10  0.07740919 -0.008394964 -0.02195612 -0.06479407 -1.868113e-01
## Qtom.JR.SB.01   0.07741221 -0.008552680 -0.02102642 -0.06214463 -2.049679e-01
## Qtom.S.LA.186  -0.14250992 -0.018043839  0.14282664 -0.05188757 -4.370300e-04
##                        PCNM6
## Qchr.A.LA.215   5.272285e-02
## Qchr.A.LA.222   5.619191e-02
## Qchr.I.LA.20    2.361035e-01
## Qchr.I.LA.25    1.123721e-02
## Qchr.I.SB.02   -7.704581e-04
## Qchr.I.SB.03   -7.306966e-04
## Qchr.JJK.SB.13 -7.859479e-04
## Qchr.JR.SB.01  -1.328475e-03
## Qtom.A.LA.169  -2.102612e-01
## Qtom.A.LA.175  -4.994774e-01
## Qtom.A.LA.176  -4.971017e-01
## Qtom.A.LA.180   7.885911e-02
## Qtom.A.LA.181   7.830612e-02
## Qtom.A.LA.183   7.649342e-02
## Qtom.A.LA.190   7.425711e-02
## Qtom.A.LA.191   7.568690e-02
## Qtom.A.LA.198   6.988973e-02
## Qtom.A.LA.200   8.091574e-02
## Qtom.A.LA.204   9.234517e-02
## Qtom.A.LA.207  -8.597461e-07
## Qtom.A.LA.208   9.999014e-02
## Qtom.A.LA.209   7.751503e-02
## Qtom.A.LA.218   5.610050e-02
## Qtom.A.LA.221   9.152552e-02
## Qtom.A.SB.324   9.211613e-04
## Qtom.A.SB.325   9.327591e-04
## Qtom.A.SB.327   8.584389e-04
## Qtom.A.SB.329   1.406238e-03
## Qtom.A.SB.330   1.407290e-03
## Qtom.A.SB.331   1.378804e-03
## Qtom.A.SB.332   1.421669e-03
## Qtom.A.SB.333   1.111486e-03
## Qtom.A.SB.335  -3.085166e-04
## Qtom.A.SB.336  -3.788808e-04
## Qtom.A.SB.337  -3.329102e-04
## Qtom.A.SB.338  -4.424689e-04
## Qtom.A.SB.339  -4.466611e-04
## Qtom.A.SB.341  -3.903145e-04
## Qtom.A.SB.342  -4.265185e-04
## Qtom.A.SB.343  -5.028240e-04
## Qtom.A.SB.344  -2.789508e-04
## Qtom.A.SB.345  -3.117142e-04
## Qtom.A.SB.346  -3.044922e-04
## Qtom.A.SB.347  -3.419912e-04
## Qtom.A.SB.348  -3.272910e-04
## Qtom.A.SB.352  -4.949456e-04
## Qtom.A.SB.353  -7.879627e-04
## Qtom.A.SB.355  -8.541228e-04
## Qtom.I.LA.05   -1.587731e-01
## Qtom.I.LA.06   -2.020655e-01
## Qtom.I.LA.08   -1.087617e-01
## Qtom.I.LA.10    2.873398e-01
## Qtom.I.LA.11    1.504600e-01
## Qtom.I.LA.12    2.597139e-02
## Qtom.I.LA.13   -8.151271e-02
## Qtom.I.LA.21    2.329465e-01
## Qtom.I.LA.22   -4.130724e-02
## Qtom.I.LA.23   -1.872657e-02
## Qtom.I.LA.24   -2.612462e-01
## Qtom.I.SB.01    4.815127e-04
## Qtom.I.SB.04   -7.752547e-04
## Qtom.I.SB.09    1.077977e-03
## Qtom.I.SB.10    1.063970e-03
## Qtom.I.SB.11    1.023785e-03
## Qtom.I.SB.12    1.019211e-03
## Qtom.I.SB.28    3.286247e-04
## Qtom.I.SB.33    4.021265e-04
## Qtom.I.SB.35    4.083234e-04
## Qtom.I.SB.36    4.450563e-04
## Qtom.I.SB.37    4.812351e-04
## Qtom.I.SB.38    4.947284e-04
## Qtom.I.SB.40    6.050918e-04
## Qtom.I.SB.41    5.887023e-04
## Qtom.I.SB.42    5.786873e-04
## Qtom.I.SB.44    4.257495e-04
## Qtom.I.SB.45    4.048662e-04
## Qtom.I.SB.73   -1.139755e-04
## Qtom.I.SB.74   -1.212148e-04
## Qtom.I.VEN.01   4.841492e-04
## Qtom.I.VEN.02   4.990129e-04
## Qtom.I.VEN.03   5.066688e-04
## Qtom.I.VEN.04   5.095219e-04
## Qtom.I.VEN.05  -1.161357e-04
## Qtom.JJK.SB.01 -1.240155e-03
## Qtom.JJK.SB.03 -1.247745e-03
## Qtom.JJK.SB.07 -1.339958e-03
## Qtom.JJK.SB.08 -1.453513e-03
## Qtom.JJK.SB.09 -1.294530e-03
## Qtom.JJK.SB.10 -1.181260e-03
## Qtom.JR.SB.01  -1.350573e-03
## Qtom.S.LA.186   7.389018e-02
## 
## $values
## [1]  4.471213e+06  1.005011e+05  2.214546e+04  2.147256e+04  2.301026e+02
## [6]  1.263256e+02 -3.601749e+00 -9.831838e+04
## 
## $weights
##  [1] 1 1 1 1 1 1 1 1 1 1 1 1 1 1 1 1 1 1 1 1 1 1 1 1 1 1 1 1 1 1 1 1 1 1 1 1 1 1
## [39] 1 1 1 1 1 1 1 1 1 1 1 1 1 1 1 1 1 1 1 1 1 1 1 1 1 1 1 1 1 1 1 1 1 1 1 1 1 1
## [77] 1 1 1 1 1 1 1 1 1 1 1 1 1 1 1
## 
## $threshold
## [1] 116.3832
## 
## attr(,"class")
## [1] "pcnm"
```

```
# visualize

palette(c('black', "#3c4a8b", "#009c85", "#84bc5f", "#edb829", "#f57404","#b30000"))

ordisplom(mem, col = clim$island)
```

```
par(mfrow = c(2,4))
for(n in 1:ncol(mem$vectors)){
  
  plot(mem$vectors[,n], col = clim$island)
  
}

# Testing different threshold values
# see ?pcnm
# "The selection of truncation distance has a huge influence on the PCNM vectors. The default is to use the longest distance to keep data connected. The distances above truncation threshold are given an arbitrary value of 4 times threshold. For regular data, the first PCNM vectors show a wide scale variation and later PCNM vectors show smaller scale variation (Borcard & Legendre 2002), but for irregular data the interpretation is not as clear."
# this is pretty irregular/clustered data

# look at mem 2
# which separates out 2 individuals: Qtom.I.VEN.05 and Qtom.A.LA.207
# these two are the closest of the northern/southern island pairs, so this distance is used as the truncation distance, and they cluster together along MEM2. This doesn't really make sense biologically, so let's test other threshold values
mem$vectors[order(mem$vectors[,2]), ]
```

```
##                      PCNM1        PCNM2       PCNM3       PCNM4         PCNM5
## Qchr.I.LA.25   -0.14213194 -0.029402015 -0.21761228  0.07393269  1.567352e-03
## Qtom.I.LA.24   -0.14214700 -0.029314289 -0.21409989  0.07269799  2.098736e-03
## Qtom.I.LA.06   -0.14217667 -0.029150274 -0.20899026  0.07091903  1.329181e-03
## Qtom.I.LA.05   -0.14218339 -0.029113777 -0.20793038  0.07055120  1.063610e-03
## Qtom.I.LA.08   -0.14219478 -0.029048378 -0.20596182  0.06986680  6.867055e-04
## Qtom.I.LA.13   -0.14219696 -0.029035359 -0.20561454  0.06974675  5.609874e-04
## Qtom.I.LA.22   -0.14222265 -0.028881313 -0.20078088  0.06806304 -6.429852e-05
## Qtom.I.LA.23   -0.14223377 -0.028812737 -0.19864185  0.06731810 -3.497008e-04
## Qtom.I.LA.12   -0.14224084 -0.028765908 -0.19725753  0.06683707 -6.213162e-04
## Qtom.I.LA.11   -0.14227894 -0.028508973 -0.18935364  0.06408562 -1.728475e-03
## Qtom.I.LA.21   -0.14229803 -0.028365923 -0.18499169  0.06256746 -2.335691e-03
## Qchr.I.LA.20   -0.14230681 -0.028307399 -0.18312638  0.06191715 -2.513762e-03
## Qtom.I.LA.10   -0.14231707 -0.028223684 -0.18058342  0.06103210 -2.856616e-03
## Qtom.A.SB.327   0.07713688 -0.020781542  0.05512716  0.15561988  1.624415e-01
## Qtom.A.SB.324   0.07713776 -0.020763127  0.05502893  0.15534065  1.685544e-01
## Qtom.A.SB.325   0.07713921 -0.020741844  0.05489989  0.15497163  1.691599e-01
## Qtom.A.SB.329   0.07717013 -0.020247595  0.05195581  0.14655897  2.052858e-01
## Qtom.A.SB.330   0.07717013 -0.020247567  0.05195591  0.14655928  2.053969e-01
## Qtom.A.SB.332   0.07718734 -0.019988644  0.05035434  0.14197546  1.997634e-01
## Qtom.A.SB.331   0.07718808 -0.019982189  0.05030340  0.14182827  1.950316e-01
## Qtom.A.SB.333   0.07719028 -0.019977615  0.05020696  0.14154373  1.665369e-01
## Qtom.A.SB.348   0.07724451 -0.019201253  0.04503050  0.12668513 -4.726529e-03
## Qtom.A.SB.347   0.07724586 -0.019178482  0.04488570  0.12627034 -6.812654e-03
## Qtom.A.SB.346   0.07724909 -0.019120653  0.04453688  0.12527326 -4.296082e-03
## Qtom.A.SB.345   0.07725057 -0.019095056  0.04437646  0.12481401 -5.664521e-03
## Qtom.A.SB.344   0.07725257 -0.019058449  0.04415794  0.12418967 -3.122255e-03
## Qtom.A.SB.343   0.07725341 -0.019051420  0.04405783  0.12389664 -2.662710e-02
## Qtom.A.SB.342   0.07725590 -0.019004675  0.04378750  0.12312532 -1.978814e-02
## Qtom.A.SB.341   0.07725692 -0.018985349  0.04367691  0.12280993 -1.647268e-02
## Qtom.A.SB.338   0.07725993 -0.018932898  0.04333880  0.12184090 -2.314171e-02
## Qtom.A.SB.339   0.07726054 -0.018922140  0.04327109  0.12164705 -2.383059e-02
## Qtom.A.SB.336   0.07726294 -0.018876485  0.04300538  0.12088872 -1.783715e-02
## Qtom.A.SB.337   0.07726325 -0.018869203  0.04297188  0.12079422 -1.321619e-02
## Qtom.A.SB.335   0.07726695 -0.018800789  0.04255423  0.11959988 -1.226211e-02
## Qtom.A.SB.352   0.07727324 -0.018690083  0.04182139  0.11749753 -3.421319e-02
## Qtom.A.SB.355   0.07727549 -0.018654072  0.04150774  0.11658974 -7.233028e-02
## Qtom.A.SB.353   0.07727557 -0.018651837  0.04151057  0.11659975 -6.551879e-02
## Qtom.A.LA.176  -0.14252344 -0.018363120  0.13421116 -0.04889820  7.873179e-04
## Qtom.A.LA.175  -0.14252288 -0.018359372  0.13433471 -0.04894149  8.098752e-04
## Qtom.A.LA.180  -0.14251255 -0.018067749  0.14206247 -0.05162026 -5.279459e-04
## Qtom.A.LA.221  -0.14251252 -0.018067659  0.14203272 -0.05160942 -5.656843e-04
## Qtom.A.LA.181  -0.14251245 -0.018066818  0.14209315 -0.05163101 -5.233273e-04
## Qtom.A.LA.218  -0.14251170 -0.018060184  0.14235864 -0.05172459 -4.344511e-04
## Qchr.A.LA.215  -0.14251161 -0.018059393  0.14239216 -0.05173643 -4.214916e-04
## Qchr.A.LA.222  -0.14251133 -0.018056812  0.14246436 -0.05176154 -4.240382e-04
## Qtom.A.LA.183  -0.14251076 -0.018051411  0.14258199 -0.05180195 -4.689880e-04
## Qtom.A.LA.191  -0.14251068 -0.018050691  0.14260668 -0.05181061 -4.642580e-04
## Qtom.A.LA.190  -0.14251045 -0.018048586  0.14267651 -0.05183507 -4.532342e-04
## Qtom.A.LA.198  -0.14251044 -0.018048502  0.14269038 -0.05184007 -4.396806e-04
## Qtom.A.LA.200  -0.14251024 -0.018046726  0.14271785 -0.05184929 -4.674702e-04
## Qtom.A.LA.208  -0.14250989 -0.018043930  0.14275663 -0.05186218 -5.155655e-04
## Qtom.S.LA.186  -0.14250992 -0.018043839  0.14282664 -0.05188757 -4.370300e-04
## Qtom.A.LA.209  -0.14250828 -0.018028957  0.14328497 -0.05204766 -4.006351e-04
## Qtom.A.LA.204  -0.14250740 -0.018021223  0.14348988 -0.05211877 -4.205222e-04
## Qtom.A.LA.169  -0.14239604 -0.017150926  0.17158158 -0.06195092  3.626812e-03
## Qtom.I.SB.45    0.07744505 -0.009891873 -0.01233085 -0.03726794 -3.121505e-02
## Qtom.I.SB.44    0.07744498 -0.009885099 -0.01236738 -0.03737165 -2.905003e-02
## Qtom.I.SB.41    0.07744458 -0.009842937 -0.01258644 -0.03799245 -1.216419e-02
## Qtom.I.SB.42    0.07744457 -0.009842624 -0.01259089 -0.03800545 -1.319611e-02
## Qtom.I.SB.40    0.07744446 -0.009833325 -0.01264160 -0.03814949 -1.045189e-02
## Qtom.I.SB.38    0.07744065 -0.009597064 -0.01412626 -0.04239222 -2.119801e-02
## Qtom.I.SB.37    0.07743983 -0.009549079 -0.01442554 -0.04324716 -2.244059e-02
## Qtom.I.SB.01    0.07743950 -0.009529523 -0.01454605 -0.04359127 -2.235013e-02
## Qtom.I.SB.36    0.07743795 -0.009443730 -0.01508420 -0.04512897 -2.581970e-02
## Qtom.I.SB.35    0.07743696 -0.009391663 -0.01541446 -0.04607302 -2.941780e-02
## Qtom.I.SB.33    0.07743679 -0.009382703 -0.01547126 -0.04623539 -3.002326e-02
## Qchr.I.SB.02    0.07742797 -0.009114040 -0.01742144 -0.05183681 -1.489870e-01
## Qtom.I.SB.28    0.07743131 -0.009104629 -0.01720408 -0.05118511 -3.646958e-02
## Qtom.I.SB.04    0.07742362 -0.008910369 -0.01867814 -0.05542488 -1.485094e-01
## Qchr.I.SB.03    0.07742346 -0.008893868 -0.01876871 -0.05568224 -1.438874e-01
## Qtom.JJK.SB.01  0.07741402 -0.008599908 -0.02070780 -0.06123207 -1.940574e-01
## Qtom.JJK.SB.03  0.07741318 -0.008567355 -0.02091032 -0.06181042 -1.946329e-01
## Qtom.JR.SB.01   0.07741221 -0.008552680 -0.02102642 -0.06214463 -2.049679e-01
## Qchr.JR.SB.01   0.07741210 -0.008542671 -0.02108259 -0.06230439 -2.026678e-01
## Qtom.JJK.SB.07  0.07741112 -0.008506335 -0.02130938 -0.06295214 -2.036085e-01
## Qtom.JJK.SB.08  0.07740963 -0.008474984 -0.02153091 -0.06358760 -2.149142e-01
## Qtom.JJK.SB.09  0.07740917 -0.008419498 -0.02183319 -0.06444623 -1.984567e-01
## Qtom.JJK.SB.10  0.07740919 -0.008394964 -0.02195612 -0.06479407 -1.868113e-01
## Qchr.JJK.SB.13  0.07740500 -0.008162865 -0.02328759 -0.06858405 -1.450610e-01
## Qtom.I.SB.73    0.07732607 -0.005908152 -0.03700471 -0.10771111 -5.338843e-02
## Qtom.I.SB.74    0.07732604 -0.005907934 -0.03700786 -0.10772030 -5.412615e-02
## Qtom.I.SB.11    0.07714194 -0.002660137 -0.05669383 -0.16383136  1.221415e-01
## Qtom.I.SB.09    0.07713858 -0.002612343 -0.05697386 -0.16462804  1.288372e-01
## Qtom.I.SB.12    0.07713681 -0.002587180 -0.05714328 -0.16511286  1.233295e-01
## Qtom.I.SB.10    0.07713507 -0.002562511 -0.05728358 -0.16551147  1.285272e-01
## Qtom.I.VEN.04   0.07663020  0.002915077 -0.09103310 -0.26168498  2.399836e-01
## Qtom.I.VEN.01   0.07662951  0.002921444 -0.09107851 -0.26181511  2.375939e-01
## Qtom.I.VEN.02   0.07662894  0.002926313 -0.09110460 -0.26188889  2.393276e-01
## Qtom.I.VEN.03   0.07662552  0.002956233 -0.09128599 -0.26240511  2.412865e-01
## Qtom.A.LA.207  -0.14083853  0.691960245  0.01369803  0.06042168 -5.611915e-04
## Qtom.I.VEN.05   0.07342745  0.702381623 -0.02832824  0.02379929 -3.485856e-07
##                        PCNM6
## Qchr.I.LA.25    1.123721e-02
## Qtom.I.LA.24   -2.612462e-01
## Qtom.I.LA.06   -2.020655e-01
## Qtom.I.LA.05   -1.587731e-01
## Qtom.I.LA.08   -1.087617e-01
## Qtom.I.LA.13   -8.151271e-02
## Qtom.I.LA.22   -4.130724e-02
## Qtom.I.LA.23   -1.872657e-02
## Qtom.I.LA.12    2.597139e-02
## Qtom.I.LA.11    1.504600e-01
## Qtom.I.LA.21    2.329465e-01
## Qchr.I.LA.20    2.361035e-01
## Qtom.I.LA.10    2.873398e-01
## Qtom.A.SB.327   8.584389e-04
## Qtom.A.SB.324   9.211613e-04
## Qtom.A.SB.325   9.327591e-04
## Qtom.A.SB.329   1.406238e-03
## Qtom.A.SB.330   1.407290e-03
## Qtom.A.SB.332   1.421669e-03
## Qtom.A.SB.331   1.378804e-03
## Qtom.A.SB.333   1.111486e-03
## Qtom.A.SB.348  -3.272910e-04
## Qtom.A.SB.347  -3.419912e-04
## Qtom.A.SB.346  -3.044922e-04
## Qtom.A.SB.345  -3.117142e-04
## Qtom.A.SB.344  -2.789508e-04
## Qtom.A.SB.343  -5.028240e-04
## Qtom.A.SB.342  -4.265185e-04
## Qtom.A.SB.341  -3.903145e-04
## Qtom.A.SB.338  -4.424689e-04
## Qtom.A.SB.339  -4.466611e-04
## Qtom.A.SB.336  -3.788808e-04
## Qtom.A.SB.337  -3.329102e-04
## Qtom.A.SB.335  -3.085166e-04
## Qtom.A.SB.352  -4.949456e-04
## Qtom.A.SB.355  -8.541228e-04
## Qtom.A.SB.353  -7.879627e-04
## Qtom.A.LA.176  -4.971017e-01
## Qtom.A.LA.175  -4.994774e-01
## Qtom.A.LA.180   7.885911e-02
## Qtom.A.LA.221   9.152552e-02
## Qtom.A.LA.181   7.830612e-02
## Qtom.A.LA.218   5.610050e-02
## Qchr.A.LA.215   5.272285e-02
## Qchr.A.LA.222   5.619191e-02
## Qtom.A.LA.183   7.649342e-02
## Qtom.A.LA.191   7.568690e-02
## Qtom.A.LA.190   7.425711e-02
## Qtom.A.LA.198   6.988973e-02
## Qtom.A.LA.200   8.091574e-02
## Qtom.A.LA.208   9.999014e-02
## Qtom.S.LA.186   7.389018e-02
## Qtom.A.LA.209   7.751503e-02
## Qtom.A.LA.204   9.234517e-02
## Qtom.A.LA.169  -2.102612e-01
## Qtom.I.SB.45    4.048662e-04
## Qtom.I.SB.44    4.257495e-04
## Qtom.I.SB.41    5.887023e-04
## Qtom.I.SB.42    5.786873e-04
## Qtom.I.SB.40    6.050918e-04
## Qtom.I.SB.38    4.947284e-04
## Qtom.I.SB.37    4.812351e-04
## Qtom.I.SB.01    4.815127e-04
## Qtom.I.SB.36    4.450563e-04
## Qtom.I.SB.35    4.083234e-04
## Qtom.I.SB.33    4.021265e-04
## Qchr.I.SB.02   -7.704581e-04
## Qtom.I.SB.28    3.286247e-04
## Qtom.I.SB.04   -7.752547e-04
## Qchr.I.SB.03   -7.306966e-04
## Qtom.JJK.SB.01 -1.240155e-03
## Qtom.JJK.SB.03 -1.247745e-03
## Qtom.JR.SB.01  -1.350573e-03
## Qchr.JR.SB.01  -1.328475e-03
## Qtom.JJK.SB.07 -1.339958e-03
## Qtom.JJK.SB.08 -1.453513e-03
## Qtom.JJK.SB.09 -1.294530e-03
## Qtom.JJK.SB.10 -1.181260e-03
## Qchr.JJK.SB.13 -7.859479e-04
## Qtom.I.SB.73   -1.139755e-04
## Qtom.I.SB.74   -1.212148e-04
## Qtom.I.SB.11    1.023785e-03
## Qtom.I.SB.09    1.077977e-03
## Qtom.I.SB.12    1.019211e-03
## Qtom.I.SB.10    1.063970e-03
## Qtom.I.VEN.04   5.095219e-04
## Qtom.I.VEN.01   4.841492e-04
## Qtom.I.VEN.02   4.990129e-04
## Qtom.I.VEN.03   5.066688e-04
## Qtom.A.LA.207  -8.597461e-07
## Qtom.I.VEN.05  -1.161357e-04
```

```
dist['Qtom.I.VEN.05', 'Qtom.A.LA.207']
```

```
## 116.3832 [m]
```

```
# default threshold value
mem$threshold # 116.3832 km
```

```
## [1] 116.3832
```

```
max(dist) # 200.831
```

```
## [1] 198.9725
```

```
# distance between samples is pretty bimodal since they are clustered within islands, then within northern/southern islands
par(mfrow = c(1,1))
```

```
hist(as.matrix(dist), breaks = 'fd')
abline(v = mem$threshold) # threshold near low end of the northern/southern break
```

```
# try increasing the threshold values (by km)
thresh <- c(120, 150, 170, 200, 200, 205, 250, 300) # thresholds to test

for(n in 1:length(thresh)){
  
  mtest <- pcnm(dist, threshold = thresh[n])
  
  # plot
  par(mfrow = c(2,4))
  for(v in 1:ncol(mtest$vectors)){
    
    plot(mtest$vectors[,v], col = clim$island,
         main = mtest$threshold)
    
  }

  
}
```

```
# 150 seems reasonable - axes 1 and 2 are basically north/south and east/west, higher axes are more fine-scale

# final version - calculate MEM

mem <- pcnm(dist, threshold = 150)


# add these to the climate dataframe
# only use the first half of the positive eigenvectors, following previous studies (eg Fitzpatrick & Keller 2014, Gugger et al 2017 koa paper)
pos_mem <- mem$values[mem$values>0] # the positive values
# keep half (round up if it's an odd number)
n_mem <- ceiling(length(pos_mem)/2)

# save both mem and climate variables to gradient forest input object
gfin <- cbind(bclim, mem$vectors[,1:n_mem])
colnames(gfin)
```

```
##  [1] "bio5"  "bio6"  "bio15" "bio18" "bio19" "elev"  "PCNM1" "PCNM2" "PCNM3"
## [10] "PCNM4" "PCNM5"
```

```
str(gfin)
```

```
## 'data.frame':    91 obs. of  11 variables:
##  $ bio5 : num  25.4 25.4 24.8 24.7 25.1 ...
##  $ bio6 : num  8.3 8.3 8.6 8.5 5.7 ...
##  $ bio15: num  98.6 98.6 98.7 96.1 97.5 ...
##  $ bio18: num  9 9 7 10 12 12 11 11 8 10 ...
##  $ bio19: num  199 199 154 175 215 215 275 276 177 194 ...
##  $ elev : num  370 370 361 475 594 594 480 478 140 304 ...
##  $ PCNM1: num  -0.1139 -0.1139 -0.1653 -0.1652 0.0533 ...
##  $ PCNM2: num  0.1115 0.1115 -0.1567 -0.157 0.0832 ...
##  $ PCNM3: num  -0.0187 -0.0187 -0.0741 -0.0744 -0.0803 ...
##  $ PCNM4: num  0.000942 0.00094 -0.032088 -0.031847 0.011255 ...
##  $ PCNM5: num  1.55e-03 5.36e-05 -3.04e-04 -2.55e-03 1.40e-02 ...
```

```
#save(gfin, snps, clim, file = 'gradient_forest_input_candidate_noGuad.rda')

# remove intermediate files
rm(snps, mtest)
```

### 2 Run Gradient Forest

Input files:

`gfin` is a dataframe with the climate and PCNM/MEM
variables as columns and samples as rows.

`snps.cand` are the candidate SNPs

```
# most of this code is modified from Fitzpatrick and Keller 2015, available on Dryad:
# https://datadryad.org/stash/dataset/doi:10.5061/dryad.2s6f9
 

maxLevel <- log2(0.368*nrow(gfin)/2) #account for correlations, see ?gradientForest 

# run gradient forest!

# gives warning
# "The response has five or fewer unique values.  Are you sure you want to do regression?"
# because values are just 0/1/2, not population-level allele freqs

gf.cand <- gradientForest(cbind(gfin, snps.cand), 
                     predictor.vars=colnames(gfin),
                     response.vars=colnames(snps.cand), 
                     ntree=500,
                     trace=T, 
                     corr.threshold=0.50,
                     maxLevel = maxLevel)
```

```
## Calculating forests for 560 species
## ...............................................................................
## ................................................................................
## ................................................................................
## ................................................................................
## ................................................................................
## ................................................................................
## ................................................................................
## .
```

```
gf.cand
```

```
## A forest of 500 regression trees for each of 533 species
## 
## Call:
## 
## gradientForest(data = cbind(gfin, snps.cand), predictor.vars = colnames(gfin), 
##     response.vars = colnames(snps.cand), ntree = 500, maxLevel = maxLevel, 
##     corr.threshold = 0.5, trace = T)
## 
## 
## 
## Important variables:
## [1] PCNM3 bio6  elev  PCNM4 PCNM1
```

```
# Important variables:
# [1] PCNM3 bio6  elev  PCNM4 PCNM1

#png(file = '/home/alayna/Documents/research/projects/2022_island_oak/results/gradient_forest/gradient_forest_importance_candidate_noGuadAna.png', height = 5, width = 7, res = 300, units = 'in')
plot(gf.cand, plot.type='O')
```

```
#dev.off()

plot(gf.cand, plot.type = 'S')
```

```
plot(gf.cand, plot.type = 'C')
```

```
plot(gf.cand, plot.type = 'C', show.species = F)
```

```
plot(gf.cand, plot.type = 'P')
```

```
# plot cumulative importance for all variables

#png(file = 'gradient_forest_cumulative_importance_candidate_SNPs.png', height = 9, width = 12, units = 'in', res = 300)
par(mfrow = c(3,4))
for(n in 1:ncol(gfin)){
  
  pvar <- colnames(gfin)[n]
  plot(cumimp(gf.cand, pvar),
       type = 'l',
       xlab = pvar,
       ylab = 'cumulative importance',
       main = pvar,
       ylim = c(0, 0.1))
  
  # add points to show density of actual values
  points(gfin[,n], rep(0, nrow(gfin)), pch = 16, col = rgb(0,0,0,0.2))
  
}
#dev.off()

# save
#save(gf.cand, file = 'data/clean/gradient_forest_results_candidate_SNPs_noGuadAna.rda')
```

### 3 Setup for mapping

#### 3.1 Load climate rasters

These are used to predict genomic composition at a given climate,
based on gf model

```
# first load one raster to get CRS info
bio1 <- rast('climate/bioclim/worldclim_2.1_30sec/wc2.1_30s_bio_1.tif')

# crop raster

# see example in crop()
# get extent - includes california islands
ext.ca <- ext(c(-121,-117.5,32.5,35))
bio1 <- crop(bio1, ext.ca)

plot(bio1)
```

```
# now make extent objects to crop out mainland and get only values for the islands

# northern
isl.n <- ext(c(-120.3, -119.3, 33.8, 34.1))
bio1.n <- crop(bio1, isl.n)
plot(bio1.n)
```

```
# southern
isl.s <- ext(c(-118.7, -118.1, 32.7, 33.5))
bio1.s <- crop(bio1, isl.s)
plot(bio1.s)
```

```
# combine northern and southern
bio1 <- merge(bio1.n, bio1.s)
plot(bio1)
```

```
# bio1 is a SpatRaster object that only includes the California islands

# now get all the clim rasters used in the gf model

dput(vars)
```

```
## c("bio5", "bio6", "bio15", "bio18", "bio19", "elev")
```

```
# files include underscore, add by hand
vars_file <- c("bio_5", "bio_6", "bio_15", "bio_18", "bio_19", "elev")

cells_list <- list()

for(n in 1:length(vars_file)){
  
  ra <- rast(paste('climate/bioclim/worldclim_2.1_30sec/wc2.1_30s_', vars_file[n], '.tif', sep = ''))
  
  # crop to northern and southern islands, then re-combine them
  # northern
  ra.n <- crop(ra, isl.n)
  #plot(ra.n)
  # southern
  ra.s <- crop(ra, isl.s)
  #plot(ra.s)
   
  ra <- merge(ra.n, ra.s)
  
  plot(ra, main = vars_file[n])
  
  # extract values for islands from raster
  df <- extract(ra, ext(ra), cells = T, xy = F)
  colnames(df)[2] <- vars[n]
  
  cells_list[[n]] <- df
  
}
```

```
#extract another copy, this time with the lat/lon of the center of the cell
cells_with_xy <- terra::extract(ra, ext(ra), cells = T, xy = T)

# check that they all have the same number of cells
lapply(cells_list, dim)
```

```
## [[1]]
## [1] 44352     2
## 
## [[2]]
## [1] 44352     2
## 
## [[3]]
## [1] 44352     2
## 
## [[4]]
## [1] 44352     2
## 
## [[5]]
## [1] 44352     2
## 
## [[6]]
## [1] 44352     2
```

```
# [1] 44352     2

# combine the extracted climate values into a dataframe
# first two
cells <- merge(cells_list[[1]], cells_list[[2]], by = 'cell')
# then loop through rest

for(n in 3:length(cells_list)){
  
  cells <- merge(cells, cells_list[[n]], by = 'cell')
  
}

# add xy values
cells$x <- cells_with_xy$x
cells$y <- cells_with_xy$y

# look at correlations
plot(cells)
```

```
# cell numbers we want to use
cellNums <- cells$cell[complete.cases(cells)]

head(cells[cellNums,])
```

```
##     cell bio5 bio6     bio15 bio18 bio19 elev         x        y
## 838  838 21.3  8.0 100.10389    19   242   63 -119.9208 34.07083
## 839  839 21.6  7.6  99.37767    21   252  140 -119.9125 34.07083
## 840  840 22.0  7.5 100.49244    18   249  125 -119.9042 34.07083
## 841  841 22.3  7.4 100.91532    18   247   99 -119.8958 34.07083
## 842  842 22.6  7.3 101.01292    18   235  115 -119.8875 34.07083
## 843  843 22.9  7.3 101.37901    18   234  103 -119.8792 34.07083
```

```
# set up empty raster, for saving turnover results to

# using the most recently read raster from the loop above as a template
# values are the cell numbers with values = T
#rast_blank <- rasterFromCells(ra, cellNums, values = T)
rast_blank <- rast(ra, vals = NA)


# check number of cells
dim(cells)
```

```
## [1] 44352     9
```

```
ncell(rast_blank)
```

```
## [1] 44352
```

```
# cells is a dataframe with the climate values for the islands, the cell number, and the lat/lon of the cell center
str(cells)
```

```
## 'data.frame':    44352 obs. of  9 variables:
##  $ cell : num  1 2 3 4 5 6 7 8 9 10 ...
##  $ bio5 : num  NA NA NA NA NA NA NA NA NA NA ...
##  $ bio6 : num  NA NA NA NA NA NA NA NA NA NA ...
##  $ bio15: num  NA NA NA NA NA NA NA NA NA NA ...
##  $ bio18: num  NA NA NA NA NA NA NA NA NA NA ...
##  $ bio19: num  NA NA NA NA NA NA NA NA NA NA ...
##  $ elev : int  NA NA NA NA NA NA NA NA NA NA ...
##  $ x    : num  -120 -120 -120 -120 -120 ...
##  $ y    : num  34.1 34.1 34.1 34.1 34.1 ...
```

```
# rast_blank is an empty raster with the same extent, for saving the genomic turnover to
str(rast_blank)
```

```
## S4 class 'SpatRaster' [package "terra"]
```

```
# cellNums is a vector containing the cell numbers that we have climate for (they are on the islands and not NA)
# get only these cells by subsetting cells df
str(cells[cellNums,])
```

```
## 'data.frame':    1157 obs. of  9 variables:
##  $ cell : num  838 839 840 841 842 ...
##  $ bio5 : num  21.3 21.6 22 22.3 22.6 ...
##  $ bio6 : num  8 7.6 7.5 7.4 7.3 ...
##  $ bio15: num  100.1 99.4 100.5 100.9 101 ...
##  $ bio18: num  19 21 18 18 18 18 18 18 8 19 ...
##  $ bio19: num  242 252 249 247 235 234 229 228 226 225 ...
##  $ elev : int  63 140 125 99 115 103 72 58 45 47 ...
##  $ x    : num  -120 -120 -120 -120 -120 ...
##  $ y    : num  34.1 34.1 34.1 34.1 34.1 ...
```

```
# add island name to cells

# I tried to do this using spatial objects but couldn't get the SpatExtent and coordinates to work together
# instead, just check if each cell coordinate is within the island extents
# this will assign some ocean cells to an island, but those are NAs and get removed anyway

# extents for islands
# this is duplicated below, need to remove when finished
# Santa Rosa
ext.sri <- ext(c(-120.25, -119.95, 33.85, 34.05))
# Santa Cruz
ext.sci <- ext(c(-119.95, -119.5, 33.7, 34.1))
# Anacapa
ext.ana <- ext(c(-119.455, -119.35, 33.9, 34.1))
# Catalina
ext.cat <- ext(c(-118.65, -118.25, 33.25, 33.5))
# San Clemente
ext.scl <- ext(c(-118.65, -118.25, 32.7, 33.1))

ext.all <- list(ext.sri, ext.sci, ext.ana, ext.cat, ext.scl)
names(ext.all) <- c('Santa_Rosa', 'Santa_Cruz', 'Anacapa', 'Catalina', 'San_Clemente')


island <- rep(NA, length = nrow(cells))
names(island) <- cells$cell

for(n in 1:length(ext.all)){
  
  bord <- as.vector(ext.all[[n]])
  # check which cells are within this border
  
  index <- which(with(cells, y < bord['ymax'] & y > bord['ymin'] & x > bord['xmin'] & x < bord['xmax']))
  
  island[index] <- names(ext.all)[n]

}

cells$island <- island
rm(island)
```

#### 3.2 Mapping functions

```
# again modified from Fitzpatrick and Keller 2015 code

# Mapping spatial genetic variation --------------------------------------------
###### functions to support mapping #####
# builds RGB raster from transformed environment
# snpPreds = dataframe of transformed variables from gf or gdm model
# rast = a raster mask to which RGB values are to be mapped
# cellNums = cell IDs to which RGB values should be assigned


pcaToRaster <- function(snpPreds, rast, mapCells){
  #require(raster)
  
  pca <- prcomp(snpPreds, center=TRUE, scale.=FALSE)
    
  ##assigns to colors, edit as needed to maximize color contrast, etc.
  a1 <- pca$x[,1]; a2 <- pca$x[,2]; a3 <- pca$x[,3]
  # original scaling from F&K script:
  #r <- a1+a2; g <- -a2; b <- a3+a2-a1
  # here just sent them to values of pca
  r <- a1
  g <- a2
  b <- a3
  
  ##scales colors
  scalR <- (r-min(r))/(max(r)-min(r))*255
  scalG <- (g-min(g))/(max(g)-min(g))*255
  scalB <- (b-min(b))/(max(b)-min(b))*255
  
  ##assigns color to raster
  rast1 <- rast2 <- rast3 <- rast
  rast1[mapCells] <- scalR
  rast2[mapCells] <- scalG
  rast3[mapCells] <- scalB
  
  # vector of colors
  cols <- rgb(scalR, scalG, scalB, maxColorValue = 255)
  ##stacks color rasters
  outRast <- rast(list(rast1, rast2, rast3))
  #return(outRast)
  return(list(raster = outRast, cols = cols, pca = pca))
}


# Function to map difference between spatial genetic predictions
# predMap1 = dataframe of transformed variables from gf or gdm model for first set of SNPs
# predMap2 = dataframe of transformed variables from gf or gdm model for second set of SNPs
# rast = a raster mask to which Procrustes residuals are to be mapped
# mapCells = cell IDs to which Procrustes residuals values should be assigned
RGBdiffMap <- function(predMap1, predMap2, rast, mapCells){
  require(vegan)
  PCA1 <- prcomp(predMap1, center=TRUE, scale.=FALSE)
  PCA2 <- prcomp(predMap2, center=TRUE, scale.=FALSE)
  diffProcrust <- procrustes(PCA1, PCA2, scale=TRUE, symmetrical=FALSE)
  residMap <- residuals(diffProcrust)
  rast[mapCells] <- residMap
  return(list(max(residMap), rast))
}
```

### 4 Mapping

#### 4.1 Genomic turnover

```
# OK, on to mapping. Script assumes:
# (1) a dataframe named env_trns containing extracted raster data (w/ cell IDs)
# and env. variables used in the models & with columns as follows: cell, bio1, bio2, etc.
#
# (2) a raster mask of the study region to which the RGB data will be written

# just get cells with no NAs for climate
env_trns <- cells[complete.cases(cells),]

# transform env using gf models, see ?predict.gradientForest
pred <- predict(gf.cand, env_trns[,vars])

# map continuous variation
color_map <- pcaToRaster(pred, rast_blank, cellNums)
map <- color_map$raster

# the plot!

# get shapefile for the island coastlines
coast <- read_sf('shapefiles/channel_islands_shapefile/xw602fs2985.shp')

# png(file = 'results/gradient_forest/noGuadAna/turnover_map_candSNPs_CAislands_gradient_forest.png', height = 8, width = 10, res = 300, units = 'in')
# pdf(file = 'results/gradient_forest/noGuadAna/turnover_map_candSNPs_CAislands_gradient_forest.pdf', height = 8, width = 10)

par(mfrow = c(1,1), mar = c(5,4,4,3))

# first plot borders to set the axes
plot(coast$geometry)
# add cell colors
plotRGB(map, smooth = F, add = T)
# add borders again since cells covered them
plot(coast$geometry, add = T)

# add island labels
text_size <- 1.3
text(-120.12, 33.85, labels = 'Santa Rosa', cex = text_size)
text(-119.75, 33.85, 'Santa Cruz', cex = text_size)
text(-119.4, 33.85, 'Anacapa', cex = text_size)
text(-118.7, 33.4, 'Catalina', cex = text_size)
text(-118.8, 32.9, 'San Clemente', cex = text_size)

# add sample locations
points(clim$lon, clim$lat, pch = 1, cex = 0.7, col = rgb(0,0,0,0.5))
```

```
#dev.off()

# function to plot as panels


# set up extents for islands
# Santa Rosa
ext.sri <- ext(c(-120.25, -119.95, 33.85, 34.05))
# Santa Cruz
ext.sci <- ext(c(-119.93, -119.5, 33.9, 34.1))
# Anacapa
ext.ana <- ext(c(-119.455, -119.35, 33.98, 34.04))
# Catalina
ext.cat <- ext(c(-118.65, -118.27, 33.27, 33.5))
# San Clemente
ext.scl <- ext(c(-118.65, -118.3, 32.76, 33.05))

ext.all <- list(ext.sri, ext.sci, ext.ana, ext.cat, ext.scl)
names(ext.all) <- c('Santa Rosa', 'Santa Cruz', 'Anacapa', 'Catalina', 'San Clemente')

plot_map_panel_rgb <- function(map, extents){
  
  n_isl <- length(extents)
  
  par(mfrow = c(1, n_isl), mar = c(5,4.5,4,2))
  
  for(n in 1:n_isl){
    
    extent <- extents[[n]]
    
    # first plot island borders to set the axes
    plot(coast$geometry, 
         extent = extent, 
         main = names(ext.all)[n], 
         axes = T,
         cex.main = 2, 
         cex.axis = 1,
         las = 2)
    #plot(map, add = T)
    # add cell colors
    terra::plotRGB(map,
          r=1, g=2, b=3,
         smooth = F,
         add = T,
         legend = ifelse(n = 1, T, F))
   # add borders again since cells covered them
    plot(coast$geometry, add = T)
    # add sample locations
    points(clim$lon, clim$lat, pch = 1, cex = 2, col = rgb(0,0,0,0.5))
    
    # attempting to add a scalebar, but 
    # sbar is from terra and doesn't plot when sf is used to plot first
    #sbar(10, xy = 'bottom', type = 'bar', below = 'km', cex = 1.5, adj = c(0.5, 1.5))
    
  }  
  
  # reset mfrow
  par(mfrow = c(1,1))
}


# note: this looks horrendous in the plot window but works when saving to a file
# probably because it uses both sf and terra to plot and they don't work well together

# png(file = 'results/gradient_forest/noGuadAna/map_genomic_turnover_candidate_SNPs.png', height = 4, width = 16, res = 300, units = 'in')
# pdf(file = 'results/gradient_forest/noGuadAna/map_genomic_turnover_candidate_SNPs.pdf', height = 4, width = 16)

plot_map_panel_rgb(map, extent = ext.all)
```

```
#dev.off()

# writeRaster(refRGBmap, "/.../refSNPs_map.tif", format="GTiff", overwrite=TRUE)
```

```
# ggplot version of map panel above
# I didn't end up using this version

library(ggspatial)
library(gridExtra) #grid.arrange
library(gtable)  #grid.draw
library(cowplot)  #plot_grid

# plot each of three layers separately
names(map) <- c(1,2,3)
ggplot() +  geom_spatraster(data = map) + facet_wrap(~lyr, nrow = 1)

ggplot() + geom_spatraster_rgb(data = map, interpolate = F) + geom_sf(data = coast$geometry, col = 'black', fill = 'transparent', lwd = 0.5) + geom_point(aes(x = clim$lon, y = clim$lat), pch = 1, col = rgb(0,0,0,0.5), cex = 1.5)

# just one island
# santa cruz
p1 <- ggplot() + geom_spatraster_rgb(data = map, interpolate = F,) + geom_sf(data = coast$geometry, col = 'black', fill = 'transparent', lwd = 0.5) + geom_point(aes(x = clim$lon, y = clim$lat), pch = 1, col = rgb(0,0,0,0.5), cex = 1.5) + coord_sf(xlim = c(-119.93, -119.5), ylim = c(33.9, 34.1)) + annotation_scale()

#catalina
p2 <- ggplot() + geom_spatraster_rgb(data = map, interpolate = F,) + geom_sf(data = coast$geometry, col = 'black', fill = 'transparent', lwd = 0.5) + geom_point(aes(x = clim$lon, y = clim$lat), pch = 1, col = rgb(0,0,0,0.5), cex = 1.5) + coord_sf(xlim = c(-118.65, -118.25), ylim = c(33.25, 33.5)) + annotation_scale()

#plot the islands as panels
grid.arrange(p1, p2, nrow = 1)

g1 <- ggplotGrob(p1)
g2 <- ggplotGrob(p2)

two_plots <- cbind(g1,g2, size = 'first')

grid.newpage()
grid.draw(two_plots)

plot_grid(p1, p2, align = 'v', nrow = 2)


#ggplot(map) + geom_spatraster_rgb(data = map, interpolate = F) + geom_sf(coast)


# function
# this takes extents because I already have them set up, but it's not really needed, just need to give ggplot xlim and ylim
ggplot_map_panel_rgb <- function(map, extents){
  
  n_isl <- length(extents)
  
  # list to save plots to
  maps <- list()
  
  for(n in 1:n_isl){
    
    extent <- extents[[n]]
    
    p <- ggplot() + 
      geom_spatraster_rgb(data = map, interpolate = F,) + 
      geom_sf(data = coast$geometry, col = 'black', fill = 'transparent', lwd = 0.5) +
      geom_point(aes(x = clim$lon, y = clim$lat), pch = 1, col = rgb(0,0,0,0.5), cex = 1.5) +
      coord_sf(xlim = c(extent[1], extent[2]), ylim = c(extent[3], extent[4])) + 
      annotation_scale() +
      ggtitle(names(extents)[n]) +
      xlab(NULL) +
      ylab(NULL)
    
    maps[[n]] <- ggplotGrob(p)

  }  
  
return(grid.arrange(grobs = maps, nrow = 1))

}


ggplot_map_panel_rgb(map, extent = ext.all)

g <- ggplot_map_panel_rgb(map, extent = ext.all)

#ggsave(file = 'results/gradient_forest/noGuadAna/map_genomic_turnover_candidate_SNPs_ggplot.png', g, height = 4, width = 16, dpi = 300, units = 'in')
```

#### 4.2 Climate PCA

```
# plot PCA with predicted values for each cell, and loadings of scaled climate variables

pca <- color_map$pca
cols <- color_map$cols

# function for plotting
plot_clim_pca <- function(pca, axis_x, axis_y, varnames = vars, text_cex = 1.5){
  
  clim_scaled_x <- scales::rescale(pca$rotation[,axis_x], to = range(pca$x[,axis_x]))
  clim_scaled_y <- scales::rescale(pca$rotation[,axis_y], to = range(pca$x[,axis_y]))
  
  # add padding to axes
  pad <- 0.2
  xmin <- min(pca$x[,axis_x]) + pad*min(pca$x[,axis_x])
  xmax <- max(pca$x[,axis_x]) + pad*max(pca$x[,axis_x])
  ymin <- min(pca$x[,axis_y]) + pad*min(pca$x[,axis_y])
  ymax <- max(pca$x[,axis_y]) + pad*max(pca$x[,axis_y])
  
  plot(pca$x[,axis_x], pca$x[,axis_y], 
       col = cols, 
       pch = 16, 
       cex = 1.5,
       axes = T, 
       xlim = c(xmin, xmax),
       ylim = c(ymin, ymax),
       xlab = paste('PC', axis_x, sep = ''), 
       ylab = paste('PC', axis_y, sep = ''), 
       cex.axis = 1.2, 
       cex.lab = 1.5)
  
  text(clim_scaled_x, 
       clim_scaled_y,
       varnames,
       cex = text_cex,
       pos = 3,
       family = 'bold')
  # add a line from the origin to climate
  for(n in 1:length(clim_scaled_x)){
    arrows(0, 0, clim_scaled_x[n], clim_scaled_y[n], code = 2, length = 0.1, lwd = 1.5)
  }
  
}


par(mar = c(5,4,4,3), mfrow = c(1,2))
plot_clim_pca(pca, 1, 2)
plot_clim_pca(pca, 1, 3)
```

```
plot_clim_pca(pca, 1, 4)
plot_clim_pca(pca, 1, 5)
```

```
plot_clim_pca(pca, 1, 6)

# plot to save
#png(file = 'results/gradient_forest/noGuadAna/gf_scaled_climate_PCA_noGuadAna_PC-1-3_vertical.png', height = 8, width = 4, res = 300, units = 'in')
par(mfrow = c(2,1), mar = c(4,5,1,1), cex = 0.8)
```

```
plot_clim_pca(pca, 1, 2)
plot_clim_pca(pca, 1, 3)
```

```
#dev.off()

# with longer names
# rename in PCA
dput(rownames(pca$rotation))
```

```
## c("bio5", "bio6", "bio15", "bio18", "bio19", "elev")
```

```
pca_rename <- pca
rownames(pca_rename$rotation) <- c("Max Temp Warmest Month", "Min Temp Coldest Month", "Precip Seasonality", "Precip Warmest Quarter", "Precip Coldes Quarter", "Elevation")

# with newlines
varnames <- c("Max Temp\nWarmest Month", "Min Temp\nColdest Month", "Precipitation\nSeasonality", "Precipitation\nWarmest Quarter", "Precipitation\nColdest Quarter", "Elevation")

#varnames <- c("Max Temp Warmest Month", "Min Temp Coldest Month", "Precip Seasonality", "Precip Warmest Quarter", "Precip Coldest Quarter", "Elevation")

#png(file = 'results/gradient_forest/noGuadAna/gf_scaled_climate_PCA_noGuadAna_PC-1-3_longClimNames_vertical.png', height = 8, width = 4, res = 300, units = 'in')

# bold text isn't working with PDF
#pdf(file = 'results/gradient_forest/noGuadAna/gf_scaled_climate_PCA_noGuadAna_PC-1-3_longClimNames_vertical.pdf', height = 8, width = 4)
# Error in text.default(clim_scaled_x, clim_scaled_y, varnames, cex = text_cex,  : 
#   invalid font type
# In addition: Warning messages:
# 1: In text.default(clim_scaled_x, clim_scaled_y, varnames, cex = text_cex,  :
#   font family 'bold' not found in PostScript font database

par(mfrow = c(2,1), mar = c(4,5,1,1), cex = 0.8)
plot_clim_pca(pca, 1, 2, varnames = varnames, text_cex = 0.8)
plot_clim_pca(pca, 1, 3, varnames = varnames, text_cex = 0.8)
```

```
dev.off()
```

```
## null device 
##           1
```

```
#png(file = 'results/gradient_forest/noGuadAna/gf_scaled_climate_PCA_noGuadAna_PC-1-3_longClimNames_horizontal.png', height = 4, width = 8, res = 300, units = 'in')
par(mfrow = c(1,2), mar = c(4,5,1,1), cex = 0.8)
plot_clim_pca(pca, 1, 2, varnames = varnames, text_cex = 0.8)
```

```
## Warning in text.default(clim_scaled_x, clim_scaled_y, varnames, cex = text_cex,
## : font family 'bold' not found in PostScript font database

## Warning in text.default(clim_scaled_x, clim_scaled_y, varnames, cex = text_cex,
## : font family 'bold' not found in PostScript font database

## Warning in text.default(clim_scaled_x, clim_scaled_y, varnames, cex = text_cex,
## : font family 'bold' not found in PostScript font database

## Warning in text.default(clim_scaled_x, clim_scaled_y, varnames, cex = text_cex,
## : font family 'bold' not found in PostScript font database

## Warning in text.default(clim_scaled_x, clim_scaled_y, varnames, cex = text_cex,
## : font family 'bold' not found in PostScript font database

## Warning in text.default(clim_scaled_x, clim_scaled_y, varnames, cex = text_cex,
## : font family 'bold' not found in PostScript font database

## Warning in text.default(clim_scaled_x, clim_scaled_y, varnames, cex = text_cex,
## : font family 'bold' not found in PostScript font database

## Warning in text.default(clim_scaled_x, clim_scaled_y, varnames, cex = text_cex,
## : font family 'bold' not found in PostScript font database

## Warning in text.default(clim_scaled_x, clim_scaled_y, varnames, cex = text_cex,
## : font family 'bold' not found in PostScript font database

## Warning in text.default(clim_scaled_x, clim_scaled_y, varnames, cex = text_cex,
## : font family 'bold' not found in PostScript font database

## Warning in text.default(clim_scaled_x, clim_scaled_y, varnames, cex = text_cex,
## : font family 'bold' not found in PostScript font database

## Warning in text.default(clim_scaled_x, clim_scaled_y, varnames, cex = text_cex,
## : font family 'bold' not found in PostScript font database
```

```
plot_clim_pca(pca, 1, 3, varnames = varnames, text_cex = 0.8)
```

```
## Warning in text.default(clim_scaled_x, clim_scaled_y, varnames, cex = text_cex,
## : font family 'bold' not found in PostScript font database

## Warning in text.default(clim_scaled_x, clim_scaled_y, varnames, cex = text_cex,
## : font family 'bold' not found in PostScript font database

## Warning in text.default(clim_scaled_x, clim_scaled_y, varnames, cex = text_cex,
## : font family 'bold' not found in PostScript font database

## Warning in text.default(clim_scaled_x, clim_scaled_y, varnames, cex = text_cex,
## : font family 'bold' not found in PostScript font database

## Warning in text.default(clim_scaled_x, clim_scaled_y, varnames, cex = text_cex,
## : font family 'bold' not found in PostScript font database

## Warning in text.default(clim_scaled_x, clim_scaled_y, varnames, cex = text_cex,
## : font family 'bold' not found in PostScript font database

## Warning in text.default(clim_scaled_x, clim_scaled_y, varnames, cex = text_cex,
## : font family 'bold' not found in PostScript font database

## Warning in text.default(clim_scaled_x, clim_scaled_y, varnames, cex = text_cex,
## : font family 'bold' not found in PostScript font database

## Warning in text.default(clim_scaled_x, clim_scaled_y, varnames, cex = text_cex,
## : font family 'bold' not found in PostScript font database

## Warning in text.default(clim_scaled_x, clim_scaled_y, varnames, cex = text_cex,
## : font family 'bold' not found in PostScript font database

## Warning in text.default(clim_scaled_x, clim_scaled_y, varnames, cex = text_cex,
## : font family 'bold' not found in PostScript font database

## Warning in text.default(clim_scaled_x, clim_scaled_y, varnames, cex = text_cex,
## : font family 'bold' not found in PostScript font database
```

```
#dev.off()
```

#### 4.3 Genomic Offsets

```
# setup for offsets by getting rasters for future climate

# the four models I'm using
# MIROC and CNRM, 4.5 and 8.5, for 2050
future_string <- c('mr45bi50', 'mr85bi50', 'cn45bi50', 'cn85bi50')
future_names <- c('MIROC 4.5 2050', 'MIROC 8.5 2050', 'CNRM 4.5 2050', 'CNRM 8.5 2050')

# first load one raster to get CRS info
bio1 <- rast(paste('climate/bioclim/future/cmip5/', future_string[1], '1.tif', sep = ''))

# crop raster

# see example in crop()
# note: the bioclim raster still has to be mounted even though it's loaded into R
# get extent - includes california islands
ext.ca <- ext(c(-121,-117.5,32.5,35))
bio1 <- crop(bio1, ext.ca)

plot(bio1)
```

```
# now make extent objects to crop out mainland and get only values for the islands

# northern
isl.n <- ext(c(-120.3, -119.3, 33.8, 34.1))
bio1.n <- crop(bio1, isl.n)
plot(bio1.n)
```

```
# southern
isl.s <- ext(c(-118.7, -118.1, 32.7, 33.5))
bio1.s <- crop(bio1, isl.s)
plot(bio1.s)
```

```
# combine northern and southern
bio1 <- merge(bio1.n, bio1.s)
plot(bio1)
```

```
# bio1 is a SpatRaster object that only includes the California islands

# now get all the clim rasters used in the gf model

dput(vars)
```

```
## c("bio5", "bio6", "bio15", "bio18", "bio19", "elev")
```

```
# c("bio5", "bio6", "bio15", "bio18", "bio19", "elev")

# the future climates just have the bioclim number at the end
# also remove elevation since it isn't in the future projections
vars_file <- gsub('bio', '', vars)
vars_file <- vars_file[vars_file != 'elev']
dput(vars_file)
```

```
## c("5", "6", "15", "18", "19")
```

```
# c("5", "6", "15", "18", "19")


setup_climate <- function(future_string, vars_file){
  
  cells_list <- list()
  
  # first get all the climate rasters
  for(n in 1:length(vars_file)){
    
    ra <- rast(paste('climate/bioclim/future/cmip5/', future_string, vars_file[n], '.tif', sep = ''))
    
    # crop to northern and southern islands, then re-combine them
    # northern
    ra.n <- crop(ra, isl.n)
    #plot(ra.n)
    # southern
    ra.s <- crop(ra, isl.s)
    #plot(ra.s)
    
    ra <- merge(ra.n, ra.s)
    
    plot(ra, main = paste('bio', vars_file[n], '\n', future_string, sep = ''))
    
    # extract values for islands from raster
    df <- extract(ra, ext(ra), cells = T, xy = F)
    colnames(df)[2] <- vars[n]
    
    cells_list[[n]] <- df
    
  }
  
  #extract another copy, this time with the lat/lon of the center of the cell
  cells_with_xy <- terra::extract(ra, ext(ra), cells = T, xy = T)
  
  # add elevation separately, from current climate
  
  ra <- rast('climate/bioclim/worldclim_2.1_30sec/wc2.1_30s_elev.tif')
  
  # crop to northern and southern islands, then re-combine them
  # northern
  ra.n <- crop(ra, isl.n)
  #plot(ra.n)
  # southern
  ra.s <- crop(ra, isl.s)
  #plot(ra.s)
  
  ra <- merge(ra.n, ra.s)
  
  plot(ra, main = 'elevation')
  
  # extract values for islands from raster
  df <- extract(ra, ext(ra), cells = T, xy = F)
  colnames(df)[2] <- 'elev'
  
  # add as last element of list
  cells_list[[length(cells_list)+1]] <- df
  
  
  
  # check that they all have the same number of cells
  lapply(cells_list, dim)
  # [1] 44352     2
  
  # combine the extracted climate values into a dataframe
  # first two
  cells <- merge(cells_list[[1]], cells_list[[2]], by = 'cell')
  # then loop through rest
  
  for(n in 3:length(cells_list)){
    
    cells <- merge(cells, cells_list[[n]], by = 'cell')
    
  }
  
  # add xy values
  cells$x <- cells_with_xy$x
  cells$y <- cells_with_xy$y
  
  # look at correlations
  # this is slow, can comment out
  #plot(cells)
  
  return(cells)
}

# now loop through each future scenario and get cell info
futures <- list() 
for(n in 1:length(future_string)){
  
  futures[[n]] <- setup_climate(future_string[n], vars_file)
  names(futures)[n] <- gsub(' ', '_', future_names[n])
  
}
```

```
# set up empty raster, for saving turnover results to

# do the same as above - using elevation file as a template
# values are the cell numbers with values = T
ra <- rast('climate/bioclim/worldclim_2.1_30sec/wc2.1_30s_elev.tif')

# crop to northern and southern islands, then re-combine them
# northern
ra.n <- crop(ra, isl.n)
plot(ra.n)
```

```
# southern
ra.s <- crop(ra, isl.s)
plot(ra.s)
```

```
ra <- merge(ra.n, ra.s)

plot(ra)
```

```
# save to blank raster
rast_blank <- rast(ra, vals = NA)


# cell numbers we want to use - the ones without NAs
# here just doing it for the first scenario - I think it should be the same for all of them but check if this causes problems
# do this in plotting section?
cellNums <- futures[[1]]$cell[complete.cases(futures[[1]])]

head(futures[[1]][cellNums,])
```

```
##     cell bio5 bio6 bio15 bio18 bio19 elev         x        y
## 839  839  251   82    97     5   220  140 -119.9125 34.07083
## 840  840  252   83    98     4   216  125 -119.9042 34.07083
## 841  841  252   84    98     4   216   99 -119.8958 34.07083
## 842  842  252   82    99     5   219  115 -119.8875 34.07083
## 843  843  252   84    98     4   217  103 -119.8792 34.07083
## 844  844  252   87    97     4   195   72 -119.8708 34.07083
```

```
################################
# description of objects

# cellNums is a vector containing the cell numbers that we have climate for (they are on the islands and not NA)
# get only these cells by subsetting cells df
str(futures[[1]][cellNums,])
```

```
## 'data.frame':    1101 obs. of  9 variables:
##  $ cell : num  839 840 841 842 843 ...
##  $ bio5 : int  251 252 252 252 252 252 252 253 252 251 ...
##  $ bio6 : int  82 83 84 82 84 87 88 87 86 86 ...
##  $ bio15: int  97 98 98 99 98 97 97 97 99 99 ...
##  $ bio18: int  5 4 4 5 4 4 4 4 4 4 ...
##  $ bio19: int  220 216 216 219 217 195 195 175 198 204 ...
##  $ elev : int  140 125 99 115 103 72 58 45 47 89 ...
##  $ x    : num  -120 -120 -120 -120 -120 ...
##  $ y    : num  34.1 34.1 34.1 34.1 34.1 ...
```

```
# check number of cells
lapply(futures, dim)
```

```
## $MIROC_4.5_2050
## [1] 44352     9
## 
## $MIROC_8.5_2050
## [1] 44352     9
## 
## $CNRM_4.5_2050
## [1] 44352     9
## 
## $CNRM_8.5_2050
## [1] 44352     9
```

```
ncell(rast_blank)
```

```
## [1] 44352
```

```
# futures is a list of the climate scenarios
# each one is a dataframe with the future climate values for the islands under that scenario, the cell number, and the lat/lon of the cell center
lapply(futures, str)
```

```
## 'data.frame':    44352 obs. of  9 variables:
##  $ cell : num  1 2 3 4 5 6 7 8 9 10 ...
##  $ bio5 : int  NA NA NA NA NA NA NA NA NA NA ...
##  $ bio6 : int  NA NA NA NA NA NA NA NA NA NA ...
##  $ bio15: int  NA NA NA NA NA NA NA NA NA NA ...
##  $ bio18: int  NA NA NA NA NA NA NA NA NA NA ...
##  $ bio19: int  NA NA NA NA NA NA NA NA NA NA ...
##  $ elev : int  NA NA NA NA NA NA NA NA NA NA ...
##  $ x    : num  -120 -120 -120 -120 -120 ...
##  $ y    : num  34.1 34.1 34.1 34.1 34.1 ...
## 'data.frame':    44352 obs. of  9 variables:
##  $ cell : num  1 2 3 4 5 6 7 8 9 10 ...
##  $ bio5 : int  NA NA NA NA NA NA NA NA NA NA ...
##  $ bio6 : int  NA NA NA NA NA NA NA NA NA NA ...
##  $ bio15: int  NA NA NA NA NA NA NA NA NA NA ...
##  $ bio18: int  NA NA NA NA NA NA NA NA NA NA ...
##  $ bio19: int  NA NA NA NA NA NA NA NA NA NA ...
##  $ elev : int  NA NA NA NA NA NA NA NA NA NA ...
##  $ x    : num  -120 -120 -120 -120 -120 ...
##  $ y    : num  34.1 34.1 34.1 34.1 34.1 ...
## 'data.frame':    44352 obs. of  9 variables:
##  $ cell : num  1 2 3 4 5 6 7 8 9 10 ...
##  $ bio5 : int  NA NA NA NA NA NA NA NA NA NA ...
##  $ bio6 : int  NA NA NA NA NA NA NA NA NA NA ...
##  $ bio15: int  NA NA NA NA NA NA NA NA NA NA ...
##  $ bio18: int  NA NA NA NA NA NA NA NA NA NA ...
##  $ bio19: int  NA NA NA NA NA NA NA NA NA NA ...
##  $ elev : int  NA NA NA NA NA NA NA NA NA NA ...
##  $ x    : num  -120 -120 -120 -120 -120 ...
##  $ y    : num  34.1 34.1 34.1 34.1 34.1 ...
## 'data.frame':    44352 obs. of  9 variables:
##  $ cell : num  1 2 3 4 5 6 7 8 9 10 ...
##  $ bio5 : int  NA NA NA NA NA NA NA NA NA NA ...
##  $ bio6 : int  NA NA NA NA NA NA NA NA NA NA ...
##  $ bio15: int  NA NA NA NA NA NA NA NA NA NA ...
##  $ bio18: int  NA NA NA NA NA NA NA NA NA NA ...
##  $ bio19: int  NA NA NA NA NA NA NA NA NA NA ...
##  $ elev : int  NA NA NA NA NA NA NA NA NA NA ...
##  $ x    : num  -120 -120 -120 -120 -120 ...
##  $ y    : num  34.1 34.1 34.1 34.1 34.1 ...
```

```
## $MIROC_4.5_2050
## NULL
## 
## $MIROC_8.5_2050
## NULL
## 
## $CNRM_4.5_2050
## NULL
## 
## $CNRM_8.5_2050
## NULL
```

```
# rast_blank is an empty raster with the same extent, for saving the genomic turnover to
str(rast_blank)
```

```
## S4 class 'SpatRaster' [package "terra"]
```

```
# From Fitzpatrick and Keller script 

# Calculate and map "genetic offset" under climate change ----------------------
# Script assumes:
  # (1) a dataframe of transformed env. variables for CURRENT climate 
  # (e.g., predGI5 from above).
  #
  # (2) a dataframe named env_trns_future containing extracted raster data of 
  # env. variables for FUTURE a climate scenario, same structure as env_trns

# loop through future scenarios

offsets <- list()

scaled_gf_clim <- list()

for(n in 1:length(futures)){
  
  fut <- futures[[n]]
  
  env_trns_future <- fut[complete.cases(fut),]
  
  # some of the cells with present data are missing in the future data
  env_trns[! env_trns$cell %in% env_trns_future$cell,]
  # remove them
  env_trns <- env_trns[env_trns$cell %in% env_trns_future$cell, ]

  
  # check for mismatches
  sum(env_trns$cell != env_trns_future$cell)
  
  # convert rownames (cell numbers) from character to numeric
  # this isn't working?
  # rownames(env_trns) <- as.numeric(rownames(env_trns))
  # rownames(env_trns_future) <- as.numeric(rownames(env_trns_future))
  
  # rerun prediction for present climate
  hist.cand <- predict(gf.cand, env_trns[,vars])
  
  # prediction for future climate
  fut.cand <- predict(gf.cand, env_trns_future[,vars])
  # save for using later
  scaled_gf_clim[[n]] <- fut.cand
  
  # calculate euclidean distance between current and future genetic spaces  
  genOffset <- sqrt((fut.cand[,1]-hist.cand[,1])^2+(fut.cand[,2]-hist.cand[,2])^2
                    +(fut.cand[,3]-hist.cand[,3])^2+(fut.cand[,4]-hist.cand[,4])^2
                    +(fut.cand[,5]-hist.cand[,5])^2+(fut.cand[,6]-hist.cand[,6])^2)
  
  # assign values to raster - can be tricky if current/future climate
  # rasters are not identical in terms of # cells, extent, etc.
  offset <- rast_blank
  offset[cellNums] <- genOffset
  
  offsets[[n]] <- offset
  
}

# add historic climate to the list of scaled climates
scaled_gf_clim[[length(scaled_gf_clim)+1]] <- hist.cand
names(scaled_gf_clim) <- c(names(futures), 'historic')

names(offsets) <- names(futures)

pal <- colorRampPalette(brewer.pal(9, 'YlOrRd'))

for(n in 1:length(offsets)){
  
  offset <- offsets[[n]]
  
  #png(file = 'genetic_offset_map_cand_SNPs_MIROC85.png', height = 8, width = 12, res = 300, units = 'in')
  
  par(mfrow = c(1,1), mar = c(5,4,4,3))
  
  # first plot island borders to set the axes
  plot(coast$geometry, main = future_names[n])
  # add cell colors
  plot(offset, smooth = F,  col = pal(100), add = T)
  # add borders again since cells covered them
  plot(coast$geometry, add = T)
  # add sample locations
  points(clim$lon, clim$lat, pch = 1, cex = 0.7, col = rgb(0,0,0,0.5))
  
  # add island labels
  text_size <- 1.3
  text(-120.12, 33.85, labels = 'Santa Rosa', cex = text_size)
  text(-119.75, 33.85, 'Santa Cruz', cex = text_size)
  text(-119.4, 33.85, 'Anacapa', cex = text_size)
  text(-118.7, 33.4, 'Catalina', cex = text_size)
  text(-118.8, 32.9, 'San Clemente', cex = text_size)
  
  sbar(40, xy = 'bottomleft', divs = 4, lonlat = T, type = 'bar', below = 'km', cex = 1.5, adj = c(0.5, 1.5))
  
  #dev.off()
  
  
}
```

```
# version with the same color scale for all climate projections, for easier comparison
# get min and max when all rasters are considered
min <- min(sapply(1:length(offsets), function(n) minmax(offsets[[n]])[1]))
max <- max(sapply(1:length(offsets), function(n) minmax(offsets[[n]])[2]))

# colf <- colorRamp2(breaks = seq(from = min, to = max, length.out = 9),
#                    colors = brewer.pal(9, 'YlOrRd'))


for(n in 1:length(offsets)){
  
  offset <- offsets[[n]]
  
  # png(file = paste('results/gradient_forest/noGuadAna/genetic_offset_full_map_cand_SNPs_', names(offsets)[n], '.png'),
  #     height = 8, 
  #     width = 12, 
  #     res = 300, 
  #     units = 'in')
  
  par(mfrow = c(1,1), mar = c(5,4,4,3))
  
  # first plot island borders to set the axes
  plot(coast$geometry, main = future_names[n], axes = T)
  # add cell colors
  plot(offset, smooth = F,  
       col = pal(100), 
       breaks = seq(from = min, to = max, length.out = 100), 
       add = T, 
       legend = F)
  # add borders again since cells covered them
  plot(coast$geometry, add = T)
  # add sample locations
  points(clim$lon, clim$lat, pch = 1, cex = 0.7, col = rgb(0,0,0,0.5))
  
  # add island labels
  text_size <- 1.3
  text(-120.12, 33.85, labels = 'Santa Rosa', cex = text_size)
  text(-119.75, 33.85, 'Santa Cruz', cex = text_size)
  text(-119.4, 33.85, 'Anacapa', cex = text_size)
  text(-118.7, 33.4, 'Catalina', cex = text_size)
  text(-118.8, 32.9, 'San Clemente', cex = text_size)
  
  
  # hacky way to add legend
  # source: https://stackoverflow.com/a/70522655
  legend_image <- as.raster(matrix(rev(pal(100)), ncol=1))
  
  #figSet <- c(grconvertX(c(120.5, 120), from="user", to="ndc"), grconvertY(c(33, 34), from="user", to="ndc"))
  figSet <- c(0.1, 0.35, 0.15, 0.6)
  
  ## layer 2, legend inside
  op <- par(  ## set and store par
    fig = figSet, 
    mar = c(1, 1, 1, 9.5), ## set margins
    new = TRUE) ## set new for overplot w/ next plot
  
  plot(c(0, 2), c(0, 1), type='n', axes=F, xlab='', ylab='')  ## ini plot2
  rasterImage(legend_image, 0, 0, 1, 1) ## the gradient
  lbsq <- seq.int(0, 1, l=5)   ## seq. for labels
  axis(4, at=lbsq, pos=1, labels=F, col=0, col.ticks=1, tck=-.1)  ## axis ticks
  mtext(round(seq.int(min, max, l=5), 2), 4, -.5, at=lbsq, las=2, cex=.8)  ## tick labels
  mtext('Offset', 3, -.125, cex=1, adj=.1, font=2)  ## legend title
  
  par(op)  ## reset par
  
  
  #dev.off()
  
  
}
```

```
# function to plot as panels


# set up extents for islands
# Santa Rosa
ext.sri <- ext(c(-120.25, -119.95, 33.85, 34.05))
# Santa Cruz
ext.sci <- ext(c(-119.93, -119.5, 33.9, 34.1))
# Anacapa
ext.ana <- ext(c(-119.455, -119.35, 33.98, 34.04))
# Catalina
ext.cat <- ext(c(-118.65, -118.27, 33.27, 33.5))
# San Clemente
ext.scl <- ext(c(-118.65, -118.3, 32.76, 33.05))

ext.all <- list(ext.sri, ext.sci, ext.ana, ext.cat, ext.scl)
names <- c('Santa Rosa', 'Santa Cruz', 'Anacapa', 'Catalina', 'San Clemente')


plot_map_panel <- function(map, extents){
  
  n_isl <- length(extents)
  
  par(mfrow = c(1, n_isl), mar = c(5,4.5,4,2))
  
  for(n in 1:n_isl){
    
    extent <- extents[[n]]
    
    # first plot island borders to set the axes
    plot(coast$geometry, 
         extent = extent, 
         main = names[n], 
         axes = T,
         cex.main = 2, 
         cex.axis = 1, 
         las = 2)
    # add cell colors
    plot(map, 
         smooth = F,  
         col = pal(100),
         breaks = seq(from = min, to = max, length.out = 100),
         add = T,
         legend = ifelse(n = 1, T, F))
    # add borders again since cells covered them
    plot(coast$geometry, add = T)
    # add sample locations
    points(clim$lon, clim$lat, pch = 1, cex = 2, col = rgb(0,0,0,0.5))
    
    
    # add legend to first plot
    if(n == n_isl){
      
      # hacky way to add legend
      # source: https://stackoverflow.com/a/70522655
      legend_image <- as.raster(matrix(rev(pal(100)), ncol=1))
      
      # position of the legend, which is plotted as an inset plot
      # had to play with these values
      figSet <- c(0.85, 0.95, 0.2, 0.55)
      
      ## layer 2, legend inside
      op <- par(  ## set and store par
        fig = figSet, 
        mar = c(0.5, 0.5, 0.5, 9.5), ## set margins
        new = TRUE) ## set new for overplot w/ next plot
      
      plot(c(0, 2), c(0, 1), type='n', axes=F, xlab='', ylab='')  ## ini plot2
      rasterImage(legend_image, 0, 0, 1, 1) ## the gradient
      lbsq <- seq.int(0, 1, l=5)   ## seq. for labels
      axis(4, at=lbsq, pos=1, labels=F, col=0, col.ticks=1, tck=-.1)  ## axis ticks
      mtext(round(seq.int(min, max, l=5), 2), 4, -.5, at=lbsq, las=2, cex=0.7)  ## tick labels
      mtext('Offset', 3, -.12, cex=0.8, adj=1, font=2)  ## legend title
      
      par(op)  ## reset par
      
      
    }
    
  }  
  # reset mfrow
  par(mfrow = c(1,1))

}


# version with the same color scale
# get min and max when all rasters are considered
min <- min(sapply(1:length(offsets), function(n) minmax(offsets[[n]])[1]))
max <- max(sapply(1:length(offsets), function(n) minmax(offsets[[n]])[2]))

for(n in 1:length(offsets)){
  
  offset <- offsets[[n]]
  
  #png(file = paste('results/gradient_forest/noGuadAna/offset_panel_', names(offsets)[n], '_2050.png', sep = ''), height = 4, width = 16, res = 300, units = 'in')
  #pdf(file = paste('results/gradient_forest/noGuadAna/offset_panel_', names(offsets)[n], '_2050.pdf', sep = ''), height = 4, width = 16)

  # comment out this line in order to knit
  #plot_map_panel(offsets[[n]], extents = ext.all)
  #dev.off()
  
}


# look at using this 
#https://stackoverflow.com/questions/76072271/manage-subplots-titles-in-multiple-rastervis-levelplot
#library(rasterVis)
```

### 5 Offset calculations

#### 5.1 Offsets for specific populations, without movement

```
# look at offsets for specific 'populations' (cells in raster where trees were collected)


# get coords
clim.sub <- clim[clim$island %in% c("Santa Rosa Island", "Santa Cruz Island", "Anacapa Island", "Catalina Island", "San Clemente Island"),]

df <- clim.sub[,c('lat', 'lon')]
coords <- vect(df, geom=c("lon", "lat"))

off.pop <- as.data.frame(matrix(nrow = nrow(clim.sub), ncol = c(10)))
rownames(off.pop) <- clim.sub$ID_vcf
colnames(off.pop) <- c('island', 'lat', 'lon', 'cell', 'cell_x', 'cell_y', names(offsets))

off.pop$island <- clim.sub$island
off.pop$lat <- clim.sub$lat
off.pop$lon <- clim.sub$lon


for(n in 1:length(offsets)){
  
  offset <- offsets[[n]]
  offset_name <- names(offsets)[n]

  # extract offsets for those coords
  
  off.tmp <- extract(offset, coords, xy = T, cells = T)
  colnames(off.tmp) <- c('ID',  offset_name, 'cell', 'x', 'y')
  
  off.pop[,offset_name] <- off.tmp[,offset_name]
  #  add cell info - this overwrites each time which isn't ideal but should be fine since all offset rasters are the same
  off.pop[,'cell'] <- off.tmp[,'cell']
  off.pop[,'cell_x'] <- off.tmp[,'x']
  off.pop[,'cell_y'] <- off.tmp[,'y']
  
}
  

for(n in 1:length(offsets)){
  
  offset <- names(offsets)[n]
  
  #png(file = paste('results/gradient_forest/offset_boxplot_by_island_pops_', offset, '.png', sep = ''), height = 6, width = 7, res = 300, units = 'in')
  par(mar = c(10,5,4,3))
  boxplot(off.pop[,offset] ~ clim.sub$island, las = 2, drop = T,
          border = c("#3c4a8b", "#009c85", "#84bc5f", "#edb829", "#f57404"),
          col = 'white',
          xlab = '',
          ylab = 'Offset',
          main = future_names[n])
  #dev.off()
  
}
```

#### 5.2 Offsets with assisted gene flow

```
# Calculate pairwise offsets between potential seed sources (historic climates of sampled populations) and planting sites (all cells on the islands, including sampled sites)

# get the environmental variables for our collection sites
# the rownames of the env_trns objects are the cell numbers, but as a character

# get the cells where we collected trees, removing duplicates
off.pop.sort <- off.pop[order(off.pop$island),]

# get row index of unique cells
unq <- !duplicated(off.pop.sort$cell)
pop_cells <- as.character(off.pop.sort[unq, 'cell'])
pop_islands <- off.pop.sort[unq, 'island']

# function for calculating offset
# this needs to be the scaled gf values, not raw climate values
calc_offset <- function(current, future){
  
  sqrt((future[,1]-current[,1])^2+(future[,2]-current[,2])^2
                    +(future[,3]-current[,3])^2+(future[,4]-current[,4])^2
                    +(future[,5]-current[,5])^2+(future[,6]-current[,6])^2)
  
}


# check that it's working and gives same results as above

off <- calc_offset(scaled_gf_clim$historic[pop_cells,], scaled_gf_clim$MIROC_4.5_2050[pop_cells,])
boxplot(off ~ pop_islands, las = 2)
```

```
# predict the offset for each 'population' (cell with trees present) if planted in a different site
# each column is a 'population'
# each row is a possible planting site
# offset is if a population were planted into a given site under its future climate
# for the rows, we include all cells on the islands as possible planting sites
# columns are just places we collected trees

# set up cleaner column names
colnames_off <- paste(gsub(' ', '_', pop_islands), pop_cells, sep = '_')
colnames_off <- gsub('Island_', '', colnames_off)

off.agf <- list()

# loop through our 4 climate models and calculate offset values
# this may take a few minutes to run

for(n in 1:length(futures)){
  
  # set up empty dataframe to save offsets
  off <- as.data.frame(matrix(nrow = nrow(scaled_gf_clim$historic), ncol = length(pop_cells)))
  colnames(off) <- colnames_off
  # add island names to cell names
  # first make sure names match
  sum(rownames(scaled_gf_clim$historic) != rownames(env_trns))
  rownames(off) <- paste(env_trns$island, '_', env_trns$cell, sep = '')
  
  
  # calculate offsets for each population and planting site
  
  # first loop through current sites
  for(p in 1:length(pop_cells)){
    
    # get the historic climate variables for population
    pop <- scaled_gf_clim$historic[pop_cells[p], vars]
    
    # for each site, calculate offset to other locations in the future
    # by looping through rows of future climate
    for(f in 1:nrow(off)){
      
      fut <- scaled_gf_clim[[n]][rownames(env_trns)[f], vars]
      off[f,p] <- calc_offset(pop, fut)
    }
  }
  off.agf[[n]] <- off
  cat('done with', n, '\n')
}
```

```
## done with 1 
## done with 2 
## done with 3 
## done with 4
```

```
names(off.agf) <- names(futures)


# plot offsets as a heatmap
# columns are a population
# rows are offset for future climate at sites where it could be planted 
# san clemente is best for most pops because bio5 (max temp warmest month) will actually decrease in the future under this climate model

for(n in 1:length(off.agf)){
  
  #png(file = 'results/gradient_forest/offset_with_agf_heatmap_miroc85_2050.png', height = 8, width = 8, res = 300, units = 'in')
  heatmap(as.matrix(off.agf[[n]]), Rowv = NA, Colv = NA, margins = c(12, 10), scale = 'none', xlab = 'source site', ylab = 'planting site', main = paste('offset with AGF', names(off.agf)[n]))
  #dev.off()
  
}
```

```
#######################################################

# Compare offsets under different management scenarios

# SUMMARY PLOT
# format data to compare across three options: 
# no movement, movement to best site, and planting from best site
# this calculates the lowest offset (best scenario) for a parciular 'population' (cell) under each scenario

# check order
cbind(colnames(off.agf$MIROC_4.5_2050), paste(pop_islands, pop_cells))
```

```
##       [,1]                 [,2]                       
##  [1,] "Santa_Rosa_4773"    "Santa Rosa Island 4773"   
##  [2,] "Santa_Rosa_4774"    "Santa Rosa Island 4774"   
##  [3,] "Santa_Rosa_4775"    "Santa Rosa Island 4775"   
##  [4,] "Santa_Rosa_4511"    "Santa Rosa Island 4511"   
##  [5,] "Santa_Rosa_3723"    "Santa Rosa Island 3723"   
##  [6,] "Santa_Rosa_3722"    "Santa Rosa Island 3722"   
##  [7,] "Santa_Rosa_3724"    "Santa Rosa Island 3724"   
##  [8,] "Santa_Rosa_3460"    "Santa Rosa Island 3460"   
##  [9,] "Santa_Cruz_2174"    "Santa Cruz Island 2174"   
## [10,] "Santa_Cruz_2177"    "Santa Cruz Island 2177"   
## [11,] "Santa_Cruz_1911"    "Santa Cruz Island 1911"   
## [12,] "Santa_Cruz_2700"    "Santa Cruz Island 2700"   
## [13,] "Santa_Cruz_2725"    "Santa Cruz Island 2725"   
## [14,] "Santa_Cruz_2702"    "Santa Cruz Island 2702"   
## [15,] "Santa_Cruz_2701"    "Santa Cruz Island 2701"   
## [16,] "Santa_Cruz_2699"    "Santa Cruz Island 2699"   
## [17,] "Santa_Cruz_2185"    "Santa Cruz Island 2185"   
## [18,] "Santa_Cruz_1912"    "Santa Cruz Island 1912"   
## [19,] "Anacapa_2745"       "Anacapa Island 2745"      
## [20,] "Catalina_23193"     "Catalina Island 23193"    
## [21,] "Catalina_21878"     "Catalina Island 21878"    
## [22,] "Catalina_23465"     "Catalina Island 23465"    
## [23,] "Catalina_22929"     "Catalina Island 22929"    
## [24,] "San_Clemente_37707" "San Clemente Island 37707"
## [25,] "San_Clemente_39294" "San Clemente Island 39294"
## [26,] "San_Clemente_38768" "San Clemente Island 38768"
## [27,] "San_Clemente_38769" "San Clemente Island 38769"
## [28,] "San_Clemente_37442" "San Clemente Island 37442"
## [29,] "San_Clemente_37972" "San Clemente Island 37972"
## [30,] "San_Clemente_38238" "San Clemente Island 38238"
## [31,] "San_Clemente_38767" "San Clemente Island 38767"
## [32,] "San_Clemente_38503" "San Clemente Island 38503"
## [33,] "San_Clemente_38502" "San Clemente Island 38502"
## [34,] "San_Clemente_39034" "San Clemente Island 39034"
```

```
# setup list to save results for each future climate
offset.compare <- list()
offset.best <- list()

# look through future climate, get the best planting site or source site for each population we collected
for(fut in 1:length(off.agf)){
  
  off <- off.agf[[fut]]
  
  
  # set up df for saving the offset values
  off.compare <- as.data.frame(matrix(nrow = ncol(off), ncol = 5))
  rownames(off.compare) <- colnames(off)
  colnames(off.compare) <- c('island', 'no_movement', 'from_best_pop', 'to_best_site', 'to_best_current_site')
  off.compare$island <- pop_islands
  
  # set up similar df for saving the cell IDs of the best site for each scenario
  off.best <- as.data.frame(matrix(nrow = ncol(off), ncol = 4))
  rownames(off.best) <- colnames(off)
  colnames(off.best) <- c('island', 'best_source_pop', 'best_planting_site', 'best_planting_site_current_pop')
  off.best$island <- pop_islands
  
  for(n in 1:nrow(off.compare)){
    
    # in off.agf, columns are populations and are named with both island and cell number
    # rows are the same, but include all cells on the islands
    site <- rownames(off.compare)[n]
    # the cell is the last number of the string
    #site <- rev(strsplit(site, '_')[[1]])[1]

    
    # no movement - what is offset from the same site
    off.compare[n, 'no_movement'] <- off[site,site]
    
    # to best site - lowest offset in species range for this population
    # in off.agf, columns are source pops, rows are planting site
    # get the minimum for this column/population
    off.compare[n, 'to_best_site'] <- min(off[,site])
    # which cell is this?
    off.best[n, 'best_planting_site'] <- rownames(off)[which.min(off[,site])]
    # which cell is best, including only sites where there are currently trees that we sampled?
    # colnames(off) here is grabbing the cells where we collected by name
    off.compare[n, 'to_best_current_site'] <- min(off[colnames(off),site])
    off.best[n, 'best_planting_site_current_pop'] <- rownames(off[colnames(off),])[which.min(off[colnames(off),site])]
    
    
    # from best site - plant another population at this site
    # get the minimum from this row/planting site
    off.compare[n, 'from_best_pop'] <- min(off[site,])
    # which cell is this?
    off.best[n, 'best_source_pop'] <- colnames(off)[which.min(off[site,])]
    
  }
  
  offset.compare[[fut]] <- off.compare
  offset.best[[fut]] <- off.best
  
}


names(offset.compare) <- names(off.agf)
names(offset.best) <- names(off.agf)


# PLOTS

off.compare.long <- list() # for saving results

for(n in 1:length(offset.compare)){
  
  off.compare <- offset.compare[[n]]
  
  # first convert to long format for plotting a boxplot
  off.compare$id <- rownames(off.compare)
  
  off.compare.long[[n]] <-reshape(off.compare, direction = 'long', idvar = 'id', varying = c("no_movement", "to_best_current_site", "to_best_site", "from_best_pop"), times = c("no_movement", "to_best_current_site", "to_best_site", "from_best_pop"), timevar = 'strategy',  v.names = 'offset')
  
  off.compare.long[[n]]$strategy <- factor(off.compare.long[[n]]$strategy, levels = c("no_movement", "from_best_pop", "to_best_current_site", "to_best_site"))
  
  
  #png(file = 'results/gradient_forest/offset_comparison_3strategies_byIsland.png', height = 10, width = 10, res = 300, units = 'in')
  par(mfrow = c(1,1), mar = c(16,5,4,3))
  boxplot(off.compare.long[[n]]$offset ~ off.compare.long[[n]]$strategy * off.compare.long[[n]]$island,
          main = names(offset.compare)[n],
          names = rep(c('Status quo',  'Ecosystem preservation', 'Species preservation (sampled sites)', 'Species preservation (any site)'), 5),
          las = 2, 
          cex.lab = 1.5,
          cex.main = 1.5,
          cex.axis = 1.2,
          xlab = '',
          ylab = 'Offset', 
          drop = T, 
          col = c('grey40', 'grey80', 'grey90', 'white'), 
          border = rep(c("#3c4a8b", "#009c85", "#84bc5f", "#edb829", "#f57404"), each = 4))
  legend('topright', fill = c('grey40', 'grey80', 'grey90', 'white'), legend = c('Status quo',  'Ecosystem preservation', 'Species preservation (sampled sites)', 'Species preservation (any site)'), cex = 0.7)
  #dev.off()
  
}
```

```
names(off.compare.long) <- names(offset.compare)


# make nice tables with best planting/source site
# note: lat/lons are for the center of the cell, not the exact site of trees

for(n in 1:length(offset.best)){
  
  off.best <- offset.best[[n]]
  
  # add lat/lons for cells
  
  # planting site
  # get just the cell number without the island name to match to 'cells' object
  # sorry about the readability 
  # this just splits the string by underscore, then gets the last one using rev()
  best_cell <- sapply(1:nrow(off.best), function(x) rev(strsplit(off.best$best_planting_site[x], '_')[[1]])[1])
  off.best$best_planting_site_lat <- cells[best_cell, c('y')]
  off.best$best_planting_site_lon <- cells[best_cell, c('x')]
  
  # planting site, current pop
  # get just the cell number without the island name to match to 'cells' object
   best_cell <- sapply(1:nrow(off.best), function(x) rev(strsplit(off.best$best_planting_site_current_pop[x], '_')[[1]])[1])
  off.best$best_planting_site_current_lat <- cells[best_cell, c('y')]
  off.best$best_planting_site_current_lon <- cells[best_cell, c('x')]
  
  # source pop
  # get just the cell number without the island name to match to 'cells' object
  best_cell <- sapply(1:nrow(off.best), function(x) rev(strsplit(off.best$best_source_pop[x], '_')[[1]])[1])
  off.best$best_source_pop_lat <- cells[best_cell, c('y')]
  off.best$best_source_pop_lon <- cells[best_cell, c('x')]
  
  # rearrange columns
off.best <- off.best[,c("island", "best_source_pop", "best_source_pop_lat", "best_source_pop_lon", "best_planting_site", "best_planting_site_lat", "best_planting_site_lon", "best_planting_site_current_pop", "best_planting_site_current_lat", "best_planting_site_current_lon")]
  
  offset.best[[n]] <- off.best
}


knitr::kable(offset.best[[1]], caption = names(offset.best)[1])
```

MIROC\_4.5\_2050

|  | island | best\_source\_pop | best\_source\_pop\_lat | best\_source\_pop\_lon | best\_planting\_site | best\_planting\_site\_lat | best\_planting\_site\_lon | best\_planting\_site\_current\_pop | best\_planting\_site\_current\_lat | best\_planting\_site\_current\_lon |
| --- | --- | --- | --- | --- | --- | --- | --- | --- | --- | --- |
| Santa\_Rosa\_4773 | Santa Rosa Island | Catalina\_23465 | 33.3625 | -118.3625 | Santa\_Rosa\_5566 | 33.92083 | -120.1208 | Santa\_Rosa\_4773 | 33.94583 | -120.1292 |
| Santa\_Rosa\_4774 | Santa Rosa Island | Catalina\_23465 | 33.3625 | -118.3625 | Santa\_Rosa\_5566 | 33.92083 | -120.1208 | Santa\_Rosa\_4773 | 33.94583 | -120.1292 |
| Santa\_Rosa\_4775 | Santa Rosa Island | Catalina\_23465 | 33.3625 | -118.3625 | Santa\_Rosa\_5566 | 33.92083 | -120.1208 | Santa\_Rosa\_4773 | 33.94583 | -120.1292 |
| Santa\_Rosa\_4511 | Santa Rosa Island | Catalina\_23465 | 33.3625 | -118.3625 | Santa\_Rosa\_5566 | 33.92083 | -120.1208 | Santa\_Rosa\_4773 | 33.94583 | -120.1292 |
| Santa\_Rosa\_3723 | Santa Rosa Island | Catalina\_23465 | 33.3625 | -118.3625 | Santa\_Rosa\_5566 | 33.92083 | -120.1208 | Santa\_Rosa\_4773 | 33.94583 | -120.1292 |
| Santa\_Rosa\_3722 | Santa Rosa Island | Catalina\_23465 | 33.3625 | -118.3625 | Santa\_Rosa\_5566 | 33.92083 | -120.1208 | Santa\_Rosa\_4773 | 33.94583 | -120.1292 |
| Santa\_Rosa\_3724 | Santa Rosa Island | Catalina\_23465 | 33.3625 | -118.3625 | Santa\_Rosa\_5566 | 33.92083 | -120.1208 | Santa\_Rosa\_4773 | 33.94583 | -120.1292 |
| Santa\_Rosa\_3460 | Santa Rosa Island | Catalina\_23465 | 33.3625 | -118.3625 | Santa\_Rosa\_5566 | 33.92083 | -120.1208 | Santa\_Rosa\_4773 | 33.94583 | -120.1292 |
| Santa\_Cruz\_2174 | Santa Cruz Island | Catalina\_23465 | 33.3625 | -118.3625 | Santa\_Rosa\_5566 | 33.92083 | -120.1208 | Santa\_Rosa\_4773 | 33.94583 | -120.1292 |
| Santa\_Cruz\_2177 | Santa Cruz Island | Catalina\_23465 | 33.3625 | -118.3625 | Santa\_Rosa\_5566 | 33.92083 | -120.1208 | Santa\_Rosa\_4773 | 33.94583 | -120.1292 |
| Santa\_Cruz\_1911 | Santa Cruz Island | Catalina\_23465 | 33.3625 | -118.3625 | Santa\_Rosa\_5566 | 33.92083 | -120.1208 | Santa\_Rosa\_4773 | 33.94583 | -120.1292 |
| Santa\_Cruz\_2700 | Santa Cruz Island | Catalina\_23465 | 33.3625 | -118.3625 | Santa\_Rosa\_5566 | 33.92083 | -120.1208 | Santa\_Rosa\_4773 | 33.94583 | -120.1292 |
| Santa\_Cruz\_2725 | Santa Cruz Island | Catalina\_23465 | 33.3625 | -118.3625 | Santa\_Rosa\_5566 | 33.92083 | -120.1208 | Santa\_Rosa\_4773 | 33.94583 | -120.1292 |
| Santa\_Cruz\_2702 | Santa Cruz Island | Catalina\_23465 | 33.3625 | -118.3625 | Santa\_Rosa\_5566 | 33.92083 | -120.1208 | Santa\_Rosa\_4773 | 33.94583 | -120.1292 |
| Santa\_Cruz\_2701 | Santa Cruz Island | Catalina\_23465 | 33.3625 | -118.3625 | Santa\_Rosa\_5566 | 33.92083 | -120.1208 | Santa\_Rosa\_4773 | 33.94583 | -120.1292 |
| Santa\_Cruz\_2699 | Santa Cruz Island | Catalina\_23465 | 33.3625 | -118.3625 | Santa\_Rosa\_5566 | 33.92083 | -120.1208 | Santa\_Rosa\_4773 | 33.94583 | -120.1292 |
| Santa\_Cruz\_2185 | Santa Cruz Island | Catalina\_23465 | 33.3625 | -118.3625 | Santa\_Rosa\_5566 | 33.92083 | -120.1208 | Santa\_Rosa\_4773 | 33.94583 | -120.1292 |
| Santa\_Cruz\_1912 | Santa Cruz Island | Catalina\_23465 | 33.3625 | -118.3625 | Santa\_Rosa\_5566 | 33.92083 | -120.1208 | Santa\_Rosa\_4773 | 33.94583 | -120.1292 |
| Anacapa\_2745 | Anacapa Island | Catalina\_23465 | 33.3625 | -118.3625 | Santa\_Rosa\_5566 | 33.92083 | -120.1208 | Santa\_Rosa\_4773 | 33.94583 | -120.1292 |
| Catalina\_23193 | Catalina Island | Catalina\_23465 | 33.3625 | -118.3625 | Santa\_Rosa\_5566 | 33.92083 | -120.1208 | Santa\_Rosa\_4773 | 33.94583 | -120.1292 |
| Catalina\_21878 | Catalina Island | Catalina\_23465 | 33.3625 | -118.3625 | Santa\_Rosa\_5566 | 33.92083 | -120.1208 | Santa\_Rosa\_4773 | 33.94583 | -120.1292 |
| Catalina\_23465 | Catalina Island | Catalina\_23465 | 33.3625 | -118.3625 | Santa\_Rosa\_5566 | 33.92083 | -120.1208 | Santa\_Rosa\_4773 | 33.94583 | -120.1292 |
| Catalina\_22929 | Catalina Island | Catalina\_23465 | 33.3625 | -118.3625 | Santa\_Rosa\_5566 | 33.92083 | -120.1208 | Santa\_Rosa\_4773 | 33.94583 | -120.1292 |
| San\_Clemente\_37707 | San Clemente Island | Catalina\_23465 | 33.3625 | -118.3625 | Santa\_Rosa\_5566 | 33.92083 | -120.1208 | Santa\_Rosa\_4773 | 33.94583 | -120.1292 |
| San\_Clemente\_39294 | San Clemente Island | Catalina\_23465 | 33.3625 | -118.3625 | Santa\_Rosa\_5566 | 33.92083 | -120.1208 | Santa\_Rosa\_4773 | 33.94583 | -120.1292 |
| San\_Clemente\_38768 | San Clemente Island | Catalina\_23465 | 33.3625 | -118.3625 | Santa\_Rosa\_5566 | 33.92083 | -120.1208 | Santa\_Rosa\_4773 | 33.94583 | -120.1292 |
| San\_Clemente\_38769 | San Clemente Island | Catalina\_23465 | 33.3625 | -118.3625 | Santa\_Rosa\_5566 | 33.92083 | -120.1208 | Santa\_Rosa\_4773 | 33.94583 | -120.1292 |
| San\_Clemente\_37442 | San Clemente Island | Catalina\_23465 | 33.3625 | -118.3625 | Santa\_Rosa\_5566 | 33.92083 | -120.1208 | Santa\_Rosa\_4773 | 33.94583 | -120.1292 |
| San\_Clemente\_37972 | San Clemente Island | Catalina\_23465 | 33.3625 | -118.3625 | Santa\_Rosa\_5566 | 33.92083 | -120.1208 | Santa\_Rosa\_4773 | 33.94583 | -120.1292 |
| San\_Clemente\_38238 | San Clemente Island | Catalina\_23465 | 33.3625 | -118.3625 | Santa\_Rosa\_5566 | 33.92083 | -120.1208 | Santa\_Rosa\_4773 | 33.94583 | -120.1292 |
| San\_Clemente\_38767 | San Clemente Island | Catalina\_23465 | 33.3625 | -118.3625 | Santa\_Rosa\_5566 | 33.92083 | -120.1208 | Santa\_Rosa\_4773 | 33.94583 | -120.1292 |
| San\_Clemente\_38503 | San Clemente Island | Catalina\_23465 | 33.3625 | -118.3625 | Santa\_Rosa\_5566 | 33.92083 | -120.1208 | Santa\_Rosa\_4773 | 33.94583 | -120.1292 |
| San\_Clemente\_38502 | San Clemente Island | Catalina\_23465 | 33.3625 | -118.3625 | Santa\_Rosa\_5566 | 33.92083 | -120.1208 | Santa\_Rosa\_4773 | 33.94583 | -120.1292 |
| San\_Clemente\_39034 | San Clemente Island | Catalina\_23465 | 33.3625 | -118.3625 | Santa\_Rosa\_5566 | 33.92083 | -120.1208 | Santa\_Rosa\_4773 | 33.94583 | -120.1292 |

```
knitr::kable(offset.best[[2]], caption = names(offset.best)[2])
```

MIROC\_8.5\_2050

|  | island | best\_source\_pop | best\_source\_pop\_lat | best\_source\_pop\_lon | best\_planting\_site | best\_planting\_site\_lat | best\_planting\_site\_lon | best\_planting\_site\_current\_pop | best\_planting\_site\_current\_lat | best\_planting\_site\_current\_lon |
| --- | --- | --- | --- | --- | --- | --- | --- | --- | --- | --- |
| Santa\_Rosa\_4773 | Santa Rosa Island | Catalina\_23465 | 33.3625 | -118.3625 | Santa\_Rosa\_5035 | 33.93750 | -120.1458 | Santa\_Rosa\_4773 | 33.94583 | -120.1292 |
| Santa\_Rosa\_4774 | Santa Rosa Island | Catalina\_23465 | 33.3625 | -118.3625 | Santa\_Rosa\_5035 | 33.93750 | -120.1458 | Santa\_Rosa\_4773 | 33.94583 | -120.1292 |
| Santa\_Rosa\_4775 | Santa Rosa Island | Catalina\_23465 | 33.3625 | -118.3625 | Santa\_Rosa\_5035 | 33.93750 | -120.1458 | Santa\_Rosa\_4773 | 33.94583 | -120.1292 |
| Santa\_Rosa\_4511 | Santa Rosa Island | Catalina\_23465 | 33.3625 | -118.3625 | Santa\_Rosa\_5035 | 33.93750 | -120.1458 | Santa\_Rosa\_4773 | 33.94583 | -120.1292 |
| Santa\_Rosa\_3723 | Santa Rosa Island | Catalina\_23465 | 33.3625 | -118.3625 | Santa\_Rosa\_5035 | 33.93750 | -120.1458 | Santa\_Rosa\_4773 | 33.94583 | -120.1292 |
| Santa\_Rosa\_3722 | Santa Rosa Island | Catalina\_23465 | 33.3625 | -118.3625 | Santa\_Rosa\_4505 | 33.95417 | -120.1625 | Santa\_Rosa\_4773 | 33.94583 | -120.1292 |
| Santa\_Rosa\_3724 | Santa Rosa Island | Catalina\_23465 | 33.3625 | -118.3625 | Santa\_Rosa\_5035 | 33.93750 | -120.1458 | Santa\_Rosa\_4773 | 33.94583 | -120.1292 |
| Santa\_Rosa\_3460 | Santa Rosa Island | Catalina\_23465 | 33.3625 | -118.3625 | Santa\_Rosa\_4505 | 33.95417 | -120.1625 | Santa\_Rosa\_4773 | 33.94583 | -120.1292 |
| Santa\_Cruz\_2174 | Santa Cruz Island | Catalina\_23465 | 33.3625 | -118.3625 | Santa\_Rosa\_5035 | 33.93750 | -120.1458 | Santa\_Rosa\_4773 | 33.94583 | -120.1292 |
| Santa\_Cruz\_2177 | Santa Cruz Island | Catalina\_23465 | 33.3625 | -118.3625 | Santa\_Rosa\_5035 | 33.93750 | -120.1458 | Santa\_Rosa\_4773 | 33.94583 | -120.1292 |
| Santa\_Cruz\_1911 | Santa Cruz Island | Catalina\_23465 | 33.3625 | -118.3625 | Santa\_Rosa\_5035 | 33.93750 | -120.1458 | Santa\_Rosa\_4773 | 33.94583 | -120.1292 |
| Santa\_Cruz\_2700 | Santa Cruz Island | Catalina\_23465 | 33.3625 | -118.3625 | Santa\_Rosa\_5035 | 33.93750 | -120.1458 | Santa\_Rosa\_4773 | 33.94583 | -120.1292 |
| Santa\_Cruz\_2725 | Santa Cruz Island | Catalina\_23465 | 33.3625 | -118.3625 | Santa\_Rosa\_5035 | 33.93750 | -120.1458 | Santa\_Rosa\_4773 | 33.94583 | -120.1292 |
| Santa\_Cruz\_2702 | Santa Cruz Island | Catalina\_23465 | 33.3625 | -118.3625 | Santa\_Rosa\_5035 | 33.93750 | -120.1458 | Santa\_Rosa\_4773 | 33.94583 | -120.1292 |
| Santa\_Cruz\_2701 | Santa Cruz Island | Catalina\_23465 | 33.3625 | -118.3625 | Santa\_Rosa\_5035 | 33.93750 | -120.1458 | Santa\_Rosa\_4773 | 33.94583 | -120.1292 |
| Santa\_Cruz\_2699 | Santa Cruz Island | Catalina\_23465 | 33.3625 | -118.3625 | Santa\_Rosa\_5035 | 33.93750 | -120.1458 | Santa\_Rosa\_4773 | 33.94583 | -120.1292 |
| Santa\_Cruz\_2185 | Santa Cruz Island | Catalina\_23465 | 33.3625 | -118.3625 | Santa\_Rosa\_4505 | 33.95417 | -120.1625 | Santa\_Rosa\_4773 | 33.94583 | -120.1292 |
| Santa\_Cruz\_1912 | Santa Cruz Island | Catalina\_23465 | 33.3625 | -118.3625 | Santa\_Rosa\_5035 | 33.93750 | -120.1458 | Santa\_Rosa\_4773 | 33.94583 | -120.1292 |
| Anacapa\_2745 | Anacapa Island | Catalina\_23465 | 33.3625 | -118.3625 | Santa\_Rosa\_4505 | 33.95417 | -120.1625 | Santa\_Rosa\_4773 | 33.94583 | -120.1292 |
| Catalina\_23193 | Catalina Island | Catalina\_23465 | 33.3625 | -118.3625 | Santa\_Rosa\_5035 | 33.93750 | -120.1458 | Santa\_Rosa\_4773 | 33.94583 | -120.1292 |
| Catalina\_21878 | Catalina Island | Catalina\_23465 | 33.3625 | -118.3625 | Santa\_Rosa\_4505 | 33.95417 | -120.1625 | Santa\_Rosa\_4773 | 33.94583 | -120.1292 |
| Catalina\_23465 | Catalina Island | Catalina\_23465 | 33.3625 | -118.3625 | Santa\_Rosa\_5035 | 33.93750 | -120.1458 | Santa\_Rosa\_4773 | 33.94583 | -120.1292 |
| Catalina\_22929 | Catalina Island | Catalina\_23465 | 33.3625 | -118.3625 | Santa\_Rosa\_5035 | 33.93750 | -120.1458 | Santa\_Rosa\_4773 | 33.94583 | -120.1292 |
| San\_Clemente\_37707 | San Clemente Island | Catalina\_23465 | 33.3625 | -118.3625 | Santa\_Rosa\_5035 | 33.93750 | -120.1458 | Santa\_Rosa\_4773 | 33.94583 | -120.1292 |
| San\_Clemente\_39294 | San Clemente Island | Catalina\_23465 | 33.3625 | -118.3625 | Santa\_Rosa\_5035 | 33.93750 | -120.1458 | Santa\_Rosa\_4773 | 33.94583 | -120.1292 |
| San\_Clemente\_38768 | San Clemente Island | Catalina\_23465 | 33.3625 | -118.3625 | Santa\_Rosa\_5035 | 33.93750 | -120.1458 | Santa\_Rosa\_4773 | 33.94583 | -120.1292 |
| San\_Clemente\_38769 | San Clemente Island | Catalina\_23465 | 33.3625 | -118.3625 | Santa\_Rosa\_4505 | 33.95417 | -120.1625 | Santa\_Rosa\_4773 | 33.94583 | -120.1292 |
| San\_Clemente\_37442 | San Clemente Island | Catalina\_23465 | 33.3625 | -118.3625 | Santa\_Rosa\_4505 | 33.95417 | -120.1625 | Santa\_Rosa\_4773 | 33.94583 | -120.1292 |
| San\_Clemente\_37972 | San Clemente Island | Catalina\_23465 | 33.3625 | -118.3625 | Santa\_Rosa\_5035 | 33.93750 | -120.1458 | Santa\_Rosa\_4773 | 33.94583 | -120.1292 |
| San\_Clemente\_38238 | San Clemente Island | Catalina\_23465 | 33.3625 | -118.3625 | Santa\_Rosa\_4505 | 33.95417 | -120.1625 | Santa\_Rosa\_4773 | 33.94583 | -120.1292 |
| San\_Clemente\_38767 | San Clemente Island | Catalina\_23465 | 33.3625 | -118.3625 | Santa\_Rosa\_5035 | 33.93750 | -120.1458 | Santa\_Rosa\_4773 | 33.94583 | -120.1292 |
| San\_Clemente\_38503 | San Clemente Island | Catalina\_23465 | 33.3625 | -118.3625 | Santa\_Rosa\_5035 | 33.93750 | -120.1458 | Santa\_Rosa\_4773 | 33.94583 | -120.1292 |
| San\_Clemente\_38502 | San Clemente Island | Catalina\_23465 | 33.3625 | -118.3625 | Santa\_Rosa\_5035 | 33.93750 | -120.1458 | Santa\_Rosa\_4773 | 33.94583 | -120.1292 |
| San\_Clemente\_39034 | San Clemente Island | Catalina\_23465 | 33.3625 | -118.3625 | Santa\_Rosa\_4505 | 33.95417 | -120.1625 | Santa\_Rosa\_4773 | 33.94583 | -120.1292 |

```
knitr::kable(offset.best[[3]], caption = names(offset.best)[3])
```

CNRM\_4.5\_2050

|  | island | best\_source\_pop | best\_source\_pop\_lat | best\_source\_pop\_lon | best\_planting\_site | best\_planting\_site\_lat | best\_planting\_site\_lon | best\_planting\_site\_current\_pop | best\_planting\_site\_current\_lat | best\_planting\_site\_current\_lon |
| --- | --- | --- | --- | --- | --- | --- | --- | --- | --- | --- |
| Santa\_Rosa\_4773 | Santa Rosa Island | Catalina\_23465 | 33.3625 | -118.3625 | Santa\_Rosa\_5566 | 33.92083 | -120.1208 | Santa\_Rosa\_4773 | 33.94583 | -120.1292 |
| Santa\_Rosa\_4774 | Santa Rosa Island | Catalina\_23465 | 33.3625 | -118.3625 | Santa\_Rosa\_5566 | 33.92083 | -120.1208 | Santa\_Rosa\_4773 | 33.94583 | -120.1292 |
| Santa\_Rosa\_4775 | Santa Rosa Island | Catalina\_23465 | 33.3625 | -118.3625 | Santa\_Rosa\_5566 | 33.92083 | -120.1208 | Santa\_Rosa\_4773 | 33.94583 | -120.1292 |
| Santa\_Rosa\_4511 | Santa Rosa Island | Catalina\_23465 | 33.3625 | -118.3625 | Santa\_Rosa\_5566 | 33.92083 | -120.1208 | Santa\_Rosa\_4773 | 33.94583 | -120.1292 |
| Santa\_Rosa\_3723 | Santa Rosa Island | Catalina\_23465 | 33.3625 | -118.3625 | Santa\_Rosa\_5566 | 33.92083 | -120.1208 | Santa\_Rosa\_4773 | 33.94583 | -120.1292 |
| Santa\_Rosa\_3722 | Santa Rosa Island | Catalina\_23465 | 33.3625 | -118.3625 | Santa\_Rosa\_5566 | 33.92083 | -120.1208 | Santa\_Rosa\_4773 | 33.94583 | -120.1292 |
| Santa\_Rosa\_3724 | Santa Rosa Island | Catalina\_23465 | 33.3625 | -118.3625 | Santa\_Rosa\_5566 | 33.92083 | -120.1208 | Santa\_Rosa\_4773 | 33.94583 | -120.1292 |
| Santa\_Rosa\_3460 | Santa Rosa Island | Catalina\_23465 | 33.3625 | -118.3625 | Santa\_Rosa\_5566 | 33.92083 | -120.1208 | Santa\_Rosa\_4773 | 33.94583 | -120.1292 |
| Santa\_Cruz\_2174 | Santa Cruz Island | Catalina\_23465 | 33.3625 | -118.3625 | Santa\_Rosa\_5566 | 33.92083 | -120.1208 | Santa\_Rosa\_4773 | 33.94583 | -120.1292 |
| Santa\_Cruz\_2177 | Santa Cruz Island | Catalina\_23465 | 33.3625 | -118.3625 | Santa\_Rosa\_5566 | 33.92083 | -120.1208 | Santa\_Rosa\_4773 | 33.94583 | -120.1292 |
| Santa\_Cruz\_1911 | Santa Cruz Island | Catalina\_23465 | 33.3625 | -118.3625 | Santa\_Rosa\_5566 | 33.92083 | -120.1208 | Santa\_Rosa\_4773 | 33.94583 | -120.1292 |
| Santa\_Cruz\_2700 | Santa Cruz Island | Catalina\_23465 | 33.3625 | -118.3625 | Santa\_Rosa\_5566 | 33.92083 | -120.1208 | Santa\_Rosa\_4773 | 33.94583 | -120.1292 |
| Santa\_Cruz\_2725 | Santa Cruz Island | Catalina\_23465 | 33.3625 | -118.3625 | Santa\_Rosa\_5566 | 33.92083 | -120.1208 | Santa\_Rosa\_4773 | 33.94583 | -120.1292 |
| Santa\_Cruz\_2702 | Santa Cruz Island | Catalina\_23465 | 33.3625 | -118.3625 | Santa\_Rosa\_5566 | 33.92083 | -120.1208 | Santa\_Rosa\_4773 | 33.94583 | -120.1292 |
| Santa\_Cruz\_2701 | Santa Cruz Island | Catalina\_23465 | 33.3625 | -118.3625 | Santa\_Rosa\_5566 | 33.92083 | -120.1208 | Santa\_Rosa\_4773 | 33.94583 | -120.1292 |
| Santa\_Cruz\_2699 | Santa Cruz Island | Catalina\_23465 | 33.3625 | -118.3625 | Santa\_Rosa\_5566 | 33.92083 | -120.1208 | Santa\_Rosa\_4773 | 33.94583 | -120.1292 |
| Santa\_Cruz\_2185 | Santa Cruz Island | Catalina\_23465 | 33.3625 | -118.3625 | Santa\_Rosa\_5566 | 33.92083 | -120.1208 | Santa\_Rosa\_4773 | 33.94583 | -120.1292 |
| Santa\_Cruz\_1912 | Santa Cruz Island | Catalina\_23465 | 33.3625 | -118.3625 | Santa\_Rosa\_5566 | 33.92083 | -120.1208 | Santa\_Rosa\_4773 | 33.94583 | -120.1292 |
| Anacapa\_2745 | Anacapa Island | Catalina\_23465 | 33.3625 | -118.3625 | Santa\_Rosa\_5566 | 33.92083 | -120.1208 | Santa\_Rosa\_4773 | 33.94583 | -120.1292 |
| Catalina\_23193 | Catalina Island | Catalina\_23465 | 33.3625 | -118.3625 | Santa\_Rosa\_5566 | 33.92083 | -120.1208 | Santa\_Rosa\_4773 | 33.94583 | -120.1292 |
| Catalina\_21878 | Catalina Island | Catalina\_23465 | 33.3625 | -118.3625 | Santa\_Rosa\_5566 | 33.92083 | -120.1208 | Santa\_Rosa\_4773 | 33.94583 | -120.1292 |
| Catalina\_23465 | Catalina Island | Catalina\_23465 | 33.3625 | -118.3625 | Santa\_Rosa\_5566 | 33.92083 | -120.1208 | Santa\_Rosa\_4773 | 33.94583 | -120.1292 |
| Catalina\_22929 | Catalina Island | Catalina\_23465 | 33.3625 | -118.3625 | Santa\_Rosa\_5566 | 33.92083 | -120.1208 | Santa\_Rosa\_4773 | 33.94583 | -120.1292 |
| San\_Clemente\_37707 | San Clemente Island | Catalina\_23465 | 33.3625 | -118.3625 | Santa\_Rosa\_5566 | 33.92083 | -120.1208 | Santa\_Rosa\_4773 | 33.94583 | -120.1292 |
| San\_Clemente\_39294 | San Clemente Island | Catalina\_23465 | 33.3625 | -118.3625 | Santa\_Rosa\_5566 | 33.92083 | -120.1208 | Santa\_Rosa\_4773 | 33.94583 | -120.1292 |
| San\_Clemente\_38768 | San Clemente Island | Catalina\_23465 | 33.3625 | -118.3625 | Santa\_Rosa\_5566 | 33.92083 | -120.1208 | Santa\_Rosa\_4773 | 33.94583 | -120.1292 |
| San\_Clemente\_38769 | San Clemente Island | Catalina\_23465 | 33.3625 | -118.3625 | Santa\_Rosa\_5566 | 33.92083 | -120.1208 | Santa\_Rosa\_4773 | 33.94583 | -120.1292 |
| San\_Clemente\_37442 | San Clemente Island | Catalina\_23465 | 33.3625 | -118.3625 | Santa\_Rosa\_5566 | 33.92083 | -120.1208 | Santa\_Rosa\_4773 | 33.94583 | -120.1292 |
| San\_Clemente\_37972 | San Clemente Island | Catalina\_23465 | 33.3625 | -118.3625 | Santa\_Rosa\_5566 | 33.92083 | -120.1208 | Santa\_Rosa\_4773 | 33.94583 | -120.1292 |
| San\_Clemente\_38238 | San Clemente Island | Catalina\_23465 | 33.3625 | -118.3625 | Santa\_Rosa\_5566 | 33.92083 | -120.1208 | Santa\_Rosa\_4773 | 33.94583 | -120.1292 |
| San\_Clemente\_38767 | San Clemente Island | Catalina\_23465 | 33.3625 | -118.3625 | Santa\_Rosa\_5566 | 33.92083 | -120.1208 | Santa\_Rosa\_4773 | 33.94583 | -120.1292 |
| San\_Clemente\_38503 | San Clemente Island | Catalina\_23465 | 33.3625 | -118.3625 | Santa\_Rosa\_5566 | 33.92083 | -120.1208 | Santa\_Rosa\_4773 | 33.94583 | -120.1292 |
| San\_Clemente\_38502 | San Clemente Island | Catalina\_23465 | 33.3625 | -118.3625 | Santa\_Rosa\_5566 | 33.92083 | -120.1208 | Santa\_Rosa\_4773 | 33.94583 | -120.1292 |
| San\_Clemente\_39034 | San Clemente Island | Catalina\_23465 | 33.3625 | -118.3625 | Santa\_Rosa\_5566 | 33.92083 | -120.1208 | Santa\_Rosa\_4773 | 33.94583 | -120.1292 |

```
knitr::kable(offset.best[[4]], caption = names(offset.best)[4])
```

CNRM\_8.5\_2050

|  | island | best\_source\_pop | best\_source\_pop\_lat | best\_source\_pop\_lon | best\_planting\_site | best\_planting\_site\_lat | best\_planting\_site\_lon | best\_planting\_site\_current\_pop | best\_planting\_site\_current\_lat | best\_planting\_site\_current\_lon |
| --- | --- | --- | --- | --- | --- | --- | --- | --- | --- | --- |
| Santa\_Rosa\_4773 | Santa Rosa Island | Catalina\_23465 | 33.3625 | -118.3625 | San\_Clemente\_39296 | 32.8625 | -118.4375 | San\_Clemente\_39294 | 32.8625 | -118.4542 |
| Santa\_Rosa\_4774 | Santa Rosa Island | Catalina\_23465 | 33.3625 | -118.3625 | San\_Clemente\_39296 | 32.8625 | -118.4375 | San\_Clemente\_39294 | 32.8625 | -118.4542 |
| Santa\_Rosa\_4775 | Santa Rosa Island | Catalina\_23465 | 33.3625 | -118.3625 | San\_Clemente\_39296 | 32.8625 | -118.4375 | San\_Clemente\_39294 | 32.8625 | -118.4542 |
| Santa\_Rosa\_4511 | Santa Rosa Island | Catalina\_23465 | 33.3625 | -118.3625 | San\_Clemente\_39296 | 32.8625 | -118.4375 | San\_Clemente\_39294 | 32.8625 | -118.4542 |
| Santa\_Rosa\_3723 | Santa Rosa Island | Catalina\_23465 | 33.3625 | -118.3625 | San\_Clemente\_39296 | 32.8625 | -118.4375 | San\_Clemente\_39294 | 32.8625 | -118.4542 |
| Santa\_Rosa\_3722 | Santa Rosa Island | Catalina\_23465 | 33.3625 | -118.3625 | San\_Clemente\_39296 | 32.8625 | -118.4375 | San\_Clemente\_39294 | 32.8625 | -118.4542 |
| Santa\_Rosa\_3724 | Santa Rosa Island | Catalina\_23465 | 33.3625 | -118.3625 | San\_Clemente\_39296 | 32.8625 | -118.4375 | San\_Clemente\_39294 | 32.8625 | -118.4542 |
| Santa\_Rosa\_3460 | Santa Rosa Island | Catalina\_23465 | 33.3625 | -118.3625 | San\_Clemente\_39296 | 32.8625 | -118.4375 | San\_Clemente\_39294 | 32.8625 | -118.4542 |
| Santa\_Cruz\_2174 | Santa Cruz Island | Catalina\_23465 | 33.3625 | -118.3625 | San\_Clemente\_39296 | 32.8625 | -118.4375 | San\_Clemente\_39294 | 32.8625 | -118.4542 |
| Santa\_Cruz\_2177 | Santa Cruz Island | Catalina\_23465 | 33.3625 | -118.3625 | San\_Clemente\_39296 | 32.8625 | -118.4375 | San\_Clemente\_39294 | 32.8625 | -118.4542 |
| Santa\_Cruz\_1911 | Santa Cruz Island | Catalina\_23465 | 33.3625 | -118.3625 | San\_Clemente\_39296 | 32.8625 | -118.4375 | San\_Clemente\_39294 | 32.8625 | -118.4542 |
| Santa\_Cruz\_2700 | Santa Cruz Island | Catalina\_23465 | 33.3625 | -118.3625 | San\_Clemente\_39296 | 32.8625 | -118.4375 | San\_Clemente\_39294 | 32.8625 | -118.4542 |
| Santa\_Cruz\_2725 | Santa Cruz Island | Catalina\_23465 | 33.3625 | -118.3625 | San\_Clemente\_39296 | 32.8625 | -118.4375 | San\_Clemente\_39294 | 32.8625 | -118.4542 |
| Santa\_Cruz\_2702 | Santa Cruz Island | Catalina\_23465 | 33.3625 | -118.3625 | San\_Clemente\_39296 | 32.8625 | -118.4375 | San\_Clemente\_39294 | 32.8625 | -118.4542 |
| Santa\_Cruz\_2701 | Santa Cruz Island | Catalina\_23465 | 33.3625 | -118.3625 | San\_Clemente\_39296 | 32.8625 | -118.4375 | San\_Clemente\_39294 | 32.8625 | -118.4542 |
| Santa\_Cruz\_2699 | Santa Cruz Island | Catalina\_23465 | 33.3625 | -118.3625 | San\_Clemente\_39296 | 32.8625 | -118.4375 | San\_Clemente\_39294 | 32.8625 | -118.4542 |
| Santa\_Cruz\_2185 | Santa Cruz Island | Catalina\_23465 | 33.3625 | -118.3625 | San\_Clemente\_39296 | 32.8625 | -118.4375 | San\_Clemente\_39294 | 32.8625 | -118.4542 |
| Santa\_Cruz\_1912 | Santa Cruz Island | Catalina\_23465 | 33.3625 | -118.3625 | San\_Clemente\_39296 | 32.8625 | -118.4375 | San\_Clemente\_39294 | 32.8625 | -118.4542 |
| Anacapa\_2745 | Anacapa Island | Catalina\_23465 | 33.3625 | -118.3625 | San\_Clemente\_39296 | 32.8625 | -118.4375 | San\_Clemente\_39294 | 32.8625 | -118.4542 |
| Catalina\_23193 | Catalina Island | Catalina\_23465 | 33.3625 | -118.3625 | San\_Clemente\_39296 | 32.8625 | -118.4375 | San\_Clemente\_39294 | 32.8625 | -118.4542 |
| Catalina\_21878 | Catalina Island | Catalina\_23465 | 33.3625 | -118.3625 | San\_Clemente\_39296 | 32.8625 | -118.4375 | San\_Clemente\_39294 | 32.8625 | -118.4542 |
| Catalina\_23465 | Catalina Island | Catalina\_23465 | 33.3625 | -118.3625 | San\_Clemente\_39296 | 32.8625 | -118.4375 | San\_Clemente\_39294 | 32.8625 | -118.4542 |
| Catalina\_22929 | Catalina Island | Catalina\_23465 | 33.3625 | -118.3625 | San\_Clemente\_39296 | 32.8625 | -118.4375 | San\_Clemente\_39294 | 32.8625 | -118.4542 |
| San\_Clemente\_37707 | San Clemente Island | Catalina\_23465 | 33.3625 | -118.3625 | San\_Clemente\_39296 | 32.8625 | -118.4375 | San\_Clemente\_39294 | 32.8625 | -118.4542 |
| San\_Clemente\_39294 | San Clemente Island | Catalina\_23465 | 33.3625 | -118.3625 | San\_Clemente\_39296 | 32.8625 | -118.4375 | San\_Clemente\_39294 | 32.8625 | -118.4542 |
| San\_Clemente\_38768 | San Clemente Island | Catalina\_23465 | 33.3625 | -118.3625 | San\_Clemente\_39296 | 32.8625 | -118.4375 | San\_Clemente\_39294 | 32.8625 | -118.4542 |
| San\_Clemente\_38769 | San Clemente Island | Catalina\_23465 | 33.3625 | -118.3625 | San\_Clemente\_39296 | 32.8625 | -118.4375 | San\_Clemente\_39294 | 32.8625 | -118.4542 |
| San\_Clemente\_37442 | San Clemente Island | Catalina\_23465 | 33.3625 | -118.3625 | San\_Clemente\_39296 | 32.8625 | -118.4375 | San\_Clemente\_39294 | 32.8625 | -118.4542 |
| San\_Clemente\_37972 | San Clemente Island | Catalina\_23465 | 33.3625 | -118.3625 | San\_Clemente\_39296 | 32.8625 | -118.4375 | San\_Clemente\_39294 | 32.8625 | -118.4542 |
| San\_Clemente\_38238 | San Clemente Island | Catalina\_23465 | 33.3625 | -118.3625 | San\_Clemente\_39296 | 32.8625 | -118.4375 | San\_Clemente\_39294 | 32.8625 | -118.4542 |
| San\_Clemente\_38767 | San Clemente Island | Catalina\_23465 | 33.3625 | -118.3625 | San\_Clemente\_39296 | 32.8625 | -118.4375 | San\_Clemente\_39294 | 32.8625 | -118.4542 |
| San\_Clemente\_38503 | San Clemente Island | Catalina\_23465 | 33.3625 | -118.3625 | San\_Clemente\_39296 | 32.8625 | -118.4375 | San\_Clemente\_39294 | 32.8625 | -118.4542 |
| San\_Clemente\_38502 | San Clemente Island | Catalina\_23465 | 33.3625 | -118.3625 | San\_Clemente\_39296 | 32.8625 | -118.4375 | San\_Clemente\_39294 | 32.8625 | -118.4542 |
| San\_Clemente\_39034 | San Clemente Island | Catalina\_23465 | 33.3625 | -118.3625 | San\_Clemente\_39296 | 32.8625 | -118.4375 | San\_Clemente\_39294 | 32.8625 | -118.4542 |

```
# write.csv(off.pop.sort, file = 'results/gradient_forest/noGuadAna/offsets_by_indiv_no_AGF.csv', row.names = T)
# 
# write.csv(off.best, file = 'results/gradient_forest/noGuadAna/offsets_by_indiv_with_best_AGF.csv')
# save(offset.best, file = 'results/gradient_forest/noGuadAna/offsets_by_indiv_with_best_AGF.rda')
# save(off.compare.long, file = 'results/gradient_forest/noGuadAna/offsets_by_strategy_long_format_for_boxplots.rda')
```

### 6 Climate suitability

```
# calculate climate suitability index from genomic offset
# where 1 = same climate as home, 0 = maximum offset

range(unlist(off.agf))
```

```
## [1] 1.747200 2.270656
```

```
#[1] 1.744947 2.267777

# climate suitability index
csi <- off.agf

maxoff <- max(unlist(off.agf))

for(n in 1:length(off.agf)){
  
  csi[[n]] <- 1 - off.agf[[n]]/maxoff
  
}

hist(unlist(csi))
```

```
# quick heatmap to check
for(n in 1:length(csi)){
  
  heatmap(as.matrix(csi[[n]]), Rowv = NA, Colv = NA, margins = c(12, 10), scale = 'none', xlab = 'source site', ylab = 'planting site', main = paste('climate suitability', names(off.agf)[n]))

  
}
```

```
# another heatmap, subset 
# see below for a nicer heatmap plot
colf <- colorRampPalette(brewer.pal(9, 'YlGn'))

for(n in 1:length(csi)){
  
  mat <- as.matrix(csi[[n]][colnames(csi[[n]]),])
  # order by best planting site (overall)
  ord <- order(rowSums(mat))
  
 #png(file = paste('results/gradient_forest/climate_suitability_heatmap_noGuadAna_', names(csi)[n], '.png', sep = ''), height = 8, width = 8, res = 300, units = 'in')
  heatmap(mat[ord,], col = colf(100), Rowv = NA, Colv = NA, margins = c(10, 10), scale = 'none', xlab = 'source site', ylab = 'planting site', main = paste('climate suitability', names(off.agf)[n]))
  #dev.off()

  
}
```

```
# get island name of each row for aggregating values by island
# just use env_trns object, but first check that they match using a random subset
rand <- sample(1:nrow(env_trns), 30)
cbind(rownames(csi$MIROC_4.5_2050)[rand], env_trns$island[rand])
```

```
##       [,1]                 [,2]          
##  [1,] "Santa_Cruz_2439"    "Santa_Cruz"  
##  [2,] "San_Clemente_39559" "San_Clemente"
##  [3,] "Santa_Cruz_2182"    "Santa_Cruz"  
##  [4,] "Santa_Cruz_3762"    "Santa_Cruz"  
##  [5,] "San_Clemente_40360" "San_Clemente"
##  [6,] "Catalina_24248"     "Catalina"    
##  [7,] "Santa_Cruz_3754"    "Santa_Cruz"  
##  [8,] "Catalina_23199"     "Catalina"    
##  [9,] "San_Clemente_37436" "San_Clemente"
## [10,] "Santa_Rosa_3176"    "Santa_Rosa"  
## [11,] "Santa_Cruz_3508"    "Santa_Cruz"  
## [12,] "Santa_Cruz_1377"    "Santa_Cruz"  
## [13,] "Santa_Cruz_3239"    "Santa_Cruz"  
## [14,] "Santa_Cruz_2692"    "Santa_Cruz"  
## [15,] "Santa_Cruz_1409"    "Santa_Cruz"  
## [16,] "Catalina_22668"     "Catalina"    
## [17,] "Santa_Rosa_4519"    "Santa_Rosa"  
## [18,] "Santa_Cruz_3225"    "Santa_Cruz"  
## [19,] "Santa_Cruz_1939"    "Santa_Cruz"  
## [20,] "San_Clemente_36907" "San_Clemente"
## [21,] "San_Clemente_40616" "San_Clemente"
## [22,] "Santa_Cruz_2462"    "Santa_Cruz"  
## [23,] "Santa_Rosa_2140"    "Santa_Rosa"  
## [24,] "Santa_Rosa_6095"    "Santa_Rosa"  
## [25,] "Santa_Rosa_5296"    "Santa_Rosa"  
## [26,] "Santa_Rosa_4525"    "Santa_Rosa"  
## [27,] "Catalina_21338"     "Catalina"    
## [28,] "Catalina_22137"     "Catalina"    
## [29,] "San_Clemente_39822" "San_Clemente"
## [30,] "San_Clemente_39028" "San_Clemente"
```

```
island <- env_trns$island
island <- as.factor(island)
island <- factor(island, levels = c('Santa_Rosa', 'Santa_Cruz', 'Anacapa', 'Catalina', 'San_Clemente'))
# another check
cbind(rownames(csi$MIROC_4.5_2050)[rand], as.character(island[rand]))
```

```
##       [,1]                 [,2]          
##  [1,] "Santa_Cruz_2439"    "Santa_Cruz"  
##  [2,] "San_Clemente_39559" "San_Clemente"
##  [3,] "Santa_Cruz_2182"    "Santa_Cruz"  
##  [4,] "Santa_Cruz_3762"    "Santa_Cruz"  
##  [5,] "San_Clemente_40360" "San_Clemente"
##  [6,] "Catalina_24248"     "Catalina"    
##  [7,] "Santa_Cruz_3754"    "Santa_Cruz"  
##  [8,] "Catalina_23199"     "Catalina"    
##  [9,] "San_Clemente_37436" "San_Clemente"
## [10,] "Santa_Rosa_3176"    "Santa_Rosa"  
## [11,] "Santa_Cruz_3508"    "Santa_Cruz"  
## [12,] "Santa_Cruz_1377"    "Santa_Cruz"  
## [13,] "Santa_Cruz_3239"    "Santa_Cruz"  
## [14,] "Santa_Cruz_2692"    "Santa_Cruz"  
## [15,] "Santa_Cruz_1409"    "Santa_Cruz"  
## [16,] "Catalina_22668"     "Catalina"    
## [17,] "Santa_Rosa_4519"    "Santa_Rosa"  
## [18,] "Santa_Cruz_3225"    "Santa_Cruz"  
## [19,] "Santa_Cruz_1939"    "Santa_Cruz"  
## [20,] "San_Clemente_36907" "San_Clemente"
## [21,] "San_Clemente_40616" "San_Clemente"
## [22,] "Santa_Cruz_2462"    "Santa_Cruz"  
## [23,] "Santa_Rosa_2140"    "Santa_Rosa"  
## [24,] "Santa_Rosa_6095"    "Santa_Rosa"  
## [25,] "Santa_Rosa_5296"    "Santa_Rosa"  
## [26,] "Santa_Rosa_4525"    "Santa_Rosa"  
## [27,] "Catalina_21338"     "Catalina"    
## [28,] "Catalina_22137"     "Catalina"    
## [29,] "San_Clemente_39822" "San_Clemente"
## [30,] "San_Clemente_39028" "San_Clemente"
```

```
# what sites have the best future conditions for conserving the genetic composition of a given population?
# plot a boxplot of offset for planting sites in each population

# loop through each climate model
for(n in 1:length(csi)){
  
  # loop through each population
  # pdf(file = paste('results/gradient_forest/best_planting_site_boxplots_by_island_noGuadAna_', names(csi)[n], '.pdf', sep = ''),
  #     height = 12, width = 16)
  par(mar = c(8,5,5,2), mfrow = c(3,4))
  for(p in 1:ncol(csi[[n]])){
    
    boxplot(csi[[n]][,p] ~ island, 
            main = paste('Planting sites for\n' ,colnames(csi[[n]][p]), '\n', names(csi)[n], sep = ''),
            las = 2, 
            border = c("#3c4a8b", "#009c85", "#84bc5f", "#edb829", "#f57404"), 
            col = 'white', 
            xlab = '', 
            ylab = 'Climate Suitability Index',
            ylim = range(csi))
    
  }
  #dev.off()
  
}
```

```
# now do the same, but for seed sourcing
# which population is the best seed source for a site?
# subset to sites with trees

# get islands for column names
cbind(colnames(csi$MIROC_4.5_2050), as.character(pop_islands))
```

```
##       [,1]                 [,2]                 
##  [1,] "Santa_Rosa_4773"    "Santa Rosa Island"  
##  [2,] "Santa_Rosa_4774"    "Santa Rosa Island"  
##  [3,] "Santa_Rosa_4775"    "Santa Rosa Island"  
##  [4,] "Santa_Rosa_4511"    "Santa Rosa Island"  
##  [5,] "Santa_Rosa_3723"    "Santa Rosa Island"  
##  [6,] "Santa_Rosa_3722"    "Santa Rosa Island"  
##  [7,] "Santa_Rosa_3724"    "Santa Rosa Island"  
##  [8,] "Santa_Rosa_3460"    "Santa Rosa Island"  
##  [9,] "Santa_Cruz_2174"    "Santa Cruz Island"  
## [10,] "Santa_Cruz_2177"    "Santa Cruz Island"  
## [11,] "Santa_Cruz_1911"    "Santa Cruz Island"  
## [12,] "Santa_Cruz_2700"    "Santa Cruz Island"  
## [13,] "Santa_Cruz_2725"    "Santa Cruz Island"  
## [14,] "Santa_Cruz_2702"    "Santa Cruz Island"  
## [15,] "Santa_Cruz_2701"    "Santa Cruz Island"  
## [16,] "Santa_Cruz_2699"    "Santa Cruz Island"  
## [17,] "Santa_Cruz_2185"    "Santa Cruz Island"  
## [18,] "Santa_Cruz_1912"    "Santa Cruz Island"  
## [19,] "Anacapa_2745"       "Anacapa Island"     
## [20,] "Catalina_23193"     "Catalina Island"    
## [21,] "Catalina_21878"     "Catalina Island"    
## [22,] "Catalina_23465"     "Catalina Island"    
## [23,] "Catalina_22929"     "Catalina Island"    
## [24,] "San_Clemente_37707" "San Clemente Island"
## [25,] "San_Clemente_39294" "San Clemente Island"
## [26,] "San_Clemente_38768" "San Clemente Island"
## [27,] "San_Clemente_38769" "San Clemente Island"
## [28,] "San_Clemente_37442" "San Clemente Island"
## [29,] "San_Clemente_37972" "San Clemente Island"
## [30,] "San_Clemente_38238" "San Clemente Island"
## [31,] "San_Clemente_38767" "San Clemente Island"
## [32,] "San_Clemente_38503" "San Clemente Island"
## [33,] "San_Clemente_38502" "San Clemente Island"
## [34,] "San_Clemente_39034" "San Clemente Island"
```

```
island <- pop_islands
island <- gsub(' Island', '', island)
island <- gsub(' ', '_', island)
island <- as.factor(island)
island <- factor(island, levels = c('Santa_Rosa', 'Santa_Cruz', 'Anacapa', 'Catalina', 'San_Clemente'))

# loop through each climate model
for(n in 1:length(csi)){
  
  # loop through each population
  #pdf(file = paste('results/gradient_forest/best_seed_source_boxplots_by_island_noGuadAna_', names(csi)[n], '.pdf', sep = ''),
   #   height = 12, width = 16)
  par(mar = c(8,5,5,2), mfrow = c(3,4))
  for(p in 1:ncol(csi[[n]])){
    
    # use column names, which are actual pops, to get corresponding rows
    boxplot(as.numeric(csi[[n]][p,]) ~ island, 
            main = paste('Seed sources for\n' , colnames(csi[[n]][p]), '\n', names(csi)[n], sep = ''),
            las = 2, 
            border = c("#3c4a8b", "#009c85", "#84bc5f", "#edb829", "#f57404"), 
            col = 'white', 
            xlab = '', 
            ylab = 'Climate Suitability Index',
            ylim = range(csi))
    
  }
  #dev.off()
  
}
```

```
#################
# heatmap - merge similar/nearby cells
pop_cells
```

```
##  [1] "4773"  "4774"  "4775"  "4511"  "3723"  "3722"  "3724"  "3460"  "2174" 
## [10] "2177"  "1911"  "2700"  "2725"  "2702"  "2701"  "2699"  "2185"  "1912" 
## [19] "2745"  "23193" "21878" "23465" "22929" "37707" "39294" "38768" "38769"
## [28] "37442" "37972" "38238" "38767" "38503" "38502" "39034"
```

```
pop_cells_merge_nearby <- as.data.frame(matrix(ncol = 4, nrow = length(pop_cells)))
colnames(pop_cells_merge_nearby) <- c('cell', 'island', 'island_cell', 'group')
pop_cells_merge_nearby$cell <- pop_cells
pop_cells_merge_nearby$island <- pop_islands
pop_cells_merge_nearby$island_cell <- gsub(' ', '_', paste(pop_islands, pop_cells))

pop_cells_merge_nearby$island_cell <- gsub('_Island', '', pop_cells_merge_nearby$island_cell)

# manually merge
pop_cells_merge_nearby[pop_cells_merge_nearby$cell %in% c(4773,4511,4775,4774), 'group'] <- 'Santa Rosa (SW)'
pop_cells_merge_nearby[pop_cells_merge_nearby$cell %in% c(3722, 3723, 3724), 'group'] <- 'Santa Rosa (NE)'
pop_cells_merge_nearby[pop_cells_merge_nearby$cell %in% c(3460), 'group'] <- 'Santa Rosa (NE, cell 3460)'
pop_cells_merge_nearby[pop_cells_merge_nearby$cell %in% c(39294,38502), 'group'] <- 'San Clemente (two central cells)'
pop_cells_merge_nearby[pop_cells_merge_nearby$cell %in% c(38238, 39034, 38767, 38503, 37707, 38769, 38768, 37972, 37442 ), 'group'] <- 'San Clemente (all other)'
pop_cells_merge_nearby[pop_cells_merge_nearby$cell %in% c(2725), 'group'] <- 'Santa Cruz (E)'
pop_cells_merge_nearby[pop_cells_merge_nearby$cell %in% c(2185), 'group'] <- 'Santa Cruz (Pelican Bay Trail)'
pop_cells_merge_nearby[pop_cells_merge_nearby$cell %in% c(2699, 2700, 2701, 2702), 'group'] <- 'Santa Cruz (W)'
pop_cells_merge_nearby[pop_cells_merge_nearby$cell %in% c(1911, 1912, 2174, 2177), 'group'] <- 'Santa Cruz (central)'
pop_cells_merge_nearby[pop_cells_merge_nearby$cell %in% c(2745), 'group'] <- 'Anacapa'
pop_cells_merge_nearby[pop_cells_merge_nearby$cell %in% c(23193, 21878, 23465, 22929), 'group'] <- 'Catalina'
pop_cells_merge_nearby
```

```
##     cell              island        island_cell
## 1   4773   Santa Rosa Island    Santa_Rosa_4773
## 2   4774   Santa Rosa Island    Santa_Rosa_4774
## 3   4775   Santa Rosa Island    Santa_Rosa_4775
## 4   4511   Santa Rosa Island    Santa_Rosa_4511
## 5   3723   Santa Rosa Island    Santa_Rosa_3723
## 6   3722   Santa Rosa Island    Santa_Rosa_3722
## 7   3724   Santa Rosa Island    Santa_Rosa_3724
## 8   3460   Santa Rosa Island    Santa_Rosa_3460
## 9   2174   Santa Cruz Island    Santa_Cruz_2174
## 10  2177   Santa Cruz Island    Santa_Cruz_2177
## 11  1911   Santa Cruz Island    Santa_Cruz_1911
## 12  2700   Santa Cruz Island    Santa_Cruz_2700
## 13  2725   Santa Cruz Island    Santa_Cruz_2725
## 14  2702   Santa Cruz Island    Santa_Cruz_2702
## 15  2701   Santa Cruz Island    Santa_Cruz_2701
## 16  2699   Santa Cruz Island    Santa_Cruz_2699
## 17  2185   Santa Cruz Island    Santa_Cruz_2185
## 18  1912   Santa Cruz Island    Santa_Cruz_1912
## 19  2745      Anacapa Island       Anacapa_2745
## 20 23193     Catalina Island     Catalina_23193
## 21 21878     Catalina Island     Catalina_21878
## 22 23465     Catalina Island     Catalina_23465
## 23 22929     Catalina Island     Catalina_22929
## 24 37707 San Clemente Island San_Clemente_37707
## 25 39294 San Clemente Island San_Clemente_39294
## 26 38768 San Clemente Island San_Clemente_38768
## 27 38769 San Clemente Island San_Clemente_38769
## 28 37442 San Clemente Island San_Clemente_37442
## 29 37972 San Clemente Island San_Clemente_37972
## 30 38238 San Clemente Island San_Clemente_38238
## 31 38767 San Clemente Island San_Clemente_38767
## 32 38503 San Clemente Island San_Clemente_38503
## 33 38502 San Clemente Island San_Clemente_38502
## 34 39034 San Clemente Island San_Clemente_39034
##                               group
## 1                   Santa Rosa (SW)
## 2                   Santa Rosa (SW)
## 3                   Santa Rosa (SW)
## 4                   Santa Rosa (SW)
## 5                   Santa Rosa (NE)
## 6                   Santa Rosa (NE)
## 7                   Santa Rosa (NE)
## 8        Santa Rosa (NE, cell 3460)
## 9              Santa Cruz (central)
## 10             Santa Cruz (central)
## 11             Santa Cruz (central)
## 12                   Santa Cruz (W)
## 13                   Santa Cruz (E)
## 14                   Santa Cruz (W)
## 15                   Santa Cruz (W)
## 16                   Santa Cruz (W)
## 17   Santa Cruz (Pelican Bay Trail)
## 18             Santa Cruz (central)
## 19                          Anacapa
## 20                         Catalina
## 21                         Catalina
## 22                         Catalina
## 23                         Catalina
## 24         San Clemente (all other)
## 25 San Clemente (two central cells)
## 26         San Clemente (all other)
## 27         San Clemente (all other)
## 28         San Clemente (all other)
## 29         San Clemente (all other)
## 30         San Clemente (all other)
## 31         San Clemente (all other)
## 32         San Clemente (all other)
## 33 San Clemente (two central cells)
## 34         San Clemente (all other)
```

```
groups <- unique(pop_cells_merge_nearby$group)


# new list
csi.merge <- list()

# loop through the four climates and merge similar sites
for(n in 1:length(csi)){

  full <- csi[[n]]
  
  # make new matrix
  csi.merge[[n]] <- as.data.frame(matrix(nrow = length(groups), ncol = length(groups)))
  colnames(csi.merge[[n]]) <- groups
  rownames(csi.merge[[n]]) <- groups
  
  # calculate average for each group - loop through each combination of groups
  for(a in 1:length(groups)){
    
    group1 <- groups[a]
    indivs1 <- pop_cells_merge_nearby[pop_cells_merge_nearby$group == group1, 'island_cell']
    
    for(b in 1:length(groups)){
      
      group2 <- groups[b]
      indivs2 <- pop_cells_merge_nearby[pop_cells_merge_nearby$group == group2, 'island_cell']
      
      sub <- full[indivs1, indivs2]
      
      csi.merge[[n]][group1, group2] <- mean(unlist(sub))
      
    }
  }

}

names(csi.merge) <- names(csi)


colf <- colorRampPalette(brewer.pal(9, 'YlGn'))
#heatmap(as.matrix(csi.merge$MIROC_4.5_2050), col = colf(100), Rowv = NA, Colv = NA, margins = c(12, 10), scale = 'none', xlab = 'source site', ylab = 'planting site')


for(n in 1:length(csi.merge)){
  
  mat <- as.matrix(csi.merge[[n]][colnames(csi.merge[[n]]),])
  # order by best planting site (overall)
  row_ord <- order(rowSums(mat))
  
 #png(file = paste('results/gradient_forest/climate_suitability_heatmap_mergedSites_noGuadAna_', names(csi.merge)[n], '.png', sep = ''), height = 8, width = 8, res = 300, units = 'in')
  heatmap(mat[row_ord,], col = colf(100), Rowv = NA, Colv = NA, margins = c(15, 15), scale = 'none', xlab = 'source site', ylab = 'planting site', main = paste('climate suitability', names(off.agf)[n]))
  #dev.off()

}
```

```
# prettier version with with complex heatmap

# order columns geographically
dput(colnames(mat))
```

```
## c("Santa Rosa (SW)", "Santa Rosa (NE)", "Santa Rosa (NE, cell 3460)", 
## "Santa Cruz (central)", "Santa Cruz (W)", "Santa Cruz (E)", "Santa Cruz (Pelican Bay Trail)", 
## "Anacapa", "Catalina", "San Clemente (all other)", "San Clemente (two central cells)"
## )
```

```
col_ord <- c("Santa Rosa (SW)", "Santa Rosa (NE)", "Santa Rosa (NE, cell 3460)",  "Santa Cruz (W)", "Santa Cruz (central)",  "Santa Cruz (Pelican Bay Trail)", "Santa Cruz (E)", "Anacapa", "Catalina", "San Clemente (two central cells)", "San Clemente (all other)")

for(n in 1:length(csi.merge)){
  
  mat <- as.matrix(csi.merge[[n]][colnames(csi.merge[[n]]),])
  # order rows by best planting site (overall)
  row_ord <- rev(order(rowSums(mat)))
  
  # order columns by geography
  col_ord <- c("Santa Rosa (SW)", "Santa Rosa (NE)", "Santa Rosa (NE, cell 3460)",  "Santa Cruz (W)", "Santa Cruz (central)",  "Santa Cruz (Pelican Bay Trail)", "Santa Cruz (E)", "Anacapa", "Catalina", "San Clemente (two central cells)", "San Clemente (all other)")
  
  
  hm <- mat[row_ord,col_ord]
  
  #png(file = paste('results/gradient_forest/noGuadAna/climate_suitability_heatmap_mergedSites_noGuadAna_', names(csi.merge)[n], '.png', sep = ''), height = 8, width = 8, res = 300, units = 'in')
  
    #pdf(file = paste('results/gradient_forest/noGuadAna/climate_suitability_heatmap_mergedSites_noGuadAna_', names(csi.merge)[n], '.pdf', sep = ''), height = 8, width = 8)
  
  par(mar = c(5,10,6,10))
  hmplot <- Heatmap(hm, 
          col = colf(100),
          cluster_rows = F, 
          cluster_columns = F,
          #main = 'climate model',
          row_title = 'Planting Site',
          column_title = 'Source Site',
          row_title_gp = gpar(fontsize = 20, fontface = 'bold'),
          column_title_gp = gpar(fontsize = 20, fontface = 'bold'),
          row_names_side = 'left',
          column_names_side = 'top',
          column_names_rot = 65,
          heatmap_legend_param = list(title = 'Relative\nClimate\nSuitability', title_gp = gpar(fontsize = 12)))
  
  plot(hmplot)
  
  #dev.off()
  
}
```
