## Supplementary Rmarkdown Outputs for "Comparison of conservation strategies for California Channel Island Oak (*Quercus tomentella*) using climate suitability predicted from genomic data": read_and_plot_fast-wcfst.html

Plot Fst


### Plot Fst

#### 2024-09-11

- 1 Fst by island
- 2 Fst by island and ancestry

Plot Fst results output from fast-wcfst

### 1 Fst by island

```
library(ComplexHeatmap)
```

```
## Loading required package: grid
```

```
## ========================================
## ComplexHeatmap version 2.20.0
## Bioconductor page: http://bioconductor.org/packages/ComplexHeatmap/
## Github page: https://github.com/jokergoo/ComplexHeatmap
## Documentation: http://jokergoo.github.io/ComplexHeatmap-reference
## 
## If you use it in published research, please cite either one:
## - Gu, Z. Complex Heatmap Visualization. iMeta 2022.
## - Gu, Z. Complex heatmaps reveal patterns and correlations in multidimensional 
##     genomic data. Bioinformatics 2016.
## 
## 
## The new InteractiveComplexHeatmap package can directly export static 
## complex heatmaps into an interactive Shiny app with zero effort. Have a try!
## 
## This message can be suppressed by:
##   suppressPackageStartupMessages(library(ComplexHeatmap))
## ========================================
```

```
library(RColorBrewer)

fst <- read.csv('/home/alayna/Documents/research/projects/2022_island_oak/results/popgen_stats/fast-wcfst_output_island.tsv', sep = '\t')
str(fst)
```

```
## 'data.frame':    21 obs. of  5 variables:
##  $ Population.A : chr  "Catalina Island" "San Clemente Island" "San Clemente Island" "Santa Cruz Island" ...
##  $ Population.B : chr  "Mainland" "Mainland" "Catalina Island" "Mainland" ...
##  $ Mean.Fst     : num  0.02415 0.02436 0.00918 0.03837 0.01553 ...
##  $ Mean.Variance: num  0.00633 0.00678 0.00171 0.01 0.00204 ...
##  $ Weighted.Fst : num  0.0349 0.0399 0.0113 0.0591 0.0194 ...
```

```
sessionInfo()
```

```
## R version 4.4.1 (2024-06-14)
## Platform: x86_64-pc-linux-gnu
## Running under: Arch Linux
## 
## Matrix products: default
## BLAS:   /usr/lib/libblas.so.3.12.0 
## LAPACK: /usr/lib/liblapack.so.3.12.0
## 
## locale:
##  [1] LC_CTYPE=en_US.UTF-8       LC_NUMERIC=C              
##  [3] LC_TIME=en_US.UTF-8        LC_COLLATE=en_US.UTF-8    
##  [5] LC_MONETARY=en_US.UTF-8    LC_MESSAGES=en_US.UTF-8   
##  [7] LC_PAPER=en_US.UTF-8       LC_NAME=C                 
##  [9] LC_ADDRESS=C               LC_TELEPHONE=C            
## [11] LC_MEASUREMENT=en_US.UTF-8 LC_IDENTIFICATION=C       
## 
## time zone: America/New_York
## tzcode source: system (glibc)
## 
## attached base packages:
## [1] grid      stats     graphics  grDevices utils     datasets  methods  
## [8] base     
## 
## other attached packages:
## [1] RColorBrewer_1.1-3    ComplexHeatmap_2.20.0
## 
## loaded via a namespace (and not attached):
##  [1] crayon_1.5.2        doParallel_1.0.17   cli_3.6.2          
##  [4] knitr_1.46          rlang_1.1.3         xfun_0.44          
##  [7] circlize_0.4.16     png_0.1-8           clue_0.3-65        
## [10] jsonlite_1.8.8      S4Vectors_0.42.0    rjson_0.2.22       
## [13] colorspace_2.1-0    htmltools_0.5.8.1   GlobalOptions_0.1.2
## [16] stats4_4.4.1        sass_0.4.9          rmarkdown_2.27     
## [19] evaluate_0.23       jquerylib_0.1.4     fastmap_1.2.0      
## [22] IRanges_2.38.0      yaml_2.3.8          foreach_1.5.2      
## [25] lifecycle_1.0.4     cluster_2.1.6       compiler_4.4.1     
## [28] codetools_0.2-20    rstudioapi_0.16.0   digest_0.6.35      
## [31] R6_2.5.1            shape_1.4.6.1       parallel_4.4.1     
## [34] bslib_0.7.0         GetoptLong_1.0.5    tools_4.4.1        
## [37] iterators_1.0.14    matrixStats_1.3.0   BiocGenerics_0.50.0
## [40] cachem_1.1.0
```

```
# get list of islands from fst file
isl <- unique(c(fst$Population.A, fst$Population.B))

# reorder
dput(isl)
```

```
## c("Catalina Island", "San Clemente Island", "Santa Cruz Island", 
## "Santa Rosa Island", "Guadalupe Island", "Anacapa Island", "Mainland"
## )
```

```
isl <- c( "Mainland", "Santa Rosa Island", "Santa Cruz Island", "Anacapa Island", "Catalina Island", "San Clemente Island", "Guadalupe Island")


fst.df <- as.data.frame(matrix(nrow = length(isl), ncol = length(isl)))
colnames(fst.df) <- isl
rownames(fst.df) <- isl

pairs <- combn(isl, 2)


for(n in 1:nrow(fst)){
  
  pop1 <- fst$Population.A[n]
  pop2 <- fst$Population.B[n]
  
  # add Fst to both sides of diagonal
  fst.df[pop1, pop2] <- fst$Mean.Fst[n]
  fst.df[pop2, pop1] <- fst$Mean.Fst[n]
  
}

# make version with rounded values for nicer display
fst.df.round <- round(fst.df, 3)

#############################
# get weighted value 
fst.weight <- as.data.frame(matrix(nrow = length(isl), ncol = length(isl)))
colnames(fst.weight) <- isl
rownames(fst.weight) <- isl

for(n in 1:nrow(fst)){
  
  pop1 <- fst$Population.A[n]
  pop2 <- fst$Population.B[n]
  
  # add Fst to both sides of diagonal
  fst.weight[pop1, pop2] <- fst$Weighted.Fst[n]
  fst.weight[pop2, pop1] <- fst$Weighted.Fst[n]
  
}

# make version with rounded values for nicer display
fst.weight.round <- round(fst.weight, 3)


#################################
# variance

fst.var <- as.data.frame(matrix(nrow = length(isl), ncol = length(isl)))
colnames(fst.var) <- isl
rownames(fst.var) <- isl

for(n in 1:nrow(fst)){
  
  pop1 <- fst$Population.A[n]
  pop2 <- fst$Population.B[n]
  
  # add Fst to both sides of diagonal
  fst.var[pop1, pop2] <- fst$Mean.Variance[n]
  fst.var[pop2, pop1] <- fst$Mean.Variance[n]
  
}

# make version with rounded values for nicer display
fst.var.round <- round(fst.var, 3)


########
# make new dataframe including fst and the upper and lower limits

# calculate upper and lower
fst.low <- fst.df - sqrt(fst.var)
fst.low.round <- round(fst.low, 3)

fst.high <- fst.df + sqrt(fst.var)
fst.high.round <- round(fst.high, 3)

fst.text <- fst.df

pairs <- combn(colnames(fst.df), 2)
for(n in 1:ncol(pairs)){
  
  pop1 <- pairs[1,n]
  pop2 <- pairs[2,n]
  
  text <- paste(fst.df.round[pop1,pop2], ' (', fst.low.round[pop1, pop2], ', ', fst.high.round[pop1, pop2], ')', sep = '')
  
  fst.text[pop1, pop2] <- text
  fst.text[pop2, pop1] <- text
  
}

fst.text
```

```
##                                  Mainland     Santa Rosa Island
## Mainland                             <NA>  0.063 (-0.064, 0.19)
## Santa Rosa Island    0.063 (-0.064, 0.19)                  <NA>
## Santa Cruz Island   0.038 (-0.062, 0.138) 0.015 (-0.026, 0.057)
## Anacapa Island      0.065 (-0.098, 0.227) 0.036 (-0.095, 0.167)
## Catalina Island     0.024 (-0.055, 0.104) 0.029 (-0.035, 0.093)
## San Clemente Island 0.024 (-0.058, 0.107) 0.021 (-0.033, 0.075)
## Guadalupe Island    0.053 (-0.056, 0.163) 0.059 (-0.048, 0.166)
##                         Santa Cruz Island        Anacapa Island
## Mainland            0.038 (-0.062, 0.138) 0.065 (-0.098, 0.227)
## Santa Rosa Island   0.015 (-0.026, 0.057) 0.036 (-0.095, 0.167)
## Santa Cruz Island                    <NA> 0.014 (-0.094, 0.121)
## Anacapa Island      0.014 (-0.094, 0.121)                  <NA>
## Catalina Island      0.016 (-0.03, 0.061)  0.03 (-0.093, 0.152)
## San Clemente Island  0.01 (-0.029, 0.049)  0.021 (-0.09, 0.132)
## Guadalupe Island    0.045 (-0.046, 0.136) 0.075 (-0.098, 0.247)
##                           Catalina Island   San Clemente Island
## Mainland            0.024 (-0.055, 0.104) 0.024 (-0.058, 0.107)
## Santa Rosa Island   0.029 (-0.035, 0.093) 0.021 (-0.033, 0.075)
## Santa Cruz Island    0.016 (-0.03, 0.061)  0.01 (-0.029, 0.049)
## Anacapa Island       0.03 (-0.093, 0.152)  0.021 (-0.09, 0.132)
## Catalina Island                      <NA> 0.009 (-0.032, 0.051)
## San Clemente Island 0.009 (-0.032, 0.051)                  <NA>
## Guadalupe Island    0.043 (-0.042, 0.127) 0.041 (-0.043, 0.125)
##                          Guadalupe Island
## Mainland            0.053 (-0.056, 0.163)
## Santa Rosa Island   0.059 (-0.048, 0.166)
## Santa Cruz Island   0.045 (-0.046, 0.136)
## Anacapa Island      0.075 (-0.098, 0.247)
## Catalina Island     0.043 (-0.042, 0.127)
## San Clemente Island 0.041 (-0.043, 0.125)
## Guadalupe Island                     <NA>
```

```
# add column for average
fst.text$average <- rowMeans(fst.df, na.rm = T)
# exclude mainland
fst.text$average_islands <- c(NA, rowMeans(fst.df[rownames(fst.df)!= 'Mainland',colnames(fst.df)!= 'Mainland'], na.rm = T))
fst.text$average <- round(fst.text$average, 3)
fst.text$average_islands <- round(fst.text$average_islands, 3)

fst.text
```

```
##                                  Mainland     Santa Rosa Island
## Mainland                             <NA>  0.063 (-0.064, 0.19)
## Santa Rosa Island    0.063 (-0.064, 0.19)                  <NA>
## Santa Cruz Island   0.038 (-0.062, 0.138) 0.015 (-0.026, 0.057)
## Anacapa Island      0.065 (-0.098, 0.227) 0.036 (-0.095, 0.167)
## Catalina Island     0.024 (-0.055, 0.104) 0.029 (-0.035, 0.093)
## San Clemente Island 0.024 (-0.058, 0.107) 0.021 (-0.033, 0.075)
## Guadalupe Island    0.053 (-0.056, 0.163) 0.059 (-0.048, 0.166)
##                         Santa Cruz Island        Anacapa Island
## Mainland            0.038 (-0.062, 0.138) 0.065 (-0.098, 0.227)
## Santa Rosa Island   0.015 (-0.026, 0.057) 0.036 (-0.095, 0.167)
## Santa Cruz Island                    <NA> 0.014 (-0.094, 0.121)
## Anacapa Island      0.014 (-0.094, 0.121)                  <NA>
## Catalina Island      0.016 (-0.03, 0.061)  0.03 (-0.093, 0.152)
## San Clemente Island  0.01 (-0.029, 0.049)  0.021 (-0.09, 0.132)
## Guadalupe Island    0.045 (-0.046, 0.136) 0.075 (-0.098, 0.247)
##                           Catalina Island   San Clemente Island
## Mainland            0.024 (-0.055, 0.104) 0.024 (-0.058, 0.107)
## Santa Rosa Island   0.029 (-0.035, 0.093) 0.021 (-0.033, 0.075)
## Santa Cruz Island    0.016 (-0.03, 0.061)  0.01 (-0.029, 0.049)
## Anacapa Island       0.03 (-0.093, 0.152)  0.021 (-0.09, 0.132)
## Catalina Island                      <NA> 0.009 (-0.032, 0.051)
## San Clemente Island 0.009 (-0.032, 0.051)                  <NA>
## Guadalupe Island    0.043 (-0.042, 0.127) 0.041 (-0.043, 0.125)
##                          Guadalupe Island average average_islands
## Mainland            0.053 (-0.056, 0.163)   0.045              NA
## Santa Rosa Island   0.059 (-0.048, 0.166)   0.037           0.032
## Santa Cruz Island   0.045 (-0.046, 0.136)   0.023           0.020
## Anacapa Island      0.075 (-0.098, 0.247)   0.040           0.035
## Catalina Island     0.043 (-0.042, 0.127)   0.025           0.025
## San Clemente Island 0.041 (-0.043, 0.125)   0.021           0.020
## Guadalupe Island                     <NA>   0.053           0.052
```

```
#write.csv(fst.text, file = 'results/popgen_stats/Fst_island_table.csv')
```

```
# heatmap

# make version with rounded values for nicer display
fst.df.round <- round(fst.df, 3)

#png(file = 'results/popgen_stats/Fst_heatmap_fast-wcfst.png', width = 8, height = 8, res = 300, units = 'in')
par(mar = c(5,4,4,10))
Heatmap(as.matrix(fst.df),
        cluster_rows = F, cluster_columns = F,
        row_names_side = 'left',
        column_names_side = 'top',
        column_names_gp = gpar(fontsize = 14),
        row_names_gp = gpar(fontsize = 14),
        column_names_rot = 90,
        col = brewer.pal(9, 'YlOrRd'),
        show_heatmap_legend = F,
        cell_fun = function(j, i, x, y, width, height, fill) {
          grid.text(fst.df.round[i,j], x, y, gp = gpar(fontsize = 14))
        })
```

```
#dev.off()

# weighted

#png(file = 'results/popgen_stats/Fst_weighted_heatmap_fast-wcfst.png', width = 8, height = 8, res = 300, units = 'in')
par(mar = c(5,4,4,10))
Heatmap(as.matrix(fst.weight),
        cluster_rows = F, cluster_columns = F,
        row_names_side = 'left',
        column_names_side = 'top',
        column_names_gp = gpar(fontsize = 14),
        row_names_gp = gpar(fontsize = 14),
        column_names_rot = 90,
        col = brewer.pal(9, 'YlOrRd'),
        show_heatmap_legend = F,
        cell_fun = function(j, i, x, y, width, height, fill) {
          grid.text(fst.weight.round[i,j], x, y, gp = gpar(fontsize = 14))
        })
```

```
#dev.off()


# variance

#png(file = 'results/popgen_stats/Fst_variance_heatmap_fast-wcfst.png', width = 8, height = 8, res = 300, units = 'in')
par(mar = c(5,4,4,10))
Heatmap(as.matrix(fst.var),
        cluster_rows = F, cluster_columns = F,
        row_names_side = 'left',
        column_names_side = 'top',
        column_names_gp = gpar(fontsize = 14),
        row_names_gp = gpar(fontsize = 14),
        column_names_rot = 90,
        col = brewer.pal(9, 'YlOrRd'),
        show_heatmap_legend = F,
        cell_fun = function(j, i, x, y, width, height, fill) {
          grid.text(fst.var.round[i,j], x, y, gp = gpar(fontsize = 14))
        })
```

```
#dev.off()
```

### 2 Fst by island and ancestry

```
rm(fst)

fst <- read.csv('/home/alayna/Documents/research/projects/2022_island_oak/results/popgen_stats/fast-wcfst_output_ancestry.tsv', sep = '\t')
str(fst)
```

```
## 'data.frame':    55 obs. of  5 variables:
##  $ Population.A : chr  "Catalina Island_hybrid" "San Clemente Island_hybrid" "San Clemente Island_hybrid" "Santa Cruz Island_hybrid" ...
##  $ Population.B : chr  "Mainland_Qchr" "Mainland_Qchr" "Catalina Island_hybrid" "Mainland_Qchr" ...
##  $ Mean.Fst     : num  0.02722 0.02595 0.00963 0.02777 0.01349 ...
##  $ Mean.Variance: num  0.00711 0.00719 0.00206 0.00749 0.00266 ...
##  $ Weighted.Fst : num  0.0409 0.0427 0.0122 0.0447 0.0171 ...
```

```
# get list of islands from fst file
isl <- unique(c(fst$Population.A, fst$Population.B))

# reorder
dput(isl)
```

```
## c("Catalina Island_hybrid", "San Clemente Island_hybrid", "Santa Cruz Island_hybrid", 
## "Catalina Island_Qchr", "Santa Rosa Island_Qtom", "Guadalupe Island_Guadalupe", 
## "Guadalupe Island_hybrid", "San Clemente Island_Qchr", "Santa Cruz Island_Qtom", 
## "Anacapa Island_Qtom", "Mainland_Qchr")
```

```
isl <- c( "Mainland", "Santa Rosa Island", "Santa Cruz Island", "Anacapa Island", "Catalina Island", "San Clemente Island", "Guadalupe Island")

isl <- c("Mainland_Qchr", "Santa Rosa Island_Qtom", "Santa Cruz Island_Qtom", "Santa Cruz Island_hybrid", "Anacapa Island_Qtom", "Catalina Island_hybrid", "Catalina Island_Qchr", "San Clemente Island_hybrid","San Clemente Island_Qchr", "Guadalupe Island_Guadalupe", "Guadalupe Island_hybrid")


fst.df <- as.data.frame(matrix(nrow = length(isl), ncol = length(isl)))
colnames(fst.df) <- isl
rownames(fst.df) <- isl

for(n in 1:nrow(fst)){
  
  pop1 <- fst$Population.A[n]
  pop2 <- fst$Population.B[n]
  
  # add Fst to both sides of diagonal
  fst.df[pop1, pop2] <- fst$Mean.Fst[n]
  fst.df[pop2, pop1] <- fst$Mean.Fst[n]
  
}

# make version with rounded values for nicer display
fst.df.round <- round(fst.df, 3)

#############################
# get weighted value 
fst.weight <- as.data.frame(matrix(nrow = length(isl), ncol = length(isl)))
colnames(fst.weight) <- isl
rownames(fst.weight) <- isl

for(n in 1:nrow(fst)){
  
  pop1 <- fst$Population.A[n]
  pop2 <- fst$Population.B[n]
  
  # add Fst to both sides of diagonal
  fst.weight[pop1, pop2] <- fst$Weighted.Fst[n]
  fst.weight[pop2, pop1] <- fst$Weighted.Fst[n]
  
}

# make version with rounded values for nicer display
fst.weight.round <- round(fst.weight, 3)


#################################
# variance

fst.var <- as.data.frame(matrix(nrow = length(isl), ncol = length(isl)))
colnames(fst.var) <- isl
rownames(fst.var) <- isl

for(n in 1:nrow(fst)){
  
  pop1 <- fst$Population.A[n]
  pop2 <- fst$Population.B[n]
  
  # add Fst to both sides of diagonal
  fst.var[pop1, pop2] <- fst$Mean.Variance[n]
  fst.var[pop2, pop1] <- fst$Mean.Variance[n]
  
}

# make version with rounded values for nicer display
fst.var.round <- round(fst.var, 3)


########
# make new dataframe including fst and the upper and lower limits

# calculate upper and lower
fst.low <- fst.df - sqrt(fst.var)
fst.low.round <- round(fst.low, 3)

fst.high <- fst.df + sqrt(fst.var)
fst.high.round <- round(fst.high, 3)

fst.text <- fst.df

pairs <- combn(colnames(fst.df), 2)
for(n in 1:ncol(pairs)){
  
  pop1 <- pairs[1,n]
  pop2 <- pairs[2,n]
  
  text <- paste(fst.df.round[pop1,pop2], ' (', fst.low.round[pop1, pop2], ', ', fst.high.round[pop1, pop2], ')', sep = '')
  
  fst.text[pop1, pop2] <- text
  fst.text[pop2, pop1] <- text
  
}

fst.text
```

```
##                                     Mainland_Qchr Santa Rosa Island_Qtom
## Mainland_Qchr                                <NA>   0.063 (-0.064, 0.19)
## Santa Rosa Island_Qtom       0.063 (-0.064, 0.19)                   <NA>
## Santa Cruz Island_Qtom       0.05 (-0.064, 0.164)  0.017 (-0.034, 0.069)
## Santa Cruz Island_hybrid    0.028 (-0.059, 0.114)  0.025 (-0.038, 0.088)
## Anacapa Island_Qtom         0.065 (-0.098, 0.227)  0.036 (-0.095, 0.167)
## Catalina Island_hybrid      0.027 (-0.057, 0.112)  0.029 (-0.038, 0.095)
## Catalina Island_Qchr        0.014 (-0.121, 0.148)   0.059 (-0.13, 0.249)
## San Clemente Island_hybrid  0.026 (-0.059, 0.111)  0.021 (-0.034, 0.075)
## San Clemente Island_Qchr   -0.099 (-0.433, 0.234) -0.055 (-0.408, 0.298)
## Guadalupe Island_Guadalupe  0.056 (-0.056, 0.169)   0.061 (-0.05, 0.171)
## Guadalupe Island_hybrid    -0.019 (-0.266, 0.229)  0.034 (-0.259, 0.327)
##                            Santa Cruz Island_Qtom Santa Cruz Island_hybrid
## Mainland_Qchr                0.05 (-0.064, 0.164)    0.028 (-0.059, 0.114)
## Santa Rosa Island_Qtom      0.017 (-0.034, 0.069)    0.025 (-0.038, 0.088)
## Santa Cruz Island_Qtom                       <NA>    0.012 (-0.038, 0.063)
## Santa Cruz Island_hybrid    0.012 (-0.038, 0.063)                     <NA>
## Anacapa Island_Qtom          0.022 (-0.086, 0.13)    0.024 (-0.089, 0.136)
## Catalina Island_hybrid        0.02 (-0.039, 0.08)    0.013 (-0.038, 0.065)
## Catalina Island_Qchr        0.043 (-0.121, 0.207)    0.017 (-0.122, 0.156)
## San Clemente Island_hybrid  0.013 (-0.037, 0.063)    0.009 (-0.037, 0.055)
## San Clemente Island_Qchr   -0.061 (-0.399, 0.278)   -0.102 (-0.415, 0.211)
## Guadalupe Island_Guadalupe   0.057 (-0.05, 0.163)    0.047 (-0.047, 0.141)
## Guadalupe Island_hybrid      0.021 (-0.25, 0.291)   -0.015 (-0.261, 0.232)
##                              Anacapa Island_Qtom Catalina Island_hybrid
## Mainland_Qchr              0.065 (-0.098, 0.227)  0.027 (-0.057, 0.112)
## Santa Rosa Island_Qtom     0.036 (-0.095, 0.167)  0.029 (-0.038, 0.095)
## Santa Cruz Island_Qtom      0.022 (-0.086, 0.13)    0.02 (-0.039, 0.08)
## Santa Cruz Island_hybrid   0.024 (-0.089, 0.136)  0.013 (-0.038, 0.065)
## Anacapa Island_Qtom                         <NA>     0.03 (-0.09, 0.15)
## Catalina Island_hybrid        0.03 (-0.09, 0.15)                   <NA>
## Catalina Island_Qchr       0.078 (-0.101, 0.257)  0.011 (-0.121, 0.143)
## San Clemente Island_hybrid 0.021 (-0.089, 0.132)   0.01 (-0.036, 0.055)
## San Clemente Island_Qchr   0.063 (-0.323, 0.449)   -0.1 (-0.409, 0.209)
## Guadalupe Island_Guadalupe 0.079 (-0.095, 0.252)  0.047 (-0.045, 0.139)
## Guadalupe Island_hybrid    0.094 (-0.185, 0.372) -0.012 (-0.259, 0.235)
##                              Catalina Island_Qchr San Clemente Island_hybrid
## Mainland_Qchr               0.014 (-0.121, 0.148)      0.026 (-0.059, 0.111)
## Santa Rosa Island_Qtom       0.059 (-0.13, 0.249)      0.021 (-0.034, 0.075)
## Santa Cruz Island_Qtom      0.043 (-0.121, 0.207)      0.013 (-0.037, 0.063)
## Santa Cruz Island_hybrid    0.017 (-0.122, 0.156)      0.009 (-0.037, 0.055)
## Anacapa Island_Qtom         0.078 (-0.101, 0.257)      0.021 (-0.089, 0.132)
## Catalina Island_hybrid      0.011 (-0.121, 0.143)       0.01 (-0.036, 0.055)
## Catalina Island_Qchr                         <NA>       0.013 (-0.124, 0.15)
## San Clemente Island_hybrid   0.013 (-0.124, 0.15)                       <NA>
## San Clemente Island_Qchr   -0.037 (-0.421, 0.346)     -0.112 (-0.417, 0.194)
## Guadalupe Island_Guadalupe  0.065 (-0.118, 0.248)      0.044 (-0.046, 0.133)
## Guadalupe Island_hybrid      0.016 (-0.25, 0.281)      -0.02 (-0.264, 0.224)
##                            San Clemente Island_Qchr Guadalupe Island_Guadalupe
## Mainland_Qchr                -0.099 (-0.433, 0.234)      0.056 (-0.056, 0.169)
## Santa Rosa Island_Qtom       -0.055 (-0.408, 0.298)       0.061 (-0.05, 0.171)
## Santa Cruz Island_Qtom       -0.061 (-0.399, 0.278)       0.057 (-0.05, 0.163)
## Santa Cruz Island_hybrid     -0.102 (-0.415, 0.211)      0.047 (-0.047, 0.141)
## Anacapa Island_Qtom           0.063 (-0.323, 0.449)      0.079 (-0.095, 0.252)
## Catalina Island_hybrid         -0.1 (-0.409, 0.209)      0.047 (-0.045, 0.139)
## Catalina Island_Qchr         -0.037 (-0.421, 0.346)      0.065 (-0.118, 0.248)
## San Clemente Island_hybrid   -0.112 (-0.417, 0.194)      0.044 (-0.046, 0.133)
## San Clemente Island_Qchr                       <NA>     -0.044 (-0.389, 0.302)
## Guadalupe Island_Guadalupe   -0.044 (-0.389, 0.302)                       <NA>
## Guadalupe Island_hybrid      -0.045 (-0.513, 0.424)     -0.014 (-0.244, 0.217)
##                            Guadalupe Island_hybrid
## Mainland_Qchr               -0.019 (-0.266, 0.229)
## Santa Rosa Island_Qtom       0.034 (-0.259, 0.327)
## Santa Cruz Island_Qtom        0.021 (-0.25, 0.291)
## Santa Cruz Island_hybrid    -0.015 (-0.261, 0.232)
## Anacapa Island_Qtom          0.094 (-0.185, 0.372)
## Catalina Island_hybrid      -0.012 (-0.259, 0.235)
## Catalina Island_Qchr          0.016 (-0.25, 0.281)
## San Clemente Island_hybrid   -0.02 (-0.264, 0.224)
## San Clemente Island_Qchr    -0.045 (-0.513, 0.424)
## Guadalupe Island_Guadalupe  -0.014 (-0.244, 0.217)
## Guadalupe Island_hybrid                       <NA>
```

```
# add column for average
fst.text$average <- rowMeans(fst.df, na.rm = T)
# exclude mainland
fst.text$average_islands <- c(NA, rowMeans(fst.df[rownames(fst.df)!= 'Mainland_Qchr',colnames(fst.df)!= 'Mainland_Qchr'], na.rm = T))
fst.text$average <- round(fst.text$average, 3)
fst.text$average_islands <- round(fst.text$average_islands, 3)

fst.text
```

```
##                                     Mainland_Qchr Santa Rosa Island_Qtom
## Mainland_Qchr                                <NA>   0.063 (-0.064, 0.19)
## Santa Rosa Island_Qtom       0.063 (-0.064, 0.19)                   <NA>
## Santa Cruz Island_Qtom       0.05 (-0.064, 0.164)  0.017 (-0.034, 0.069)
## Santa Cruz Island_hybrid    0.028 (-0.059, 0.114)  0.025 (-0.038, 0.088)
## Anacapa Island_Qtom         0.065 (-0.098, 0.227)  0.036 (-0.095, 0.167)
## Catalina Island_hybrid      0.027 (-0.057, 0.112)  0.029 (-0.038, 0.095)
## Catalina Island_Qchr        0.014 (-0.121, 0.148)   0.059 (-0.13, 0.249)
## San Clemente Island_hybrid  0.026 (-0.059, 0.111)  0.021 (-0.034, 0.075)
## San Clemente Island_Qchr   -0.099 (-0.433, 0.234) -0.055 (-0.408, 0.298)
## Guadalupe Island_Guadalupe  0.056 (-0.056, 0.169)   0.061 (-0.05, 0.171)
## Guadalupe Island_hybrid    -0.019 (-0.266, 0.229)  0.034 (-0.259, 0.327)
##                            Santa Cruz Island_Qtom Santa Cruz Island_hybrid
## Mainland_Qchr                0.05 (-0.064, 0.164)    0.028 (-0.059, 0.114)
## Santa Rosa Island_Qtom      0.017 (-0.034, 0.069)    0.025 (-0.038, 0.088)
## Santa Cruz Island_Qtom                       <NA>    0.012 (-0.038, 0.063)
## Santa Cruz Island_hybrid    0.012 (-0.038, 0.063)                     <NA>
## Anacapa Island_Qtom          0.022 (-0.086, 0.13)    0.024 (-0.089, 0.136)
## Catalina Island_hybrid        0.02 (-0.039, 0.08)    0.013 (-0.038, 0.065)
## Catalina Island_Qchr        0.043 (-0.121, 0.207)    0.017 (-0.122, 0.156)
## San Clemente Island_hybrid  0.013 (-0.037, 0.063)    0.009 (-0.037, 0.055)
## San Clemente Island_Qchr   -0.061 (-0.399, 0.278)   -0.102 (-0.415, 0.211)
## Guadalupe Island_Guadalupe   0.057 (-0.05, 0.163)    0.047 (-0.047, 0.141)
## Guadalupe Island_hybrid      0.021 (-0.25, 0.291)   -0.015 (-0.261, 0.232)
##                              Anacapa Island_Qtom Catalina Island_hybrid
## Mainland_Qchr              0.065 (-0.098, 0.227)  0.027 (-0.057, 0.112)
## Santa Rosa Island_Qtom     0.036 (-0.095, 0.167)  0.029 (-0.038, 0.095)
## Santa Cruz Island_Qtom      0.022 (-0.086, 0.13)    0.02 (-0.039, 0.08)
## Santa Cruz Island_hybrid   0.024 (-0.089, 0.136)  0.013 (-0.038, 0.065)
## Anacapa Island_Qtom                         <NA>     0.03 (-0.09, 0.15)
## Catalina Island_hybrid        0.03 (-0.09, 0.15)                   <NA>
## Catalina Island_Qchr       0.078 (-0.101, 0.257)  0.011 (-0.121, 0.143)
## San Clemente Island_hybrid 0.021 (-0.089, 0.132)   0.01 (-0.036, 0.055)
## San Clemente Island_Qchr   0.063 (-0.323, 0.449)   -0.1 (-0.409, 0.209)
## Guadalupe Island_Guadalupe 0.079 (-0.095, 0.252)  0.047 (-0.045, 0.139)
## Guadalupe Island_hybrid    0.094 (-0.185, 0.372) -0.012 (-0.259, 0.235)
##                              Catalina Island_Qchr San Clemente Island_hybrid
## Mainland_Qchr               0.014 (-0.121, 0.148)      0.026 (-0.059, 0.111)
## Santa Rosa Island_Qtom       0.059 (-0.13, 0.249)      0.021 (-0.034, 0.075)
## Santa Cruz Island_Qtom      0.043 (-0.121, 0.207)      0.013 (-0.037, 0.063)
## Santa Cruz Island_hybrid    0.017 (-0.122, 0.156)      0.009 (-0.037, 0.055)
## Anacapa Island_Qtom         0.078 (-0.101, 0.257)      0.021 (-0.089, 0.132)
## Catalina Island_hybrid      0.011 (-0.121, 0.143)       0.01 (-0.036, 0.055)
## Catalina Island_Qchr                         <NA>       0.013 (-0.124, 0.15)
## San Clemente Island_hybrid   0.013 (-0.124, 0.15)                       <NA>
## San Clemente Island_Qchr   -0.037 (-0.421, 0.346)     -0.112 (-0.417, 0.194)
## Guadalupe Island_Guadalupe  0.065 (-0.118, 0.248)      0.044 (-0.046, 0.133)
## Guadalupe Island_hybrid      0.016 (-0.25, 0.281)      -0.02 (-0.264, 0.224)
##                            San Clemente Island_Qchr Guadalupe Island_Guadalupe
## Mainland_Qchr                -0.099 (-0.433, 0.234)      0.056 (-0.056, 0.169)
## Santa Rosa Island_Qtom       -0.055 (-0.408, 0.298)       0.061 (-0.05, 0.171)
## Santa Cruz Island_Qtom       -0.061 (-0.399, 0.278)       0.057 (-0.05, 0.163)
## Santa Cruz Island_hybrid     -0.102 (-0.415, 0.211)      0.047 (-0.047, 0.141)
## Anacapa Island_Qtom           0.063 (-0.323, 0.449)      0.079 (-0.095, 0.252)
## Catalina Island_hybrid         -0.1 (-0.409, 0.209)      0.047 (-0.045, 0.139)
## Catalina Island_Qchr         -0.037 (-0.421, 0.346)      0.065 (-0.118, 0.248)
## San Clemente Island_hybrid   -0.112 (-0.417, 0.194)      0.044 (-0.046, 0.133)
## San Clemente Island_Qchr                       <NA>     -0.044 (-0.389, 0.302)
## Guadalupe Island_Guadalupe   -0.044 (-0.389, 0.302)                       <NA>
## Guadalupe Island_hybrid      -0.045 (-0.513, 0.424)     -0.014 (-0.244, 0.217)
##                            Guadalupe Island_hybrid average average_islands
## Mainland_Qchr               -0.019 (-0.266, 0.229)   0.021              NA
## Santa Rosa Island_Qtom       0.034 (-0.259, 0.327)   0.029           0.025
## Santa Cruz Island_Qtom        0.021 (-0.25, 0.291)   0.019           0.016
## Santa Cruz Island_hybrid    -0.015 (-0.261, 0.232)   0.006           0.003
## Anacapa Island_Qtom          0.094 (-0.185, 0.372)   0.051           0.050
## Catalina Island_hybrid      -0.012 (-0.259, 0.235)   0.008           0.005
## Catalina Island_Qchr          0.016 (-0.25, 0.281)   0.028           0.029
## San Clemente Island_hybrid   -0.02 (-0.264, 0.224)   0.003           0.000
## San Clemente Island_Qchr    -0.045 (-0.513, 0.424)  -0.059          -0.055
## Guadalupe Island_Guadalupe  -0.014 (-0.244, 0.217)   0.040           0.038
## Guadalupe Island_hybrid                       <NA>   0.004           0.007
```

```
#write.csv(fst.text, file = 'results/popgen_stats/Fst_island_ancestry_table.csv')
```

```
# heatmap

#png(file = 'results/popgen_stats/Fst_heatmap_fast-wcfst_ancestry.png', width = 10, height = 10, res = 300, units = 'in')
par(mar = c(5,4,4,10))
Heatmap(as.matrix(fst.df),
        cluster_rows = F, cluster_columns = F,
        row_names_side = 'left',
        column_names_side = 'top',
        column_names_gp = gpar(fontsize = 14),
        row_names_gp = gpar(fontsize = 14),
        column_names_rot = 90,
        col = brewer.pal(9, 'YlOrRd'),
        show_heatmap_legend = F,
        cell_fun = function(j, i, x, y, width, height, fill) {
          grid.text(fst.df.round[i,j], x, y, gp = gpar(fontsize = 14))
        })
```

```
#dev.off()

# weighted

#png(file = 'results/popgen_stats/Fst_weighted_heatmap_fast-wcfst_ancestry.png', width = 10, height = 10, res = 300, units = 'in')
par(mar = c(5,4,4,10))
Heatmap(as.matrix(fst.weight),
        cluster_rows = F, cluster_columns = F,
        row_names_side = 'left',
        column_names_side = 'top',
        column_names_gp = gpar(fontsize = 14),
        row_names_gp = gpar(fontsize = 14),
        column_names_rot = 90,
        col = brewer.pal(9, 'YlOrRd'),
        show_heatmap_legend = F,
        cell_fun = function(j, i, x, y, width, height, fill) {
          grid.text(fst.weight.round[i,j], x, y, gp = gpar(fontsize = 14))
        })
```

```
#dev.off()


# variance

#png(file = 'results/popgen_stats/Fst_variance_heatmap_fast-wcfst_ancestry.png', width = 10, height = 10, res = 300, units = 'in')
par(mar = c(5,4,4,10))
Heatmap(as.matrix(fst.var),
        cluster_rows = F, cluster_columns = F,
        row_names_side = 'left',
        column_names_side = 'top',
        column_names_gp = gpar(fontsize = 14),
        row_names_gp = gpar(fontsize = 14),
        column_names_rot = 90,
        col = brewer.pal(9, 'YlOrRd'),
        show_heatmap_legend = F,
        cell_fun = function(j, i, x, y, width, height, fill) {
          grid.text(fst.var.round[i,j], x, y, gp = gpar(fontsize = 14))
        })
```

```
#dev.off()
```
