## Supplementary Rmarkdown Outputs for "Comparison of conservation strategies for California Channel Island Oak (*Quercus tomentella*) using climate suitability predicted from genomic data": redundancy_analysis_SNPs.html


### Redundancy analysis

#### 2024-05-16

- 1 Setup
  - 1.1 load packages and data
  - 1.2 SNP data setup
  - 1.3 sample data setup
  - 1.4 impute SNP data
- 2 Simple PCA
  - 2.1 plot functions
  - 2.2 All samples
  - 2.3 without Guadalupe
- 3 RDA with climate
  - 3.1 Basic RDA with all individuals and
    climate variables to visualize
  - 3.2 Basic RDA, visualize without
    Guadalupe
  - 3.3 Basic RDA, visualize without
    Guadalupe or Mainland Qchr
  - 3.4 Basic RDA, visualize with only
    Guadalupe
  - 3.5 Reduce Climate Variables
- 4 Variance partitioning - all
  individuals
- 5 GEA analysis, all Q. tomentella
  - 5.1 RDA of SNPs and climate
  - 5.2 identify candidate SNPs from RDA
    outliers
- 6 GEA analysis, without Guadalupe or
  Mainaland Qchr
  - 6.1 RDA of SNPs and climate
  - 6.2 Variance partitioning - only
    Channel Islands samples
  - 6.3 identify candidate SNPs from RDA
    outliers
- 7 GEA analysis, only Guadalupe
  - 7.1 RDA of SNPs and climate
    - 7.1.1 identify candidate SNPs from RDA
      outliers
- 8 GEA analysis without Guadalupe,
  Qchr, or Anacapa
  - 8.1 RDA of SNPs and climate
  - 8.2 identify candidate SNPs from RDA
    outliers

Redundancy analysis with vegan to identify candidate
climate-associated SNPs

### 1 Setup

#### 1.1 load packages and data

```
library('vegan') # redundancy analysis
```

```
## Loading required package: permute
```

```
## Loading required package: lattice
```

```
## This is vegan 2.6-4
```

```
library('adegenet') # read.PLINK()
```

```
## Loading required package: ade4
```

```
## 
##    /// adegenet 2.1.10 is loaded ////////////
## 
##    > overview: '?adegenet'
##    > tutorials/doc/questions: 'adegenetWeb()' 
##    > bug reports/feature requests: adegenetIssues()
```

```
library('psych') # pairs.panels()

# going to follow this for a quick analysis!
#https://popgen.nescent.org/2018-03-27_RDA_GEA.html

# site data - named dat, needs to be renamed
load('data/clean/CCGP_samples_channel_islands_clean.rda')
info <- dat

# use this to read SNPs!
dat <- read.PLINK('data/raw/Qtom107.Qchr17.Qssp3.20220906.qlob.ef.repeatsOut.renamedChrsVars.biallelicSNPs.meanDP5.genoDP5.MAF0.01.missing0.9.ldPruned.additive.raw', n.cores = 2, quiet = F)
```

```
## 
##  Reading PLINK raw format into a genlight object... 
## 
## 
##  Reading loci information... 
## 
##  Reading and converting genotypes... 
## .
##  Building final object... 
## 
## ...done.
```

```
# climate data
load('/home/alayna/Documents/research/projects/2022_island_oak/data/clean/bioclim_locality_dataframe.rda')
clim <- bclim.loc
rm(bclim.loc)


sessionInfo()
```

```
## R version 4.3.3 (2024-02-29)
## Platform: x86_64-pc-linux-gnu (64-bit)
## Running under: Arch Linux
## 
## Matrix products: default
## BLAS:   /usr/lib/libblas.so.3.12.0 
## LAPACK: /usr/lib/liblapack.so.3.12.0
## 
## locale:
##  [1] LC_CTYPE=en_US.UTF-8       LC_NUMERIC=C              
##  [3] LC_TIME=en_US.UTF-8        LC_COLLATE=en_US.UTF-8    
##  [5] LC_MONETARY=en_US.UTF-8    LC_MESSAGES=en_US.UTF-8   
##  [7] LC_PAPER=en_US.UTF-8       LC_NAME=C                 
##  [9] LC_ADDRESS=C               LC_TELEPHONE=C            
## [11] LC_MEASUREMENT=en_US.UTF-8 LC_IDENTIFICATION=C       
## 
## time zone: America/New_York
## tzcode source: system (glibc)
## 
## attached base packages:
## [1] stats     graphics  grDevices utils     datasets  methods   base     
## 
## other attached packages:
## [1] psych_2.3.6     adegenet_2.1.10 ade4_1.7-22     vegan_2.6-4    
## [5] lattice_0.22-5  permute_0.9-7  
## 
## loaded via a namespace (and not attached):
##  [1] sass_0.4.5       utf8_1.2.3       generics_0.1.3   stringi_1.8.3   
##  [5] digest_0.6.31    magrittr_2.0.3   evaluate_0.20    grid_4.3.3      
##  [9] fastmap_1.1.1    seqinr_4.2-30    plyr_1.8.8       jsonlite_1.8.4  
## [13] Matrix_1.6-5     ape_5.7-1        promises_1.2.0.1 mgcv_1.9-1      
## [17] fansi_1.0.4      scales_1.2.1     jquerylib_0.1.4  mnormt_2.1.1    
## [21] cli_3.6.1        shiny_1.7.4.1    rlang_1.1.0      ellipsis_0.3.2  
## [25] munsell_0.5.0    splines_4.3.3    cachem_1.0.7     yaml_2.3.7      
## [29] tools_4.3.3      parallel_4.3.3   reshape2_1.4.4   dplyr_1.1.2     
## [33] colorspace_2.1-0 ggplot2_3.4.2    httpuv_1.6.11    mime_0.12       
## [37] vctrs_0.6.2      R6_2.5.1         lifecycle_1.0.3  stringr_1.5.0   
## [41] MASS_7.3-60.0.1  cluster_2.1.6    pkgconfig_2.0.3  later_1.3.1     
## [45] pillar_1.9.0     bslib_0.4.2      gtable_0.3.3     glue_1.6.2      
## [49] Rcpp_1.0.10      xfun_0.39        tibble_3.2.1     tidyselect_1.2.0
## [53] rstudioapi_0.14  knitr_1.42       xtable_1.8-4     htmltools_0.5.5 
## [57] nlme_3.1-164     igraph_1.5.0     rmarkdown_2.21   compiler_4.3.3
```

```
knitr::opts_chunk$set(fig.width = 10, fig.height = 8)
```

#### 1.2 SNP data setup

```
# look at dataset
# it omits the non-SNP info
dat
```

```
##  /// GENLIGHT OBJECT /////////
## 
##  // 127 genotypes,  585,298 binary SNPs, size: 80.1 Mb
##  3736056 (5.03 %) missing data
## 
##  // Basic content
##    @gen: list of 127 SNPbin
##    @ploidy: ploidy of each individual  (range: 2-2)
## 
##  // Optional content
##    @ind.names:  127 individual labels
##    @loc.names:  585298 locus labels
##    @pop: population of each individual (group size range: 1-1)
##    @other: a list containing: sex  phenotype  pat  mat
```

```
dim(dat)
```

```
## [1]    127 585298
```

```
snps <-  as.matrix(dat)
snps[1:10, 1:10]
```

```
##               1_39789_A_G_G 1_52176_G_A_A 1_52195_G_A_A 1_52213_C_T_T
## Qchr.A.232                1             0             0             1
## Qchr.A.275               NA             0             0             0
## Qchr.A.ED.102             1             0             0             0
## Qchr.A.ELD.53             1             0             0             1
## Qchr.A.KER.16             1             0             0             0
## Qchr.A.LA.02              1             0             0             0
## Qchr.A.LA.215            NA             0             0             1
## Qchr.A.LA.222             1             0             0             0
## Qchr.A.LA.309             0             0             0             0
## Qchr.A.PLU.65            NA             0             0             0
##               1_52248_T_C_T 1_52331_G_T_T 1_53841_G_A_A 1_53940_T_C_T
## Qchr.A.232                0             0             0             1
## Qchr.A.275                0             0             0             1
## Qchr.A.ED.102             0             0             1             0
## Qchr.A.ELD.53             0            NA             0             1
## Qchr.A.KER.16             0             1             0             1
## Qchr.A.LA.02              0             0             0             1
## Qchr.A.LA.215             0             0             0             1
## Qchr.A.LA.222             0             0             0             1
## Qchr.A.LA.309             0             0             0             1
## Qchr.A.PLU.65             0             0             0             1
##               1_53944_A_G_A 1_53966_G_A_A
## Qchr.A.232                0             0
## Qchr.A.275                0             0
## Qchr.A.ED.102             0             0
## Qchr.A.ELD.53             0             0
## Qchr.A.KER.16             0             0
## Qchr.A.LA.02              1             0
## Qchr.A.LA.215             0             0
## Qchr.A.LA.222             0             0
## Qchr.A.LA.309             1             0
## Qchr.A.PLU.65             1             0
```

```
dim(snps)
```

```
## [1]    127 585298
```

```
# optional - get a subset of SNPs for faster analysis, then run on whole dataset
# snps <- snps[, sample(1:ncol(snps), 10000, replace = F)]
```

#### 1.3 sample data setup

```
# get info only for the ones we have SNP data for

# first check for any mismatches
rownames(snps)[! rownames(snps) %in% info$ID_vcf]
```

```
## character(0)
```

```
# get subset
info <- info[match(rownames(snps), info$ID_vcf),]

# check
cbind(rownames(snps), info$ID_vcf, rownames(info))
```

```
##        [,1]             [,2]             [,3]             
##   [1,] "Qchr.A.232"     "Qchr.A.232"     "Qchr.A.232"     
##   [2,] "Qchr.A.275"     "Qchr.A.275"     "Qchr.A.275"     
##   [3,] "Qchr.A.ED.102"  "Qchr.A.ED.102"  "Qchr.A.ED.102"  
##   [4,] "Qchr.A.ELD.53"  "Qchr.A.ELD.53"  "Qchr.A.ELD.53"  
##   [5,] "Qchr.A.KER.16"  "Qchr.A.KER.16"  "Qchr.A.KER.16"  
##   [6,] "Qchr.A.LA.02"   "Qchr.A.LA.02"   "Qchr.A.LA.02"   
##   [7,] "Qchr.A.LA.215"  "Qchr.A.LA.215"  "Qchr.A.LA.215"  
##   [8,] "Qchr.A.LA.222"  "Qchr.A.LA.222"  "Qchr.A.LA.222"  
##   [9,] "Qchr.A.LA.309"  "Qchr.A.LA.309"  "Qchr.A.SIS.309" 
##  [10,] "Qchr.A.PLU.65"  "Qchr.A.PLU.65"  "Qchr.A.PLU.65"  
##  [11,] "Qchr.A.SBD.83"  "Qchr.A.SBD.83"  "Qchr.A.SBD.83"  
##  [12,] "Qchr.I.LA.20"   "Qchr.I.LA.20"   "Qchr.I.LA.20"   
##  [13,] "Qchr.I.LA.25"   "Qchr.I.LA.25"   "Qchr.I.LA.25"   
##  [14,] "Qchr.I.SB.02"   "Qchr.I.SB.02"   "Qchr.I.SB.02"   
##  [15,] "Qchr.I.SB.03"   "Qchr.I.SB.03"   "Qchr.I.SB.03"   
##  [16,] "Qchr.JJK.SB.13" "Qchr.JJK.SB.13" "Qchr.JJK.SB.13" 
##  [17,] "Qchr.JR.SB.01"  "Qchr.JR.SB.01"  "Qchr.JR.SB.01"  
##  [18,] "Qssp.I.LA.03"   "Qssp.I.LA.03"   "Qspp.I.LA.03"   
##  [19,] "Qssp.I.LA.04"   "Qssp.I.LA.04"   "Qspp.I.LA.04"   
##  [20,] "Qssp.I.LA.14"   "Qssp.I.LA.14"   "Qspp.I.LA.14"   
##  [21,] "Qtom.A.LA.169"  "Qtom.A.LA.169"  "Qtom.A.LA.169"  
##  [22,] "Qtom.A.LA.175"  "Qtom.A.LA.175"  "Qtom.A.LA.175"  
##  [23,] "Qtom.A.LA.176"  "Qtom.A.LA.176"  "Qtom.A.LA.176"  
##  [24,] "Qtom.A.LA.180"  "Qtom.A.LA.180"  "Qtom.A.LA.180"  
##  [25,] "Qtom.A.LA.181"  "Qtom.A.LA.181"  "Qtom.A.LA.181"  
##  [26,] "Qtom.A.LA.183"  "Qtom.A.LA.183"  "Qtom.A.LA.183"  
##  [27,] "Qtom.A.LA.190"  "Qtom.A.LA.190"  "Qtom.A.LA.190"  
##  [28,] "Qtom.A.LA.191"  "Qtom.A.LA.191"  "Qtom.A.LA.191"  
##  [29,] "Qtom.A.LA.198"  "Qtom.A.LA.198"  "Qtom.A.LA.198"  
##  [30,] "Qtom.A.LA.200"  "Qtom.A.LA.200"  "Qtom.A.LA.200"  
##  [31,] "Qtom.A.LA.204"  "Qtom.A.LA.204"  "Qtom.A.LA.204"  
##  [32,] "Qtom.A.LA.207"  "Qtom.A.LA.207"  "Qtom.A.LA.207"  
##  [33,] "Qtom.A.LA.208"  "Qtom.A.LA.208"  "Qtom.A.LA.208"  
##  [34,] "Qtom.A.LA.209"  "Qtom.A.LA.209"  "Qtom.A.LA.209"  
##  [35,] "Qtom.A.LA.218"  "Qtom.A.LA.218"  "Qtom.A.LA.218"  
##  [36,] "Qtom.A.LA.221"  "Qtom.A.LA.221"  "Qtom.A.LA.221"  
##  [37,] "Qtom.A.SB.324"  "Qtom.A.SB.324"  "Qtom.A.SB.324"  
##  [38,] "Qtom.A.SB.325"  "Qtom.A.SB.325"  "Qtom.A.SB.325"  
##  [39,] "Qtom.A.SB.327"  "Qtom.A.SB.327"  "Qtom.A.SB.327"  
##  [40,] "Qtom.A.SB.329"  "Qtom.A.SB.329"  "Qtom.A.SB.329"  
##  [41,] "Qtom.A.SB.330"  "Qtom.A.SB.330"  "Qtom.A.SB.330"  
##  [42,] "Qtom.A.SB.331"  "Qtom.A.SB.331"  "Qtom.A.SB.331"  
##  [43,] "Qtom.A.SB.332"  "Qtom.A.SB.332"  "Qtom.A.SB.332"  
##  [44,] "Qtom.A.SB.333"  "Qtom.A.SB.333"  "Qtom.A.SB.333"  
##  [45,] "Qtom.A.SB.335"  "Qtom.A.SB.335"  "Qtom.A.SB.335"  
##  [46,] "Qtom.A.SB.336"  "Qtom.A.SB.336"  "Qtom.A.SB.336"  
##  [47,] "Qtom.A.SB.337"  "Qtom.A.SB.337"  "Qtom.A.SB.337"  
##  [48,] "Qtom.A.SB.338"  "Qtom.A.SB.338"  "Qtom.A.SB.338"  
##  [49,] "Qtom.A.SB.339"  "Qtom.A.SB.339"  "Qtom.A.SB.339"  
##  [50,] "Qtom.A.SB.341"  "Qtom.A.SB.341"  "Qtom.A.SB.341"  
##  [51,] "Qtom.A.SB.342"  "Qtom.A.SB.342"  "Qtom.A.SB.342"  
##  [52,] "Qtom.A.SB.343"  "Qtom.A.SB.343"  "Qtom.A.SB.343"  
##  [53,] "Qtom.A.SB.344"  "Qtom.A.SB.344"  "Qtom.A.SB.344"  
##  [54,] "Qtom.A.SB.345"  "Qtom.A.SB.345"  "Qtom.A.SB.345"  
##  [55,] "Qtom.A.SB.346"  "Qtom.A.SB.346"  "Qtom.A.SB.346"  
##  [56,] "Qtom.A.SB.347"  "Qtom.A.SB.347"  "Qtom.A.SB.347"  
##  [57,] "Qtom.A.SB.348"  "Qtom.A.SB.348"  "Qtom.A.SB.348"  
##  [58,] "Qtom.A.SB.352"  "Qtom.A.SB.352"  "Qtom.A.SB.352"  
##  [59,] "Qtom.A.SB.353"  "Qtom.A.SB.353"  "Qtom.A.SB.353"  
##  [60,] "Qtom.A.SB.355"  "Qtom.A.SB.355"  "Qtom.A.SB.355"  
##  [61,] "Qtom.I.BC.10"   "Qtom.I.BC.10"   "Qtom.I.BC.10"   
##  [62,] "Qtom.I.BC.11"   "Qtom.I.BC.11"   "Qtom.I.BC.11"   
##  [63,] "Qtom.I.BC.12"   "Qtom.I.BC.12"   "Qtom.I.BC.12"   
##  [64,] "Qtom.I.BC.13"   "Qtom.I.BC.13"   "Qtom.I.BC.13"   
##  [65,] "Qtom.I.BC.18"   "Qtom.I.BC.18"   "Qtom.I.BC.18"   
##  [66,] "Qtom.I.BC.20"   "Qtom.I.BC.20"   "Qtom.I.BC.20"   
##  [67,] "Qtom.I.BC.21"   "Qtom.I.BC.21"   "Qtom.I.BC.21"   
##  [68,] "Qtom.I.BC.23"   "Qtom.I.BC.23"   "Qtom.I.BC.23"   
##  [69,] "Qtom.I.BC.24"   "Qtom.I.BC.24"   "Qtom.I.BC.24"   
##  [70,] "Qtom.I.BC.32"   "Qtom.I.BC.32"   "Qtom.I.BC.32"   
##  [71,] "Qtom.I.BC.33"   "Qtom.I.BC.33"   "Qtom.I.BC.33"   
##  [72,] "Qtom.I.BC.35"   "Qtom.I.BC.35"   "Qtom.I.BC.35"   
##  [73,] "Qtom.I.BC.39"   "Qtom.I.BC.39"   "Qtom.I.BC.39"   
##  [74,] "Qtom.I.BC.4"    "Qtom.I.BC.4"    "Qtom.I.BC.4"    
##  [75,] "Qtom.I.BC.40"   "Qtom.I.BC.40"   "Qtom.I.BC.40"   
##  [76,] "Qtom.I.BC.41"   "Qtom.I.BC.41"   "Qtom.I.BC.41"   
##  [77,] "Qtom.I.BC.42"   "Qtom.I.BC.42"   "Qtom.I.BC.42"   
##  [78,] "Qtom.I.BC.43"   "Qtom.I.BC.43"   "Qtom.I.BC.43"   
##  [79,] "Qtom.I.BC.44"   "Qtom.I.BC.44"   "Qtom.I.BC.44"   
##  [80,] "Qtom.I.BC.45"   "Qtom.I.BC.45"   "Qtom.I.BC.45"   
##  [81,] "Qtom.I.BC.46"   "Qtom.I.BC.46"   "Qtom.I.BC.46"   
##  [82,] "Qtom.I.BC.48"   "Qtom.I.BC.48"   "Qtom.I.BC.48"   
##  [83,] "Qtom.I.LA.01b"  "Qtom.I.LA.01b"  "Qtom.I.LA.SC.01"
##  [84,] "Qtom.I.LA.02"   "Qtom.I.LA.02"   "Qtom.I.LA.02"   
##  [85,] "Qtom.I.LA.05"   "Qtom.I.LA.05"   "Qtom.I.LA.05"   
##  [86,] "Qtom.I.LA.06"   "Qtom.I.LA.06"   "Qtom.I.LA.06"   
##  [87,] "Qtom.I.LA.08"   "Qtom.I.LA.08"   "Qtom.I.LA.08"   
##  [88,] "Qtom.I.LA.10"   "Qtom.I.LA.10"   "Qtom.I.LA.10"   
##  [89,] "Qtom.I.LA.11"   "Qtom.I.LA.11"   "Qtom.I.LA.11"   
##  [90,] "Qtom.I.LA.12"   "Qtom.I.LA.12"   "Qtom.I.LA.12"   
##  [91,] "Qtom.I.LA.13"   "Qtom.I.LA.13"   "Qtom.I.LA.13"   
##  [92,] "Qtom.I.LA.21"   "Qtom.I.LA.21"   "Qtom.I.LA.21"   
##  [93,] "Qtom.I.LA.22"   "Qtom.I.LA.22"   "Qtom.I.LA.22"   
##  [94,] "Qtom.I.LA.23"   "Qtom.I.LA.23"   "Qtom.I.LA.23"   
##  [95,] "Qtom.I.LA.24"   "Qtom.I.LA.24"   "Qtom.I.LA.24"   
##  [96,] "Qtom.I.SB.01"   "Qtom.I.SB.01"   "Qtom.I.SB.01"   
##  [97,] "Qtom.I.SB.04"   "Qtom.I.SB.04"   "Qtom.I.SB.04"   
##  [98,] "Qtom.I.SB.09"   "Qtom.I.SB.09"   "Qtom.I.SB.09"   
##  [99,] "Qtom.I.SB.10"   "Qtom.I.SB.10"   "Qtom.I.SB.10"   
## [100,] "Qtom.I.SB.11"   "Qtom.I.SB.11"   "Qtom.I.SB.11"   
## [101,] "Qtom.I.SB.12"   "Qtom.I.SB.12"   "Qtom.I.SB.12"   
## [102,] "Qtom.I.SB.28"   "Qtom.I.SB.28"   "Qtom.I.SB.28"   
## [103,] "Qtom.I.SB.33"   "Qtom.I.SB.33"   "Qtom.I.SB.33"   
## [104,] "Qtom.I.SB.35"   "Qtom.I.SB.35"   "Qtom.I.SB.35"   
## [105,] "Qtom.I.SB.36"   "Qtom.I.SB.36"   "Qtom.I.SB.36"   
## [106,] "Qtom.I.SB.37"   "Qtom.I.SB.37"   "Qtom.I.SB.37"   
## [107,] "Qtom.I.SB.38"   "Qtom.I.SB.38"   "Qtom.I.SB.38"   
## [108,] "Qtom.I.SB.40"   "Qtom.I.SB.40"   "Qtom.I.SB.40"   
## [109,] "Qtom.I.SB.41"   "Qtom.I.SB.41"   "Qtom.I.SB.41"   
## [110,] "Qtom.I.SB.42"   "Qtom.I.SB.42"   "Qtom.I.SB.42"   
## [111,] "Qtom.I.SB.44"   "Qtom.I.SB.44"   "Qtom.I.SB.44"   
## [112,] "Qtom.I.SB.45"   "Qtom.I.SB.45"   "Qtom.I.SB.45"   
## [113,] "Qtom.I.SB.73"   "Qtom.I.SB.73"   "Qtom.I.SB.73"   
## [114,] "Qtom.I.SB.74"   "Qtom.I.SB.74"   "Qtom.I.SB.74"   
## [115,] "Qtom.I.VEN.01"  "Qtom.I.VEN.01"  "Qtom.I.VEN.01"  
## [116,] "Qtom.I.VEN.02"  "Qtom.I.VEN.02"  "Qtom.I.VEN.02"  
## [117,] "Qtom.I.VEN.03"  "Qtom.I.VEN.03"  "Qtom.I.VEN.03"  
## [118,] "Qtom.I.VEN.04"  "Qtom.I.VEN.04"  "Qtom.I.VEN.04"  
## [119,] "Qtom.I.VEN.05"  "Qtom.I.VEN.05"  "Qtom.I.VEN.05"  
## [120,] "Qtom.JJK.SB.01" "Qtom.JJK.SB.01" "Qtom.JJK.SB.01" 
## [121,] "Qtom.JJK.SB.03" "Qtom.JJK.SB.03" "Qtom.JJK.SB.03" 
## [122,] "Qtom.JJK.SB.07" "Qtom.JJK.SB.07" "Qtom.JJK.SB.07" 
## [123,] "Qtom.JJK.SB.08" "Qtom.JJK.SB.08" "Qtom.JJK.SB.08" 
## [124,] "Qtom.JJK.SB.09" "Qtom.JJK.SB.09" "Qtom.JJK.SB.09" 
## [125,] "Qtom.JJK.SB.10" "Qtom.JJK.SB.10" "Qtom.JJK.SB.10" 
## [126,] "Qtom.JR.SB.01"  "Qtom.JR.SB.01"  "Qtom.JR.SB.01"  
## [127,] "Qtom.S.LA.186"  "Qtom.S.LA.186"  "Qtom.S.LA.186"
```

```
# rename SNP rows to match corrected names (info rownames)
rownames(snps) <- rownames(info)

# also get species abbreviation from name
info$sp <- sapply(1:nrow(snps), function(x) strsplit(rownames(snps)[x], split = '.', fixed = T)[[1]][1])
info$sp <- factor(info$sp)
```

#### 1.4 impute SNP data

```
# need to impute missing data

# proportion missing data - about 5%
sum(is.na(snps))/length(unlist(snps))
```

```
## [1] 0.05026117
```

```
# missing data by individual
sapply(1:nrow(snps), function(x) sum(is.na(snps[x,])))
```

```
##   [1]  20983  44944  46343  21462  94170  33554 102082  31387  26942  14045
##  [11]  67790  15550  18109   2612   3158   6849  44727  32195   7037   3407
##  [21]  56524  13715  32344  57563  39197  74444  85828  45213  14671   3409
##  [31]  12135   6741    816  26804  33685  49597 106186  17891   9112  19991
##  [41]  21032  50060  11536  13195   4364   5811   2167   7595   2067  99840
##  [51]  10712   5847  23971  60724  42372   1660   6708   8761   7872   3889
##  [61]  62652   5174  16710  22374  52223 101481  30350   9754    798  12526
##  [71]  47669  12144 130814   7709  15040  58472 118808   2268  18733   8360
##  [81]  17513  43143  12901  54424  56736   8854   7944 101447   1050   1587
##  [91]  10494   9737 111125  19100   2201   1313   2940  63956    928   2070
## [101]   4639   4466  14191  11829   3974   2585  12137  51742  20337   1500
## [111]   4086   2393  44002  21655   8442   7386  13220   4319   8398  20256
## [121] 159940  37917  50023 146983  38691  23885  20143
```

```
miss <- sapply(1:nrow(snps), function(x) sum(is.na(snps[x,]))/ncol(snps))
missdf <- cbind(rownames(snps), miss)
missdf[order(missdf[,2]),]
```

```
##                          miss                 
##   [1,] "Qtom.I.BC.24"    "0.0013634080417155" 
##   [2,] "Qtom.A.LA.208"   "0.00139416160656623"
##   [3,] "Qtom.I.SB.10"    "0.00158551712119296"
##   [4,] "Qtom.I.LA.11"    "0.00179395794962566"
##   [5,] "Qtom.I.SB.01"    "0.00224330170272237"
##   [6,] "Qtom.I.SB.42"    "0.0025627970708938" 
##   [7,] "Qtom.I.LA.12"    "0.00271143930100564"
##   [8,] "Qtom.A.SB.347"   "0.00283616209178914"
##   [9,] "Qtom.A.SB.339"   "0.00353153436369166"
##  [10,] "Qtom.I.SB.11"    "0.00353665995783345"
##  [11,] "Qtom.A.SB.337"   "0.00370238750175124"
##  [12,] "Qtom.I.LA.24"    "0.0037604775686915" 
##  [13,] "Qtom.I.BC.43"    "0.00387494917119143"
##  [14,] "Qtom.I.SB.45"    "0.00408851559376591"
##  [15,] "Qtom.I.SB.37"    "0.00441655361884032"
##  [16,] "Qchr.I.SB.02"    "0.00446268396611641"
##  [17,] "Qtom.I.SB.04"    "0.00502308225895185"
##  [18,] "Qchr.I.SB.03"    "0.00539554209992175"
##  [19,] "Qspp.I.LA.14"    "0.00582096641369012"
##  [20,] "Qtom.A.LA.200"   "0.00582438347645131"
##  [21,] "Qtom.A.SB.355"   "0.00664447853913733"
##  [22,] "Qtom.I.SB.36"    "0.00678970370648798"
##  [23,] "Qtom.I.SB.44"    "0.00698105922111471"
##  [24,] "Qtom.I.VEN.04"   "0.00737914703279355"
##  [25,] "Qtom.A.SB.335"   "0.00745603094492037"
##  [26,] "Qtom.I.SB.28"    "0.00763030114574114"
##  [27,] "Qtom.I.SB.12"    "0.00792587707458423"
##  [28,] "Qtom.I.BC.11"    "0.00883994136320302"
##  [29,] "Qtom.A.SB.336"   "0.00992827585264259"
##  [30,] "Qtom.A.SB.343"   "0.00998978298234404"
##  [31,] "Qtom.A.SB.348"   "0.0114608285010371" 
##  [32,] "Qtom.A.LA.207"   "0.0115172100365967" 
##  [33,] "Qchr.JJK.SB.13"  "0.0117017314257011" 
##  [34,] "Qspp.I.LA.04"    "0.0120229353252531" 
##  [35,] "Qtom.I.VEN.02"   "0.0126192127770811" 
##  [36,] "Qtom.A.SB.338"   "0.0129762958356256" 
##  [37,] "Qtom.I.BC.4"     "0.0131710684130135" 
##  [38,] "Qtom.A.SB.353"   "0.0134495590280507" 
##  [39,] "Qtom.I.LA.08"    "0.0135725732874536" 
##  [40,] "Qtom.I.BC.45"    "0.0142833223417815" 
##  [41,] "Qtom.I.VEN.05"   "0.0143482465342441" 
##  [42,] "Qtom.I.VEN.01"   "0.0144234219149903" 
##  [43,] "Qtom.A.SB.352"   "0.0149684434254004" 
##  [44,] "Qtom.I.LA.06"    "0.0151273368437958" 
##  [45,] "Qtom.A.SB.327"   "0.0155681379399895" 
##  [46,] "Qtom.I.LA.21"    "0.016635970052862"  
##  [47,] "Qtom.I.BC.23"    "0.0166650150863321" 
##  [48,] "Qtom.I.LA.13"    "0.017929328307973"  
##  [49,] "Qtom.A.SB.342"   "0.0183017881489429" 
##  [50,] "Qtom.A.SB.332"   "0.0197096180065539" 
##  [51,] "Qtom.I.SB.35"    "0.0202102177010685" 
##  [52,] "Qtom.A.LA.204"   "0.0207330283035309" 
##  [53,] "Qtom.I.SB.38"    "0.020736445366292"  
##  [54,] "Qtom.I.BC.35"    "0.0207484050859562" 
##  [55,] "Qtom.I.BC.32"    "0.0214010640733438" 
##  [56,] "Qtom.I.LA.SC.01" "0.0220417633410673" 
##  [57,] "Qtom.A.SB.333"   "0.0225440715669625" 
##  [58,] "Qtom.I.VEN.03"   "0.0225867848514774" 
##  [59,] "Qtom.A.LA.175"   "0.0234325078848723" 
##  [60,] "Qchr.A.PLU.65"   "0.023996323240469"  
##  [61,] "Qtom.I.SB.33"    "0.024245768822036"  
##  [62,] "Qtom.A.LA.198"   "0.025065863884722"  
##  [63,] "Qtom.I.BC.40"    "0.0256963119641618" 
##  [64,] "Qchr.I.LA.20"    "0.0265676629682657" 
##  [65,] "Qtom.I.BC.12"    "0.0285495593697569" 
##  [66,] "Qtom.I.BC.46"    "0.0299215100683754" 
##  [67,] "Qtom.A.SB.325"   "0.0305673349302407" 
##  [68,] "Qchr.I.LA.25"    "0.0309397947712106" 
##  [69,] "Qtom.I.BC.44"    "0.0320059183527024" 
##  [70,] "Qtom.I.LA.23"    "0.0326329493693811" 
##  [71,] "Qtom.A.SB.329"   "0.034155250829492"  
##  [72,] "Qtom.S.LA.186"   "0.0344149475993426" 
##  [73,] "Qtom.JJK.SB.01"  "0.0346080116453499" 
##  [74,] "Qtom.I.SB.41"    "0.0347464026871782" 
##  [75,] "Qchr.A.232"      "0.0358501139590431" 
##  [76,] "Qtom.A.SB.330"   "0.0359338319966923" 
##  [77,] "Qchr.A.ELD.53"   "0.0366685004903485" 
##  [78,] "Qtom.I.SB.74"    "0.0369982470468035" 
##  [79,] "Qtom.I.BC.13"    "0.0382266811094519" 
##  [80,] "Qtom.JR.SB.01"   "0.0408082720255323" 
##  [81,] "Qtom.A.SB.344"   "0.0409552057242635" 
##  [82,] "Qtom.A.LA.209"   "0.0457954751254916" 
##  [83,] "Qchr.A.SIS.309"  "0.0460312524560139" 
##  [84,] "Qtom.I.BC.21"    "0.0518539274010846" 
##  [85,] "Qchr.A.LA.222"   "0.0536256744427625" 
##  [86,] "Qspp.I.LA.03"    "0.055006167798284"  
##  [87,] "Qtom.A.LA.176"   "0.0552607389739927" 
##  [88,] "Qchr.A.LA.02"    "0.0573280619445137" 
##  [89,] "Qtom.A.LA.218"   "0.0575518795553718" 
##  [90,] "Qtom.JJK.SB.07"  "0.0647823843580535" 
##  [91,] "Qtom.JJK.SB.10"  "0.0661047876466347" 
##  [92,] "Qtom.A.LA.181"   "0.0669693045252162" 
##  [93,] "Qtom.A.SB.346"   "0.0723938916586081" 
##  [94,] "Qtom.I.BC.48"    "0.0737111693530475" 
##  [95,] "Qtom.I.SB.73"    "0.0751787978089794" 
##  [96,] "Qchr.JR.SB.01"   "0.0764174830599114" 
##  [97,] "Qchr.A.275"      "0.0767882343695007" 
##  [98,] "Qtom.A.LA.191"   "0.0772478293108809" 
##  [99,] "Qchr.A.ED.102"   "0.0791784697709543" 
## [100,] "Qtom.I.BC.33"    "0.0814439823816244" 
## [101,] "Qtom.A.LA.221"   "0.0847380308834132" 
## [102,] "Qtom.JJK.SB.08"  "0.0854658652515471" 
## [103,] "Qtom.A.SB.331"   "0.0855290809126291" 
## [104,] "Qtom.I.SB.40"    "0.0884028306947914" 
## [105,] "Qtom.I.BC.18"    "0.089224634288858"  
## [106,] "Qtom.I.LA.02"    "0.0929851118575495" 
## [107,] "Qtom.A.LA.169"   "0.0965730277568008" 
## [108,] "Qtom.I.LA.05"    "0.0969352364094871" 
## [109,] "Qtom.A.LA.180"   "0.0983481918612399" 
## [110,] "Qtom.I.BC.41"    "0.0999012468862016" 
## [111,] "Qtom.A.SB.345"   "0.103748859555303"  
## [112,] "Qtom.I.BC.10"    "0.107042908057092"  
## [113,] "Qtom.I.SB.09"    "0.109270832977389"  
## [114,] "Qchr.A.SBD.83"   "0.115821342290594"  
## [115,] "Qtom.A.LA.183"   "0.127189910097079"  
## [116,] "Qtom.A.LA.190"   "0.146639831333782"  
## [117,] "Qchr.A.KER.16"   "0.160892400110713"  
## [118,] "Qtom.A.SB.341"   "0.170579773038691"  
## [119,] "Qtom.I.LA.10"    "0.173325382967309"  
## [120,] "Qtom.I.BC.20"    "0.173383473034249"  
## [121,] "Qchr.A.LA.215"   "0.174410300393987"  
## [122,] "Qtom.A.SB.324"   "0.181422113179953"  
## [123,] "Qtom.I.LA.22"    "0.189860549668716"  
## [124,] "Qtom.I.BC.42"    "0.202987196265834"  
## [125,] "Qtom.I.BC.39"    "0.223499824021268"  
## [126,] "Qtom.JJK.SB.09"  "0.251125067914122"  
## [127,] "Qtom.JJK.SB.03"  "0.273262509012503"
```

```
# impute by most common snp
imp <- apply(snps, 2, function(x) replace(x, is.na(x), as.numeric(names(which.max(table(x))))))
sum(is.na(imp))
```

```
## [1] 0
```

```
#save(imp, file = 'data/clean/Qtom107.Qchr17.Qssp3.20220906.qlob.ef.repeatsOut.renamedChrsVars.biallelicSNPs.meanDP5.genoDP5.MAF0.01.missing0.9.ldPruned.additive_imputed.rda')
#write.csv(imp, file = 'data/clean/Qtom107.Qchr17.Qssp3.20220906.qlob.ef.repeatsOut.renamedChrsVars.biallelicSNPs.meanDP5.genoDP5.MAF0.01.missing0.9.ldPruned.additive_imputed.csv', quote = F)
```

```
# one last check that rows are in the same order...
cbind(rownames(info), rownames(snps), rownames(imp))
```

```
##        [,1]              [,2]              [,3]             
##   [1,] "Qchr.A.232"      "Qchr.A.232"      "Qchr.A.232"     
##   [2,] "Qchr.A.275"      "Qchr.A.275"      "Qchr.A.275"     
##   [3,] "Qchr.A.ED.102"   "Qchr.A.ED.102"   "Qchr.A.ED.102"  
##   [4,] "Qchr.A.ELD.53"   "Qchr.A.ELD.53"   "Qchr.A.ELD.53"  
##   [5,] "Qchr.A.KER.16"   "Qchr.A.KER.16"   "Qchr.A.KER.16"  
##   [6,] "Qchr.A.LA.02"    "Qchr.A.LA.02"    "Qchr.A.LA.02"   
##   [7,] "Qchr.A.LA.215"   "Qchr.A.LA.215"   "Qchr.A.LA.215"  
##   [8,] "Qchr.A.LA.222"   "Qchr.A.LA.222"   "Qchr.A.LA.222"  
##   [9,] "Qchr.A.SIS.309"  "Qchr.A.SIS.309"  "Qchr.A.SIS.309" 
##  [10,] "Qchr.A.PLU.65"   "Qchr.A.PLU.65"   "Qchr.A.PLU.65"  
##  [11,] "Qchr.A.SBD.83"   "Qchr.A.SBD.83"   "Qchr.A.SBD.83"  
##  [12,] "Qchr.I.LA.20"    "Qchr.I.LA.20"    "Qchr.I.LA.20"   
##  [13,] "Qchr.I.LA.25"    "Qchr.I.LA.25"    "Qchr.I.LA.25"   
##  [14,] "Qchr.I.SB.02"    "Qchr.I.SB.02"    "Qchr.I.SB.02"   
##  [15,] "Qchr.I.SB.03"    "Qchr.I.SB.03"    "Qchr.I.SB.03"   
##  [16,] "Qchr.JJK.SB.13"  "Qchr.JJK.SB.13"  "Qchr.JJK.SB.13" 
##  [17,] "Qchr.JR.SB.01"   "Qchr.JR.SB.01"   "Qchr.JR.SB.01"  
##  [18,] "Qspp.I.LA.03"    "Qspp.I.LA.03"    "Qspp.I.LA.03"   
##  [19,] "Qspp.I.LA.04"    "Qspp.I.LA.04"    "Qspp.I.LA.04"   
##  [20,] "Qspp.I.LA.14"    "Qspp.I.LA.14"    "Qspp.I.LA.14"   
##  [21,] "Qtom.A.LA.169"   "Qtom.A.LA.169"   "Qtom.A.LA.169"  
##  [22,] "Qtom.A.LA.175"   "Qtom.A.LA.175"   "Qtom.A.LA.175"  
##  [23,] "Qtom.A.LA.176"   "Qtom.A.LA.176"   "Qtom.A.LA.176"  
##  [24,] "Qtom.A.LA.180"   "Qtom.A.LA.180"   "Qtom.A.LA.180"  
##  [25,] "Qtom.A.LA.181"   "Qtom.A.LA.181"   "Qtom.A.LA.181"  
##  [26,] "Qtom.A.LA.183"   "Qtom.A.LA.183"   "Qtom.A.LA.183"  
##  [27,] "Qtom.A.LA.190"   "Qtom.A.LA.190"   "Qtom.A.LA.190"  
##  [28,] "Qtom.A.LA.191"   "Qtom.A.LA.191"   "Qtom.A.LA.191"  
##  [29,] "Qtom.A.LA.198"   "Qtom.A.LA.198"   "Qtom.A.LA.198"  
##  [30,] "Qtom.A.LA.200"   "Qtom.A.LA.200"   "Qtom.A.LA.200"  
##  [31,] "Qtom.A.LA.204"   "Qtom.A.LA.204"   "Qtom.A.LA.204"  
##  [32,] "Qtom.A.LA.207"   "Qtom.A.LA.207"   "Qtom.A.LA.207"  
##  [33,] "Qtom.A.LA.208"   "Qtom.A.LA.208"   "Qtom.A.LA.208"  
##  [34,] "Qtom.A.LA.209"   "Qtom.A.LA.209"   "Qtom.A.LA.209"  
##  [35,] "Qtom.A.LA.218"   "Qtom.A.LA.218"   "Qtom.A.LA.218"  
##  [36,] "Qtom.A.LA.221"   "Qtom.A.LA.221"   "Qtom.A.LA.221"  
##  [37,] "Qtom.A.SB.324"   "Qtom.A.SB.324"   "Qtom.A.SB.324"  
##  [38,] "Qtom.A.SB.325"   "Qtom.A.SB.325"   "Qtom.A.SB.325"  
##  [39,] "Qtom.A.SB.327"   "Qtom.A.SB.327"   "Qtom.A.SB.327"  
##  [40,] "Qtom.A.SB.329"   "Qtom.A.SB.329"   "Qtom.A.SB.329"  
##  [41,] "Qtom.A.SB.330"   "Qtom.A.SB.330"   "Qtom.A.SB.330"  
##  [42,] "Qtom.A.SB.331"   "Qtom.A.SB.331"   "Qtom.A.SB.331"  
##  [43,] "Qtom.A.SB.332"   "Qtom.A.SB.332"   "Qtom.A.SB.332"  
##  [44,] "Qtom.A.SB.333"   "Qtom.A.SB.333"   "Qtom.A.SB.333"  
##  [45,] "Qtom.A.SB.335"   "Qtom.A.SB.335"   "Qtom.A.SB.335"  
##  [46,] "Qtom.A.SB.336"   "Qtom.A.SB.336"   "Qtom.A.SB.336"  
##  [47,] "Qtom.A.SB.337"   "Qtom.A.SB.337"   "Qtom.A.SB.337"  
##  [48,] "Qtom.A.SB.338"   "Qtom.A.SB.338"   "Qtom.A.SB.338"  
##  [49,] "Qtom.A.SB.339"   "Qtom.A.SB.339"   "Qtom.A.SB.339"  
##  [50,] "Qtom.A.SB.341"   "Qtom.A.SB.341"   "Qtom.A.SB.341"  
##  [51,] "Qtom.A.SB.342"   "Qtom.A.SB.342"   "Qtom.A.SB.342"  
##  [52,] "Qtom.A.SB.343"   "Qtom.A.SB.343"   "Qtom.A.SB.343"  
##  [53,] "Qtom.A.SB.344"   "Qtom.A.SB.344"   "Qtom.A.SB.344"  
##  [54,] "Qtom.A.SB.345"   "Qtom.A.SB.345"   "Qtom.A.SB.345"  
##  [55,] "Qtom.A.SB.346"   "Qtom.A.SB.346"   "Qtom.A.SB.346"  
##  [56,] "Qtom.A.SB.347"   "Qtom.A.SB.347"   "Qtom.A.SB.347"  
##  [57,] "Qtom.A.SB.348"   "Qtom.A.SB.348"   "Qtom.A.SB.348"  
##  [58,] "Qtom.A.SB.352"   "Qtom.A.SB.352"   "Qtom.A.SB.352"  
##  [59,] "Qtom.A.SB.353"   "Qtom.A.SB.353"   "Qtom.A.SB.353"  
##  [60,] "Qtom.A.SB.355"   "Qtom.A.SB.355"   "Qtom.A.SB.355"  
##  [61,] "Qtom.I.BC.10"    "Qtom.I.BC.10"    "Qtom.I.BC.10"   
##  [62,] "Qtom.I.BC.11"    "Qtom.I.BC.11"    "Qtom.I.BC.11"   
##  [63,] "Qtom.I.BC.12"    "Qtom.I.BC.12"    "Qtom.I.BC.12"   
##  [64,] "Qtom.I.BC.13"    "Qtom.I.BC.13"    "Qtom.I.BC.13"   
##  [65,] "Qtom.I.BC.18"    "Qtom.I.BC.18"    "Qtom.I.BC.18"   
##  [66,] "Qtom.I.BC.20"    "Qtom.I.BC.20"    "Qtom.I.BC.20"   
##  [67,] "Qtom.I.BC.21"    "Qtom.I.BC.21"    "Qtom.I.BC.21"   
##  [68,] "Qtom.I.BC.23"    "Qtom.I.BC.23"    "Qtom.I.BC.23"   
##  [69,] "Qtom.I.BC.24"    "Qtom.I.BC.24"    "Qtom.I.BC.24"   
##  [70,] "Qtom.I.BC.32"    "Qtom.I.BC.32"    "Qtom.I.BC.32"   
##  [71,] "Qtom.I.BC.33"    "Qtom.I.BC.33"    "Qtom.I.BC.33"   
##  [72,] "Qtom.I.BC.35"    "Qtom.I.BC.35"    "Qtom.I.BC.35"   
##  [73,] "Qtom.I.BC.39"    "Qtom.I.BC.39"    "Qtom.I.BC.39"   
##  [74,] "Qtom.I.BC.4"     "Qtom.I.BC.4"     "Qtom.I.BC.4"    
##  [75,] "Qtom.I.BC.40"    "Qtom.I.BC.40"    "Qtom.I.BC.40"   
##  [76,] "Qtom.I.BC.41"    "Qtom.I.BC.41"    "Qtom.I.BC.41"   
##  [77,] "Qtom.I.BC.42"    "Qtom.I.BC.42"    "Qtom.I.BC.42"   
##  [78,] "Qtom.I.BC.43"    "Qtom.I.BC.43"    "Qtom.I.BC.43"   
##  [79,] "Qtom.I.BC.44"    "Qtom.I.BC.44"    "Qtom.I.BC.44"   
##  [80,] "Qtom.I.BC.45"    "Qtom.I.BC.45"    "Qtom.I.BC.45"   
##  [81,] "Qtom.I.BC.46"    "Qtom.I.BC.46"    "Qtom.I.BC.46"   
##  [82,] "Qtom.I.BC.48"    "Qtom.I.BC.48"    "Qtom.I.BC.48"   
##  [83,] "Qtom.I.LA.SC.01" "Qtom.I.LA.SC.01" "Qtom.I.LA.SC.01"
##  [84,] "Qtom.I.LA.02"    "Qtom.I.LA.02"    "Qtom.I.LA.02"   
##  [85,] "Qtom.I.LA.05"    "Qtom.I.LA.05"    "Qtom.I.LA.05"   
##  [86,] "Qtom.I.LA.06"    "Qtom.I.LA.06"    "Qtom.I.LA.06"   
##  [87,] "Qtom.I.LA.08"    "Qtom.I.LA.08"    "Qtom.I.LA.08"   
##  [88,] "Qtom.I.LA.10"    "Qtom.I.LA.10"    "Qtom.I.LA.10"   
##  [89,] "Qtom.I.LA.11"    "Qtom.I.LA.11"    "Qtom.I.LA.11"   
##  [90,] "Qtom.I.LA.12"    "Qtom.I.LA.12"    "Qtom.I.LA.12"   
##  [91,] "Qtom.I.LA.13"    "Qtom.I.LA.13"    "Qtom.I.LA.13"   
##  [92,] "Qtom.I.LA.21"    "Qtom.I.LA.21"    "Qtom.I.LA.21"   
##  [93,] "Qtom.I.LA.22"    "Qtom.I.LA.22"    "Qtom.I.LA.22"   
##  [94,] "Qtom.I.LA.23"    "Qtom.I.LA.23"    "Qtom.I.LA.23"   
##  [95,] "Qtom.I.LA.24"    "Qtom.I.LA.24"    "Qtom.I.LA.24"   
##  [96,] "Qtom.I.SB.01"    "Qtom.I.SB.01"    "Qtom.I.SB.01"   
##  [97,] "Qtom.I.SB.04"    "Qtom.I.SB.04"    "Qtom.I.SB.04"   
##  [98,] "Qtom.I.SB.09"    "Qtom.I.SB.09"    "Qtom.I.SB.09"   
##  [99,] "Qtom.I.SB.10"    "Qtom.I.SB.10"    "Qtom.I.SB.10"   
## [100,] "Qtom.I.SB.11"    "Qtom.I.SB.11"    "Qtom.I.SB.11"   
## [101,] "Qtom.I.SB.12"    "Qtom.I.SB.12"    "Qtom.I.SB.12"   
## [102,] "Qtom.I.SB.28"    "Qtom.I.SB.28"    "Qtom.I.SB.28"   
## [103,] "Qtom.I.SB.33"    "Qtom.I.SB.33"    "Qtom.I.SB.33"   
## [104,] "Qtom.I.SB.35"    "Qtom.I.SB.35"    "Qtom.I.SB.35"   
## [105,] "Qtom.I.SB.36"    "Qtom.I.SB.36"    "Qtom.I.SB.36"   
## [106,] "Qtom.I.SB.37"    "Qtom.I.SB.37"    "Qtom.I.SB.37"   
## [107,] "Qtom.I.SB.38"    "Qtom.I.SB.38"    "Qtom.I.SB.38"   
## [108,] "Qtom.I.SB.40"    "Qtom.I.SB.40"    "Qtom.I.SB.40"   
## [109,] "Qtom.I.SB.41"    "Qtom.I.SB.41"    "Qtom.I.SB.41"   
## [110,] "Qtom.I.SB.42"    "Qtom.I.SB.42"    "Qtom.I.SB.42"   
## [111,] "Qtom.I.SB.44"    "Qtom.I.SB.44"    "Qtom.I.SB.44"   
## [112,] "Qtom.I.SB.45"    "Qtom.I.SB.45"    "Qtom.I.SB.45"   
## [113,] "Qtom.I.SB.73"    "Qtom.I.SB.73"    "Qtom.I.SB.73"   
## [114,] "Qtom.I.SB.74"    "Qtom.I.SB.74"    "Qtom.I.SB.74"   
## [115,] "Qtom.I.VEN.01"   "Qtom.I.VEN.01"   "Qtom.I.VEN.01"  
## [116,] "Qtom.I.VEN.02"   "Qtom.I.VEN.02"   "Qtom.I.VEN.02"  
## [117,] "Qtom.I.VEN.03"   "Qtom.I.VEN.03"   "Qtom.I.VEN.03"  
## [118,] "Qtom.I.VEN.04"   "Qtom.I.VEN.04"   "Qtom.I.VEN.04"  
## [119,] "Qtom.I.VEN.05"   "Qtom.I.VEN.05"   "Qtom.I.VEN.05"  
## [120,] "Qtom.JJK.SB.01"  "Qtom.JJK.SB.01"  "Qtom.JJK.SB.01" 
## [121,] "Qtom.JJK.SB.03"  "Qtom.JJK.SB.03"  "Qtom.JJK.SB.03" 
## [122,] "Qtom.JJK.SB.07"  "Qtom.JJK.SB.07"  "Qtom.JJK.SB.07" 
## [123,] "Qtom.JJK.SB.08"  "Qtom.JJK.SB.08"  "Qtom.JJK.SB.08" 
## [124,] "Qtom.JJK.SB.09"  "Qtom.JJK.SB.09"  "Qtom.JJK.SB.09" 
## [125,] "Qtom.JJK.SB.10"  "Qtom.JJK.SB.10"  "Qtom.JJK.SB.10" 
## [126,] "Qtom.JR.SB.01"   "Qtom.JR.SB.01"   "Qtom.JR.SB.01"  
## [127,] "Qtom.S.LA.186"   "Qtom.S.LA.186"   "Qtom.S.LA.186"
```

### 2 Simple PCA

#### 2.1 plot functions

```
# functions to make different types of plots of RDA results

# setup colors and points

# color by island
bg <- c('black', "#3c4a8b","#009c85","#84bc5f","#edb829","#f57404","#b30000")

# point shape by species
sp <- c(23, 22, 21)

###
# just plot the individuals (sites)
plot_simple <- function(dat, info, choices = c(1,2), save = F, file_string, legend = F, legend_pos = 'topright'){
  
  if(save == T){
    filename <- paste('rda_plot_indivs_', file_string, '_PC', choices[1], '_PC', choices[2], '.png', sep = '')
    png(file = filename, width = 10, height = 9, res = 300, units = 'in')
  }
  
  # the plot
  plot(dat, type = 'n', choices = choices)
  points(dat, pch = sp[info$sp], cex = 2, display = 'sites', bg = bg[info$island], col = 'black', choices = choices)
  if(legend == T){
    legend(legend_pos, legend = c(levels(info$sp), levels(info$island)), pch = c(sp, rep(15, 8)), col = c(rep('black', 3), bg), cex = 1)
  }
  
  
  if(save == T){
    dev.off()
  }
  
}

# plot individuals with climate

plot_clim <- function(dat, info, choices = c(1,2), save = F, file_string, legend = F, legend_pos = 'topright'){

  if(save == T){
    filename <- paste('rda_plot_indivs_clim_', file_string, '_PC', choices[1], '_PC', choices[2], '.png', sep = '')
    png(file = filename, width = 10, height = 9, res = 300, units = 'in')
  }

  # the plot
  plot(dat, type = 'n', choices = choices)
  points(dat, pch = sp[info$sp], cex = 2, display = 'sites', bg = bg[info$island], col = 'black', choices = choices)
  text(dat, display = 'bp', choices = choices)
  if(legend == T){
    legend(legend_pos, legend = c(levels(info$sp), levels(info$island)), pch = c(sp, rep(15, 8)), col = c(rep('black', 3), bg), cex = 1)
  }


  if(save == T){
    dev.off()
  }

}
```

#### 2.2 All samples

```
# RDA with just SNP data
# since this is not partial or constrained, this just runs a regular PCA
# from help(rda): "If both matrices Z and Y are missing, the data matrix is analysed by ordinary correspondence analysis (or principal components analysis)."

rda <- rda(imp, scale = T)

rda
```

```
## Call: rda(X = imp, scale = T)
## 
##               Inertia Rank
## Total          585298     
## Unconstrained  585298  126
## Inertia is correlations 
## 
## Eigenvalues for unconstrained axes:
##   PC1   PC2   PC3   PC4   PC5   PC6   PC7   PC8 
## 16078 12277  8152  7876  7338  7186  6959  6772 
## (Showing 8 of 126 unconstrained eigenvalues)
```

```
#summary(rda)
rda.info <- summary(rda)
head(rda.info)
```

```
## 
## Call:
## rda(X = imp, scale = T) 
## 
## Partitioning of correlations:
##               Inertia Proportion
## Total          585298          1
## Unconstrained  585298          1
## 
## Eigenvalues, and their contribution to the correlations 
## 
## Importance of components:
##                             PC1       PC2       PC3       PC4       PC5
## Eigenvalue            1.608e+04 1.228e+04 8.152e+03 7.876e+03 7.338e+03
## Proportion Explained  2.747e-02 2.097e-02 1.393e-02 1.346e-02 1.254e-02
## Cumulative Proportion 2.747e-02 4.844e-02 6.237e-02 7.583e-02 8.837e-02
##                             PC6       PC7       PC8       PC9      PC10
## Eigenvalue            7.186e+03 6.959e+03 6.772e+03 6.578e+03 6.565e+03
## Proportion Explained  1.228e-02 1.189e-02 1.157e-02 1.124e-02 1.122e-02
## Cumulative Proportion 1.006e-01 1.125e-01 1.241e-01 1.353e-01 1.466e-01
##                            PC11      PC12      PC13      PC14      PC15
## Eigenvalue            6378.9495 6.141e+03 6.084e+03 6.065e+03 5.976e+03
## Proportion Explained     0.0109 1.049e-02 1.039e-02 1.036e-02 1.021e-02
## Cumulative Proportion    0.1575 1.679e-01 1.783e-01 1.887e-01 1.989e-01
##                            PC16      PC17      PC18      PC19      PC20
## Eigenvalue            5.961e+03 5.867e+03 5.850e+03 5.727e+03 5.691e+03
## Proportion Explained  1.018e-02 1.002e-02 9.994e-03 9.784e-03 9.724e-03
## Cumulative Proportion 2.091e-01 2.191e-01 2.291e-01 2.389e-01 2.486e-01
##                            PC21      PC22      PC23      PC24      PC25
## Eigenvalue            5.663e+03 5.651e+03 5.637e+03 5.614e+03 5.568e+03
## Proportion Explained  9.675e-03 9.654e-03 9.631e-03 9.592e-03 9.513e-03
## Cumulative Proportion 2.583e-01 2.680e-01 2.776e-01 2.872e-01 2.967e-01
##                            PC26      PC27      PC28      PC29      PC30
## Eigenvalue            5.564e+03 5.524e+03 5.502e+03 5.460e+03 5.425e+03
## Proportion Explained  9.507e-03 9.438e-03 9.401e-03 9.328e-03 9.269e-03
## Cumulative Proportion 3.062e-01 3.156e-01 3.250e-01 3.344e-01 3.436e-01
##                            PC31      PC32      PC33      PC34      PC35
## Eigenvalue            5384.9606 5.364e+03 5.269e+03 5.166e+03 5.123e+03
## Proportion Explained     0.0092 9.165e-03 9.001e-03 8.826e-03 8.753e-03
## Cumulative Proportion    0.3528 3.620e-01 3.710e-01 3.798e-01 3.886e-01
##                            PC36      PC37      PC38      PC39      PC40
## Eigenvalue            5.102e+03 5.069e+03 5.045e+03 5.020e+03 4.933e+03
## Proportion Explained  8.718e-03 8.660e-03 8.620e-03 8.577e-03 8.427e-03
## Cumulative Proportion 3.973e-01 4.060e-01 4.146e-01 4.232e-01 4.316e-01
##                            PC41      PC42      PC43      PC44      PC45
## Eigenvalue            4.915e+03 4.860e+03 4.821e+03 4.786e+03 4.757e+03
## Proportion Explained  8.398e-03 8.304e-03 8.237e-03 8.178e-03 8.127e-03
## Cumulative Proportion 4.400e-01 4.483e-01 4.565e-01 4.647e-01 4.728e-01
##                            PC46      PC47      PC48      PC49      PC50
## Eigenvalue            4.737e+03 4.715e+03 4.691e+03 4.648e+03 4.611e+03
## Proportion Explained  8.093e-03 8.056e-03 8.015e-03 7.941e-03 7.878e-03
## Cumulative Proportion 4.809e-01 4.890e-01 4.970e-01 5.049e-01 5.128e-01
##                            PC51      PC52      PC53      PC54      PC55
## Eigenvalue            4.575e+03 4.548e+03 4.535e+03 4.531e+03 4.495e+03
## Proportion Explained  7.816e-03 7.771e-03 7.748e-03 7.742e-03 7.679e-03
## Cumulative Proportion 5.206e-01 5.284e-01 5.361e-01 5.439e-01 5.516e-01
##                            PC56      PC57      PC58      PC59      PC60
## Eigenvalue            4.471e+03 4.414e+03 4.386e+03 4.371e+03 4.350e+03
## Proportion Explained  7.638e-03 7.542e-03 7.493e-03 7.468e-03 7.432e-03
## Cumulative Proportion 5.592e-01 5.667e-01 5.742e-01 5.817e-01 5.891e-01
##                            PC61      PC62      PC63      PC64      PC65
## Eigenvalue            4.337e+03 4.327e+03 4.271e+03 4.256e+03 4.247e+03
## Proportion Explained  7.410e-03 7.393e-03 7.296e-03 7.271e-03 7.256e-03
## Cumulative Proportion 5.965e-01 6.039e-01 6.112e-01 6.185e-01 6.258e-01
##                            PC66      PC67      PC68      PC69      PC70
## Eigenvalue            4.223e+03 4.210e+03 4.199e+03 4.171e+03 4.147e+03
## Proportion Explained  7.214e-03 7.193e-03 7.174e-03 7.125e-03 7.086e-03
## Cumulative Proportion 6.330e-01 6.402e-01 6.473e-01 6.545e-01 6.616e-01
##                            PC71      PC72      PC73      PC74      PC75
## Eigenvalue            4.132e+03 4.096e+03 4.084e+03 4.069e+03 4.055e+03
## Proportion Explained  7.060e-03 6.998e-03 6.978e-03 6.952e-03 6.928e-03
## Cumulative Proportion 6.686e-01 6.756e-01 6.826e-01 6.895e-01 6.965e-01
##                            PC76      PC77      PC78      PC79      PC80
## Eigenvalue            4.042e+03 4.018e+03 3.998e+03 3.971e+03 3.949e+03
## Proportion Explained  6.906e-03 6.864e-03 6.830e-03 6.784e-03 6.747e-03
## Cumulative Proportion 7.034e-01 7.102e-01 7.171e-01 7.239e-01 7.306e-01
##                            PC81      PC82      PC83      PC84      PC85
## Eigenvalue            3.937e+03 3.898e+03 3.872e+03 3.844e+03 3.828e+03
## Proportion Explained  6.727e-03 6.660e-03 6.615e-03 6.568e-03 6.541e-03
## Cumulative Proportion 7.373e-01 7.440e-01 7.506e-01 7.572e-01 7.637e-01
##                            PC86      PC87      PC88      PC89      PC90
## Eigenvalue            3.819e+03 3.798e+03 3.784e+03 3.773e+03 3.766e+03
## Proportion Explained  6.524e-03 6.488e-03 6.465e-03 6.446e-03 6.435e-03
## Cumulative Proportion 7.702e-01 7.767e-01 7.832e-01 7.896e-01 7.961e-01
##                            PC91      PC92      PC93      PC94      PC95
## Eigenvalue            3.713e+03 3.701e+03 3.696e+03 3.682e+03 3.666e+03
## Proportion Explained  6.344e-03 6.323e-03 6.315e-03 6.290e-03 6.263e-03
## Cumulative Proportion 8.024e-01 8.087e-01 8.151e-01 8.213e-01 8.276e-01
##                            PC96      PC97      PC98      PC99     PC100
## Eigenvalue            3.653e+03 3.643e+03 3.637e+03 3.619e+03 3.590e+03
## Proportion Explained  6.241e-03 6.225e-03 6.213e-03 6.182e-03 6.133e-03
## Cumulative Proportion 8.338e-01 8.401e-01 8.463e-01 8.525e-01 8.586e-01
##                           PC101     PC102     PC103     PC104     PC105
## Eigenvalue            3.587e+03 3.578e+03 3.567e+03 3.558e+03 3.549e+03
## Proportion Explained  6.129e-03 6.114e-03 6.095e-03 6.080e-03 6.064e-03
## Cumulative Proportion 8.647e-01 8.708e-01 8.769e-01 8.830e-01 8.891e-01
##                           PC106     PC107     PC108     PC109     PC110
## Eigenvalue            3.481e+03 3.471e+03 3.450e+03 3.432e+03 3.412e+03
## Proportion Explained  5.948e-03 5.930e-03 5.895e-03 5.863e-03 5.829e-03
## Cumulative Proportion 8.950e-01 9.010e-01 9.069e-01 9.127e-01 9.185e-01
##                           PC111     PC112     PC113     PC114     PC115
## Eigenvalue            3.388e+03 3.378e+03 3.347e+03 3.304e+03 3.247e+03
## Proportion Explained  5.789e-03 5.771e-03 5.718e-03 5.644e-03 5.548e-03
## Cumulative Proportion 9.243e-01 9.301e-01 9.358e-01 9.415e-01 9.470e-01
##                           PC116     PC117     PC118     PC119     PC120
## Eigenvalue            3.233e+03 3.128e+03 3.122e+03 3.017e+03 2.993e+03
## Proportion Explained  5.524e-03 5.345e-03 5.334e-03 5.155e-03 5.113e-03
## Cumulative Proportion 9.525e-01 9.579e-01 9.632e-01 9.684e-01 9.735e-01
##                           PC121     PC122     PC123     PC124     PC125
## Eigenvalue            2.888e+03 2.825e+03 2.803e+03 2.770e+03 2.564e+03
## Proportion Explained  4.935e-03 4.826e-03 4.789e-03 4.732e-03 4.381e-03
## Cumulative Proportion 9.784e-01 9.833e-01 9.880e-01 9.928e-01 9.972e-01
##                           PC126
## Eigenvalue            1.666e+03
## Proportion Explained  2.847e-03
## Cumulative Proportion 1.000e+00
## 
## Scaling 2 for species and site scores
## * Species are scaled proportional to eigenvalues
## * Sites are unscaled: weighted dispersion equal on all dimensions
## * General scaling constant of scores:  92.66952 
## 
## 
## Species scores
## 
##                     PC1       PC2       PC3       PC4        PC5      PC6
## 1_39789_A_G_G  0.016254  0.006390  0.010577  0.031286  0.0063812 0.011929
## 1_52176_G_A_A  0.017263 -0.015926 -0.014821  0.029731 -0.0053177 0.005304
## 1_52195_G_A_A -0.005537 -0.003335 -0.003549 -0.003055  0.0014905 0.001922
## 1_52213_C_T_T  0.020633  0.004017 -0.012757 -0.013708 -0.0008197 0.006172
## 1_52248_T_C_T -0.003496 -0.006368  0.005046  0.007231  0.0022652 0.004594
## 1_52331_G_T_T  0.007229  0.015860  0.014081 -0.011938  0.0006462 0.008550
## ....                                                                     
## 
## 
## Site scores (weighted sums of species scores)
## 
##                  PC1   PC2       PC3    PC4    PC5     PC6
## Qchr.A.232    -4.667 16.51 -19.76969 1.5843 -3.565 10.8109
## Qchr.A.275    -5.026 16.15 -18.28864 2.9243 -1.509 10.1214
## Qchr.A.ED.102 -6.218 19.58 -16.33083 3.5002 -1.231  9.9266
## Qchr.A.ELD.53 -6.397 20.59 -20.36504 3.4481 -3.281 12.1158
## Qchr.A.KER.16 -4.952 14.06  -5.34261 2.8803  2.650  4.9942
## Qchr.A.LA.02  -6.543 17.98   0.03709 0.9812  0.270 -0.8393
## ....
```

```
screeplot(rda)
```

```
# plot with functions!
plot_simple(rda, info=info, legend = T, legend_pos = 'topleft', save = F, file_string = 'Qtom107.Qchr17.Qssp3')
```

```
plot_simple(rda, info = info, choices = c(1,3), legend = F, legend_pos = 'topleft', save = F, file_string = 'Qtom107.Qchr17.Qssp3')
```

```
# plot with text only

#png(file = 'rda_plot_indivs_Qtom107.Qchr17.Qssp3_textIDs_PC1_PC2.png', width = 10, height = 9, res = 300, units = 'in')
plot(rda, type = 'n')
text(rda, cex = 1, display = 'sites', col = bg[info$island])
legend('topleft', legend = levels(info$island), fill = bg, cex = 1)
```

```
#dev.off()

#png(file = 'rda_plot_indivs_Qtom107.Qchr17.Qssp3_textIDs_PC1_PC3.png', width = 10, height = 9, res = 300, units = 'in')
plot(rda, type = 'n', choices = c(1,3))
text(rda, choices = c(1,3), cex = 1, display = 'sites', col = bg[info$island])
legend('topleft', legend = levels(info$island), fill = bg, cex = 1)
```

```
#dev.off()
```

#### 2.3 without Guadalupe

```
# since guadalupe appears divergent, make a PCA without it to show the relationships among other islands

imp.sub <- imp[info$island != 'Guadalupe Island',]
info.sub <- info[info$island != 'Guadalupe Island',]
```

```
# run and plot PCA

# make sure they're all in the same order
cbind(rownames(imp.sub), rownames(info.sub))
```

```
##        [,1]              [,2]             
##   [1,] "Qchr.A.232"      "Qchr.A.232"     
##   [2,] "Qchr.A.275"      "Qchr.A.275"     
##   [3,] "Qchr.A.ED.102"   "Qchr.A.ED.102"  
##   [4,] "Qchr.A.ELD.53"   "Qchr.A.ELD.53"  
##   [5,] "Qchr.A.KER.16"   "Qchr.A.KER.16"  
##   [6,] "Qchr.A.LA.02"    "Qchr.A.LA.02"   
##   [7,] "Qchr.A.LA.215"   "Qchr.A.LA.215"  
##   [8,] "Qchr.A.LA.222"   "Qchr.A.LA.222"  
##   [9,] "Qchr.A.SIS.309"  "Qchr.A.SIS.309" 
##  [10,] "Qchr.A.PLU.65"   "Qchr.A.PLU.65"  
##  [11,] "Qchr.A.SBD.83"   "Qchr.A.SBD.83"  
##  [12,] "Qchr.I.LA.20"    "Qchr.I.LA.20"   
##  [13,] "Qchr.I.LA.25"    "Qchr.I.LA.25"   
##  [14,] "Qchr.I.SB.02"    "Qchr.I.SB.02"   
##  [15,] "Qchr.I.SB.03"    "Qchr.I.SB.03"   
##  [16,] "Qchr.JJK.SB.13"  "Qchr.JJK.SB.13" 
##  [17,] "Qchr.JR.SB.01"   "Qchr.JR.SB.01"  
##  [18,] "Qspp.I.LA.03"    "Qspp.I.LA.03"   
##  [19,] "Qspp.I.LA.04"    "Qspp.I.LA.04"   
##  [20,] "Qspp.I.LA.14"    "Qspp.I.LA.14"   
##  [21,] "Qtom.A.LA.169"   "Qtom.A.LA.169"  
##  [22,] "Qtom.A.LA.175"   "Qtom.A.LA.175"  
##  [23,] "Qtom.A.LA.176"   "Qtom.A.LA.176"  
##  [24,] "Qtom.A.LA.180"   "Qtom.A.LA.180"  
##  [25,] "Qtom.A.LA.181"   "Qtom.A.LA.181"  
##  [26,] "Qtom.A.LA.183"   "Qtom.A.LA.183"  
##  [27,] "Qtom.A.LA.190"   "Qtom.A.LA.190"  
##  [28,] "Qtom.A.LA.191"   "Qtom.A.LA.191"  
##  [29,] "Qtom.A.LA.198"   "Qtom.A.LA.198"  
##  [30,] "Qtom.A.LA.200"   "Qtom.A.LA.200"  
##  [31,] "Qtom.A.LA.204"   "Qtom.A.LA.204"  
##  [32,] "Qtom.A.LA.207"   "Qtom.A.LA.207"  
##  [33,] "Qtom.A.LA.208"   "Qtom.A.LA.208"  
##  [34,] "Qtom.A.LA.209"   "Qtom.A.LA.209"  
##  [35,] "Qtom.A.LA.218"   "Qtom.A.LA.218"  
##  [36,] "Qtom.A.LA.221"   "Qtom.A.LA.221"  
##  [37,] "Qtom.A.SB.324"   "Qtom.A.SB.324"  
##  [38,] "Qtom.A.SB.325"   "Qtom.A.SB.325"  
##  [39,] "Qtom.A.SB.327"   "Qtom.A.SB.327"  
##  [40,] "Qtom.A.SB.329"   "Qtom.A.SB.329"  
##  [41,] "Qtom.A.SB.330"   "Qtom.A.SB.330"  
##  [42,] "Qtom.A.SB.331"   "Qtom.A.SB.331"  
##  [43,] "Qtom.A.SB.332"   "Qtom.A.SB.332"  
##  [44,] "Qtom.A.SB.333"   "Qtom.A.SB.333"  
##  [45,] "Qtom.A.SB.335"   "Qtom.A.SB.335"  
##  [46,] "Qtom.A.SB.336"   "Qtom.A.SB.336"  
##  [47,] "Qtom.A.SB.337"   "Qtom.A.SB.337"  
##  [48,] "Qtom.A.SB.338"   "Qtom.A.SB.338"  
##  [49,] "Qtom.A.SB.339"   "Qtom.A.SB.339"  
##  [50,] "Qtom.A.SB.341"   "Qtom.A.SB.341"  
##  [51,] "Qtom.A.SB.342"   "Qtom.A.SB.342"  
##  [52,] "Qtom.A.SB.343"   "Qtom.A.SB.343"  
##  [53,] "Qtom.A.SB.344"   "Qtom.A.SB.344"  
##  [54,] "Qtom.A.SB.345"   "Qtom.A.SB.345"  
##  [55,] "Qtom.A.SB.346"   "Qtom.A.SB.346"  
##  [56,] "Qtom.A.SB.347"   "Qtom.A.SB.347"  
##  [57,] "Qtom.A.SB.348"   "Qtom.A.SB.348"  
##  [58,] "Qtom.A.SB.352"   "Qtom.A.SB.352"  
##  [59,] "Qtom.A.SB.353"   "Qtom.A.SB.353"  
##  [60,] "Qtom.A.SB.355"   "Qtom.A.SB.355"  
##  [61,] "Qtom.I.LA.SC.01" "Qtom.I.LA.SC.01"
##  [62,] "Qtom.I.LA.02"    "Qtom.I.LA.02"   
##  [63,] "Qtom.I.LA.05"    "Qtom.I.LA.05"   
##  [64,] "Qtom.I.LA.06"    "Qtom.I.LA.06"   
##  [65,] "Qtom.I.LA.08"    "Qtom.I.LA.08"   
##  [66,] "Qtom.I.LA.10"    "Qtom.I.LA.10"   
##  [67,] "Qtom.I.LA.11"    "Qtom.I.LA.11"   
##  [68,] "Qtom.I.LA.12"    "Qtom.I.LA.12"   
##  [69,] "Qtom.I.LA.13"    "Qtom.I.LA.13"   
##  [70,] "Qtom.I.LA.21"    "Qtom.I.LA.21"   
##  [71,] "Qtom.I.LA.22"    "Qtom.I.LA.22"   
##  [72,] "Qtom.I.LA.23"    "Qtom.I.LA.23"   
##  [73,] "Qtom.I.LA.24"    "Qtom.I.LA.24"   
##  [74,] "Qtom.I.SB.01"    "Qtom.I.SB.01"   
##  [75,] "Qtom.I.SB.04"    "Qtom.I.SB.04"   
##  [76,] "Qtom.I.SB.09"    "Qtom.I.SB.09"   
##  [77,] "Qtom.I.SB.10"    "Qtom.I.SB.10"   
##  [78,] "Qtom.I.SB.11"    "Qtom.I.SB.11"   
##  [79,] "Qtom.I.SB.12"    "Qtom.I.SB.12"   
##  [80,] "Qtom.I.SB.28"    "Qtom.I.SB.28"   
##  [81,] "Qtom.I.SB.33"    "Qtom.I.SB.33"   
##  [82,] "Qtom.I.SB.35"    "Qtom.I.SB.35"   
##  [83,] "Qtom.I.SB.36"    "Qtom.I.SB.36"   
##  [84,] "Qtom.I.SB.37"    "Qtom.I.SB.37"   
##  [85,] "Qtom.I.SB.38"    "Qtom.I.SB.38"   
##  [86,] "Qtom.I.SB.40"    "Qtom.I.SB.40"   
##  [87,] "Qtom.I.SB.41"    "Qtom.I.SB.41"   
##  [88,] "Qtom.I.SB.42"    "Qtom.I.SB.42"   
##  [89,] "Qtom.I.SB.44"    "Qtom.I.SB.44"   
##  [90,] "Qtom.I.SB.45"    "Qtom.I.SB.45"   
##  [91,] "Qtom.I.SB.73"    "Qtom.I.SB.73"   
##  [92,] "Qtom.I.SB.74"    "Qtom.I.SB.74"   
##  [93,] "Qtom.I.VEN.01"   "Qtom.I.VEN.01"  
##  [94,] "Qtom.I.VEN.02"   "Qtom.I.VEN.02"  
##  [95,] "Qtom.I.VEN.03"   "Qtom.I.VEN.03"  
##  [96,] "Qtom.I.VEN.04"   "Qtom.I.VEN.04"  
##  [97,] "Qtom.I.VEN.05"   "Qtom.I.VEN.05"  
##  [98,] "Qtom.JJK.SB.01"  "Qtom.JJK.SB.01" 
##  [99,] "Qtom.JJK.SB.03"  "Qtom.JJK.SB.03" 
## [100,] "Qtom.JJK.SB.07"  "Qtom.JJK.SB.07" 
## [101,] "Qtom.JJK.SB.08"  "Qtom.JJK.SB.08" 
## [102,] "Qtom.JJK.SB.09"  "Qtom.JJK.SB.09" 
## [103,] "Qtom.JJK.SB.10"  "Qtom.JJK.SB.10" 
## [104,] "Qtom.JR.SB.01"   "Qtom.JR.SB.01"  
## [105,] "Qtom.S.LA.186"   "Qtom.S.LA.186"
```

```
rda <- rda(imp.sub, scale = T)

plot_simple(rda, info = info.sub, legend = T, legend_pos = 'topleft', save = F, file_string = 'Qtom107.Qchr17.Qssp3_noGuadalupe')
```

```
plot_simple(rda, info = info.sub, choices = c(1,3), legend = F, legend_pos = 'topright', save = F, file_string = 'Qtom107.Qchr17.Qssp3_noGuadalupe')
```

```
#png(file = 'rda_plot_indivs_Qtom107.Qchr17.Qssp3_noGuadalupe_textIDs_PC1_PC2.png', width = 10, height = 9, res = 300, units = 'in')
plot(rda, type = 'n')
text(rda, cex = 1, display = 'sites', col = bg[info.sub$island])
legend('topright', legend = levels(info$island), fill = bg, cex = 1)
```

```
#dev.off()

#png(file = 'rda_plot_indivs_textIDs_Qtom107.Qchr17.Qssp3_noGuadalupe_PC1_PC3.png', width = 10, height = 9, res = 300, units = 'in')
plot(rda, type = 'n', choices = c(1,3))
text(rda, choices = c(1,3), cex = 1, display = 'sites', col = bg[info.sub$island])
```

```
#legend('bottomright', legend = levels(info$island), fill = bg, cex = 1)
#dev.off()
```

### 3 RDA with climate

Now include both genetic and climate data

#### 3.1 Basic RDA with all individuals and climate variables to visualize

```
# subsetting to get all individuals with snp and climate data
# note: climate dataset has to be matched to rownames(info), rownames(snps) uses the info$id_vcf naming, which is different for some samples

# get only individuals with climate data (some are missing)
tmp.info <- info[rownames(info) %in% rownames(clim),]

dim(tmp.info)
```

```
## [1] 126  20
```

```
dim(clim)
```

```
## [1] 295  20
```

```
# get climate for the individuals with snp data
tmp.clim <- clim[match(rownames(tmp.info), rownames(clim)),]

# they are in the same order
dim(tmp.info)
```

```
## [1] 126  20
```

```
dim(tmp.clim)
```

```
## [1] 126  20
```

```
cbind(rownames(tmp.info), rownames(tmp.clim))
```

```
##        [,1]              [,2]             
##   [1,] "Qchr.A.232"      "Qchr.A.232"     
##   [2,] "Qchr.A.275"      "Qchr.A.275"     
##   [3,] "Qchr.A.ED.102"   "Qchr.A.ED.102"  
##   [4,] "Qchr.A.ELD.53"   "Qchr.A.ELD.53"  
##   [5,] "Qchr.A.KER.16"   "Qchr.A.KER.16"  
##   [6,] "Qchr.A.LA.02"    "Qchr.A.LA.02"   
##   [7,] "Qchr.A.LA.215"   "Qchr.A.LA.215"  
##   [8,] "Qchr.A.LA.222"   "Qchr.A.LA.222"  
##   [9,] "Qchr.A.SIS.309"  "Qchr.A.SIS.309" 
##  [10,] "Qchr.A.PLU.65"   "Qchr.A.PLU.65"  
##  [11,] "Qchr.A.SBD.83"   "Qchr.A.SBD.83"  
##  [12,] "Qchr.I.LA.20"    "Qchr.I.LA.20"   
##  [13,] "Qchr.I.LA.25"    "Qchr.I.LA.25"   
##  [14,] "Qchr.I.SB.02"    "Qchr.I.SB.02"   
##  [15,] "Qchr.I.SB.03"    "Qchr.I.SB.03"   
##  [16,] "Qchr.JJK.SB.13"  "Qchr.JJK.SB.13" 
##  [17,] "Qchr.JR.SB.01"   "Qchr.JR.SB.01"  
##  [18,] "Qspp.I.LA.03"    "Qspp.I.LA.03"   
##  [19,] "Qspp.I.LA.04"    "Qspp.I.LA.04"   
##  [20,] "Qspp.I.LA.14"    "Qspp.I.LA.14"   
##  [21,] "Qtom.A.LA.169"   "Qtom.A.LA.169"  
##  [22,] "Qtom.A.LA.175"   "Qtom.A.LA.175"  
##  [23,] "Qtom.A.LA.176"   "Qtom.A.LA.176"  
##  [24,] "Qtom.A.LA.180"   "Qtom.A.LA.180"  
##  [25,] "Qtom.A.LA.181"   "Qtom.A.LA.181"  
##  [26,] "Qtom.A.LA.183"   "Qtom.A.LA.183"  
##  [27,] "Qtom.A.LA.190"   "Qtom.A.LA.190"  
##  [28,] "Qtom.A.LA.191"   "Qtom.A.LA.191"  
##  [29,] "Qtom.A.LA.198"   "Qtom.A.LA.198"  
##  [30,] "Qtom.A.LA.200"   "Qtom.A.LA.200"  
##  [31,] "Qtom.A.LA.204"   "Qtom.A.LA.204"  
##  [32,] "Qtom.A.LA.207"   "Qtom.A.LA.207"  
##  [33,] "Qtom.A.LA.208"   "Qtom.A.LA.208"  
##  [34,] "Qtom.A.LA.209"   "Qtom.A.LA.209"  
##  [35,] "Qtom.A.LA.218"   "Qtom.A.LA.218"  
##  [36,] "Qtom.A.LA.221"   "Qtom.A.LA.221"  
##  [37,] "Qtom.A.SB.324"   "Qtom.A.SB.324"  
##  [38,] "Qtom.A.SB.325"   "Qtom.A.SB.325"  
##  [39,] "Qtom.A.SB.327"   "Qtom.A.SB.327"  
##  [40,] "Qtom.A.SB.329"   "Qtom.A.SB.329"  
##  [41,] "Qtom.A.SB.330"   "Qtom.A.SB.330"  
##  [42,] "Qtom.A.SB.331"   "Qtom.A.SB.331"  
##  [43,] "Qtom.A.SB.332"   "Qtom.A.SB.332"  
##  [44,] "Qtom.A.SB.333"   "Qtom.A.SB.333"  
##  [45,] "Qtom.A.SB.335"   "Qtom.A.SB.335"  
##  [46,] "Qtom.A.SB.336"   "Qtom.A.SB.336"  
##  [47,] "Qtom.A.SB.337"   "Qtom.A.SB.337"  
##  [48,] "Qtom.A.SB.338"   "Qtom.A.SB.338"  
##  [49,] "Qtom.A.SB.339"   "Qtom.A.SB.339"  
##  [50,] "Qtom.A.SB.341"   "Qtom.A.SB.341"  
##  [51,] "Qtom.A.SB.342"   "Qtom.A.SB.342"  
##  [52,] "Qtom.A.SB.343"   "Qtom.A.SB.343"  
##  [53,] "Qtom.A.SB.344"   "Qtom.A.SB.344"  
##  [54,] "Qtom.A.SB.345"   "Qtom.A.SB.345"  
##  [55,] "Qtom.A.SB.346"   "Qtom.A.SB.346"  
##  [56,] "Qtom.A.SB.347"   "Qtom.A.SB.347"  
##  [57,] "Qtom.A.SB.348"   "Qtom.A.SB.348"  
##  [58,] "Qtom.A.SB.352"   "Qtom.A.SB.352"  
##  [59,] "Qtom.A.SB.353"   "Qtom.A.SB.353"  
##  [60,] "Qtom.A.SB.355"   "Qtom.A.SB.355"  
##  [61,] "Qtom.I.BC.10"    "Qtom.I.BC.10"   
##  [62,] "Qtom.I.BC.11"    "Qtom.I.BC.11"   
##  [63,] "Qtom.I.BC.12"    "Qtom.I.BC.12"   
##  [64,] "Qtom.I.BC.13"    "Qtom.I.BC.13"   
##  [65,] "Qtom.I.BC.18"    "Qtom.I.BC.18"   
##  [66,] "Qtom.I.BC.20"    "Qtom.I.BC.20"   
##  [67,] "Qtom.I.BC.21"    "Qtom.I.BC.21"   
##  [68,] "Qtom.I.BC.23"    "Qtom.I.BC.23"   
##  [69,] "Qtom.I.BC.24"    "Qtom.I.BC.24"   
##  [70,] "Qtom.I.BC.32"    "Qtom.I.BC.32"   
##  [71,] "Qtom.I.BC.33"    "Qtom.I.BC.33"   
##  [72,] "Qtom.I.BC.35"    "Qtom.I.BC.35"   
##  [73,] "Qtom.I.BC.39"    "Qtom.I.BC.39"   
##  [74,] "Qtom.I.BC.4"     "Qtom.I.BC.4"    
##  [75,] "Qtom.I.BC.40"    "Qtom.I.BC.40"   
##  [76,] "Qtom.I.BC.41"    "Qtom.I.BC.41"   
##  [77,] "Qtom.I.BC.42"    "Qtom.I.BC.42"   
##  [78,] "Qtom.I.BC.43"    "Qtom.I.BC.43"   
##  [79,] "Qtom.I.BC.44"    "Qtom.I.BC.44"   
##  [80,] "Qtom.I.BC.45"    "Qtom.I.BC.45"   
##  [81,] "Qtom.I.BC.46"    "Qtom.I.BC.46"   
##  [82,] "Qtom.I.BC.48"    "Qtom.I.BC.48"   
##  [83,] "Qtom.I.LA.SC.01" "Qtom.I.LA.SC.01"
##  [84,] "Qtom.I.LA.05"    "Qtom.I.LA.05"   
##  [85,] "Qtom.I.LA.06"    "Qtom.I.LA.06"   
##  [86,] "Qtom.I.LA.08"    "Qtom.I.LA.08"   
##  [87,] "Qtom.I.LA.10"    "Qtom.I.LA.10"   
##  [88,] "Qtom.I.LA.11"    "Qtom.I.LA.11"   
##  [89,] "Qtom.I.LA.12"    "Qtom.I.LA.12"   
##  [90,] "Qtom.I.LA.13"    "Qtom.I.LA.13"   
##  [91,] "Qtom.I.LA.21"    "Qtom.I.LA.21"   
##  [92,] "Qtom.I.LA.22"    "Qtom.I.LA.22"   
##  [93,] "Qtom.I.LA.23"    "Qtom.I.LA.23"   
##  [94,] "Qtom.I.LA.24"    "Qtom.I.LA.24"   
##  [95,] "Qtom.I.SB.01"    "Qtom.I.SB.01"   
##  [96,] "Qtom.I.SB.04"    "Qtom.I.SB.04"   
##  [97,] "Qtom.I.SB.09"    "Qtom.I.SB.09"   
##  [98,] "Qtom.I.SB.10"    "Qtom.I.SB.10"   
##  [99,] "Qtom.I.SB.11"    "Qtom.I.SB.11"   
## [100,] "Qtom.I.SB.12"    "Qtom.I.SB.12"   
## [101,] "Qtom.I.SB.28"    "Qtom.I.SB.28"   
## [102,] "Qtom.I.SB.33"    "Qtom.I.SB.33"   
## [103,] "Qtom.I.SB.35"    "Qtom.I.SB.35"   
## [104,] "Qtom.I.SB.36"    "Qtom.I.SB.36"   
## [105,] "Qtom.I.SB.37"    "Qtom.I.SB.37"   
## [106,] "Qtom.I.SB.38"    "Qtom.I.SB.38"   
## [107,] "Qtom.I.SB.40"    "Qtom.I.SB.40"   
## [108,] "Qtom.I.SB.41"    "Qtom.I.SB.41"   
## [109,] "Qtom.I.SB.42"    "Qtom.I.SB.42"   
## [110,] "Qtom.I.SB.44"    "Qtom.I.SB.44"   
## [111,] "Qtom.I.SB.45"    "Qtom.I.SB.45"   
## [112,] "Qtom.I.SB.73"    "Qtom.I.SB.73"   
## [113,] "Qtom.I.SB.74"    "Qtom.I.SB.74"   
## [114,] "Qtom.I.VEN.01"   "Qtom.I.VEN.01"  
## [115,] "Qtom.I.VEN.02"   "Qtom.I.VEN.02"  
## [116,] "Qtom.I.VEN.03"   "Qtom.I.VEN.03"  
## [117,] "Qtom.I.VEN.04"   "Qtom.I.VEN.04"  
## [118,] "Qtom.I.VEN.05"   "Qtom.I.VEN.05"  
## [119,] "Qtom.JJK.SB.01"  "Qtom.JJK.SB.01" 
## [120,] "Qtom.JJK.SB.03"  "Qtom.JJK.SB.03" 
## [121,] "Qtom.JJK.SB.07"  "Qtom.JJK.SB.07" 
## [122,] "Qtom.JJK.SB.08"  "Qtom.JJK.SB.08" 
## [123,] "Qtom.JJK.SB.09"  "Qtom.JJK.SB.09" 
## [124,] "Qtom.JJK.SB.10"  "Qtom.JJK.SB.10" 
## [125,] "Qtom.JR.SB.01"   "Qtom.JR.SB.01"  
## [126,] "Qtom.S.LA.186"   "Qtom.S.LA.186"
```

```
# rename
clim <- tmp.clim
info <- tmp.info

# remove the ones we don't have climate data for from snp dataset
tmp.snps <- snps[rownames(snps) %in% rownames(info),]
tmp.imp <- imp[rownames(imp) %in% rownames(info),]

cbind(rownames(info), rownames(clim), rownames(tmp.snps), rownames(tmp.imp))
```

```
##        [,1]              [,2]              [,3]              [,4]             
##   [1,] "Qchr.A.232"      "Qchr.A.232"      "Qchr.A.232"      "Qchr.A.232"     
##   [2,] "Qchr.A.275"      "Qchr.A.275"      "Qchr.A.275"      "Qchr.A.275"     
##   [3,] "Qchr.A.ED.102"   "Qchr.A.ED.102"   "Qchr.A.ED.102"   "Qchr.A.ED.102"  
##   [4,] "Qchr.A.ELD.53"   "Qchr.A.ELD.53"   "Qchr.A.ELD.53"   "Qchr.A.ELD.53"  
##   [5,] "Qchr.A.KER.16"   "Qchr.A.KER.16"   "Qchr.A.KER.16"   "Qchr.A.KER.16"  
##   [6,] "Qchr.A.LA.02"    "Qchr.A.LA.02"    "Qchr.A.LA.02"    "Qchr.A.LA.02"   
##   [7,] "Qchr.A.LA.215"   "Qchr.A.LA.215"   "Qchr.A.LA.215"   "Qchr.A.LA.215"  
##   [8,] "Qchr.A.LA.222"   "Qchr.A.LA.222"   "Qchr.A.LA.222"   "Qchr.A.LA.222"  
##   [9,] "Qchr.A.SIS.309"  "Qchr.A.SIS.309"  "Qchr.A.SIS.309"  "Qchr.A.SIS.309" 
##  [10,] "Qchr.A.PLU.65"   "Qchr.A.PLU.65"   "Qchr.A.PLU.65"   "Qchr.A.PLU.65"  
##  [11,] "Qchr.A.SBD.83"   "Qchr.A.SBD.83"   "Qchr.A.SBD.83"   "Qchr.A.SBD.83"  
##  [12,] "Qchr.I.LA.20"    "Qchr.I.LA.20"    "Qchr.I.LA.20"    "Qchr.I.LA.20"   
##  [13,] "Qchr.I.LA.25"    "Qchr.I.LA.25"    "Qchr.I.LA.25"    "Qchr.I.LA.25"   
##  [14,] "Qchr.I.SB.02"    "Qchr.I.SB.02"    "Qchr.I.SB.02"    "Qchr.I.SB.02"   
##  [15,] "Qchr.I.SB.03"    "Qchr.I.SB.03"    "Qchr.I.SB.03"    "Qchr.I.SB.03"   
##  [16,] "Qchr.JJK.SB.13"  "Qchr.JJK.SB.13"  "Qchr.JJK.SB.13"  "Qchr.JJK.SB.13" 
##  [17,] "Qchr.JR.SB.01"   "Qchr.JR.SB.01"   "Qchr.JR.SB.01"   "Qchr.JR.SB.01"  
##  [18,] "Qspp.I.LA.03"    "Qspp.I.LA.03"    "Qspp.I.LA.03"    "Qspp.I.LA.03"   
##  [19,] "Qspp.I.LA.04"    "Qspp.I.LA.04"    "Qspp.I.LA.04"    "Qspp.I.LA.04"   
##  [20,] "Qspp.I.LA.14"    "Qspp.I.LA.14"    "Qspp.I.LA.14"    "Qspp.I.LA.14"   
##  [21,] "Qtom.A.LA.169"   "Qtom.A.LA.169"   "Qtom.A.LA.169"   "Qtom.A.LA.169"  
##  [22,] "Qtom.A.LA.175"   "Qtom.A.LA.175"   "Qtom.A.LA.175"   "Qtom.A.LA.175"  
##  [23,] "Qtom.A.LA.176"   "Qtom.A.LA.176"   "Qtom.A.LA.176"   "Qtom.A.LA.176"  
##  [24,] "Qtom.A.LA.180"   "Qtom.A.LA.180"   "Qtom.A.LA.180"   "Qtom.A.LA.180"  
##  [25,] "Qtom.A.LA.181"   "Qtom.A.LA.181"   "Qtom.A.LA.181"   "Qtom.A.LA.181"  
##  [26,] "Qtom.A.LA.183"   "Qtom.A.LA.183"   "Qtom.A.LA.183"   "Qtom.A.LA.183"  
##  [27,] "Qtom.A.LA.190"   "Qtom.A.LA.190"   "Qtom.A.LA.190"   "Qtom.A.LA.190"  
##  [28,] "Qtom.A.LA.191"   "Qtom.A.LA.191"   "Qtom.A.LA.191"   "Qtom.A.LA.191"  
##  [29,] "Qtom.A.LA.198"   "Qtom.A.LA.198"   "Qtom.A.LA.198"   "Qtom.A.LA.198"  
##  [30,] "Qtom.A.LA.200"   "Qtom.A.LA.200"   "Qtom.A.LA.200"   "Qtom.A.LA.200"  
##  [31,] "Qtom.A.LA.204"   "Qtom.A.LA.204"   "Qtom.A.LA.204"   "Qtom.A.LA.204"  
##  [32,] "Qtom.A.LA.207"   "Qtom.A.LA.207"   "Qtom.A.LA.207"   "Qtom.A.LA.207"  
##  [33,] "Qtom.A.LA.208"   "Qtom.A.LA.208"   "Qtom.A.LA.208"   "Qtom.A.LA.208"  
##  [34,] "Qtom.A.LA.209"   "Qtom.A.LA.209"   "Qtom.A.LA.209"   "Qtom.A.LA.209"  
##  [35,] "Qtom.A.LA.218"   "Qtom.A.LA.218"   "Qtom.A.LA.218"   "Qtom.A.LA.218"  
##  [36,] "Qtom.A.LA.221"   "Qtom.A.LA.221"   "Qtom.A.LA.221"   "Qtom.A.LA.221"  
##  [37,] "Qtom.A.SB.324"   "Qtom.A.SB.324"   "Qtom.A.SB.324"   "Qtom.A.SB.324"  
##  [38,] "Qtom.A.SB.325"   "Qtom.A.SB.325"   "Qtom.A.SB.325"   "Qtom.A.SB.325"  
##  [39,] "Qtom.A.SB.327"   "Qtom.A.SB.327"   "Qtom.A.SB.327"   "Qtom.A.SB.327"  
##  [40,] "Qtom.A.SB.329"   "Qtom.A.SB.329"   "Qtom.A.SB.329"   "Qtom.A.SB.329"  
##  [41,] "Qtom.A.SB.330"   "Qtom.A.SB.330"   "Qtom.A.SB.330"   "Qtom.A.SB.330"  
##  [42,] "Qtom.A.SB.331"   "Qtom.A.SB.331"   "Qtom.A.SB.331"   "Qtom.A.SB.331"  
##  [43,] "Qtom.A.SB.332"   "Qtom.A.SB.332"   "Qtom.A.SB.332"   "Qtom.A.SB.332"  
##  [44,] "Qtom.A.SB.333"   "Qtom.A.SB.333"   "Qtom.A.SB.333"   "Qtom.A.SB.333"  
##  [45,] "Qtom.A.SB.335"   "Qtom.A.SB.335"   "Qtom.A.SB.335"   "Qtom.A.SB.335"  
##  [46,] "Qtom.A.SB.336"   "Qtom.A.SB.336"   "Qtom.A.SB.336"   "Qtom.A.SB.336"  
##  [47,] "Qtom.A.SB.337"   "Qtom.A.SB.337"   "Qtom.A.SB.337"   "Qtom.A.SB.337"  
##  [48,] "Qtom.A.SB.338"   "Qtom.A.SB.338"   "Qtom.A.SB.338"   "Qtom.A.SB.338"  
##  [49,] "Qtom.A.SB.339"   "Qtom.A.SB.339"   "Qtom.A.SB.339"   "Qtom.A.SB.339"  
##  [50,] "Qtom.A.SB.341"   "Qtom.A.SB.341"   "Qtom.A.SB.341"   "Qtom.A.SB.341"  
##  [51,] "Qtom.A.SB.342"   "Qtom.A.SB.342"   "Qtom.A.SB.342"   "Qtom.A.SB.342"  
##  [52,] "Qtom.A.SB.343"   "Qtom.A.SB.343"   "Qtom.A.SB.343"   "Qtom.A.SB.343"  
##  [53,] "Qtom.A.SB.344"   "Qtom.A.SB.344"   "Qtom.A.SB.344"   "Qtom.A.SB.344"  
##  [54,] "Qtom.A.SB.345"   "Qtom.A.SB.345"   "Qtom.A.SB.345"   "Qtom.A.SB.345"  
##  [55,] "Qtom.A.SB.346"   "Qtom.A.SB.346"   "Qtom.A.SB.346"   "Qtom.A.SB.346"  
##  [56,] "Qtom.A.SB.347"   "Qtom.A.SB.347"   "Qtom.A.SB.347"   "Qtom.A.SB.347"  
##  [57,] "Qtom.A.SB.348"   "Qtom.A.SB.348"   "Qtom.A.SB.348"   "Qtom.A.SB.348"  
##  [58,] "Qtom.A.SB.352"   "Qtom.A.SB.352"   "Qtom.A.SB.352"   "Qtom.A.SB.352"  
##  [59,] "Qtom.A.SB.353"   "Qtom.A.SB.353"   "Qtom.A.SB.353"   "Qtom.A.SB.353"  
##  [60,] "Qtom.A.SB.355"   "Qtom.A.SB.355"   "Qtom.A.SB.355"   "Qtom.A.SB.355"  
##  [61,] "Qtom.I.BC.10"    "Qtom.I.BC.10"    "Qtom.I.BC.10"    "Qtom.I.BC.10"   
##  [62,] "Qtom.I.BC.11"    "Qtom.I.BC.11"    "Qtom.I.BC.11"    "Qtom.I.BC.11"   
##  [63,] "Qtom.I.BC.12"    "Qtom.I.BC.12"    "Qtom.I.BC.12"    "Qtom.I.BC.12"   
##  [64,] "Qtom.I.BC.13"    "Qtom.I.BC.13"    "Qtom.I.BC.13"    "Qtom.I.BC.13"   
##  [65,] "Qtom.I.BC.18"    "Qtom.I.BC.18"    "Qtom.I.BC.18"    "Qtom.I.BC.18"   
##  [66,] "Qtom.I.BC.20"    "Qtom.I.BC.20"    "Qtom.I.BC.20"    "Qtom.I.BC.20"   
##  [67,] "Qtom.I.BC.21"    "Qtom.I.BC.21"    "Qtom.I.BC.21"    "Qtom.I.BC.21"   
##  [68,] "Qtom.I.BC.23"    "Qtom.I.BC.23"    "Qtom.I.BC.23"    "Qtom.I.BC.23"   
##  [69,] "Qtom.I.BC.24"    "Qtom.I.BC.24"    "Qtom.I.BC.24"    "Qtom.I.BC.24"   
##  [70,] "Qtom.I.BC.32"    "Qtom.I.BC.32"    "Qtom.I.BC.32"    "Qtom.I.BC.32"   
##  [71,] "Qtom.I.BC.33"    "Qtom.I.BC.33"    "Qtom.I.BC.33"    "Qtom.I.BC.33"   
##  [72,] "Qtom.I.BC.35"    "Qtom.I.BC.35"    "Qtom.I.BC.35"    "Qtom.I.BC.35"   
##  [73,] "Qtom.I.BC.39"    "Qtom.I.BC.39"    "Qtom.I.BC.39"    "Qtom.I.BC.39"   
##  [74,] "Qtom.I.BC.4"     "Qtom.I.BC.4"     "Qtom.I.BC.4"     "Qtom.I.BC.4"    
##  [75,] "Qtom.I.BC.40"    "Qtom.I.BC.40"    "Qtom.I.BC.40"    "Qtom.I.BC.40"   
##  [76,] "Qtom.I.BC.41"    "Qtom.I.BC.41"    "Qtom.I.BC.41"    "Qtom.I.BC.41"   
##  [77,] "Qtom.I.BC.42"    "Qtom.I.BC.42"    "Qtom.I.BC.42"    "Qtom.I.BC.42"   
##  [78,] "Qtom.I.BC.43"    "Qtom.I.BC.43"    "Qtom.I.BC.43"    "Qtom.I.BC.43"   
##  [79,] "Qtom.I.BC.44"    "Qtom.I.BC.44"    "Qtom.I.BC.44"    "Qtom.I.BC.44"   
##  [80,] "Qtom.I.BC.45"    "Qtom.I.BC.45"    "Qtom.I.BC.45"    "Qtom.I.BC.45"   
##  [81,] "Qtom.I.BC.46"    "Qtom.I.BC.46"    "Qtom.I.BC.46"    "Qtom.I.BC.46"   
##  [82,] "Qtom.I.BC.48"    "Qtom.I.BC.48"    "Qtom.I.BC.48"    "Qtom.I.BC.48"   
##  [83,] "Qtom.I.LA.SC.01" "Qtom.I.LA.SC.01" "Qtom.I.LA.SC.01" "Qtom.I.LA.SC.01"
##  [84,] "Qtom.I.LA.05"    "Qtom.I.LA.05"    "Qtom.I.LA.05"    "Qtom.I.LA.05"   
##  [85,] "Qtom.I.LA.06"    "Qtom.I.LA.06"    "Qtom.I.LA.06"    "Qtom.I.LA.06"   
##  [86,] "Qtom.I.LA.08"    "Qtom.I.LA.08"    "Qtom.I.LA.08"    "Qtom.I.LA.08"   
##  [87,] "Qtom.I.LA.10"    "Qtom.I.LA.10"    "Qtom.I.LA.10"    "Qtom.I.LA.10"   
##  [88,] "Qtom.I.LA.11"    "Qtom.I.LA.11"    "Qtom.I.LA.11"    "Qtom.I.LA.11"   
##  [89,] "Qtom.I.LA.12"    "Qtom.I.LA.12"    "Qtom.I.LA.12"    "Qtom.I.LA.12"   
##  [90,] "Qtom.I.LA.13"    "Qtom.I.LA.13"    "Qtom.I.LA.13"    "Qtom.I.LA.13"   
##  [91,] "Qtom.I.LA.21"    "Qtom.I.LA.21"    "Qtom.I.LA.21"    "Qtom.I.LA.21"   
##  [92,] "Qtom.I.LA.22"    "Qtom.I.LA.22"    "Qtom.I.LA.22"    "Qtom.I.LA.22"   
##  [93,] "Qtom.I.LA.23"    "Qtom.I.LA.23"    "Qtom.I.LA.23"    "Qtom.I.LA.23"   
##  [94,] "Qtom.I.LA.24"    "Qtom.I.LA.24"    "Qtom.I.LA.24"    "Qtom.I.LA.24"   
##  [95,] "Qtom.I.SB.01"    "Qtom.I.SB.01"    "Qtom.I.SB.01"    "Qtom.I.SB.01"   
##  [96,] "Qtom.I.SB.04"    "Qtom.I.SB.04"    "Qtom.I.SB.04"    "Qtom.I.SB.04"   
##  [97,] "Qtom.I.SB.09"    "Qtom.I.SB.09"    "Qtom.I.SB.09"    "Qtom.I.SB.09"   
##  [98,] "Qtom.I.SB.10"    "Qtom.I.SB.10"    "Qtom.I.SB.10"    "Qtom.I.SB.10"   
##  [99,] "Qtom.I.SB.11"    "Qtom.I.SB.11"    "Qtom.I.SB.11"    "Qtom.I.SB.11"   
## [100,] "Qtom.I.SB.12"    "Qtom.I.SB.12"    "Qtom.I.SB.12"    "Qtom.I.SB.12"   
## [101,] "Qtom.I.SB.28"    "Qtom.I.SB.28"    "Qtom.I.SB.28"    "Qtom.I.SB.28"   
## [102,] "Qtom.I.SB.33"    "Qtom.I.SB.33"    "Qtom.I.SB.33"    "Qtom.I.SB.33"   
## [103,] "Qtom.I.SB.35"    "Qtom.I.SB.35"    "Qtom.I.SB.35"    "Qtom.I.SB.35"   
## [104,] "Qtom.I.SB.36"    "Qtom.I.SB.36"    "Qtom.I.SB.36"    "Qtom.I.SB.36"   
## [105,] "Qtom.I.SB.37"    "Qtom.I.SB.37"    "Qtom.I.SB.37"    "Qtom.I.SB.37"   
## [106,] "Qtom.I.SB.38"    "Qtom.I.SB.38"    "Qtom.I.SB.38"    "Qtom.I.SB.38"   
## [107,] "Qtom.I.SB.40"    "Qtom.I.SB.40"    "Qtom.I.SB.40"    "Qtom.I.SB.40"   
## [108,] "Qtom.I.SB.41"    "Qtom.I.SB.41"    "Qtom.I.SB.41"    "Qtom.I.SB.41"   
## [109,] "Qtom.I.SB.42"    "Qtom.I.SB.42"    "Qtom.I.SB.42"    "Qtom.I.SB.42"   
## [110,] "Qtom.I.SB.44"    "Qtom.I.SB.44"    "Qtom.I.SB.44"    "Qtom.I.SB.44"   
## [111,] "Qtom.I.SB.45"    "Qtom.I.SB.45"    "Qtom.I.SB.45"    "Qtom.I.SB.45"   
## [112,] "Qtom.I.SB.73"    "Qtom.I.SB.73"    "Qtom.I.SB.73"    "Qtom.I.SB.73"   
## [113,] "Qtom.I.SB.74"    "Qtom.I.SB.74"    "Qtom.I.SB.74"    "Qtom.I.SB.74"   
## [114,] "Qtom.I.VEN.01"   "Qtom.I.VEN.01"   "Qtom.I.VEN.01"   "Qtom.I.VEN.01"  
## [115,] "Qtom.I.VEN.02"   "Qtom.I.VEN.02"   "Qtom.I.VEN.02"   "Qtom.I.VEN.02"  
## [116,] "Qtom.I.VEN.03"   "Qtom.I.VEN.03"   "Qtom.I.VEN.03"   "Qtom.I.VEN.03"  
## [117,] "Qtom.I.VEN.04"   "Qtom.I.VEN.04"   "Qtom.I.VEN.04"   "Qtom.I.VEN.04"  
## [118,] "Qtom.I.VEN.05"   "Qtom.I.VEN.05"   "Qtom.I.VEN.05"   "Qtom.I.VEN.05"  
## [119,] "Qtom.JJK.SB.01"  "Qtom.JJK.SB.01"  "Qtom.JJK.SB.01"  "Qtom.JJK.SB.01" 
## [120,] "Qtom.JJK.SB.03"  "Qtom.JJK.SB.03"  "Qtom.JJK.SB.03"  "Qtom.JJK.SB.03" 
## [121,] "Qtom.JJK.SB.07"  "Qtom.JJK.SB.07"  "Qtom.JJK.SB.07"  "Qtom.JJK.SB.07" 
## [122,] "Qtom.JJK.SB.08"  "Qtom.JJK.SB.08"  "Qtom.JJK.SB.08"  "Qtom.JJK.SB.08" 
## [123,] "Qtom.JJK.SB.09"  "Qtom.JJK.SB.09"  "Qtom.JJK.SB.09"  "Qtom.JJK.SB.09" 
## [124,] "Qtom.JJK.SB.10"  "Qtom.JJK.SB.10"  "Qtom.JJK.SB.10"  "Qtom.JJK.SB.10" 
## [125,] "Qtom.JR.SB.01"   "Qtom.JR.SB.01"   "Qtom.JR.SB.01"   "Qtom.JR.SB.01"  
## [126,] "Qtom.S.LA.186"   "Qtom.S.LA.186"   "Qtom.S.LA.186"   "Qtom.S.LA.186"
```

```
# rename
snps <- tmp.snps
imp <- tmp.imp

cbind(rownames(info), rownames(clim), rownames(snps), rownames(imp))
```

```
##        [,1]              [,2]              [,3]              [,4]             
##   [1,] "Qchr.A.232"      "Qchr.A.232"      "Qchr.A.232"      "Qchr.A.232"     
##   [2,] "Qchr.A.275"      "Qchr.A.275"      "Qchr.A.275"      "Qchr.A.275"     
##   [3,] "Qchr.A.ED.102"   "Qchr.A.ED.102"   "Qchr.A.ED.102"   "Qchr.A.ED.102"  
##   [4,] "Qchr.A.ELD.53"   "Qchr.A.ELD.53"   "Qchr.A.ELD.53"   "Qchr.A.ELD.53"  
##   [5,] "Qchr.A.KER.16"   "Qchr.A.KER.16"   "Qchr.A.KER.16"   "Qchr.A.KER.16"  
##   [6,] "Qchr.A.LA.02"    "Qchr.A.LA.02"    "Qchr.A.LA.02"    "Qchr.A.LA.02"   
##   [7,] "Qchr.A.LA.215"   "Qchr.A.LA.215"   "Qchr.A.LA.215"   "Qchr.A.LA.215"  
##   [8,] "Qchr.A.LA.222"   "Qchr.A.LA.222"   "Qchr.A.LA.222"   "Qchr.A.LA.222"  
##   [9,] "Qchr.A.SIS.309"  "Qchr.A.SIS.309"  "Qchr.A.SIS.309"  "Qchr.A.SIS.309" 
##  [10,] "Qchr.A.PLU.65"   "Qchr.A.PLU.65"   "Qchr.A.PLU.65"   "Qchr.A.PLU.65"  
##  [11,] "Qchr.A.SBD.83"   "Qchr.A.SBD.83"   "Qchr.A.SBD.83"   "Qchr.A.SBD.83"  
##  [12,] "Qchr.I.LA.20"    "Qchr.I.LA.20"    "Qchr.I.LA.20"    "Qchr.I.LA.20"   
##  [13,] "Qchr.I.LA.25"    "Qchr.I.LA.25"    "Qchr.I.LA.25"    "Qchr.I.LA.25"   
##  [14,] "Qchr.I.SB.02"    "Qchr.I.SB.02"    "Qchr.I.SB.02"    "Qchr.I.SB.02"   
##  [15,] "Qchr.I.SB.03"    "Qchr.I.SB.03"    "Qchr.I.SB.03"    "Qchr.I.SB.03"   
##  [16,] "Qchr.JJK.SB.13"  "Qchr.JJK.SB.13"  "Qchr.JJK.SB.13"  "Qchr.JJK.SB.13" 
##  [17,] "Qchr.JR.SB.01"   "Qchr.JR.SB.01"   "Qchr.JR.SB.01"   "Qchr.JR.SB.01"  
##  [18,] "Qspp.I.LA.03"    "Qspp.I.LA.03"    "Qspp.I.LA.03"    "Qspp.I.LA.03"   
##  [19,] "Qspp.I.LA.04"    "Qspp.I.LA.04"    "Qspp.I.LA.04"    "Qspp.I.LA.04"   
##  [20,] "Qspp.I.LA.14"    "Qspp.I.LA.14"    "Qspp.I.LA.14"    "Qspp.I.LA.14"   
##  [21,] "Qtom.A.LA.169"   "Qtom.A.LA.169"   "Qtom.A.LA.169"   "Qtom.A.LA.169"  
##  [22,] "Qtom.A.LA.175"   "Qtom.A.LA.175"   "Qtom.A.LA.175"   "Qtom.A.LA.175"  
##  [23,] "Qtom.A.LA.176"   "Qtom.A.LA.176"   "Qtom.A.LA.176"   "Qtom.A.LA.176"  
##  [24,] "Qtom.A.LA.180"   "Qtom.A.LA.180"   "Qtom.A.LA.180"   "Qtom.A.LA.180"  
##  [25,] "Qtom.A.LA.181"   "Qtom.A.LA.181"   "Qtom.A.LA.181"   "Qtom.A.LA.181"  
##  [26,] "Qtom.A.LA.183"   "Qtom.A.LA.183"   "Qtom.A.LA.183"   "Qtom.A.LA.183"  
##  [27,] "Qtom.A.LA.190"   "Qtom.A.LA.190"   "Qtom.A.LA.190"   "Qtom.A.LA.190"  
##  [28,] "Qtom.A.LA.191"   "Qtom.A.LA.191"   "Qtom.A.LA.191"   "Qtom.A.LA.191"  
##  [29,] "Qtom.A.LA.198"   "Qtom.A.LA.198"   "Qtom.A.LA.198"   "Qtom.A.LA.198"  
##  [30,] "Qtom.A.LA.200"   "Qtom.A.LA.200"   "Qtom.A.LA.200"   "Qtom.A.LA.200"  
##  [31,] "Qtom.A.LA.204"   "Qtom.A.LA.204"   "Qtom.A.LA.204"   "Qtom.A.LA.204"  
##  [32,] "Qtom.A.LA.207"   "Qtom.A.LA.207"   "Qtom.A.LA.207"   "Qtom.A.LA.207"  
##  [33,] "Qtom.A.LA.208"   "Qtom.A.LA.208"   "Qtom.A.LA.208"   "Qtom.A.LA.208"  
##  [34,] "Qtom.A.LA.209"   "Qtom.A.LA.209"   "Qtom.A.LA.209"   "Qtom.A.LA.209"  
##  [35,] "Qtom.A.LA.218"   "Qtom.A.LA.218"   "Qtom.A.LA.218"   "Qtom.A.LA.218"  
##  [36,] "Qtom.A.LA.221"   "Qtom.A.LA.221"   "Qtom.A.LA.221"   "Qtom.A.LA.221"  
##  [37,] "Qtom.A.SB.324"   "Qtom.A.SB.324"   "Qtom.A.SB.324"   "Qtom.A.SB.324"  
##  [38,] "Qtom.A.SB.325"   "Qtom.A.SB.325"   "Qtom.A.SB.325"   "Qtom.A.SB.325"  
##  [39,] "Qtom.A.SB.327"   "Qtom.A.SB.327"   "Qtom.A.SB.327"   "Qtom.A.SB.327"  
##  [40,] "Qtom.A.SB.329"   "Qtom.A.SB.329"   "Qtom.A.SB.329"   "Qtom.A.SB.329"  
##  [41,] "Qtom.A.SB.330"   "Qtom.A.SB.330"   "Qtom.A.SB.330"   "Qtom.A.SB.330"  
##  [42,] "Qtom.A.SB.331"   "Qtom.A.SB.331"   "Qtom.A.SB.331"   "Qtom.A.SB.331"  
##  [43,] "Qtom.A.SB.332"   "Qtom.A.SB.332"   "Qtom.A.SB.332"   "Qtom.A.SB.332"  
##  [44,] "Qtom.A.SB.333"   "Qtom.A.SB.333"   "Qtom.A.SB.333"   "Qtom.A.SB.333"  
##  [45,] "Qtom.A.SB.335"   "Qtom.A.SB.335"   "Qtom.A.SB.335"   "Qtom.A.SB.335"  
##  [46,] "Qtom.A.SB.336"   "Qtom.A.SB.336"   "Qtom.A.SB.336"   "Qtom.A.SB.336"  
##  [47,] "Qtom.A.SB.337"   "Qtom.A.SB.337"   "Qtom.A.SB.337"   "Qtom.A.SB.337"  
##  [48,] "Qtom.A.SB.338"   "Qtom.A.SB.338"   "Qtom.A.SB.338"   "Qtom.A.SB.338"  
##  [49,] "Qtom.A.SB.339"   "Qtom.A.SB.339"   "Qtom.A.SB.339"   "Qtom.A.SB.339"  
##  [50,] "Qtom.A.SB.341"   "Qtom.A.SB.341"   "Qtom.A.SB.341"   "Qtom.A.SB.341"  
##  [51,] "Qtom.A.SB.342"   "Qtom.A.SB.342"   "Qtom.A.SB.342"   "Qtom.A.SB.342"  
##  [52,] "Qtom.A.SB.343"   "Qtom.A.SB.343"   "Qtom.A.SB.343"   "Qtom.A.SB.343"  
##  [53,] "Qtom.A.SB.344"   "Qtom.A.SB.344"   "Qtom.A.SB.344"   "Qtom.A.SB.344"  
##  [54,] "Qtom.A.SB.345"   "Qtom.A.SB.345"   "Qtom.A.SB.345"   "Qtom.A.SB.345"  
##  [55,] "Qtom.A.SB.346"   "Qtom.A.SB.346"   "Qtom.A.SB.346"   "Qtom.A.SB.346"  
##  [56,] "Qtom.A.SB.347"   "Qtom.A.SB.347"   "Qtom.A.SB.347"   "Qtom.A.SB.347"  
##  [57,] "Qtom.A.SB.348"   "Qtom.A.SB.348"   "Qtom.A.SB.348"   "Qtom.A.SB.348"  
##  [58,] "Qtom.A.SB.352"   "Qtom.A.SB.352"   "Qtom.A.SB.352"   "Qtom.A.SB.352"  
##  [59,] "Qtom.A.SB.353"   "Qtom.A.SB.353"   "Qtom.A.SB.353"   "Qtom.A.SB.353"  
##  [60,] "Qtom.A.SB.355"   "Qtom.A.SB.355"   "Qtom.A.SB.355"   "Qtom.A.SB.355"  
##  [61,] "Qtom.I.BC.10"    "Qtom.I.BC.10"    "Qtom.I.BC.10"    "Qtom.I.BC.10"   
##  [62,] "Qtom.I.BC.11"    "Qtom.I.BC.11"    "Qtom.I.BC.11"    "Qtom.I.BC.11"   
##  [63,] "Qtom.I.BC.12"    "Qtom.I.BC.12"    "Qtom.I.BC.12"    "Qtom.I.BC.12"   
##  [64,] "Qtom.I.BC.13"    "Qtom.I.BC.13"    "Qtom.I.BC.13"    "Qtom.I.BC.13"   
##  [65,] "Qtom.I.BC.18"    "Qtom.I.BC.18"    "Qtom.I.BC.18"    "Qtom.I.BC.18"   
##  [66,] "Qtom.I.BC.20"    "Qtom.I.BC.20"    "Qtom.I.BC.20"    "Qtom.I.BC.20"   
##  [67,] "Qtom.I.BC.21"    "Qtom.I.BC.21"    "Qtom.I.BC.21"    "Qtom.I.BC.21"   
##  [68,] "Qtom.I.BC.23"    "Qtom.I.BC.23"    "Qtom.I.BC.23"    "Qtom.I.BC.23"   
##  [69,] "Qtom.I.BC.24"    "Qtom.I.BC.24"    "Qtom.I.BC.24"    "Qtom.I.BC.24"   
##  [70,] "Qtom.I.BC.32"    "Qtom.I.BC.32"    "Qtom.I.BC.32"    "Qtom.I.BC.32"   
##  [71,] "Qtom.I.BC.33"    "Qtom.I.BC.33"    "Qtom.I.BC.33"    "Qtom.I.BC.33"   
##  [72,] "Qtom.I.BC.35"    "Qtom.I.BC.35"    "Qtom.I.BC.35"    "Qtom.I.BC.35"   
##  [73,] "Qtom.I.BC.39"    "Qtom.I.BC.39"    "Qtom.I.BC.39"    "Qtom.I.BC.39"   
##  [74,] "Qtom.I.BC.4"     "Qtom.I.BC.4"     "Qtom.I.BC.4"     "Qtom.I.BC.4"    
##  [75,] "Qtom.I.BC.40"    "Qtom.I.BC.40"    "Qtom.I.BC.40"    "Qtom.I.BC.40"   
##  [76,] "Qtom.I.BC.41"    "Qtom.I.BC.41"    "Qtom.I.BC.41"    "Qtom.I.BC.41"   
##  [77,] "Qtom.I.BC.42"    "Qtom.I.BC.42"    "Qtom.I.BC.42"    "Qtom.I.BC.42"   
##  [78,] "Qtom.I.BC.43"    "Qtom.I.BC.43"    "Qtom.I.BC.43"    "Qtom.I.BC.43"   
##  [79,] "Qtom.I.BC.44"    "Qtom.I.BC.44"    "Qtom.I.BC.44"    "Qtom.I.BC.44"   
##  [80,] "Qtom.I.BC.45"    "Qtom.I.BC.45"    "Qtom.I.BC.45"    "Qtom.I.BC.45"   
##  [81,] "Qtom.I.BC.46"    "Qtom.I.BC.46"    "Qtom.I.BC.46"    "Qtom.I.BC.46"   
##  [82,] "Qtom.I.BC.48"    "Qtom.I.BC.48"    "Qtom.I.BC.48"    "Qtom.I.BC.48"   
##  [83,] "Qtom.I.LA.SC.01" "Qtom.I.LA.SC.01" "Qtom.I.LA.SC.01" "Qtom.I.LA.SC.01"
##  [84,] "Qtom.I.LA.05"    "Qtom.I.LA.05"    "Qtom.I.LA.05"    "Qtom.I.LA.05"   
##  [85,] "Qtom.I.LA.06"    "Qtom.I.LA.06"    "Qtom.I.LA.06"    "Qtom.I.LA.06"   
##  [86,] "Qtom.I.LA.08"    "Qtom.I.LA.08"    "Qtom.I.LA.08"    "Qtom.I.LA.08"   
##  [87,] "Qtom.I.LA.10"    "Qtom.I.LA.10"    "Qtom.I.LA.10"    "Qtom.I.LA.10"   
##  [88,] "Qtom.I.LA.11"    "Qtom.I.LA.11"    "Qtom.I.LA.11"    "Qtom.I.LA.11"   
##  [89,] "Qtom.I.LA.12"    "Qtom.I.LA.12"    "Qtom.I.LA.12"    "Qtom.I.LA.12"   
##  [90,] "Qtom.I.LA.13"    "Qtom.I.LA.13"    "Qtom.I.LA.13"    "Qtom.I.LA.13"   
##  [91,] "Qtom.I.LA.21"    "Qtom.I.LA.21"    "Qtom.I.LA.21"    "Qtom.I.LA.21"   
##  [92,] "Qtom.I.LA.22"    "Qtom.I.LA.22"    "Qtom.I.LA.22"    "Qtom.I.LA.22"   
##  [93,] "Qtom.I.LA.23"    "Qtom.I.LA.23"    "Qtom.I.LA.23"    "Qtom.I.LA.23"   
##  [94,] "Qtom.I.LA.24"    "Qtom.I.LA.24"    "Qtom.I.LA.24"    "Qtom.I.LA.24"   
##  [95,] "Qtom.I.SB.01"    "Qtom.I.SB.01"    "Qtom.I.SB.01"    "Qtom.I.SB.01"   
##  [96,] "Qtom.I.SB.04"    "Qtom.I.SB.04"    "Qtom.I.SB.04"    "Qtom.I.SB.04"   
##  [97,] "Qtom.I.SB.09"    "Qtom.I.SB.09"    "Qtom.I.SB.09"    "Qtom.I.SB.09"   
##  [98,] "Qtom.I.SB.10"    "Qtom.I.SB.10"    "Qtom.I.SB.10"    "Qtom.I.SB.10"   
##  [99,] "Qtom.I.SB.11"    "Qtom.I.SB.11"    "Qtom.I.SB.11"    "Qtom.I.SB.11"   
## [100,] "Qtom.I.SB.12"    "Qtom.I.SB.12"    "Qtom.I.SB.12"    "Qtom.I.SB.12"   
## [101,] "Qtom.I.SB.28"    "Qtom.I.SB.28"    "Qtom.I.SB.28"    "Qtom.I.SB.28"   
## [102,] "Qtom.I.SB.33"    "Qtom.I.SB.33"    "Qtom.I.SB.33"    "Qtom.I.SB.33"   
## [103,] "Qtom.I.SB.35"    "Qtom.I.SB.35"    "Qtom.I.SB.35"    "Qtom.I.SB.35"   
## [104,] "Qtom.I.SB.36"    "Qtom.I.SB.36"    "Qtom.I.SB.36"    "Qtom.I.SB.36"   
## [105,] "Qtom.I.SB.37"    "Qtom.I.SB.37"    "Qtom.I.SB.37"    "Qtom.I.SB.37"   
## [106,] "Qtom.I.SB.38"    "Qtom.I.SB.38"    "Qtom.I.SB.38"    "Qtom.I.SB.38"   
## [107,] "Qtom.I.SB.40"    "Qtom.I.SB.40"    "Qtom.I.SB.40"    "Qtom.I.SB.40"   
## [108,] "Qtom.I.SB.41"    "Qtom.I.SB.41"    "Qtom.I.SB.41"    "Qtom.I.SB.41"   
## [109,] "Qtom.I.SB.42"    "Qtom.I.SB.42"    "Qtom.I.SB.42"    "Qtom.I.SB.42"   
## [110,] "Qtom.I.SB.44"    "Qtom.I.SB.44"    "Qtom.I.SB.44"    "Qtom.I.SB.44"   
## [111,] "Qtom.I.SB.45"    "Qtom.I.SB.45"    "Qtom.I.SB.45"    "Qtom.I.SB.45"   
## [112,] "Qtom.I.SB.73"    "Qtom.I.SB.73"    "Qtom.I.SB.73"    "Qtom.I.SB.73"   
## [113,] "Qtom.I.SB.74"    "Qtom.I.SB.74"    "Qtom.I.SB.74"    "Qtom.I.SB.74"   
## [114,] "Qtom.I.VEN.01"   "Qtom.I.VEN.01"   "Qtom.I.VEN.01"   "Qtom.I.VEN.01"  
## [115,] "Qtom.I.VEN.02"   "Qtom.I.VEN.02"   "Qtom.I.VEN.02"   "Qtom.I.VEN.02"  
## [116,] "Qtom.I.VEN.03"   "Qtom.I.VEN.03"   "Qtom.I.VEN.03"   "Qtom.I.VEN.03"  
## [117,] "Qtom.I.VEN.04"   "Qtom.I.VEN.04"   "Qtom.I.VEN.04"   "Qtom.I.VEN.04"  
## [118,] "Qtom.I.VEN.05"   "Qtom.I.VEN.05"   "Qtom.I.VEN.05"   "Qtom.I.VEN.05"  
## [119,] "Qtom.JJK.SB.01"  "Qtom.JJK.SB.01"  "Qtom.JJK.SB.01"  "Qtom.JJK.SB.01" 
## [120,] "Qtom.JJK.SB.03"  "Qtom.JJK.SB.03"  "Qtom.JJK.SB.03"  "Qtom.JJK.SB.03" 
## [121,] "Qtom.JJK.SB.07"  "Qtom.JJK.SB.07"  "Qtom.JJK.SB.07"  "Qtom.JJK.SB.07" 
## [122,] "Qtom.JJK.SB.08"  "Qtom.JJK.SB.08"  "Qtom.JJK.SB.08"  "Qtom.JJK.SB.08" 
## [123,] "Qtom.JJK.SB.09"  "Qtom.JJK.SB.09"  "Qtom.JJK.SB.09"  "Qtom.JJK.SB.09" 
## [124,] "Qtom.JJK.SB.10"  "Qtom.JJK.SB.10"  "Qtom.JJK.SB.10"  "Qtom.JJK.SB.10" 
## [125,] "Qtom.JR.SB.01"   "Qtom.JR.SB.01"   "Qtom.JR.SB.01"   "Qtom.JR.SB.01"  
## [126,] "Qtom.S.LA.186"   "Qtom.S.LA.186"   "Qtom.S.LA.186"   "Qtom.S.LA.186"
```

```
# first look at rda with climate
# including guadalupe to visualize climate

rda <- rda(imp ~ ., data = as.data.frame(clim), scale = T)
screeplot(rda)
```

```
plot_clim(rda, info = info, save = F, file_string = 'Qtom107.Qchr17.Qssp3')
```

```
plot_clim(rda, info = info, choices = c(1,3), save = F, file_string = 'Qtom107.Qchr17.Qssp3')
```

#### 3.2 Basic RDA, visualize without Guadalupe

```
# do same plot without Guadalupe

# get subset 
clim.sub <- clim[info$island != 'Guadalupe Island',]
snps.sub <- snps[info$island != 'Guadalupe Island',]
imp.sub <- imp[info$island != 'Guadalupe Island',]
info.sub <- info[info$island != 'Guadalupe Island',]

rda <- rda(imp.sub ~ ., data = as.data.frame(clim.sub), scale = T)
screeplot(rda)
```

```
plot_clim(rda, info = info.sub, save = F, file_string = 'Qtom107.Qchr17.Qssp3_noGuadalupe')
```

```
plot_clim(rda, info = info.sub, choices = c(1,3), save = F, file_string = 'Qtom107.Qchr17.Qssp3_noGuadalupe')
```

#### 3.3 Basic RDA, visualize without Guadalupe or Mainland Qchr

```
# same plot without Guadalupe or mainland Q. chrysolepis, only California Channel Islands

# get subset without guadalupe or mainland Qchr
clim.sub <- clim[! info$island %in% c('Guadalupe Island', "Mainland"),]
snps.sub <- snps[! info$island %in% c('Guadalupe Island', "Mainland"),]
imp.sub <- imp[! info$island %in% c('Guadalupe Island', "Mainland"),]
info.sub <- info[! info$island %in% c('Guadalupe Island', "Mainland"),]

cbind(rownames(info.sub), rownames(snps.sub), rownames(imp.sub), rownames(clim.sub))
```

```
##       [,1]              [,2]              [,3]              [,4]             
##  [1,] "Qchr.A.LA.215"   "Qchr.A.LA.215"   "Qchr.A.LA.215"   "Qchr.A.LA.215"  
##  [2,] "Qchr.A.LA.222"   "Qchr.A.LA.222"   "Qchr.A.LA.222"   "Qchr.A.LA.222"  
##  [3,] "Qchr.I.LA.20"    "Qchr.I.LA.20"    "Qchr.I.LA.20"    "Qchr.I.LA.20"   
##  [4,] "Qchr.I.LA.25"    "Qchr.I.LA.25"    "Qchr.I.LA.25"    "Qchr.I.LA.25"   
##  [5,] "Qchr.I.SB.02"    "Qchr.I.SB.02"    "Qchr.I.SB.02"    "Qchr.I.SB.02"   
##  [6,] "Qchr.I.SB.03"    "Qchr.I.SB.03"    "Qchr.I.SB.03"    "Qchr.I.SB.03"   
##  [7,] "Qchr.JJK.SB.13"  "Qchr.JJK.SB.13"  "Qchr.JJK.SB.13"  "Qchr.JJK.SB.13" 
##  [8,] "Qchr.JR.SB.01"   "Qchr.JR.SB.01"   "Qchr.JR.SB.01"   "Qchr.JR.SB.01"  
##  [9,] "Qspp.I.LA.03"    "Qspp.I.LA.03"    "Qspp.I.LA.03"    "Qspp.I.LA.03"   
## [10,] "Qspp.I.LA.04"    "Qspp.I.LA.04"    "Qspp.I.LA.04"    "Qspp.I.LA.04"   
## [11,] "Qspp.I.LA.14"    "Qspp.I.LA.14"    "Qspp.I.LA.14"    "Qspp.I.LA.14"   
## [12,] "Qtom.A.LA.169"   "Qtom.A.LA.169"   "Qtom.A.LA.169"   "Qtom.A.LA.169"  
## [13,] "Qtom.A.LA.175"   "Qtom.A.LA.175"   "Qtom.A.LA.175"   "Qtom.A.LA.175"  
## [14,] "Qtom.A.LA.176"   "Qtom.A.LA.176"   "Qtom.A.LA.176"   "Qtom.A.LA.176"  
## [15,] "Qtom.A.LA.180"   "Qtom.A.LA.180"   "Qtom.A.LA.180"   "Qtom.A.LA.180"  
## [16,] "Qtom.A.LA.181"   "Qtom.A.LA.181"   "Qtom.A.LA.181"   "Qtom.A.LA.181"  
## [17,] "Qtom.A.LA.183"   "Qtom.A.LA.183"   "Qtom.A.LA.183"   "Qtom.A.LA.183"  
## [18,] "Qtom.A.LA.190"   "Qtom.A.LA.190"   "Qtom.A.LA.190"   "Qtom.A.LA.190"  
## [19,] "Qtom.A.LA.191"   "Qtom.A.LA.191"   "Qtom.A.LA.191"   "Qtom.A.LA.191"  
## [20,] "Qtom.A.LA.198"   "Qtom.A.LA.198"   "Qtom.A.LA.198"   "Qtom.A.LA.198"  
## [21,] "Qtom.A.LA.200"   "Qtom.A.LA.200"   "Qtom.A.LA.200"   "Qtom.A.LA.200"  
## [22,] "Qtom.A.LA.204"   "Qtom.A.LA.204"   "Qtom.A.LA.204"   "Qtom.A.LA.204"  
## [23,] "Qtom.A.LA.207"   "Qtom.A.LA.207"   "Qtom.A.LA.207"   "Qtom.A.LA.207"  
## [24,] "Qtom.A.LA.208"   "Qtom.A.LA.208"   "Qtom.A.LA.208"   "Qtom.A.LA.208"  
## [25,] "Qtom.A.LA.209"   "Qtom.A.LA.209"   "Qtom.A.LA.209"   "Qtom.A.LA.209"  
## [26,] "Qtom.A.LA.218"   "Qtom.A.LA.218"   "Qtom.A.LA.218"   "Qtom.A.LA.218"  
## [27,] "Qtom.A.LA.221"   "Qtom.A.LA.221"   "Qtom.A.LA.221"   "Qtom.A.LA.221"  
## [28,] "Qtom.A.SB.324"   "Qtom.A.SB.324"   "Qtom.A.SB.324"   "Qtom.A.SB.324"  
## [29,] "Qtom.A.SB.325"   "Qtom.A.SB.325"   "Qtom.A.SB.325"   "Qtom.A.SB.325"  
## [30,] "Qtom.A.SB.327"   "Qtom.A.SB.327"   "Qtom.A.SB.327"   "Qtom.A.SB.327"  
## [31,] "Qtom.A.SB.329"   "Qtom.A.SB.329"   "Qtom.A.SB.329"   "Qtom.A.SB.329"  
## [32,] "Qtom.A.SB.330"   "Qtom.A.SB.330"   "Qtom.A.SB.330"   "Qtom.A.SB.330"  
## [33,] "Qtom.A.SB.331"   "Qtom.A.SB.331"   "Qtom.A.SB.331"   "Qtom.A.SB.331"  
## [34,] "Qtom.A.SB.332"   "Qtom.A.SB.332"   "Qtom.A.SB.332"   "Qtom.A.SB.332"  
## [35,] "Qtom.A.SB.333"   "Qtom.A.SB.333"   "Qtom.A.SB.333"   "Qtom.A.SB.333"  
## [36,] "Qtom.A.SB.335"   "Qtom.A.SB.335"   "Qtom.A.SB.335"   "Qtom.A.SB.335"  
## [37,] "Qtom.A.SB.336"   "Qtom.A.SB.336"   "Qtom.A.SB.336"   "Qtom.A.SB.336"  
## [38,] "Qtom.A.SB.337"   "Qtom.A.SB.337"   "Qtom.A.SB.337"   "Qtom.A.SB.337"  
## [39,] "Qtom.A.SB.338"   "Qtom.A.SB.338"   "Qtom.A.SB.338"   "Qtom.A.SB.338"  
## [40,] "Qtom.A.SB.339"   "Qtom.A.SB.339"   "Qtom.A.SB.339"   "Qtom.A.SB.339"  
## [41,] "Qtom.A.SB.341"   "Qtom.A.SB.341"   "Qtom.A.SB.341"   "Qtom.A.SB.341"  
## [42,] "Qtom.A.SB.342"   "Qtom.A.SB.342"   "Qtom.A.SB.342"   "Qtom.A.SB.342"  
## [43,] "Qtom.A.SB.343"   "Qtom.A.SB.343"   "Qtom.A.SB.343"   "Qtom.A.SB.343"  
## [44,] "Qtom.A.SB.344"   "Qtom.A.SB.344"   "Qtom.A.SB.344"   "Qtom.A.SB.344"  
## [45,] "Qtom.A.SB.345"   "Qtom.A.SB.345"   "Qtom.A.SB.345"   "Qtom.A.SB.345"  
## [46,] "Qtom.A.SB.346"   "Qtom.A.SB.346"   "Qtom.A.SB.346"   "Qtom.A.SB.346"  
## [47,] "Qtom.A.SB.347"   "Qtom.A.SB.347"   "Qtom.A.SB.347"   "Qtom.A.SB.347"  
## [48,] "Qtom.A.SB.348"   "Qtom.A.SB.348"   "Qtom.A.SB.348"   "Qtom.A.SB.348"  
## [49,] "Qtom.A.SB.352"   "Qtom.A.SB.352"   "Qtom.A.SB.352"   "Qtom.A.SB.352"  
## [50,] "Qtom.A.SB.353"   "Qtom.A.SB.353"   "Qtom.A.SB.353"   "Qtom.A.SB.353"  
## [51,] "Qtom.A.SB.355"   "Qtom.A.SB.355"   "Qtom.A.SB.355"   "Qtom.A.SB.355"  
## [52,] "Qtom.I.LA.SC.01" "Qtom.I.LA.SC.01" "Qtom.I.LA.SC.01" "Qtom.I.LA.SC.01"
## [53,] "Qtom.I.LA.05"    "Qtom.I.LA.05"    "Qtom.I.LA.05"    "Qtom.I.LA.05"   
## [54,] "Qtom.I.LA.06"    "Qtom.I.LA.06"    "Qtom.I.LA.06"    "Qtom.I.LA.06"   
## [55,] "Qtom.I.LA.08"    "Qtom.I.LA.08"    "Qtom.I.LA.08"    "Qtom.I.LA.08"   
## [56,] "Qtom.I.LA.10"    "Qtom.I.LA.10"    "Qtom.I.LA.10"    "Qtom.I.LA.10"   
## [57,] "Qtom.I.LA.11"    "Qtom.I.LA.11"    "Qtom.I.LA.11"    "Qtom.I.LA.11"   
## [58,] "Qtom.I.LA.12"    "Qtom.I.LA.12"    "Qtom.I.LA.12"    "Qtom.I.LA.12"   
## [59,] "Qtom.I.LA.13"    "Qtom.I.LA.13"    "Qtom.I.LA.13"    "Qtom.I.LA.13"   
## [60,] "Qtom.I.LA.21"    "Qtom.I.LA.21"    "Qtom.I.LA.21"    "Qtom.I.LA.21"   
## [61,] "Qtom.I.LA.22"    "Qtom.I.LA.22"    "Qtom.I.LA.22"    "Qtom.I.LA.22"   
## [62,] "Qtom.I.LA.23"    "Qtom.I.LA.23"    "Qtom.I.LA.23"    "Qtom.I.LA.23"   
## [63,] "Qtom.I.LA.24"    "Qtom.I.LA.24"    "Qtom.I.LA.24"    "Qtom.I.LA.24"   
## [64,] "Qtom.I.SB.01"    "Qtom.I.SB.01"    "Qtom.I.SB.01"    "Qtom.I.SB.01"   
## [65,] "Qtom.I.SB.04"    "Qtom.I.SB.04"    "Qtom.I.SB.04"    "Qtom.I.SB.04"   
## [66,] "Qtom.I.SB.09"    "Qtom.I.SB.09"    "Qtom.I.SB.09"    "Qtom.I.SB.09"   
## [67,] "Qtom.I.SB.10"    "Qtom.I.SB.10"    "Qtom.I.SB.10"    "Qtom.I.SB.10"   
## [68,] "Qtom.I.SB.11"    "Qtom.I.SB.11"    "Qtom.I.SB.11"    "Qtom.I.SB.11"   
## [69,] "Qtom.I.SB.12"    "Qtom.I.SB.12"    "Qtom.I.SB.12"    "Qtom.I.SB.12"   
## [70,] "Qtom.I.SB.28"    "Qtom.I.SB.28"    "Qtom.I.SB.28"    "Qtom.I.SB.28"   
## [71,] "Qtom.I.SB.33"    "Qtom.I.SB.33"    "Qtom.I.SB.33"    "Qtom.I.SB.33"   
## [72,] "Qtom.I.SB.35"    "Qtom.I.SB.35"    "Qtom.I.SB.35"    "Qtom.I.SB.35"   
## [73,] "Qtom.I.SB.36"    "Qtom.I.SB.36"    "Qtom.I.SB.36"    "Qtom.I.SB.36"   
## [74,] "Qtom.I.SB.37"    "Qtom.I.SB.37"    "Qtom.I.SB.37"    "Qtom.I.SB.37"   
## [75,] "Qtom.I.SB.38"    "Qtom.I.SB.38"    "Qtom.I.SB.38"    "Qtom.I.SB.38"   
## [76,] "Qtom.I.SB.40"    "Qtom.I.SB.40"    "Qtom.I.SB.40"    "Qtom.I.SB.40"   
## [77,] "Qtom.I.SB.41"    "Qtom.I.SB.41"    "Qtom.I.SB.41"    "Qtom.I.SB.41"   
## [78,] "Qtom.I.SB.42"    "Qtom.I.SB.42"    "Qtom.I.SB.42"    "Qtom.I.SB.42"   
## [79,] "Qtom.I.SB.44"    "Qtom.I.SB.44"    "Qtom.I.SB.44"    "Qtom.I.SB.44"   
## [80,] "Qtom.I.SB.45"    "Qtom.I.SB.45"    "Qtom.I.SB.45"    "Qtom.I.SB.45"   
## [81,] "Qtom.I.SB.73"    "Qtom.I.SB.73"    "Qtom.I.SB.73"    "Qtom.I.SB.73"   
## [82,] "Qtom.I.SB.74"    "Qtom.I.SB.74"    "Qtom.I.SB.74"    "Qtom.I.SB.74"   
## [83,] "Qtom.I.VEN.01"   "Qtom.I.VEN.01"   "Qtom.I.VEN.01"   "Qtom.I.VEN.01"  
## [84,] "Qtom.I.VEN.02"   "Qtom.I.VEN.02"   "Qtom.I.VEN.02"   "Qtom.I.VEN.02"  
## [85,] "Qtom.I.VEN.03"   "Qtom.I.VEN.03"   "Qtom.I.VEN.03"   "Qtom.I.VEN.03"  
## [86,] "Qtom.I.VEN.04"   "Qtom.I.VEN.04"   "Qtom.I.VEN.04"   "Qtom.I.VEN.04"  
## [87,] "Qtom.I.VEN.05"   "Qtom.I.VEN.05"   "Qtom.I.VEN.05"   "Qtom.I.VEN.05"  
## [88,] "Qtom.JJK.SB.01"  "Qtom.JJK.SB.01"  "Qtom.JJK.SB.01"  "Qtom.JJK.SB.01" 
## [89,] "Qtom.JJK.SB.03"  "Qtom.JJK.SB.03"  "Qtom.JJK.SB.03"  "Qtom.JJK.SB.03" 
## [90,] "Qtom.JJK.SB.07"  "Qtom.JJK.SB.07"  "Qtom.JJK.SB.07"  "Qtom.JJK.SB.07" 
## [91,] "Qtom.JJK.SB.08"  "Qtom.JJK.SB.08"  "Qtom.JJK.SB.08"  "Qtom.JJK.SB.08" 
## [92,] "Qtom.JJK.SB.09"  "Qtom.JJK.SB.09"  "Qtom.JJK.SB.09"  "Qtom.JJK.SB.09" 
## [93,] "Qtom.JJK.SB.10"  "Qtom.JJK.SB.10"  "Qtom.JJK.SB.10"  "Qtom.JJK.SB.10" 
## [94,] "Qtom.JR.SB.01"   "Qtom.JR.SB.01"   "Qtom.JR.SB.01"   "Qtom.JR.SB.01"  
## [95,] "Qtom.S.LA.186"   "Qtom.S.LA.186"   "Qtom.S.LA.186"   "Qtom.S.LA.186"
```

```
rda <- rda(imp.sub ~ ., data = as.data.frame(clim.sub), scale = T)
screeplot(rda)
```

```
plot_clim(rda, info = info.sub, save = F, file_string = 'Qtom107.Qchr17.Qssp3_noQchrGuadalupe')
```

```
plot_clim(rda, info = info.sub, choices = c(1,3), save = F, file_string = 'Qtom107.Qchr17.Qssp3_noQchrGuadalupe')
```

#### 3.4 Basic RDA, visualize with only Guadalupe

```
# same plot with only Guadalupe

# get subset without guadalupe or mainland Qchr
clim.sub <- clim[info$island == 'Guadalupe Island',]
snps.sub <- snps[info$island == 'Guadalupe Island',]
imp.sub <- imp[info$island == 'Guadalupe Island',]
info.sub <- info[info$island == 'Guadalupe Island',]

cbind(rownames(info.sub), rownames(snps.sub), rownames(imp.sub), rownames(clim.sub))
```

```
##       [,1]           [,2]           [,3]           [,4]          
##  [1,] "Qtom.I.BC.10" "Qtom.I.BC.10" "Qtom.I.BC.10" "Qtom.I.BC.10"
##  [2,] "Qtom.I.BC.11" "Qtom.I.BC.11" "Qtom.I.BC.11" "Qtom.I.BC.11"
##  [3,] "Qtom.I.BC.12" "Qtom.I.BC.12" "Qtom.I.BC.12" "Qtom.I.BC.12"
##  [4,] "Qtom.I.BC.13" "Qtom.I.BC.13" "Qtom.I.BC.13" "Qtom.I.BC.13"
##  [5,] "Qtom.I.BC.18" "Qtom.I.BC.18" "Qtom.I.BC.18" "Qtom.I.BC.18"
##  [6,] "Qtom.I.BC.20" "Qtom.I.BC.20" "Qtom.I.BC.20" "Qtom.I.BC.20"
##  [7,] "Qtom.I.BC.21" "Qtom.I.BC.21" "Qtom.I.BC.21" "Qtom.I.BC.21"
##  [8,] "Qtom.I.BC.23" "Qtom.I.BC.23" "Qtom.I.BC.23" "Qtom.I.BC.23"
##  [9,] "Qtom.I.BC.24" "Qtom.I.BC.24" "Qtom.I.BC.24" "Qtom.I.BC.24"
## [10,] "Qtom.I.BC.32" "Qtom.I.BC.32" "Qtom.I.BC.32" "Qtom.I.BC.32"
## [11,] "Qtom.I.BC.33" "Qtom.I.BC.33" "Qtom.I.BC.33" "Qtom.I.BC.33"
## [12,] "Qtom.I.BC.35" "Qtom.I.BC.35" "Qtom.I.BC.35" "Qtom.I.BC.35"
## [13,] "Qtom.I.BC.39" "Qtom.I.BC.39" "Qtom.I.BC.39" "Qtom.I.BC.39"
## [14,] "Qtom.I.BC.4"  "Qtom.I.BC.4"  "Qtom.I.BC.4"  "Qtom.I.BC.4" 
## [15,] "Qtom.I.BC.40" "Qtom.I.BC.40" "Qtom.I.BC.40" "Qtom.I.BC.40"
## [16,] "Qtom.I.BC.41" "Qtom.I.BC.41" "Qtom.I.BC.41" "Qtom.I.BC.41"
## [17,] "Qtom.I.BC.42" "Qtom.I.BC.42" "Qtom.I.BC.42" "Qtom.I.BC.42"
## [18,] "Qtom.I.BC.43" "Qtom.I.BC.43" "Qtom.I.BC.43" "Qtom.I.BC.43"
## [19,] "Qtom.I.BC.44" "Qtom.I.BC.44" "Qtom.I.BC.44" "Qtom.I.BC.44"
## [20,] "Qtom.I.BC.45" "Qtom.I.BC.45" "Qtom.I.BC.45" "Qtom.I.BC.45"
## [21,] "Qtom.I.BC.46" "Qtom.I.BC.46" "Qtom.I.BC.46" "Qtom.I.BC.46"
## [22,] "Qtom.I.BC.48" "Qtom.I.BC.48" "Qtom.I.BC.48" "Qtom.I.BC.48"
```

```
rda <- rda(imp.sub ~ ., data = as.data.frame(clim.sub), scale = T)
screeplot(rda)
```

```
plot_clim(rda, info = info.sub, save = F, file_string = 'Qtom107.Qchr17.Qssp3_onlyGuadalupe')
```

```
plot_clim(rda, info = info.sub, choices = c(1,3), save = F, file_string = 'Qtom107.Qchr17.Qssp3_onlyGuadalupe')
```

#### 3.5 Reduce Climate Variables

```
# reduce strongly correlated climate variables for use in final RDA

# visualize correlations - all individuals
pairs.panels(clim, scale = T)
```

```
heatmap(abs(cor(clim)), scale = 'none')
```

```
# just the california channel islands
# going to focus on this set because it's what I'll be using for landscape genomics
# get subset without guadalupe or mainland Qchr
clim.sub <- clim[! info$island %in% c('Guadalupe Island', "Mainland"),]
snps.sub <- snps[! info$island %in% c('Guadalupe Island', "Mainland"),]
imp.sub <- imp[! info$island %in% c('Guadalupe Island', "Mainland"),]
info.sub <- info[! info$island %in% c('Guadalupe Island', "Mainland"),]

pairs.panels(clim.sub, scale = T)
```

```
heatmap(abs(cor(clim.sub)), scale = 'none')
```

```
# variables chosen by looking at loadings on pca and choosing one important var from each correlated group
# these also seem biologically important and include both temp and precip vars
vars <- c('bio5', 'bio6', 'bio15', 'bio18', 'bio19', 'elev')
heatmap(abs(cor(clim.sub[,vars])), scale = 'none')
```

```
pairs.panels(clim.sub[,vars], scale = T)
```

```
# BIO5 = Max Temperature of Warmest Month
# BIO6 = Min Temperature of Coldest Month
# BIO15 = Precipitation Seasonality (Coefficient of Variation)
# BIO18 = Precipitation of Warmest Quarter
# BIO19 = Precipitation of Coldest Quarter
```

### 4 Variance partitioning - all individuals

```
##############
# this is running on all individuals


# add geographic info via lat/lon

lat <- info$lat
lon <- info$lon

geo <- cbind(lat, lon)

pairs.panels(cbind(lat, lon, clim[,vars]), scale = T)
```

```
rda <- rda(imp ~ ., data = as.data.frame(clim[,vars]) + lat + lon, scale = T)
rda
```

```
## Call: rda(formula = imp ~ bio5 + bio6 + bio15 + bio18 + bio19 + elev,
## data = as.data.frame(clim[, vars]) + lat + lon, scale = T)
## 
##                 Inertia Proportion Rank
## Total         5.853e+05  1.000e+00     
## Constrained   5.171e+04  8.836e-02    6
## Unconstrained 5.336e+05  9.116e-01  119
## Inertia is correlations 
## 
## Eigenvalues for constrained axes:
##  RDA1  RDA2  RDA3  RDA4  RDA5  RDA6 
## 15052 11076  7247  6636  5977  5725 
## 
## Eigenvalues for unconstrained axes:
##  PC1  PC2  PC3  PC4  PC5  PC6  PC7  PC8 
## 7751 7464 7384 7164 6955 6707 6499 6329 
## (Showing 8 of 119 unconstrained eigenvalues)
```

```
screeplot(rda)
```

```
# variance partitioning 
# will not run if scale = T
# Error in qr.fitted(Q, Y) : NA/NaN/Inf in foreign function call (arg 5)
part <- varpart(imp, geo, clim[,vars])


# rdas
# values are from the subset of 10000 snps
# update 11/25: run on all snps

rda.full <- rda(imp ~ clim[,vars] + geo, scale = T)
RsquareAdj(rda.full) # 0.048143
```

```
## $r.squared
## [1] 0.1090618
## 
## $adj.r.squared
## [1] 0.048143
```

```
rda.full.info <- summary(rda.full)

head(rda.full.info)
```

```
## 
## Call:
## rda(formula = imp ~ clim[, vars] + geo, scale = T) 
## 
## Partitioning of correlations:
##               Inertia Proportion
## Total          585267     1.0000
## Constrained     63830     0.1091
## Unconstrained  521437     0.8909
## 
## Eigenvalues, and their contribution to the correlations 
## 
## Importance of components:
##                            RDA1      RDA2      RDA3      RDA4      RDA5
## Eigenvalue            1.520e+04 1.108e+04 7.390e+03 6.715e+03 6.374e+03
## Proportion Explained  2.597e-02 1.893e-02 1.263e-02 1.147e-02 1.089e-02
## Cumulative Proportion 2.597e-02 4.491e-02 5.754e-02 6.901e-02 7.990e-02
##                            RDA6      RDA7      RDA8       PC1       PC2
## Eigenvalue            5.877e+03 5.624e+03 5.565e+03 7.637e+03 7.454e+03
## Proportion Explained  1.004e-02 9.609e-03 9.509e-03 1.305e-02 1.274e-02
## Cumulative Proportion 8.994e-02 9.955e-02 1.091e-01 1.221e-01 1.348e-01
##                             PC3       PC4       PC5       PC6       PC7
## Eigenvalue            7.320e+03 7.055e+03 6.915e+03 6.516e+03 6.406e+03
## Proportion Explained  1.251e-02 1.205e-02 1.182e-02 1.113e-02 1.095e-02
## Cumulative Proportion 1.474e-01 1.594e-01 1.712e-01 1.824e-01 1.933e-01
##                             PC8       PC9      PC10      PC11      PC12
## Eigenvalue            6.231e+03 6.197e+03 6.053e+03 6030.2906 5.981e+03
## Proportion Explained  1.065e-02 1.059e-02 1.034e-02    0.0103 1.022e-02
## Cumulative Proportion 2.039e-01 2.145e-01 2.249e-01    0.2352 2.454e-01
##                            PC13      PC14      PC15      PC16      PC17
## Eigenvalue            5.930e+03 5.906e+03 5.793e+03 5.774e+03 5.720e+03
## Proportion Explained  1.013e-02 1.009e-02 9.899e-03 9.866e-03 9.773e-03
## Cumulative Proportion 2.555e-01 2.656e-01 2.755e-01 2.854e-01 2.952e-01
##                            PC18      PC19      PC20      PC21      PC22
## Eigenvalue            5.692e+03 5.678e+03 5.644e+03 5.627e+03 5.589e+03
## Proportion Explained  9.725e-03 9.702e-03 9.643e-03 9.615e-03 9.549e-03
## Cumulative Proportion 3.049e-01 3.146e-01 3.242e-01 3.338e-01 3.434e-01
##                            PC23      PC24      PC25      PC26      PC27
## Eigenvalue            5.469e+03 5.443e+03 5.378e+03 5.245e+03 5.188e+03
## Proportion Explained  9.345e-03 9.299e-03 9.189e-03 8.963e-03 8.864e-03
## Cumulative Proportion 3.527e-01 3.620e-01 3.712e-01 3.802e-01 3.891e-01
##                            PC28      PC29      PC30      PC31      PC32
## Eigenvalue            5.174e+03 5.137e+03 5.102e+03 5.081e+03 5.008e+03
## Proportion Explained  8.840e-03 8.778e-03 8.717e-03 8.682e-03 8.556e-03
## Cumulative Proportion 3.979e-01 4.067e-01 4.154e-01 4.241e-01 4.326e-01
##                            PC33      PC34      PC35      PC36      PC37
## Eigenvalue            4.981e+03 4.923e+03 4.908e+03 4.861e+03 4.830e+03
## Proportion Explained  8.510e-03 8.412e-03 8.386e-03 8.306e-03 8.253e-03
## Cumulative Proportion 4.411e-01 4.496e-01 4.579e-01 4.662e-01 4.745e-01
##                            PC38      PC39      PC40      PC41      PC42
## Eigenvalue            4799.2431 4.764e+03 4.742e+03 4.694e+03 4.666e+03
## Proportion Explained     0.0082 8.140e-03 8.102e-03 8.020e-03 7.972e-03
## Cumulative Proportion    0.4827 4.908e-01 4.989e-01 5.070e-01 5.149e-01
##                            PC43      PC44      PC45      PC46      PC47
## Eigenvalue            4.647e+03 4.618e+03 4.584e+03 4.574e+03 4.560e+03
## Proportion Explained  7.940e-03 7.891e-03 7.832e-03 7.816e-03 7.791e-03
## Cumulative Proportion 5.229e-01 5.308e-01 5.386e-01 5.464e-01 5.542e-01
##                            PC48      PC49      PC50      PC51      PC52
## Eigenvalue            4.534e+03 4.452e+03 4.424e+03 4.411e+03 4.409e+03
## Proportion Explained  7.746e-03 7.607e-03 7.559e-03 7.538e-03 7.533e-03
## Cumulative Proportion 5.619e-01 5.696e-01 5.771e-01 5.846e-01 5.922e-01
##                            PC53      PC54      PC55      PC56      PC57
## Eigenvalue            4.385e+03 4.363e+03 4.337e+03 4.299e+03 4.294e+03
## Proportion Explained  7.492e-03 7.456e-03 7.411e-03 7.346e-03 7.337e-03
## Cumulative Proportion 5.997e-01 6.071e-01 6.145e-01 6.219e-01 6.292e-01
##                            PC58      PC59      PC60      PC61      PC62
## Eigenvalue            4.265e+03 4.255e+03 4.240e+03 4.218e+03 4.198e+03
## Proportion Explained  7.288e-03 7.271e-03 7.244e-03 7.207e-03 7.173e-03
## Cumulative Proportion 6.365e-01 6.438e-01 6.510e-01 6.582e-01 6.654e-01
##                            PC63      PC64      PC65      PC66      PC67
## Eigenvalue            4.189e+03 4.137e+03 4.102e+03 4.091e+03 4.080e+03
## Proportion Explained  7.157e-03 7.069e-03 7.009e-03 6.990e-03 6.971e-03
## Cumulative Proportion 6.726e-01 6.796e-01 6.866e-01 6.936e-01 7.006e-01
##                            PC68      PC69      PC70      PC71      PC72
## Eigenvalue            4.058e+03 4.026e+03 4.008e+03 3.980e+03 3.971e+03
## Proportion Explained  6.933e-03 6.879e-03 6.849e-03 6.801e-03 6.786e-03
## Cumulative Proportion 7.075e-01 7.144e-01 7.213e-01 7.281e-01 7.349e-01
##                            PC73      PC74      PC75      PC76      PC77
## Eigenvalue            3.931e+03 3.907e+03 3.885e+03 3.867e+03 3.857e+03
## Proportion Explained  6.717e-03 6.676e-03 6.637e-03 6.607e-03 6.590e-03
## Cumulative Proportion 7.416e-01 7.482e-01 7.549e-01 7.615e-01 7.681e-01
##                            PC78      PC79      PC80      PC81      PC82
## Eigenvalue            3.832e+03 3.824e+03 3.815e+03 3.795e+03 3.746e+03
## Proportion Explained  6.547e-03 6.533e-03 6.519e-03 6.484e-03 6.401e-03
## Cumulative Proportion 7.746e-01 7.812e-01 7.877e-01 7.942e-01 8.006e-01
##                            PC83      PC84      PC85      PC86      PC87
## Eigenvalue            3.736e+03 3.728e+03 3.713e+03 3.695e+03 3.692e+03
## Proportion Explained  6.383e-03 6.370e-03 6.343e-03 6.313e-03 6.309e-03
## Cumulative Proportion 8.069e-01 8.133e-01 8.197e-01 8.260e-01 8.323e-01
##                            PC88      PC89      PC90      PC91      PC92
## Eigenvalue            3.677e+03 3.671e+03 3.653e+03 3.627e+03 3.623e+03
## Proportion Explained  6.282e-03 6.272e-03 6.241e-03 6.197e-03 6.190e-03
## Cumulative Proportion 8.386e-01 8.448e-01 8.511e-01 8.573e-01 8.635e-01
##                            PC93      PC94      PC95      PC96      PC97
## Eigenvalue            3.611e+03 3.602e+03 3.586e+03 3.583e+03 3.513e+03
## Proportion Explained  6.169e-03 6.155e-03 6.128e-03 6.122e-03 6.002e-03
## Cumulative Proportion 8.696e-01 8.758e-01 8.819e-01 8.880e-01 8.940e-01
##                            PC98      PC99     PC100     PC101     PC102
## Eigenvalue            3.502e+03 3.476e+03 3.474e+03 3.439e+03 3.424e+03
## Proportion Explained  5.983e-03 5.939e-03 5.936e-03 5.876e-03 5.850e-03
## Cumulative Proportion 9.000e-01 9.060e-01 9.119e-01 9.178e-01 9.236e-01
##                           PC103     PC104     PC105     PC106     PC107
## Eigenvalue            3.405e+03 3.391e+03 3.333e+03 3.279e+03 3.260e+03
## Proportion Explained  5.817e-03 5.794e-03 5.694e-03 5.603e-03 5.571e-03
## Cumulative Proportion 9.294e-01 9.352e-01 9.409e-01 9.465e-01 9.521e-01
##                           PC108     PC109     PC110     PC111     PC112
## Eigenvalue            3.162e+03 3.143e+03 3.041e+03 3.014e+03 2.913e+03
## Proportion Explained  5.402e-03 5.370e-03 5.196e-03 5.149e-03 4.977e-03
## Cumulative Proportion 9.575e-01 9.629e-01 9.681e-01 9.732e-01 9.782e-01
##                           PC113     PC114     PC115     PC116     PC117
## Eigenvalue            2.851e+03 2.825e+03 2.791e+03 2.585e+03 1.710e+03
## Proportion Explained  4.872e-03 4.827e-03 4.768e-03 4.417e-03 2.921e-03
## Cumulative Proportion 9.831e-01 9.879e-01 9.927e-01 9.971e-01 1.000e+00
## 
## Accumulated constrained eigenvalues
## Importance of components:
##                            RDA1      RDA2      RDA3      RDA4      RDA5
## Eigenvalue            1.520e+04 1.108e+04 7390.3847 6715.3253 6.374e+03
## Proportion Explained  2.382e-01 1.736e-01    0.1158    0.1052 9.986e-02
## Cumulative Proportion 2.382e-01 4.118e-01    0.5276    0.6328 7.326e-01
##                            RDA6      RDA7      RDA8
## Eigenvalue            5.877e+03 5.624e+03 5.565e+03
## Proportion Explained  9.207e-02 8.811e-02 8.719e-02
## Cumulative Proportion 8.247e-01 9.128e-01 1.000e+00
## 
## Scaling 2 for species and site scores
## * Species are scaled proportional to eigenvalues
## * Sites are unscaled: weighted dispersion equal on all dimensions
## * General scaling constant of scores:  92.48387 
## 
## 
## Species scores
## 
##                    RDA1       RDA2      RDA3      RDA4       RDA5      RDA6
## 1_39789_A_G_G  0.020605  0.0093619  0.022020 -0.023509 -0.0021037 -0.004453
## 1_52176_G_A_A  0.014463 -0.0169802 -0.013268 -0.011891 -0.0080510  0.001014
## 1_52195_G_A_A -0.003229 -0.0023070  0.005839  0.005274  0.0013350 -0.004580
## 1_52213_C_T_T  0.019187  0.0056283 -0.003113  0.008856  0.0035800 -0.003769
## 1_52248_T_C_T -0.004532 -0.0003605  0.008253 -0.013979 -0.0001901 -0.003396
## 1_52331_G_T_T  0.006097  0.0183735  0.014852  0.021821  0.0091662  0.007266
## ....                                                                       
## 
## 
## Site scores (weighted sums of species scores)
## 
##                 RDA1  RDA2    RDA3   RDA4    RDA5     RDA6
## Qchr.A.232    -2.667 17.33 -30.151  1.902   9.796 -30.1168
## Qchr.A.275    -1.876 21.18 -26.652 -2.851  -2.605 -16.1791
## Qchr.A.ED.102 -5.351 26.40 -13.770 -6.853  10.617   0.4073
## Qchr.A.ELD.53 -3.658 22.96 -19.565 -1.947  16.129  28.0936
## Qchr.A.KER.16 -4.527 18.06  -3.222 13.708   6.671  26.5977
## Qchr.A.LA.02  -3.821 20.29   7.085  7.232 -15.173  36.1759
## ....                                                      
## 
## 
## Site constraints (linear combinations of constraining variables)
## 
##                  RDA1  RDA2    RDA3   RDA4    RDA5    RDA6
## Qchr.A.232    -2.3560 14.17 -29.948  3.129  11.085 -29.080
## Qchr.A.275     0.6451 22.54 -26.928 -2.722  -3.589 -15.621
## Qchr.A.ED.102 -7.0663 28.58 -12.140 -7.284  11.166  -1.757
## Qchr.A.ELD.53 -2.3548 20.23 -17.578 -1.557  16.674  27.833
## Qchr.A.KER.16 -6.3211 19.75  -3.550 17.693   8.243  27.853
## Qchr.A.LA.02  -2.6029 20.08   6.094  9.059 -15.093  34.877
## ....                                                      
## 
## 
## Biplot scores for constraining variables
## 
##                       RDA1     RDA2     RDA3      RDA4     RDA5      RDA6
## clim[, vars]bio5  -0.08874  0.87349 -0.05066  0.146018 -0.34329 -0.129476
## clim[, vars]bio6  -0.18966 -0.60111  0.61952 -0.017849  0.03349 -0.460180
## clim[, vars]bio15  0.91311  0.02151  0.30771  0.205749  0.02271 -0.147568
## clim[, vars]bio18 -0.23447  0.18749 -0.70879 -0.336618 -0.26571  0.348758
## clim[, vars]bio19  0.41151  0.57517 -0.53949  0.067324  0.16027  0.045327
## clim[, vars]elev  -0.54073  0.50027 -0.25392 -0.001976 -0.08979  0.541547
## geolat             0.67894  0.53106 -0.48642 -0.003084  0.13715  0.009252
## geolon            -0.45943 -0.02011  0.85008 -0.095961 -0.03655  0.217451
```

```
rda.full.info$cont$importance
```

```
## Importance of components:
##                            RDA1      RDA2      RDA3      RDA4      RDA5
## Eigenvalue            1.520e+04 1.108e+04 7.390e+03 6.715e+03 6.374e+03
## Proportion Explained  2.597e-02 1.893e-02 1.263e-02 1.147e-02 1.089e-02
## Cumulative Proportion 2.597e-02 4.491e-02 5.754e-02 6.901e-02 7.990e-02
##                            RDA6      RDA7      RDA8       PC1       PC2
## Eigenvalue            5.877e+03 5.624e+03 5.565e+03 7.637e+03 7.454e+03
## Proportion Explained  1.004e-02 9.609e-03 9.509e-03 1.305e-02 1.274e-02
## Cumulative Proportion 8.994e-02 9.955e-02 1.091e-01 1.221e-01 1.348e-01
##                             PC3       PC4       PC5       PC6       PC7
## Eigenvalue            7.320e+03 7.055e+03 6.915e+03 6.516e+03 6.406e+03
## Proportion Explained  1.251e-02 1.205e-02 1.182e-02 1.113e-02 1.095e-02
## Cumulative Proportion 1.474e-01 1.594e-01 1.712e-01 1.824e-01 1.933e-01
##                             PC8       PC9      PC10      PC11      PC12
## Eigenvalue            6.231e+03 6.197e+03 6.053e+03 6030.2906 5.981e+03
## Proportion Explained  1.065e-02 1.059e-02 1.034e-02    0.0103 1.022e-02
## Cumulative Proportion 2.039e-01 2.145e-01 2.249e-01    0.2352 2.454e-01
##                            PC13      PC14      PC15      PC16      PC17
## Eigenvalue            5.930e+03 5.906e+03 5.793e+03 5.774e+03 5.720e+03
## Proportion Explained  1.013e-02 1.009e-02 9.899e-03 9.866e-03 9.773e-03
## Cumulative Proportion 2.555e-01 2.656e-01 2.755e-01 2.854e-01 2.952e-01
##                            PC18      PC19      PC20      PC21      PC22
## Eigenvalue            5.692e+03 5.678e+03 5.644e+03 5.627e+03 5.589e+03
## Proportion Explained  9.725e-03 9.702e-03 9.643e-03 9.615e-03 9.549e-03
## Cumulative Proportion 3.049e-01 3.146e-01 3.242e-01 3.338e-01 3.434e-01
##                            PC23      PC24      PC25      PC26      PC27
## Eigenvalue            5.469e+03 5.443e+03 5.378e+03 5.245e+03 5.188e+03
## Proportion Explained  9.345e-03 9.299e-03 9.189e-03 8.963e-03 8.864e-03
## Cumulative Proportion 3.527e-01 3.620e-01 3.712e-01 3.802e-01 3.891e-01
##                            PC28      PC29      PC30      PC31      PC32
## Eigenvalue            5.174e+03 5.137e+03 5.102e+03 5.081e+03 5.008e+03
## Proportion Explained  8.840e-03 8.778e-03 8.717e-03 8.682e-03 8.556e-03
## Cumulative Proportion 3.979e-01 4.067e-01 4.154e-01 4.241e-01 4.326e-01
##                            PC33      PC34      PC35      PC36      PC37
## Eigenvalue            4.981e+03 4.923e+03 4.908e+03 4.861e+03 4.830e+03
## Proportion Explained  8.510e-03 8.412e-03 8.386e-03 8.306e-03 8.253e-03
## Cumulative Proportion 4.411e-01 4.496e-01 4.579e-01 4.662e-01 4.745e-01
##                            PC38      PC39      PC40      PC41      PC42
## Eigenvalue            4799.2431 4.764e+03 4.742e+03 4.694e+03 4.666e+03
## Proportion Explained     0.0082 8.140e-03 8.102e-03 8.020e-03 7.972e-03
## Cumulative Proportion    0.4827 4.908e-01 4.989e-01 5.070e-01 5.149e-01
##                            PC43      PC44      PC45      PC46      PC47
## Eigenvalue            4.647e+03 4.618e+03 4.584e+03 4.574e+03 4.560e+03
## Proportion Explained  7.940e-03 7.891e-03 7.832e-03 7.816e-03 7.791e-03
## Cumulative Proportion 5.229e-01 5.308e-01 5.386e-01 5.464e-01 5.542e-01
##                            PC48      PC49      PC50      PC51      PC52
## Eigenvalue            4.534e+03 4.452e+03 4.424e+03 4.411e+03 4.409e+03
## Proportion Explained  7.746e-03 7.607e-03 7.559e-03 7.538e-03 7.533e-03
## Cumulative Proportion 5.619e-01 5.696e-01 5.771e-01 5.846e-01 5.922e-01
##                            PC53      PC54      PC55      PC56      PC57
## Eigenvalue            4.385e+03 4.363e+03 4.337e+03 4.299e+03 4.294e+03
## Proportion Explained  7.492e-03 7.456e-03 7.411e-03 7.346e-03 7.337e-03
## Cumulative Proportion 5.997e-01 6.071e-01 6.145e-01 6.219e-01 6.292e-01
##                            PC58      PC59      PC60      PC61      PC62
## Eigenvalue            4.265e+03 4.255e+03 4.240e+03 4.218e+03 4.198e+03
## Proportion Explained  7.288e-03 7.271e-03 7.244e-03 7.207e-03 7.173e-03
## Cumulative Proportion 6.365e-01 6.438e-01 6.510e-01 6.582e-01 6.654e-01
##                            PC63      PC64      PC65      PC66      PC67
## Eigenvalue            4.189e+03 4.137e+03 4.102e+03 4.091e+03 4.080e+03
## Proportion Explained  7.157e-03 7.069e-03 7.009e-03 6.990e-03 6.971e-03
## Cumulative Proportion 6.726e-01 6.796e-01 6.866e-01 6.936e-01 7.006e-01
##                            PC68      PC69      PC70      PC71      PC72
## Eigenvalue            4.058e+03 4.026e+03 4.008e+03 3.980e+03 3.971e+03
## Proportion Explained  6.933e-03 6.879e-03 6.849e-03 6.801e-03 6.786e-03
## Cumulative Proportion 7.075e-01 7.144e-01 7.213e-01 7.281e-01 7.349e-01
##                            PC73      PC74      PC75      PC76      PC77
## Eigenvalue            3.931e+03 3.907e+03 3.885e+03 3.867e+03 3.857e+03
## Proportion Explained  6.717e-03 6.676e-03 6.637e-03 6.607e-03 6.590e-03
## Cumulative Proportion 7.416e-01 7.482e-01 7.549e-01 7.615e-01 7.681e-01
##                            PC78      PC79      PC80      PC81      PC82
## Eigenvalue            3.832e+03 3.824e+03 3.815e+03 3.795e+03 3.746e+03
## Proportion Explained  6.547e-03 6.533e-03 6.519e-03 6.484e-03 6.401e-03
## Cumulative Proportion 7.746e-01 7.812e-01 7.877e-01 7.942e-01 8.006e-01
##                            PC83      PC84      PC85      PC86      PC87
## Eigenvalue            3.736e+03 3.728e+03 3.713e+03 3.695e+03 3.692e+03
## Proportion Explained  6.383e-03 6.370e-03 6.343e-03 6.313e-03 6.309e-03
## Cumulative Proportion 8.069e-01 8.133e-01 8.197e-01 8.260e-01 8.323e-01
##                            PC88      PC89      PC90      PC91      PC92
## Eigenvalue            3.677e+03 3.671e+03 3.653e+03 3.627e+03 3.623e+03
## Proportion Explained  6.282e-03 6.272e-03 6.241e-03 6.197e-03 6.190e-03
## Cumulative Proportion 8.386e-01 8.448e-01 8.511e-01 8.573e-01 8.635e-01
##                            PC93      PC94      PC95      PC96      PC97
## Eigenvalue            3.611e+03 3.602e+03 3.586e+03 3.583e+03 3.513e+03
## Proportion Explained  6.169e-03 6.155e-03 6.128e-03 6.122e-03 6.002e-03
## Cumulative Proportion 8.696e-01 8.758e-01 8.819e-01 8.880e-01 8.940e-01
##                            PC98      PC99     PC100     PC101     PC102
## Eigenvalue            3.502e+03 3.476e+03 3.474e+03 3.439e+03 3.424e+03
## Proportion Explained  5.983e-03 5.939e-03 5.936e-03 5.876e-03 5.850e-03
## Cumulative Proportion 9.000e-01 9.060e-01 9.119e-01 9.178e-01 9.236e-01
##                           PC103     PC104     PC105     PC106     PC107
## Eigenvalue            3.405e+03 3.391e+03 3.333e+03 3.279e+03 3.260e+03
## Proportion Explained  5.817e-03 5.794e-03 5.694e-03 5.603e-03 5.571e-03
## Cumulative Proportion 9.294e-01 9.352e-01 9.409e-01 9.465e-01 9.521e-01
##                           PC108     PC109     PC110     PC111     PC112
## Eigenvalue            3.162e+03 3.143e+03 3.041e+03 3.014e+03 2.913e+03
## Proportion Explained  5.402e-03 5.370e-03 5.196e-03 5.149e-03 4.977e-03
## Cumulative Proportion 9.575e-01 9.629e-01 9.681e-01 9.732e-01 9.782e-01
##                           PC113     PC114     PC115     PC116     PC117
## Eigenvalue            2.851e+03 2.825e+03 2.791e+03 2.585e+03 1.710e+03
## Proportion Explained  4.872e-03 4.827e-03 4.768e-03 4.417e-03 2.921e-03
## Cumulative Proportion 9.831e-01 9.879e-01 9.927e-01 9.971e-01 1.000e+00
```

```
rda.full.info$concont$importance
```

```
## Importance of components:
##                            RDA1      RDA2      RDA3      RDA4      RDA5
## Eigenvalue            1.520e+04 1.108e+04 7390.3847 6715.3253 6.374e+03
## Proportion Explained  2.382e-01 1.736e-01    0.1158    0.1052 9.986e-02
## Cumulative Proportion 2.382e-01 4.118e-01    0.5276    0.6328 7.326e-01
##                            RDA6      RDA7      RDA8
## Eigenvalue            5.877e+03 5.624e+03 5.565e+03
## Proportion Explained  9.207e-02 8.811e-02 8.719e-02
## Cumulative Proportion 8.247e-01 9.128e-01 1.000e+00
```

```
# Partitioning of correlations:
#               Inertia Proportion
# Total          585267     1.0000
# Constrained     63830     0.1091
# Unconstrained  521437     0.8909

rda.clim_not_geo <- rda(imp ~ clim[,vars] + Condition(geo), scale = T)
RsquareAdj(rda.clim_not_geo) # 0.02905444
```

```
## $r.squared
## [1] 0.0742787
## 
## $adj.r.squared
## [1] 0.02905444
```

```
rda.geo_not_clim <- rda(imp ~ geo + Condition(clim[,vars]), scale = T)
RsquareAdj(rda.geo_not_clim) # 0.007226599
```

```
## $r.squared
## [1] 0.02210943
## 
## $adj.r.squared
## [1] 0.007226599
```

```
# sig testing
sig.full <- anova.cca(rda.full, permutations=99)
sig.full
```

```
## Permutation test for rda under reduced model
## Permutation: free
## Number of permutations: 99
## 
## Model: rda(formula = imp ~ clim[, vars] + geo, scale = T)
##           Df Variance      F Pr(>F)   
## Model      8    63830 1.7903   0.01 **
## Residual 117   521437                 
## ---
## Signif. codes:  0 '***' 0.001 '**' 0.01 '*' 0.05 '.' 0.1 ' ' 1
```

```
# Permutation test for rda under reduced model
# Permutation: free
# Number of permutations: 99
# 
# Model: rda(formula = imp ~ clim[, vars] + geo, scale = T)
#           Df Variance      F Pr(>F)   
# Model      8    63830 1.7903   0.01 **
# Residual 117   521437                 
# ---
# Signif. codes:  0 ‘***’ 0.001 ‘**’ 0.01 ‘*’ 0.05 ‘.’ 0.1 ‘ ’ 1

sig.clim_not_geo <- anova.cca(rda.clim_not_geo, permutations=99)
sig.clim_not_geo
```

```
## Permutation test for rda under reduced model
## Permutation: free
## Number of permutations: 99
## 
## Model: rda(formula = imp ~ clim[, vars] + Condition(geo), scale = T)
##           Df Variance      F Pr(>F)   
## Model      6    43473 1.6257   0.01 **
## Residual 117   521437                 
## ---
## Signif. codes:  0 '***' 0.001 '**' 0.01 '*' 0.05 '.' 0.1 ' ' 1
```

```
# Permutation test for rda under reduced model
# Permutation: free
# Number of permutations: 99
# 
# Model: rda(formula = imp ~ clim[, vars] + Condition(geo), scale = T)
#           Df Variance      F Pr(>F)   
# Model      6    43473 1.6257   0.01 **
# Residual 117   521437                 
# ---
# Signif. codes:  0 ‘***’ 0.001 ‘**’ 0.01 ‘*’ 0.05 ‘.’ 0.1 ‘ ’ 1


sig.geo_not_clim <- anova.cca(rda.geo_not_clim, permutations=99)
sig.geo_not_clim
```

```
## Permutation test for rda under reduced model
## Permutation: free
## Number of permutations: 99
## 
## Model: rda(formula = imp ~ geo + Condition(clim[, vars]), scale = T)
##           Df Variance      F Pr(>F)   
## Model      2    12940 1.4517   0.01 **
## Residual 117   521437                 
## ---
## Signif. codes:  0 '***' 0.001 '**' 0.01 '*' 0.05 '.' 0.1 ' ' 1
```

```
# Permutation test for rda under reduced model
# Permutation: free
# Number of permutations: 99
# 
# Model: rda(formula = imp ~ geo + Condition(clim[, vars]), scale = T)
#           Df Variance      F Pr(>F)   
# Model      2    12940 1.4517   0.01 **
# Residual 117   521437                 
# ---
# Signif. codes:  0 ‘***’ 0.001 ‘**’ 0.01 ‘*’ 0.05 ‘.’ 0.1 ‘ ’ 1
```

### 5 GEA analysis, all Q. tomentella

#### 5.1 RDA of SNPs and climate

```
# here we'll use the subset climate variables to do a more detailed GEA

# with all samples except Mainland Qchr
# Qchr on islands seems to be highly introgressed with Qtom, so including all island individuals together

# get subset without mainland
clim.sub <- clim[! info$island %in% c('Mainland'),]
snps.sub <- snps[! info$island %in% c('Mainland'),]
imp.sub <- imp[! info$island %in% c('Mainland'),]
info.sub <- info[! info$island %in% c('Mainland'),]

rda <- rda(imp.sub ~ ., data = as.data.frame(clim.sub[,vars]), scale = T)
screeplot(rda)
```

```
rda
```

```
## Call: rda(formula = imp.sub ~ bio5 + bio6 + bio15 + bio18 + bio19 +
## elev, data = as.data.frame(clim.sub[, vars]), scale = T)
## 
##                 Inertia Proportion Rank
## Total         5.841e+05  1.000e+00     
## Constrained   5.231e+04  8.955e-02    6
## Unconstrained 5.318e+05  9.104e-01  110
## Inertia is correlations 
## 
## Eigenvalues for constrained axes:
##  RDA1  RDA2  RDA3  RDA4  RDA5  RDA6 
## 16268  9586  7382  7081  6228  5760 
## 
## Eigenvalues for unconstrained axes:
##  PC1  PC2  PC3  PC4  PC5  PC6  PC7  PC8 
## 8815 8250 7963 7804 7356 7205 6937 6861 
## (Showing 8 of 110 unconstrained eigenvalues)
```

```
rda.info <- summary(rda)
head(rda.info)
```

```
## 
## Call:
## rda(formula = imp.sub ~ bio5 + bio6 + bio15 + bio18 + bio19 +      elev, data = as.data.frame(clim.sub[, vars]), scale = T) 
## 
## Partitioning of correlations:
##               Inertia Proportion
## Total          584105    1.00000
## Constrained     52305    0.08955
## Unconstrained  531800    0.91045
## 
## Eigenvalues, and their contribution to the correlations 
## 
## Importance of components:
##                            RDA1      RDA2      RDA3      RDA4      RDA5
## Eigenvalue            1.627e+04 9.586e+03 7.382e+03 7.081e+03 6.228e+03
## Proportion Explained  2.785e-02 1.641e-02 1.264e-02 1.212e-02 1.066e-02
## Cumulative Proportion 2.785e-02 4.426e-02 5.690e-02 6.902e-02 7.969e-02
##                            RDA6       PC1       PC2       PC3       PC4
## Eigenvalue            5.760e+03 8.815e+03 8.250e+03 7.963e+03 7.804e+03
## Proportion Explained  9.861e-03 1.509e-02 1.412e-02 1.363e-02 1.336e-02
## Cumulative Proportion 8.955e-02 1.046e-01 1.188e-01 1.324e-01 1.458e-01
##                             PC5       PC6       PC7       PC8       PC9
## Eigenvalue            7.356e+03 7.205e+03 6.937e+03 6.861e+03 6.746e+03
## Proportion Explained  1.259e-02 1.234e-02 1.188e-02 1.175e-02 1.155e-02
## Cumulative Proportion 1.583e-01 1.707e-01 1.826e-01 1.943e-01 2.059e-01
##                            PC10      PC11      PC12      PC13      PC14
## Eigenvalue            6.728e+03 6.683e+03 6.519e+03 6.487e+03 6.473e+03
## Proportion Explained  1.152e-02 1.144e-02 1.116e-02 1.111e-02 1.108e-02
## Cumulative Proportion 2.174e-01 2.288e-01 2.400e-01 2.511e-01 2.622e-01
##                            PC15      PC16      PC17      PC18      PC19
## Eigenvalue            6.325e+03 6.238e+03 6134.6625 6.116e+03 6.009e+03
## Proportion Explained  1.083e-02 1.068e-02    0.0105 1.047e-02 1.029e-02
## Cumulative Proportion 2.730e-01 2.837e-01    0.2942 3.046e-01 3.149e-01
##                            PC20      PC21      PC22      PC23      PC24
## Eigenvalue            5841.9439 5.833e+03 5.721e+03 5.681e+03 5.629e+03
## Proportion Explained     0.0100 9.987e-03 9.794e-03 9.725e-03 9.637e-03
## Cumulative Proportion    0.3249 3.349e-01 3.447e-01 3.544e-01 3.641e-01
##                            PC25      PC26      PC27      PC28      PC29
## Eigenvalue            5.588e+03 5.556e+03 5.531e+03 5.513e+03 5.436e+03
## Proportion Explained  9.567e-03 9.513e-03 9.470e-03 9.438e-03 9.307e-03
## Cumulative Proportion 3.736e-01 3.832e-01 3.926e-01 4.021e-01 4.114e-01
##                            PC30      PC31      PC32      PC33      PC34
## Eigenvalue            5.375e+03 5.352e+03 5.310e+03 5.269e+03 5.263e+03
## Proportion Explained  9.202e-03 9.163e-03 9.091e-03 9.021e-03 9.011e-03
## Cumulative Proportion 4.206e-01 4.297e-01 4.388e-01 4.479e-01 4.569e-01
##                            PC35      PC36      PC37      PC38      PC39
## Eigenvalue            5.238e+03 5.224e+03 5.173e+03 5.114e+03 5.065e+03
## Proportion Explained  8.967e-03 8.944e-03 8.855e-03 8.756e-03 8.671e-03
## Cumulative Proportion 4.658e-01 4.748e-01 4.836e-01 4.924e-01 5.011e-01
##                            PC40      PC41      PC42      PC43      PC44
## Eigenvalue            5.044e+03 5.007e+03 4.971e+03 4.910e+03 4.894e+03
## Proportion Explained  8.635e-03 8.571e-03 8.510e-03 8.406e-03 8.379e-03
## Cumulative Proportion 5.097e-01 5.183e-01 5.268e-01 5.352e-01 5.436e-01
##                            PC45      PC46      PC47      PC48      PC49
## Eigenvalue            4.870e+03 4.833e+03 4.830e+03 4.794e+03 4.748e+03
## Proportion Explained  8.338e-03 8.275e-03 8.269e-03 8.208e-03 8.129e-03
## Cumulative Proportion 5.519e-01 5.602e-01 5.684e-01 5.766e-01 5.848e-01
##                            PC50      PC51      PC52      PC53      PC54
## Eigenvalue            4.738e+03 4.722e+03 4.699e+03 4.684e+03 4.655e+03
## Proportion Explained  8.112e-03 8.084e-03 8.045e-03 8.019e-03 7.970e-03
## Cumulative Proportion 5.929e-01 6.010e-01 6.090e-01 6.170e-01 6.250e-01
##                            PC55      PC56      PC57      PC58      PC59
## Eigenvalue            4.637e+03 4.600e+03 4.562e+03 4.515e+03 4.500e+03
## Proportion Explained  7.939e-03 7.875e-03 7.810e-03 7.730e-03 7.703e-03
## Cumulative Proportion 6.329e-01 6.408e-01 6.486e-01 6.564e-01 6.641e-01
##                            PC60      PC61      PC62      PC63      PC64
## Eigenvalue            4.453e+03 4.430e+03 4.405e+03 4.383e+03 4.357e+03
## Proportion Explained  7.623e-03 7.583e-03 7.542e-03 7.504e-03 7.459e-03
## Cumulative Proportion 6.717e-01 6.793e-01 6.868e-01 6.943e-01 7.018e-01
##                            PC65      PC66      PC67      PC68      PC69
## Eigenvalue            4.339e+03 4.304e+03 4.277e+03 4.261e+03 4.241e+03
## Proportion Explained  7.428e-03 7.369e-03 7.323e-03 7.294e-03 7.261e-03
## Cumulative Proportion 7.092e-01 7.166e-01 7.239e-01 7.312e-01 7.385e-01
##                            PC70      PC71      PC72      PC73      PC74
## Eigenvalue            4.223e+03 4.194e+03 4.180e+03 4.164e+03 4.151e+03
## Proportion Explained  7.231e-03 7.180e-03 7.156e-03 7.129e-03 7.106e-03
## Cumulative Proportion 7.457e-01 7.529e-01 7.600e-01 7.671e-01 7.743e-01
##                            PC75      PC76      PC77      PC78      PC79
## Eigenvalue            4.124e+03 4.104e+03 4089.0206 4.065e+03 4.048e+03
## Proportion Explained  7.060e-03 7.026e-03    0.0070 6.959e-03 6.930e-03
## Cumulative Proportion 7.813e-01 7.883e-01    0.7953 8.023e-01 8.092e-01
##                            PC80      PC81      PC82      PC83      PC84
## Eigenvalue            4.044e+03 4.036e+03 4.025e+03 4.018e+03 4.007e+03
## Proportion Explained  6.923e-03 6.909e-03 6.891e-03 6.880e-03 6.860e-03
## Cumulative Proportion 8.162e-01 8.231e-01 8.300e-01 8.368e-01 8.437e-01
##                            PC85      PC86      PC87      PC88      PC89
## Eigenvalue            3.971e+03 3.952e+03 3.943e+03 3.923e+03 3.905e+03
## Proportion Explained  6.799e-03 6.766e-03 6.750e-03 6.716e-03 6.685e-03
## Cumulative Proportion 8.505e-01 8.573e-01 8.640e-01 8.707e-01 8.774e-01
##                            PC90      PC91      PC92      PC93      PC94
## Eigenvalue            3.851e+03 3.829e+03 3.796e+03 3.780e+03 3.748e+03
## Proportion Explained  6.592e-03 6.555e-03 6.499e-03 6.471e-03 6.416e-03
## Cumulative Proportion 8.840e-01 8.906e-01 8.971e-01 9.035e-01 9.099e-01
##                            PC95      PC96      PC97      PC98      PC99
## Eigenvalue            3.731e+03 3.710e+03 3.697e+03 3.638e+03 3.599e+03
## Proportion Explained  6.388e-03 6.351e-03 6.329e-03 6.229e-03 6.161e-03
## Cumulative Proportion 9.163e-01 9.227e-01 9.290e-01 9.352e-01 9.414e-01
##                           PC100     PC101     PC102     PC103     PC104
## Eigenvalue            3.536e+03 3.470e+03 3.405e+03 3.298e+03 3.285e+03
## Proportion Explained  6.054e-03 5.941e-03 5.829e-03 5.646e-03 5.625e-03
## Cumulative Proportion 9.475e-01 9.534e-01 9.592e-01 9.649e-01 9.705e-01
##                           PC105     PC106     PC107     PC108     PC109
## Eigenvalue            3.192e+03 3.181e+03 3.085e+03 3.040e+03 2.807e+03
## Proportion Explained  5.465e-03 5.446e-03 5.281e-03 5.204e-03 4.805e-03
## Cumulative Proportion 9.760e-01 9.814e-01 9.867e-01 9.919e-01 9.967e-01
##                           PC110
## Eigenvalue            1.931e+03
## Proportion Explained  3.306e-03
## Cumulative Proportion 1.000e+00
## 
## Accumulated constrained eigenvalues
## Importance of components:
##                            RDA1      RDA2      RDA3      RDA4      RDA5
## Eigenvalue            16267.954 9585.8405 7382.4900 7081.3044 6227.8539
## Proportion Explained      0.311    0.1833    0.1411    0.1354    0.1191
## Cumulative Proportion     0.311    0.4943    0.6354    0.7708    0.8899
##                            RDA6
## Eigenvalue            5760.0272
## Proportion Explained     0.1101
## Cumulative Proportion    1.0000
## 
## Scaling 2 for species and site scores
## * Species are scaled proportional to eigenvalues
## * Sites are unscaled: weighted dispersion equal on all dimensions
## * General scaling constant of scores:  90.72714 
## 
## 
## Species scores
## 
##                    RDA1       RDA2      RDA3       RDA4      RDA5      RDA6
## 1_39789_A_G_G -0.020231 -0.0079423  0.052272  1.997e-02  0.011271  0.006389
## 1_52176_G_A_A -0.012585  0.0242605  0.011947 -4.726e-03  0.008348 -0.016272
## 1_52195_G_A_A  0.004031 -0.0056075 -0.004503 -1.705e-05 -0.006460  0.004770
## 1_52213_C_T_T -0.020087 -0.0084879 -0.001408 -3.007e-03 -0.004246 -0.011262
## 1_52248_T_C_T  0.007030  0.0001981  0.006122 -5.751e-05  0.009474 -0.017919
## 1_52331_G_T_T -0.008508 -0.0226805 -0.012937  6.192e-03  0.003013 -0.013165
## ....                                                                       
## 
## 
## Site scores (weighted sums of species scores)
## 
##                  RDA1    RDA2    RDA3    RDA4    RDA5   RDA6
## Qchr.A.LA.215 -1.8261  -7.655   5.159   5.056   2.797  1.329
## Qchr.A.LA.222 -0.3732 -12.620   6.057   4.342   2.630  2.680
## Qchr.I.LA.20  -0.9096  -8.209   2.719  -1.172 -17.996  5.879
## Qchr.I.LA.25  -0.8537  -8.540   4.469   2.383  -8.879 14.935
## Qchr.I.SB.02  -1.7388  -9.867 -26.365 -18.356 -14.709  8.541
## Qchr.I.SB.03  -1.6613 -10.255 -27.517 -18.542 -15.009  8.419
## ....                                                        
## 
## 
## Site constraints (linear combinations of constraining variables)
## 
##                  RDA1   RDA2    RDA3       RDA4    RDA5   RDA6
## Qchr.A.LA.215 -1.3457 -9.167   5.083   4.863238   2.170  1.794
## Qchr.A.LA.222 -1.3457 -9.167   5.083   4.863238   2.170  1.794
## Qchr.I.LA.20  -1.1453 -8.025   2.982   0.008627 -17.421  5.134
## Qchr.I.LA.25  -0.2943 -8.960   5.019   3.604017  -8.014 14.884
## Qchr.I.SB.02  -3.3250 -6.304 -24.910 -15.482268 -11.731  5.184
## Qchr.I.SB.03  -3.3250 -6.304 -24.910 -15.482268 -11.731  5.184
## ....                                                          
## 
## 
## Biplot scores for constraining variables
## 
##            RDA1     RDA2      RDA3    RDA4     RDA5     RDA6
## bio5   0.001246 -0.86450  0.145855 -0.4049  0.17533 -0.19151
## bio6   0.752484 -0.26549  0.353530  0.3907 -0.26633 -0.12122
## bio15 -0.927965 -0.30239 -0.005712  0.1424 -0.09054 -0.13763
## bio18  0.230345  0.72183  0.162557 -0.4943  0.30643  0.24756
## bio19 -0.917447 -0.02684 -0.227504  0.1380  0.28511  0.07403
## elev   0.813235 -0.03396 -0.229804 -0.2890  0.19177  0.40547
```

```
RsquareAdj(rda)
```

```
## $r.squared
## [1] 0.08954806
## 
## $adj.r.squared
## [1] 0.03988705
```

```
# $r.squared
# [1] 0.08954806
# $adj.r.squared
# [1] 0.03988705

# colors
cols <- c('black', "#3c4a8b","#009c85","#84bc5f","#edb829","#f57404","#b30000")

plot_clim(rda, info = info.sub, save = F, file_string = 'Qtom107.Qchr17.Qssp3_GEA_noQchr')
```

```
plot_clim(rda, info = info.sub, choices = c(1,3), save = F, file_string = 'Qtom107.Qchr17.Qssp3_GEA_noQchr')
```

```
plot(rda, type = 'none')
points(rda, display = 'sites', pch = sp[info.sub$sp], bg = bg[info.sub$island], col = 'black', cex = 1.5)
text(rda, display = 'bp')
points(rda, display = 'species', pch = 21, cex = 1, col = 'gray32')
```

#### 5.2 identify candidate SNPs from RDA outliers

```
# again basically following Forester et al

# most variation explained by first three axes (~5.7% out of 9% unadjusted)
screeplot(rda)
```

```
summary(rda)$cont
```

```
## $importance
## Importance of components:
##                            RDA1      RDA2      RDA3      RDA4      RDA5
## Eigenvalue            1.627e+04 9.586e+03 7.382e+03 7.081e+03 6.228e+03
## Proportion Explained  2.785e-02 1.641e-02 1.264e-02 1.212e-02 1.066e-02
## Cumulative Proportion 2.785e-02 4.426e-02 5.690e-02 6.902e-02 7.969e-02
##                            RDA6       PC1       PC2       PC3       PC4
## Eigenvalue            5.760e+03 8.815e+03 8.250e+03 7.963e+03 7.804e+03
## Proportion Explained  9.861e-03 1.509e-02 1.412e-02 1.363e-02 1.336e-02
## Cumulative Proportion 8.955e-02 1.046e-01 1.188e-01 1.324e-01 1.458e-01
##                             PC5       PC6       PC7       PC8       PC9
## Eigenvalue            7.356e+03 7.205e+03 6.937e+03 6.861e+03 6.746e+03
## Proportion Explained  1.259e-02 1.234e-02 1.188e-02 1.175e-02 1.155e-02
## Cumulative Proportion 1.583e-01 1.707e-01 1.826e-01 1.943e-01 2.059e-01
##                            PC10      PC11      PC12      PC13      PC14
## Eigenvalue            6.728e+03 6.683e+03 6.519e+03 6.487e+03 6.473e+03
## Proportion Explained  1.152e-02 1.144e-02 1.116e-02 1.111e-02 1.108e-02
## Cumulative Proportion 2.174e-01 2.288e-01 2.400e-01 2.511e-01 2.622e-01
##                            PC15      PC16      PC17      PC18      PC19
## Eigenvalue            6.325e+03 6.238e+03 6134.6625 6.116e+03 6.009e+03
## Proportion Explained  1.083e-02 1.068e-02    0.0105 1.047e-02 1.029e-02
## Cumulative Proportion 2.730e-01 2.837e-01    0.2942 3.046e-01 3.149e-01
##                            PC20      PC21      PC22      PC23      PC24
## Eigenvalue            5841.9439 5.833e+03 5.721e+03 5.681e+03 5.629e+03
## Proportion Explained     0.0100 9.987e-03 9.794e-03 9.725e-03 9.637e-03
## Cumulative Proportion    0.3249 3.349e-01 3.447e-01 3.544e-01 3.641e-01
##                            PC25      PC26      PC27      PC28      PC29
## Eigenvalue            5.588e+03 5.556e+03 5.531e+03 5.513e+03 5.436e+03
## Proportion Explained  9.567e-03 9.513e-03 9.470e-03 9.438e-03 9.307e-03
## Cumulative Proportion 3.736e-01 3.832e-01 3.926e-01 4.021e-01 4.114e-01
##                            PC30      PC31      PC32      PC33      PC34
## Eigenvalue            5.375e+03 5.352e+03 5.310e+03 5.269e+03 5.263e+03
## Proportion Explained  9.202e-03 9.163e-03 9.091e-03 9.021e-03 9.011e-03
## Cumulative Proportion 4.206e-01 4.297e-01 4.388e-01 4.479e-01 4.569e-01
##                            PC35      PC36      PC37      PC38      PC39
## Eigenvalue            5.238e+03 5.224e+03 5.173e+03 5.114e+03 5.065e+03
## Proportion Explained  8.967e-03 8.944e-03 8.855e-03 8.756e-03 8.671e-03
## Cumulative Proportion 4.658e-01 4.748e-01 4.836e-01 4.924e-01 5.011e-01
##                            PC40      PC41      PC42      PC43      PC44
## Eigenvalue            5.044e+03 5.007e+03 4.971e+03 4.910e+03 4.894e+03
## Proportion Explained  8.635e-03 8.571e-03 8.510e-03 8.406e-03 8.379e-03
## Cumulative Proportion 5.097e-01 5.183e-01 5.268e-01 5.352e-01 5.436e-01
##                            PC45      PC46      PC47      PC48      PC49
## Eigenvalue            4.870e+03 4.833e+03 4.830e+03 4.794e+03 4.748e+03
## Proportion Explained  8.338e-03 8.275e-03 8.269e-03 8.208e-03 8.129e-03
## Cumulative Proportion 5.519e-01 5.602e-01 5.684e-01 5.766e-01 5.848e-01
##                            PC50      PC51      PC52      PC53      PC54
## Eigenvalue            4.738e+03 4.722e+03 4.699e+03 4.684e+03 4.655e+03
## Proportion Explained  8.112e-03 8.084e-03 8.045e-03 8.019e-03 7.970e-03
## Cumulative Proportion 5.929e-01 6.010e-01 6.090e-01 6.170e-01 6.250e-01
##                            PC55      PC56      PC57      PC58      PC59
## Eigenvalue            4.637e+03 4.600e+03 4.562e+03 4.515e+03 4.500e+03
## Proportion Explained  7.939e-03 7.875e-03 7.810e-03 7.730e-03 7.703e-03
## Cumulative Proportion 6.329e-01 6.408e-01 6.486e-01 6.564e-01 6.641e-01
##                            PC60      PC61      PC62      PC63      PC64
## Eigenvalue            4.453e+03 4.430e+03 4.405e+03 4.383e+03 4.357e+03
## Proportion Explained  7.623e-03 7.583e-03 7.542e-03 7.504e-03 7.459e-03
## Cumulative Proportion 6.717e-01 6.793e-01 6.868e-01 6.943e-01 7.018e-01
##                            PC65      PC66      PC67      PC68      PC69
## Eigenvalue            4.339e+03 4.304e+03 4.277e+03 4.261e+03 4.241e+03
## Proportion Explained  7.428e-03 7.369e-03 7.323e-03 7.294e-03 7.261e-03
## Cumulative Proportion 7.092e-01 7.166e-01 7.239e-01 7.312e-01 7.385e-01
##                            PC70      PC71      PC72      PC73      PC74
## Eigenvalue            4.223e+03 4.194e+03 4.180e+03 4.164e+03 4.151e+03
## Proportion Explained  7.231e-03 7.180e-03 7.156e-03 7.129e-03 7.106e-03
## Cumulative Proportion 7.457e-01 7.529e-01 7.600e-01 7.671e-01 7.743e-01
##                            PC75      PC76      PC77      PC78      PC79
## Eigenvalue            4.124e+03 4.104e+03 4089.0206 4.065e+03 4.048e+03
## Proportion Explained  7.060e-03 7.026e-03    0.0070 6.959e-03 6.930e-03
## Cumulative Proportion 7.813e-01 7.883e-01    0.7953 8.023e-01 8.092e-01
##                            PC80      PC81      PC82      PC83      PC84
## Eigenvalue            4.044e+03 4.036e+03 4.025e+03 4.018e+03 4.007e+03
## Proportion Explained  6.923e-03 6.909e-03 6.891e-03 6.880e-03 6.860e-03
## Cumulative Proportion 8.162e-01 8.231e-01 8.300e-01 8.368e-01 8.437e-01
##                            PC85      PC86      PC87      PC88      PC89
## Eigenvalue            3.971e+03 3.952e+03 3.943e+03 3.923e+03 3.905e+03
## Proportion Explained  6.799e-03 6.766e-03 6.750e-03 6.716e-03 6.685e-03
## Cumulative Proportion 8.505e-01 8.573e-01 8.640e-01 8.707e-01 8.774e-01
##                            PC90      PC91      PC92      PC93      PC94
## Eigenvalue            3.851e+03 3.829e+03 3.796e+03 3.780e+03 3.748e+03
## Proportion Explained  6.592e-03 6.555e-03 6.499e-03 6.471e-03 6.416e-03
## Cumulative Proportion 8.840e-01 8.906e-01 8.971e-01 9.035e-01 9.099e-01
##                            PC95      PC96      PC97      PC98      PC99
## Eigenvalue            3.731e+03 3.710e+03 3.697e+03 3.638e+03 3.599e+03
## Proportion Explained  6.388e-03 6.351e-03 6.329e-03 6.229e-03 6.161e-03
## Cumulative Proportion 9.163e-01 9.227e-01 9.290e-01 9.352e-01 9.414e-01
##                           PC100     PC101     PC102     PC103     PC104
## Eigenvalue            3.536e+03 3.470e+03 3.405e+03 3.298e+03 3.285e+03
## Proportion Explained  6.054e-03 5.941e-03 5.829e-03 5.646e-03 5.625e-03
## Cumulative Proportion 9.475e-01 9.534e-01 9.592e-01 9.649e-01 9.705e-01
##                           PC105     PC106     PC107     PC108     PC109
## Eigenvalue            3.192e+03 3.181e+03 3.085e+03 3.040e+03 2.807e+03
## Proportion Explained  5.465e-03 5.446e-03 5.281e-03 5.204e-03 4.805e-03
## Cumulative Proportion 9.760e-01 9.814e-01 9.867e-01 9.919e-01 9.967e-01
##                           PC110
## Eigenvalue            1.931e+03
## Proportion Explained  3.306e-03
## Cumulative Proportion 1.000e+00
```

```
# get SNP loadings for first 3 axes
load.rda <- scores(rda, choices=c(1:3), display="species")

hist(load.rda[,1], main="Loadings on RDA1")
```

```
hist(load.rda[,2], main="Loadings on RDA2")
```

```
hist(load.rda[,3], main="Loadings on RDA3")
```

```
outliers <- function(x,z){
  
  lims <- mean(x) + c(-1, 1) * z * sd(x) # find loadings +/-z sd from mean loading  
  x[x < lims[1] | x > lims[2]] # locus names in these tails
  
}

# start out with a cutoff of 4 std dev because there are a lot of snps
cand1 <- outliers(load.rda[,1],4)
cand2 <- outliers(load.rda[,2],4)
cand3 <- outliers(load.rda[,3],4)

length(cand1)
```

```
## [1] 715
```

```
length(cand2)
```

```
## [1] 77
```

```
length(cand3)
```

```
## [1] 561
```

```
ncand <- length(cand1) + length(cand2) + length(cand3)
ncand
```

```
## [1] 1353
```

```
# set up results

cand1 <- cbind.data.frame(rep(1,times=length(cand1)), names(cand1), unname(cand1))
cand2 <- cbind.data.frame(rep(2,times=length(cand2)), names(cand2), unname(cand2))
cand3 <- cbind.data.frame(rep(3,times=length(cand3)), names(cand3), unname(cand3))

colnames(cand1) <- colnames(cand2) <- colnames(cand3) <- c("axis","snp","loading")

cand <- rbind(cand1, cand2, cand3)
cand$snp <- as.character(cand$snp)

# get correlations of SNPs with climate
tmp <- matrix(nrow=(ncand), ncol=ncol(clim.sub[,vars]))

colnames(tmp) <- colnames(clim.sub[,vars])

for (i in 1:length(cand$snp)) {
  
  nam <- cand$snp[i] # loop through candidate snp names
  snp.gen <- imp.sub[,nam]
  tmp[i,] <- apply(clim.sub[,vars], 2, function(x) cor(x,snp.gen))
}

cand <- cbind.data.frame(cand, tmp)
head(cand)
```

```
##   axis             snp    loading        bio5      bio6      bio15     bio18
## 1    1  1_283594_C_T_T 0.08302020 -0.02932193 0.5241133 -0.6656322 0.2206943
## 2    1  1_726642_T_C_C 0.08602280 -0.06776427 0.5237827 -0.7071993 0.2643742
## 3    1  1_729052_A_G_G 0.08865271 -0.16699782 0.5262292 -0.7282886 0.2429544
## 4    1  1_794347_C_A_A 0.08778933 -0.03966496 0.5782312 -0.6967663 0.2057635
## 5    1  1_795794_T_C_C 0.09159656 -0.04381487 0.5447065 -0.7455087 0.2689535
## 6    1 1_1753473_G_T_T 0.08090963 -0.24052835 0.4487518 -0.6878142 0.2541280
##        bio19      elev
## 1 -0.6499231 0.5317201
## 2 -0.6954256 0.5726904
## 3 -0.6682801 0.5776491
## 4 -0.7203935 0.5599467
## 5 -0.7299516 0.6020261
## 6 -0.5938398 0.5655850
```

```
#  check for duplicate snps - outliers on multiple axes
length(cand$snp[duplicated(cand$snp)]) # 0 - so skip below
```

```
## [1] 0
```

```
# foo <- cbind(cand$axis, duplicated(cand$snp)) 
# table(foo[foo[,1]==1,2]) # no duplicates on axis 1
# table(foo[foo[,1]==2,2]) #  6 duplicates on axis 2
# table(foo[foo[,1]==3,2]) # no duplicates on axis 3
# 
# cand <- cand[!duplicated(cand$snp),] # remove duplicate detections

dim(cand) # 1353 SNPs
```

```
## [1] 1353    9
```

```
# get best climate correlation for each snp
# note: need to modify the columns if different climate variables are used

for (i in 1:length(cand$snp)) {
  
  row <- cand[i,]
  cand[i, 10] <- names(which.max(abs(row[vars]))) # gives the variable
  cand[i, 11] <- max(abs(row[vars]))              # gives the correlation
}

colnames(cand)[10] <- "predictor"
colnames(cand)[11] <- "correlation"

table(cand$predictor)
```

```
## 
## bio15 bio18 bio19  bio5  bio6  elev 
##   585    55   222    36   437    18
```

```
# bio15 bio18 bio19  bio5  bio6  elev 
#    47    73   174    88    23   155

# bio15 bio18 bio19  bio5  bio6  elev 
#   585    55   222    36   437    18

########
# plot!

# colors
sel <- cand$snp
env <- cand$predictor

colset <- c("#d64b15","#013479","#ffb90f","#22a27c","#b1e3ad","#7d9ceb")
env[env=="bio5"] <- colset[1] # Max Temperature of Warmest Month
env[env=="bio6"] <- colset[2] # Min Temperature of Coldest Month
env[env=="bio15"] <- colset[3] # Precipitation Seasonality (Coefficient of Variation)
env[env=="bio18"] <- colset[4] # Precipitation of Warmest Quarter
env[env=="bio19"] <- colset[5] # Precipitation of Coldest Quarter
env[env=="elev"] <- colset[6]

# color by predictor:
col.pred <- rownames(rda$CCA$v) # pull the SNP names

for (i in 1:length(sel)) {           # color code candidate SNPs
  foo <- match(sel[i],col.pred)
  col.pred[foo] <- env[i]
}

# set color for non-candidate snps
# change if they don't already have a color
col.pred[! col.pred %in% colset] <- '#f1eef6' # non-candidate SNPs
empty <- col.pred
empty[grep("#f1eef6",empty)] <- rgb(0,1,0, alpha=0) # transparent
empty.outline <- ifelse(empty=="#00FF0000","#00FF0000","gray32")
bg <- colset

# now plot

# axes 1 & 2
#png(file = 'rda_plot_SNPs_clim_Qtom107.Qchr17.Qssp3_GEA_noQchr_PC1_PC2.png', width = 10, height = 9, res = 300, units = 'in')
plot(rda, type="n", scaling=3, xlim=c(-1,1), ylim=c(-1,1))
points(rda, display="species", pch=21, cex=1, col="gray32", bg=col.pred, scaling=3)
points(rda, display="species", pch=21, cex=1, col=empty.outline, bg=empty, scaling=3)
text(rda, scaling=3, display="bp", cex=1)
legend("bottomright", legend=vars, bty="n", col="gray32", pch=21, cex=1.2, pt.bg=bg)
```

```
#dev.off()

# axes 1 & 3
#png(file = 'rda_plot_SNPs_clim_Qtom107.Qchr17.Qssp3_GEA_noQchr_PC1_PC3.png', width = 10, height = 9, res = 300, units = 'in')
plot(rda, type="n", scaling=3, xlim=c(-1,1), ylim=c(-1,1), choices=c(1,3))
points(rda, display="species", pch=21, cex=1, col="gray32", bg=col.pred, scaling=3, choices=c(1,3))
points(rda, display="species", pch=21, cex=1, col=empty.outline, bg=empty, scaling=3, choices=c(1,3))
text(rda, scaling=3, display="bp", cex=1, choices=c(1,3))
legend("bottomright", legend=vars, bty="n", col="gray32", pch=21, cex=1.2, pt.bg=bg)
```

```
#dev.off()


# save

# write.table(cand, 'results/redundancy_analysis/RDA_candidate_climate_SNP_table_noQchr.csv', row.names = F, quote = F)
# save(cand, file = 'results/redundancy_analysis/RDA_candidate_climate_SNP_table_noQchr.rda')
```

```
cand.sort <- cand[order(cand$correlation, decreasing = T),]


# colors
cols <- c('black', "#3c4a8b","#009c85","#84bc5f","#edb829","#f57404","#b30000")

# plot just one
# snp <- cand.sort$snp[1]
# cli <- cand.sort$predictor[1]
# plot(jitter(snps.sub[,snp], 0.3), clim.sub[,cli],
#      main = snp,
#      ylab = cli,
#      xlab = 'genotype',
#      bg = cols[info.sub$island],
#      col = 'black',
#      pch = 21,
#      cex = 1)


# plot more
top <- min(96, nrow(cand.sort))

#dev.new()
par(mfrow = c(4,6))
par(mar = c(3,4,2,1))

for(n in 1:top){
  
  snp <- cand.sort$snp[n]
  cli <- cand.sort$predictor[n]
  plot(jitter(snps.sub[,snp], 0.3),
       clim.sub[,cli],
       main = snp,
       ylab = cli,
       xlab = 'genotype',
       bg = cols[info.sub$island],
       col = 'black',
       pch = 21,
       cex = 1.2)
}
```

```
# to save

# set = 1
# for(n in 1:top){
#   
#   # start a new plot every 24
#   if(n %% 24 == 1){
#     
#     png(file = paste('results/redundancy_analysis/RDA_SNP-climate_cors_noQchr_set_', set, '.png', sep = ''),
#         height = 10, width = 16,
#         units = 'in',
#         res = 300)
#     
#     par(mfrow = c(4,6))
#     par(mar = c(3,4,2,1))
#   }
# 
#   
#   snp <- cand.sort$snp[n]
#   cli <- cand.sort$predictor[n]
#   plot(jitter(snps.sub[,snp], 0.3),
#        clim.sub[,cli],
#        main = snp,
#        ylab = cli,
#        xlab = 'genotype',
#        bg = cols[info.sub$island],
#        col = 'black',
#        pch = 21,
#        cex = 1)
#   
#   if(n %% 24 == 0){
#     dev.off()
#     set = set+1
#   }
#   
# }

# looking at variation among islands to see how island structure might affect candidate SNPs
# if an island has both copies of a variant (it's not fixed), it will be easier for selection to act on it
# if islands are fixed for variants, it will be harder to tell whether differentiation is neutral or adaptive

# which islands have both copies of the variant?

snps_not_fixed <- as.data.frame(matrix(nrow = nrow(cand.sort), ncol = 6))
rownames(snps_not_fixed) <- cand.sort$snp
colnames(snps_not_fixed) <- c("Santa Rosa Island", "Santa Cruz Island", "Anacapa Island", "Catalina Island", "San Clemente Island", "Guadalupe Island")

for(n in 1:nrow(cand.sort)){
  
  snp <- cand.sort$snp[n]
  
  # df with snp copies and island
  df <- data.frame(copies = snps.sub[,snp], island = info.sub$island)
  
  # relevel island factors
  df$island <- factor(df$island, exclude = 'Mainland')
  
  # loop through islands
  for(i in 1:nlevels(df$island)){
    
    island <- levels(df$island)[i]
    df.isl <- df[df$island == island,]
    
    # which copy numbers exist in this island?
    copies <- na.omit(df.isl$copies)
    
    # does the variant vary within this island?
    # first check for heterozygotes
    if(1 %in% copies){
      not_fixed_bool <- 1
      #if no heterozygotes, check whether both homozygotes exist
    } else if(0 %in% copies & 2 %in% copies){
      not_fixed_bool <- 1
      # if not, variant is fixed on island
    } else{
      not_fixed_bool <- 0
    }
    
    # save to dataframe
    snps_not_fixed[snp, island] <- not_fixed_bool
    
  }
  
}


head(snps_not_fixed)
```

```
##                   Santa Rosa Island Santa Cruz Island Anacapa Island
## 11_17582750_A_C_C                 0                 0              0
## 2_8302636_G_A_A                   1                 0              0
## 8_23959598_A_G_G                  0                 0              0
## 6_8169577_A_G_G                   0                 0              0
## 5_22241589_C_G_G                  0                 0              0
## 4_58873603_T_C_T                  1                 1              0
##                   Catalina Island San Clemente Island Guadalupe Island
## 11_17582750_A_C_C               0                   0                1
## 2_8302636_G_A_A                 0                   0                1
## 8_23959598_A_G_G                0                   0                1
## 6_8169577_A_G_G                 0                   1                1
## 5_22241589_C_G_G                0                   0                1
## 4_58873603_T_C_T                0                   1                1
```

```
par(mfrow = c(1,1))
# number of islands with variation in each candidate SNP
# most SNPs are only variable on one island
# 114 SNPs (~8%) are variable at all 3 islands
hist(rowSums(snps_not_fixed))
```

```
table(rowSums(snps_not_fixed))
```

```
## 
##   0   1   2   3   4   5   6 
##   2 700 221 143  79  94 114
```

```
# how many snps are variable at each island?
# this is pretty consistent with observed heterozygosity metrics
barplot(colSums(snps_not_fixed))
```

### 6 GEA analysis, without Guadalupe or Mainaland Qchr

#### 6.1 RDA of SNPs and climate

```
# here we'll use the subset climate variables to do a more detailed GEA

# get subset without guadalupe
clim.sub <- clim[! info$island %in% c('Guadalupe Island', "Mainland"),]
snps.sub <- snps[! info$island %in% c('Guadalupe Island', "Mainland"),]
imp.sub <- imp[! info$island %in% c('Guadalupe Island', "Mainland"),]
info.sub <- info[! info$island %in% c('Guadalupe Island', "Mainland"),]

rda <- rda(imp.sub ~ ., data = as.data.frame(clim.sub[,vars]), scale = T)
screeplot(rda)
```

```
rda
```

```
## Call: rda(formula = imp.sub ~ bio5 + bio6 + bio15 + bio18 + bio19 +
## elev, data = as.data.frame(clim.sub[, vars]), scale = T)
## 
##                 Inertia Proportion Rank
## Total         5.702e+05  1.000e+00     
## Constrained   5.211e+04  9.139e-02    6
## Unconstrained 5.181e+05  9.086e-01   88
## Inertia is correlations 
## 
## Eigenvalues for constrained axes:
##  RDA1  RDA2  RDA3  RDA4  RDA5  RDA6 
## 11869  9308  8808  8179  7520  6428 
## 
## Eigenvalues for unconstrained axes:
##   PC1   PC2   PC3   PC4   PC5   PC6   PC7   PC8 
## 10309  9874  9491  8858  8582  8430  8380  8363 
## (Showing 8 of 88 unconstrained eigenvalues)
```

```
rda.info <- summary(rda)

head(rda.info)
```

```
## 
## Call:
## rda(formula = imp.sub ~ bio5 + bio6 + bio15 + bio18 + bio19 +      elev, data = as.data.frame(clim.sub[, vars]), scale = T) 
## 
## Partitioning of correlations:
##               Inertia Proportion
## Total          570231    1.00000
## Constrained     52112    0.09139
## Unconstrained  518119    0.90861
## 
## Eigenvalues, and their contribution to the correlations 
## 
## Importance of components:
##                            RDA1      RDA2      RDA3      RDA4      RDA5
## Eigenvalue            1.187e+04 9.308e+03 8.808e+03 8.179e+03 7.520e+03
## Proportion Explained  2.082e-02 1.632e-02 1.545e-02 1.434e-02 1.319e-02
## Cumulative Proportion 2.082e-02 3.714e-02 5.258e-02 6.693e-02 8.011e-02
##                            RDA6       PC1       PC2       PC3       PC4
## Eigenvalue            6.428e+03 1.031e+04 9.874e+03 9.491e+03 8.858e+03
## Proportion Explained  1.127e-02 1.808e-02 1.732e-02 1.664e-02 1.553e-02
## Cumulative Proportion 9.139e-02 1.095e-01 1.268e-01 1.434e-01 1.590e-01
##                             PC5       PC6       PC7       PC8       PC9
## Eigenvalue            8.582e+03 8.430e+03 8380.0964 8.363e+03 8.139e+03
## Proportion Explained  1.505e-02 1.478e-02    0.0147 1.467e-02 1.427e-02
## Cumulative Proportion 1.740e-01 1.888e-01    0.2035 2.182e-01 2.324e-01
##                            PC10      PC11      PC12      PC13      PC14
## Eigenvalue            8.032e+03 7.922e+03 7.830e+03 7.727e+03 7.667e+03
## Proportion Explained  1.409e-02 1.389e-02 1.373e-02 1.355e-02 1.345e-02
## Cumulative Proportion 2.465e-01 2.604e-01 2.741e-01 2.877e-01 3.011e-01
##                            PC15      PC16      PC17      PC18      PC19
## Eigenvalue            7.574e+03 7.501e+03 7187.6820 7.049e+03 6957.0585
## Proportion Explained  1.328e-02 1.315e-02    0.0126 1.236e-02    0.0122
## Cumulative Proportion 3.144e-01 3.276e-01    0.3402 3.525e-01    0.3647
##                            PC20      PC21      PC22      PC23      PC24
## Eigenvalue            6.938e+03 6.839e+03 6.768e+03 6.752e+03 6.676e+03
## Proportion Explained  1.217e-02 1.199e-02 1.187e-02 1.184e-02 1.171e-02
## Cumulative Proportion 3.769e-01 3.889e-01 4.008e-01 4.126e-01 4.243e-01
##                            PC25      PC26      PC27      PC28      PC29
## Eigenvalue            6.597e+03 6.591e+03 6.542e+03 6.494e+03 6.471e+03
## Proportion Explained  1.157e-02 1.156e-02 1.147e-02 1.139e-02 1.135e-02
## Cumulative Proportion 4.359e-01 4.474e-01 4.589e-01 4.703e-01 4.817e-01
##                            PC30      PC31      PC32      PC33      PC34
## Eigenvalue            6.417e+03 6.289e+03 6.252e+03 6.228e+03 6.145e+03
## Proportion Explained  1.125e-02 1.103e-02 1.096e-02 1.092e-02 1.078e-02
## Cumulative Proportion 4.929e-01 5.039e-01 5.149e-01 5.258e-01 5.366e-01
##                            PC35      PC36      PC37      PC38      PC39
## Eigenvalue            6.091e+03 6.019e+03 5.957e+03 5.883e+03 5.856e+03
## Proportion Explained  1.068e-02 1.056e-02 1.045e-02 1.032e-02 1.027e-02
## Cumulative Proportion 5.473e-01 5.578e-01 5.683e-01 5.786e-01 5.889e-01
##                            PC40      PC41      PC42      PC43      PC44
## Eigenvalue            5.831e+03 5.784e+03 5.763e+03 5.721e+03 5.692e+03
## Proportion Explained  1.022e-02 1.014e-02 1.011e-02 1.003e-02 9.982e-03
## Cumulative Proportion 5.991e-01 6.092e-01 6.193e-01 6.294e-01 6.394e-01
##                            PC45      PC46      PC47      PC48      PC49
## Eigenvalue            5.580e+03 5.551e+03 5.502e+03 5.453e+03 5.322e+03
## Proportion Explained  9.785e-03 9.736e-03 9.649e-03 9.563e-03 9.332e-03
## Cumulative Proportion 6.491e-01 6.589e-01 6.685e-01 6.781e-01 6.874e-01
##                            PC50      PC51      PC52      PC53      PC54
## Eigenvalue            5.280e+03 5.253e+03 5.185e+03 5.135e+03 5.131e+03
## Proportion Explained  9.259e-03 9.212e-03 9.093e-03 9.004e-03 8.998e-03
## Cumulative Proportion 6.967e-01 7.059e-01 7.150e-01 7.240e-01 7.330e-01
##                            PC55      PC56      PC57      PC58      PC59
## Eigenvalue            5.099e+03 5.078e+03 5.050e+03 5.038e+03 5.028e+03
## Proportion Explained  8.942e-03 8.905e-03 8.856e-03 8.834e-03 8.817e-03
## Cumulative Proportion 7.419e-01 7.508e-01 7.597e-01 7.685e-01 7.773e-01
##                            PC60      PC61      PC62      PC63      PC64
## Eigenvalue            4.966e+03 4961.0483 4.910e+03 4.885e+03 4.879e+03
## Proportion Explained  8.709e-03    0.0087 8.610e-03 8.566e-03 8.556e-03
## Cumulative Proportion 7.860e-01    0.7947 8.034e-01 8.119e-01 8.205e-01
##                            PC65      PC66      PC67      PC68      PC69
## Eigenvalue            4.857e+03 4.805e+03 4.789e+03 4.762e+03 4.736e+03
## Proportion Explained  8.518e-03 8.426e-03 8.398e-03 8.352e-03 8.306e-03
## Cumulative Proportion 8.290e-01 8.374e-01 8.458e-01 8.542e-01 8.625e-01
##                            PC70      PC71      PC72      PC73      PC74
## Eigenvalue            4.723e+03 4.673e+03 4.631e+03 4.561e+03 4.516e+03
## Proportion Explained  8.283e-03 8.194e-03 8.122e-03 7.998e-03 7.920e-03
## Cumulative Proportion 8.708e-01 8.790e-01 8.871e-01 8.951e-01 9.030e-01
##                            PC75      PC76      PC77      PC78      PC79
## Eigenvalue            4.508e+03 4.450e+03 4.423e+03 4390.8010 4.314e+03
## Proportion Explained  7.906e-03 7.803e-03 7.756e-03    0.0077 7.565e-03
## Cumulative Proportion 9.109e-01 9.187e-01 9.265e-01    0.9342 9.417e-01
##                            PC80      PC81      PC82      PC83      PC84
## Eigenvalue            4.172e+03 4.053e+03 3.982e+03 3.964e+03 3.855e+03
## Proportion Explained  7.316e-03 7.108e-03 6.984e-03 6.952e-03 6.760e-03
## Cumulative Proportion 9.490e-01 9.561e-01 9.631e-01 9.701e-01 9.768e-01
##                            PC85      PC86      PC87      PC88
## Eigenvalue            3.736e+03 3.665e+03 3.416e+03 2.387e+03
## Proportion Explained  6.551e-03 6.428e-03 5.990e-03 4.186e-03
## Cumulative Proportion 9.834e-01 9.898e-01 9.958e-01 1.000e+00
## 
## Accumulated constrained eigenvalues
## Importance of components:
##                            RDA1      RDA2      RDA3      RDA4      RDA5
## Eigenvalue            1.187e+04 9307.5358 8808.0109 8178.8570 7519.7626
## Proportion Explained  2.278e-01    0.1786    0.1690    0.1569    0.1443
## Cumulative Proportion 2.278e-01    0.4064    0.5754    0.7323    0.8766
##                            RDA6
## Eigenvalue            6428.2926
## Proportion Explained     0.1234
## Cumulative Proportion    1.0000
## 
## Scaling 2 for species and site scores
## * Species are scaled proportional to eigenvalues
## * Sites are unscaled: weighted dispersion equal on all dimensions
## * General scaling constant of scores:  85.56471 
## 
## 
## Species scores
## 
##                    RDA1      RDA2       RDA3       RDA4      RDA5      RDA6
## 1_39789_A_G_G -0.008055 -0.051751  0.0379551  0.0291603 -0.015609 -0.007749
## 1_52176_G_A_A  0.024407 -0.018225  0.0007159 -0.0040872 -0.011339  0.013889
## 1_52195_G_A_A -0.009237 -0.003146 -0.0073271 -0.0053383  0.002717 -0.004005
## 1_52213_C_T_T -0.004480  0.013425  0.0009239  0.0158889  0.010047  0.024934
## 1_52248_T_C_T -0.005649 -0.018725 -0.0055945 -0.0002107 -0.017787  0.024208
## 1_52331_G_T_T -0.021127  0.010435 -0.0028580 -0.0053897 -0.008405  0.013900
## ....                                                                       
## 
## 
## Site scores (weighted sums of species scores)
## 
##                  RDA1   RDA2    RDA3    RDA4   RDA5    RDA6
## Qchr.A.LA.215  -7.262 -4.097   4.571  0.5707 -4.229  -1.692
## Qchr.A.LA.222 -12.416 -5.079   4.102  1.3857 -3.945  -2.457
## Qchr.I.LA.20   -8.756 -2.287   1.063 -4.4137 17.532  -3.880
## Qchr.I.LA.25   -8.833 -6.958   2.281 -6.6717  5.819 -19.407
## Qchr.I.SB.02   -8.192 15.041 -27.803 -5.1101 14.371  -4.592
## Qchr.I.SB.03   -8.659 15.514 -28.630 -5.6871 14.247  -4.397
## ....                                                       
## 
## 
## Site constraints (linear combinations of constraining variables)
## 
##                 RDA1   RDA2    RDA3   RDA4   RDA5    RDA6
## Qchr.A.LA.215 -8.887 -3.614   4.670  1.434 -2.964  -1.829
## Qchr.A.LA.222 -8.887 -3.614   4.670  1.434 -2.964  -1.829
## Qchr.I.LA.20  -8.493 -1.248   2.645 -4.139 16.819  -3.221
## Qchr.I.LA.25  -9.029 -6.711   3.944 -6.445  5.521 -17.872
## Qchr.I.SB.02  -4.400 13.922 -24.238 -6.171 10.323  -2.914
## Qchr.I.SB.03  -4.400 13.922 -24.238 -6.171 10.323  -2.914
## ....                                                     
## 
## 
## Biplot scores for constraining variables
## 
##          RDA1     RDA2     RDA3    RDA4     RDA5    RDA6
## bio5  -0.9007 -0.12513 -0.27088  0.2400 -0.12568  0.1620
## bio6  -0.6632 -0.30750  0.61973 -0.1563  0.20855  0.1166
## bio15 -0.2979  0.70621  0.49984  0.1633  0.06676  0.3627
## bio18  0.8157 -0.36858 -0.32160  0.2522 -0.07298 -0.1626
## bio19  0.3969  0.62387 -0.02936  0.2640 -0.58780 -0.1929
## elev  -0.3895  0.04395 -0.68740  0.1113 -0.11299 -0.5905
```

```
rda.info$cont$importance
```

```
## Importance of components:
##                            RDA1      RDA2      RDA3      RDA4      RDA5
## Eigenvalue            1.187e+04 9.308e+03 8.808e+03 8.179e+03 7.520e+03
## Proportion Explained  2.082e-02 1.632e-02 1.545e-02 1.434e-02 1.319e-02
## Cumulative Proportion 2.082e-02 3.714e-02 5.258e-02 6.693e-02 8.011e-02
##                            RDA6       PC1       PC2       PC3       PC4
## Eigenvalue            6.428e+03 1.031e+04 9.874e+03 9.491e+03 8.858e+03
## Proportion Explained  1.127e-02 1.808e-02 1.732e-02 1.664e-02 1.553e-02
## Cumulative Proportion 9.139e-02 1.095e-01 1.268e-01 1.434e-01 1.590e-01
##                             PC5       PC6       PC7       PC8       PC9
## Eigenvalue            8.582e+03 8.430e+03 8380.0964 8.363e+03 8.139e+03
## Proportion Explained  1.505e-02 1.478e-02    0.0147 1.467e-02 1.427e-02
## Cumulative Proportion 1.740e-01 1.888e-01    0.2035 2.182e-01 2.324e-01
##                            PC10      PC11      PC12      PC13      PC14
## Eigenvalue            8.032e+03 7.922e+03 7.830e+03 7.727e+03 7.667e+03
## Proportion Explained  1.409e-02 1.389e-02 1.373e-02 1.355e-02 1.345e-02
## Cumulative Proportion 2.465e-01 2.604e-01 2.741e-01 2.877e-01 3.011e-01
##                            PC15      PC16      PC17      PC18      PC19
## Eigenvalue            7.574e+03 7.501e+03 7187.6820 7.049e+03 6957.0585
## Proportion Explained  1.328e-02 1.315e-02    0.0126 1.236e-02    0.0122
## Cumulative Proportion 3.144e-01 3.276e-01    0.3402 3.525e-01    0.3647
##                            PC20      PC21      PC22      PC23      PC24
## Eigenvalue            6.938e+03 6.839e+03 6.768e+03 6.752e+03 6.676e+03
## Proportion Explained  1.217e-02 1.199e-02 1.187e-02 1.184e-02 1.171e-02
## Cumulative Proportion 3.769e-01 3.889e-01 4.008e-01 4.126e-01 4.243e-01
##                            PC25      PC26      PC27      PC28      PC29
## Eigenvalue            6.597e+03 6.591e+03 6.542e+03 6.494e+03 6.471e+03
## Proportion Explained  1.157e-02 1.156e-02 1.147e-02 1.139e-02 1.135e-02
## Cumulative Proportion 4.359e-01 4.474e-01 4.589e-01 4.703e-01 4.817e-01
##                            PC30      PC31      PC32      PC33      PC34
## Eigenvalue            6.417e+03 6.289e+03 6.252e+03 6.228e+03 6.145e+03
## Proportion Explained  1.125e-02 1.103e-02 1.096e-02 1.092e-02 1.078e-02
## Cumulative Proportion 4.929e-01 5.039e-01 5.149e-01 5.258e-01 5.366e-01
##                            PC35      PC36      PC37      PC38      PC39
## Eigenvalue            6.091e+03 6.019e+03 5.957e+03 5.883e+03 5.856e+03
## Proportion Explained  1.068e-02 1.056e-02 1.045e-02 1.032e-02 1.027e-02
## Cumulative Proportion 5.473e-01 5.578e-01 5.683e-01 5.786e-01 5.889e-01
##                            PC40      PC41      PC42      PC43      PC44
## Eigenvalue            5.831e+03 5.784e+03 5.763e+03 5.721e+03 5.692e+03
## Proportion Explained  1.022e-02 1.014e-02 1.011e-02 1.003e-02 9.982e-03
## Cumulative Proportion 5.991e-01 6.092e-01 6.193e-01 6.294e-01 6.394e-01
##                            PC45      PC46      PC47      PC48      PC49
## Eigenvalue            5.580e+03 5.551e+03 5.502e+03 5.453e+03 5.322e+03
## Proportion Explained  9.785e-03 9.736e-03 9.649e-03 9.563e-03 9.332e-03
## Cumulative Proportion 6.491e-01 6.589e-01 6.685e-01 6.781e-01 6.874e-01
##                            PC50      PC51      PC52      PC53      PC54
## Eigenvalue            5.280e+03 5.253e+03 5.185e+03 5.135e+03 5.131e+03
## Proportion Explained  9.259e-03 9.212e-03 9.093e-03 9.004e-03 8.998e-03
## Cumulative Proportion 6.967e-01 7.059e-01 7.150e-01 7.240e-01 7.330e-01
##                            PC55      PC56      PC57      PC58      PC59
## Eigenvalue            5.099e+03 5.078e+03 5.050e+03 5.038e+03 5.028e+03
## Proportion Explained  8.942e-03 8.905e-03 8.856e-03 8.834e-03 8.817e-03
## Cumulative Proportion 7.419e-01 7.508e-01 7.597e-01 7.685e-01 7.773e-01
##                            PC60      PC61      PC62      PC63      PC64
## Eigenvalue            4.966e+03 4961.0483 4.910e+03 4.885e+03 4.879e+03
## Proportion Explained  8.709e-03    0.0087 8.610e-03 8.566e-03 8.556e-03
## Cumulative Proportion 7.860e-01    0.7947 8.034e-01 8.119e-01 8.205e-01
##                            PC65      PC66      PC67      PC68      PC69
## Eigenvalue            4.857e+03 4.805e+03 4.789e+03 4.762e+03 4.736e+03
## Proportion Explained  8.518e-03 8.426e-03 8.398e-03 8.352e-03 8.306e-03
## Cumulative Proportion 8.290e-01 8.374e-01 8.458e-01 8.542e-01 8.625e-01
##                            PC70      PC71      PC72      PC73      PC74
## Eigenvalue            4.723e+03 4.673e+03 4.631e+03 4.561e+03 4.516e+03
## Proportion Explained  8.283e-03 8.194e-03 8.122e-03 7.998e-03 7.920e-03
## Cumulative Proportion 8.708e-01 8.790e-01 8.871e-01 8.951e-01 9.030e-01
##                            PC75      PC76      PC77      PC78      PC79
## Eigenvalue            4.508e+03 4.450e+03 4.423e+03 4390.8010 4.314e+03
## Proportion Explained  7.906e-03 7.803e-03 7.756e-03    0.0077 7.565e-03
## Cumulative Proportion 9.109e-01 9.187e-01 9.265e-01    0.9342 9.417e-01
##                            PC80      PC81      PC82      PC83      PC84
## Eigenvalue            4.172e+03 4.053e+03 3.982e+03 3.964e+03 3.855e+03
## Proportion Explained  7.316e-03 7.108e-03 6.984e-03 6.952e-03 6.760e-03
## Cumulative Proportion 9.490e-01 9.561e-01 9.631e-01 9.701e-01 9.768e-01
##                            PC85      PC86      PC87      PC88
## Eigenvalue            3.736e+03 3.665e+03 3.416e+03 2.387e+03
## Proportion Explained  6.551e-03 6.428e-03 5.990e-03 4.186e-03
## Cumulative Proportion 9.834e-01 9.898e-01 9.958e-01 1.000e+00
```

```
rda.info$concont$importance
```

```
## Importance of components:
##                            RDA1      RDA2      RDA3      RDA4      RDA5
## Eigenvalue            1.187e+04 9307.5358 8808.0109 8178.8570 7519.7626
## Proportion Explained  2.278e-01    0.1786    0.1690    0.1569    0.1443
## Cumulative Proportion 2.278e-01    0.4064    0.5754    0.7323    0.8766
##                            RDA6
## Eigenvalue            6428.2926
## Proportion Explained     0.1234
## Cumulative Proportion    1.0000
```

```
RsquareAdj(rda)
```

```
## $r.squared
## [1] 0.09138727
## 
## $adj.r.squared
## [1] 0.0294364
```

```
# $r.squared
# [1] 0.09138727
# $adj.r.squared
# [1] 0.0294364

bg <- c('black', "#3c4a8b","#009c85","#84bc5f","#edb829","#f57404","#b30000")

plot_clim(rda, info = info.sub, save = F, file_string = 'Qtom107.Qchr17.Qssp3_GEA_noQchrGuadalupe')
```

```
plot_clim(rda, info = info.sub, choices = c(1,3), save = F, file_string = 'Qtom107.Qchr17.Qssp3_GEA_noQchrGuadalupe')
```

```
plot(rda, type = 'none')
points(rda, display = 'sites', pch = sp[info.sub$sp], bg = bg[info.sub$island], col = 'black', cex = 1.5)
text(rda, display = 'bp')
points(rda, display = 'species', pch = 21, cex = 1, col = 'gray32')
```

#### 6.2 Variance partitioning - only Channel Islands samples

```
# rdas
# without mainland Qchr  or guadalupe
clim.sub <- clim[! info$island %in% c('Guadalupe Island', "Mainland"),]
snps.sub <- snps[! info$island %in% c('Guadalupe Island', "Mainland"),]
imp.sub <- imp[! info$island %in% c('Guadalupe Island', "Mainland"),]
info.sub <- info[! info$island %in% c('Guadalupe Island', "Mainland"),]

# 2022-11-30: values are from all snps

# geography
lat <- info.sub$lat
lon <- info.sub$lon

geo <- cbind(lat, lon)

rda.full <- rda(imp.sub ~ clim.sub[,vars] + geo, scale = T)
RsquareAdj(rda.full) # 0.03796603
```

```
## $r.squared
## [1] 0.1198413
## 
## $adj.r.squared
## [1] 0.03796603
```

```
rda.full.info <- summary(rda.full)
head(rda.full.info)
```

```
## 
## Call:
## rda(formula = imp.sub ~ clim.sub[, vars] + geo, scale = T) 
## 
## Partitioning of correlations:
##               Inertia Proportion
## Total          570231     1.0000
## Constrained     68337     0.1198
## Unconstrained  501894     0.8802
## 
## Eigenvalues, and their contribution to the correlations 
## 
## Importance of components:
##                            RDA1      RDA2      RDA3      RDA4      RDA5
## Eigenvalue            1.247e+04 9.429e+03 9.017e+03 8.227e+03 8.081e+03
## Proportion Explained  2.188e-02 1.654e-02 1.581e-02 1.443e-02 1.417e-02
## Cumulative Proportion 2.188e-02 3.841e-02 5.422e-02 6.865e-02 8.282e-02
##                            RDA6      RDA7      RDA8       PC1       PC2
## Eigenvalue            7.566e+03 7184.1150 6.360e+03 9.881e+03 9.380e+03
## Proportion Explained  1.327e-02    0.0126 1.115e-02 1.733e-02 1.645e-02
## Cumulative Proportion 9.609e-02    0.1087 1.198e-01 1.372e-01 1.536e-01
##                             PC3       PC4       PC5       PC6       PC7
## Eigenvalue            9.320e+03 8.583e+03 8.518e+03 8.410e+03 8.344e+03
## Proportion Explained  1.634e-02 1.505e-02 1.494e-02 1.475e-02 1.463e-02
## Cumulative Proportion 1.700e-01 1.850e-01 2.000e-01 2.147e-01 2.293e-01
##                             PC8       PC9      PC10      PC11      PC12
## Eigenvalue            8.191e+03 8.035e+03 7.953e+03 7.828e+03 7698.8457
## Proportion Explained  1.436e-02 1.409e-02 1.395e-02 1.373e-02    0.0135
## Cumulative Proportion 2.437e-01 2.578e-01 2.717e-01 2.855e-01    0.2990
##                            PC13      PC14      PC15      PC16      PC17
## Eigenvalue            7.644e+03 7.553e+03 7.290e+03 7.067e+03 6.982e+03
## Proportion Explained  1.341e-02 1.324e-02 1.279e-02 1.239e-02 1.224e-02
## Cumulative Proportion 3.124e-01 3.256e-01 3.384e-01 3.508e-01 3.630e-01
##                            PC18      PC19      PC20      PC21      PC22
## Eigenvalue            6959.4056 6.940e+03 6.792e+03 6.767e+03 6.721e+03
## Proportion Explained     0.0122 1.217e-02 1.191e-02 1.187e-02 1.179e-02
## Cumulative Proportion    0.3752 3.874e-01 3.993e-01 4.112e-01 4.230e-01
##                            PC23      PC24      PC25      PC26      PC27
## Eigenvalue            6.646e+03 6.597e+03 6.545e+03 6.508e+03 6.473e+03
## Proportion Explained  1.165e-02 1.157e-02 1.148e-02 1.141e-02 1.135e-02
## Cumulative Proportion 4.346e-01 4.462e-01 4.577e-01 4.691e-01 4.804e-01
##                            PC28      PC29      PC30      PC31      PC32
## Eigenvalue            6.427e+03 6.375e+03 6.294e+03 6.240e+03 6.178e+03
## Proportion Explained  1.127e-02 1.118e-02 1.104e-02 1.094e-02 1.083e-02
## Cumulative Proportion 4.917e-01 5.029e-01 5.139e-01 5.249e-01 5.357e-01
##                            PC33      PC34      PC35      PC36      PC37
## Eigenvalue            6103.0555 6.032e+03 5.958e+03 5.907e+03 5873.1143
## Proportion Explained     0.0107 1.058e-02 1.045e-02 1.036e-02    0.0103
## Cumulative Proportion    0.5464 5.570e-01 5.674e-01 5.778e-01    0.5881
##                            PC38      PC39      PC40      PC41      PC42
## Eigenvalue            5.848e+03 5.794e+03 5.781e+03 5.721e+03 5.700e+03
## Proportion Explained  1.026e-02 1.016e-02 1.014e-02 1.003e-02 9.995e-03
## Cumulative Proportion 5.984e-01 6.085e-01 6.187e-01 6.287e-01 6.387e-01
##                            PC43      PC44      PC45      PC46      PC47
## Eigenvalue            5.690e+03 5.559e+03 5.503e+03 5.456e+03 5.374e+03
## Proportion Explained  9.978e-03 9.750e-03 9.650e-03 9.567e-03 9.424e-03
## Cumulative Proportion 6.487e-01 6.584e-01 6.681e-01 6.776e-01 6.870e-01
##                            PC48      PC49      PC50      PC51      PC52
## Eigenvalue            5.280e+03 5.264e+03 5.195e+03 5.144e+03 5.135e+03
## Proportion Explained  9.260e-03 9.232e-03 9.110e-03 9.020e-03 9.005e-03
## Cumulative Proportion 6.963e-01 7.055e-01 7.146e-01 7.237e-01 7.327e-01
##                            PC53      PC54      PC55      PC56      PC57
## Eigenvalue            5.101e+03 5.080e+03 5.065e+03 5.042e+03 5.039e+03
## Proportion Explained  8.946e-03 8.909e-03 8.883e-03 8.842e-03 8.837e-03
## Cumulative Proportion 7.416e-01 7.505e-01 7.594e-01 7.683e-01 7.771e-01
##                            PC58      PC59      PC60      PC61      PC62
## Eigenvalue            4.978e+03 4.964e+03 4.927e+03 4.886e+03 4.879e+03
## Proportion Explained  8.729e-03 8.705e-03 8.640e-03 8.569e-03 8.557e-03
## Cumulative Proportion 7.858e-01 7.945e-01 8.032e-01 8.117e-01 8.203e-01
##                            PC63      PC64      PC65      PC66      PC67
## Eigenvalue            4.859e+03 4.808e+03 4.791e+03 4.763e+03 4.739e+03
## Proportion Explained  8.520e-03 8.431e-03 8.403e-03 8.352e-03 8.310e-03
## Cumulative Proportion 8.288e-01 8.372e-01 8.456e-01 8.540e-01 8.623e-01
##                            PC68      PC69      PC70      PC71      PC72
## Eigenvalue            4.726e+03 4.673e+03 4.632e+03 4.586e+03 4.517e+03
## Proportion Explained  8.287e-03 8.194e-03 8.123e-03 8.042e-03 7.922e-03
## Cumulative Proportion 8.706e-01 8.788e-01 8.869e-01 8.950e-01 9.029e-01
##                            PC73      PC74      PC75      PC76      PC77
## Eigenvalue            4.509e+03 4.450e+03 4.423e+03 4.392e+03 4.316e+03
## Proportion Explained  7.907e-03 7.803e-03 7.757e-03 7.702e-03 7.569e-03
## Cumulative Proportion 9.108e-01 9.186e-01 9.263e-01 9.340e-01 9.416e-01
##                            PC78      PC79      PC80      PC81      PC82
## Eigenvalue            4.190e+03 4.053e+03 3.982e+03 3.965e+03 3.856e+03
## Proportion Explained  7.349e-03 7.108e-03 6.984e-03 6.954e-03 6.761e-03
## Cumulative Proportion 9.490e-01 9.561e-01 9.631e-01 9.700e-01 9.768e-01
##                            PC83      PC84      PC85      PC86
## Eigenvalue            3.736e+03 3.666e+03 3.416e+03 2.429e+03
## Proportion Explained  6.551e-03 6.428e-03 5.991e-03 4.261e-03
## Cumulative Proportion 9.833e-01 9.897e-01 9.957e-01 1.000e+00
## 
## Accumulated constrained eigenvalues
## Importance of components:
##                            RDA1      RDA2      RDA3      RDA4      RDA5
## Eigenvalue            1.247e+04 9428.8326 9017.4610 8226.5681 8080.8157
## Proportion Explained  1.825e-01    0.1380    0.1320    0.1204    0.1182
## Cumulative Proportion 1.825e-01    0.3205    0.4525    0.5728    0.6911
##                            RDA6      RDA7      RDA8
## Eigenvalue            7565.5887 7184.1150 6.360e+03
## Proportion Explained     0.1107    0.1051 9.307e-02
## Cumulative Proportion    0.8018    0.9069 1.000e+00
## 
## Scaling 2 for species and site scores
## * Species are scaled proportional to eigenvalues
## * Sites are unscaled: weighted dispersion equal on all dimensions
## * General scaling constant of scores:  85.56471 
## 
## 
## Species scores
## 
##                    RDA1      RDA2      RDA3      RDA4      RDA5       RDA6
## 1_39789_A_G_G -0.005026 -0.047544  0.002835  0.004142 -0.031003  0.0407572
## 1_52176_G_A_A  0.023079 -0.010780 -0.012243 -0.006299  0.005621  0.0176426
## 1_52195_G_A_A -0.002875  0.004146 -0.013156 -0.012219 -0.015585  0.0006336
## 1_52213_C_T_T  0.001386  0.007535  0.010452  0.012762 -0.018456 -0.0142270
## 1_52248_T_C_T -0.012586 -0.022423 -0.004376  0.010222  0.020775  0.0077072
## 1_52331_G_T_T -0.017130  0.009118  0.002193 -0.006516 -0.007212  0.0025296
## ....                                                                      
## 
## 
## Site scores (weighted sums of species scores)
## 
##                  RDA1   RDA2    RDA3    RDA4    RDA5    RDA6
## Qchr.A.LA.215  -9.371 -6.541   4.155   2.782   7.134   3.161
## Qchr.A.LA.222 -14.722 -7.303   3.392   3.567   6.999   3.051
## Qchr.I.LA.20   -5.068 -1.212  -1.652  -9.950 -14.248 -14.961
## Qchr.I.LA.25   -5.327 -5.019  -3.319 -12.697 -12.021  -2.663
## Qchr.I.SB.02  -11.989 24.198 -20.762   3.846  14.033 -14.872
## Qchr.I.SB.03  -12.746 24.923 -21.344   3.819  15.281 -14.908
## ....                                                        
## 
## 
## Site constraints (linear combinations of constraining variables)
## 
##                  RDA1    RDA2     RDA3    RDA4    RDA5    RDA6
## Qchr.A.LA.215 -12.599 -6.3544   4.6301   3.590   6.708   2.352
## Qchr.A.LA.222 -12.599 -6.3465   4.6220   3.584   6.703   2.361
## Qchr.I.LA.20   -3.256 -0.8607   0.2796  -9.760 -14.107 -14.533
## Qchr.I.LA.25   -4.152 -5.0259  -1.9952 -12.799 -11.943  -2.154
## Qchr.I.SB.02  -10.120 22.7209 -18.5605   2.058  14.127 -10.910
## Qchr.I.SB.03  -10.176 22.6013 -18.4349   2.186  14.323 -11.042
## ....                                                          
## 
## 
## Biplot scores for constraining variables
## 
##                          RDA1     RDA2    RDA3     RDA4     RDA5     RDA6
## clim.sub[, vars]bio5  -0.8390 -0.03745 -0.2851  0.15840 -0.34042  0.08457
## clim.sub[, vars]bio6  -0.6117 -0.44823  0.3982 -0.36969 -0.26943 -0.09439
## clim.sub[, vars]bio15 -0.2758  0.48256  0.7192  0.06102 -0.20041 -0.04081
## clim.sub[, vars]bio18  0.7574 -0.25819 -0.3879  0.36371  0.19587  0.08643
## clim.sub[, vars]bio19  0.3611  0.57118  0.2259  0.33874  0.13620  0.53071
## clim.sub[, vars]elev  -0.3673  0.25361 -0.5981  0.18573 -0.07227  0.03757
## geolat                 0.6352  0.46816  0.1007  0.37534  0.41151  0.22146
## geolon                -0.8402 -0.35343  0.1886 -0.23782 -0.21859 -0.14675
```

```
rda.full.info$cont$importance
```

```
## Importance of components:
##                            RDA1      RDA2      RDA3      RDA4      RDA5
## Eigenvalue            1.247e+04 9.429e+03 9.017e+03 8.227e+03 8.081e+03
## Proportion Explained  2.188e-02 1.654e-02 1.581e-02 1.443e-02 1.417e-02
## Cumulative Proportion 2.188e-02 3.841e-02 5.422e-02 6.865e-02 8.282e-02
##                            RDA6      RDA7      RDA8       PC1       PC2
## Eigenvalue            7.566e+03 7184.1150 6.360e+03 9.881e+03 9.380e+03
## Proportion Explained  1.327e-02    0.0126 1.115e-02 1.733e-02 1.645e-02
## Cumulative Proportion 9.609e-02    0.1087 1.198e-01 1.372e-01 1.536e-01
##                             PC3       PC4       PC5       PC6       PC7
## Eigenvalue            9.320e+03 8.583e+03 8.518e+03 8.410e+03 8.344e+03
## Proportion Explained  1.634e-02 1.505e-02 1.494e-02 1.475e-02 1.463e-02
## Cumulative Proportion 1.700e-01 1.850e-01 2.000e-01 2.147e-01 2.293e-01
##                             PC8       PC9      PC10      PC11      PC12
## Eigenvalue            8.191e+03 8.035e+03 7.953e+03 7.828e+03 7698.8457
## Proportion Explained  1.436e-02 1.409e-02 1.395e-02 1.373e-02    0.0135
## Cumulative Proportion 2.437e-01 2.578e-01 2.717e-01 2.855e-01    0.2990
##                            PC13      PC14      PC15      PC16      PC17
## Eigenvalue            7.644e+03 7.553e+03 7.290e+03 7.067e+03 6.982e+03
## Proportion Explained  1.341e-02 1.324e-02 1.279e-02 1.239e-02 1.224e-02
## Cumulative Proportion 3.124e-01 3.256e-01 3.384e-01 3.508e-01 3.630e-01
##                            PC18      PC19      PC20      PC21      PC22
## Eigenvalue            6959.4056 6.940e+03 6.792e+03 6.767e+03 6.721e+03
## Proportion Explained     0.0122 1.217e-02 1.191e-02 1.187e-02 1.179e-02
## Cumulative Proportion    0.3752 3.874e-01 3.993e-01 4.112e-01 4.230e-01
##                            PC23      PC24      PC25      PC26      PC27
## Eigenvalue            6.646e+03 6.597e+03 6.545e+03 6.508e+03 6.473e+03
## Proportion Explained  1.165e-02 1.157e-02 1.148e-02 1.141e-02 1.135e-02
## Cumulative Proportion 4.346e-01 4.462e-01 4.577e-01 4.691e-01 4.804e-01
##                            PC28      PC29      PC30      PC31      PC32
## Eigenvalue            6.427e+03 6.375e+03 6.294e+03 6.240e+03 6.178e+03
## Proportion Explained  1.127e-02 1.118e-02 1.104e-02 1.094e-02 1.083e-02
## Cumulative Proportion 4.917e-01 5.029e-01 5.139e-01 5.249e-01 5.357e-01
##                            PC33      PC34      PC35      PC36      PC37
## Eigenvalue            6103.0555 6.032e+03 5.958e+03 5.907e+03 5873.1143
## Proportion Explained     0.0107 1.058e-02 1.045e-02 1.036e-02    0.0103
## Cumulative Proportion    0.5464 5.570e-01 5.674e-01 5.778e-01    0.5881
##                            PC38      PC39      PC40      PC41      PC42
## Eigenvalue            5.848e+03 5.794e+03 5.781e+03 5.721e+03 5.700e+03
## Proportion Explained  1.026e-02 1.016e-02 1.014e-02 1.003e-02 9.995e-03
## Cumulative Proportion 5.984e-01 6.085e-01 6.187e-01 6.287e-01 6.387e-01
##                            PC43      PC44      PC45      PC46      PC47
## Eigenvalue            5.690e+03 5.559e+03 5.503e+03 5.456e+03 5.374e+03
## Proportion Explained  9.978e-03 9.750e-03 9.650e-03 9.567e-03 9.424e-03
## Cumulative Proportion 6.487e-01 6.584e-01 6.681e-01 6.776e-01 6.870e-01
##                            PC48      PC49      PC50      PC51      PC52
## Eigenvalue            5.280e+03 5.264e+03 5.195e+03 5.144e+03 5.135e+03
## Proportion Explained  9.260e-03 9.232e-03 9.110e-03 9.020e-03 9.005e-03
## Cumulative Proportion 6.963e-01 7.055e-01 7.146e-01 7.237e-01 7.327e-01
##                            PC53      PC54      PC55      PC56      PC57
## Eigenvalue            5.101e+03 5.080e+03 5.065e+03 5.042e+03 5.039e+03
## Proportion Explained  8.946e-03 8.909e-03 8.883e-03 8.842e-03 8.837e-03
## Cumulative Proportion 7.416e-01 7.505e-01 7.594e-01 7.683e-01 7.771e-01
##                            PC58      PC59      PC60      PC61      PC62
## Eigenvalue            4.978e+03 4.964e+03 4.927e+03 4.886e+03 4.879e+03
## Proportion Explained  8.729e-03 8.705e-03 8.640e-03 8.569e-03 8.557e-03
## Cumulative Proportion 7.858e-01 7.945e-01 8.032e-01 8.117e-01 8.203e-01
##                            PC63      PC64      PC65      PC66      PC67
## Eigenvalue            4.859e+03 4.808e+03 4.791e+03 4.763e+03 4.739e+03
## Proportion Explained  8.520e-03 8.431e-03 8.403e-03 8.352e-03 8.310e-03
## Cumulative Proportion 8.288e-01 8.372e-01 8.456e-01 8.540e-01 8.623e-01
##                            PC68      PC69      PC70      PC71      PC72
## Eigenvalue            4.726e+03 4.673e+03 4.632e+03 4.586e+03 4.517e+03
## Proportion Explained  8.287e-03 8.194e-03 8.123e-03 8.042e-03 7.922e-03
## Cumulative Proportion 8.706e-01 8.788e-01 8.869e-01 8.950e-01 9.029e-01
##                            PC73      PC74      PC75      PC76      PC77
## Eigenvalue            4.509e+03 4.450e+03 4.423e+03 4.392e+03 4.316e+03
## Proportion Explained  7.907e-03 7.803e-03 7.757e-03 7.702e-03 7.569e-03
## Cumulative Proportion 9.108e-01 9.186e-01 9.263e-01 9.340e-01 9.416e-01
##                            PC78      PC79      PC80      PC81      PC82
## Eigenvalue            4.190e+03 4.053e+03 3.982e+03 3.965e+03 3.856e+03
## Proportion Explained  7.349e-03 7.108e-03 6.984e-03 6.954e-03 6.761e-03
## Cumulative Proportion 9.490e-01 9.561e-01 9.631e-01 9.700e-01 9.768e-01
##                            PC83      PC84      PC85      PC86
## Eigenvalue            3.736e+03 3.666e+03 3.416e+03 2.429e+03
## Proportion Explained  6.551e-03 6.428e-03 5.991e-03 4.261e-03
## Cumulative Proportion 9.833e-01 9.897e-01 9.957e-01 1.000e+00
```

```
rda.full.info$concont$importance
```

```
## Importance of components:
##                            RDA1      RDA2      RDA3      RDA4      RDA5
## Eigenvalue            1.247e+04 9428.8326 9017.4610 8226.5681 8080.8157
## Proportion Explained  1.825e-01    0.1380    0.1320    0.1204    0.1182
## Cumulative Proportion 1.825e-01    0.3205    0.4525    0.5728    0.6911
##                            RDA6      RDA7      RDA8
## Eigenvalue            7565.5887 7184.1150 6.360e+03
## Proportion Explained     0.1107    0.1051 9.307e-02
## Cumulative Proportion    0.8018    0.9069 1.000e+00
```

```
# Partitioning of correlations:
#               Inertia Proportion
# Total          570231     1.0000
# Constrained     68337     0.1198
# Unconstrained  501894     0.8802
sig.full <- anova.cca(rda.full, permutations = 99)
sig.full
```

```
## Permutation test for rda under reduced model
## Permutation: free
## Number of permutations: 99
## 
## Model: rda(formula = imp.sub ~ clim.sub[, vars] + geo, scale = T)
##          Df Variance      F Pr(>F)   
## Model     8    68337 1.4637   0.01 **
## Residual 86   501894                 
## ---
## Signif. codes:  0 '***' 0.001 '**' 0.01 '*' 0.05 '.' 0.1 ' ' 1
```

```
# Permutation test for rda under reduced model
# Permutation: free
# Number of permutations: 99
# 
# Model: rda(formula = imp.sub ~ clim.sub[, vars] + geo, scale = T)
#          Df Variance      F Pr(>F)   
# Model     8    68337 1.4637   0.01 **
# Residual 86   501894                 
# ---
# Signif. codes:  0 ‘***’ 0.001 ‘**’ 0.01 ‘*’ 0.05 ‘.’ 0.1 ‘ ’ 1

rda.clim_not_geo <- rda(imp.sub ~ clim.sub[,vars] + Condition(geo), scale = T)
RsquareAdj(rda.clim_not_geo) # 0.02338488
```

```
## $r.squared
## [1] 0.08429375
## 
## $adj.r.squared
## [1] 0.02338488
```

```
sig.clim_not_geo <- anova.cca(rda.clim_not_geo, permutations = 99)
sig.clim_not_geo
```

```
## Permutation test for rda under reduced model
## Permutation: free
## Number of permutations: 99
## 
## Model: rda(formula = imp.sub ~ clim.sub[, vars] + Condition(geo), scale = T)
##          Df Variance      F Pr(>F)   
## Model     6    48067 1.3727   0.01 **
## Residual 86   501894                 
## ---
## Signif. codes:  0 '***' 0.001 '**' 0.01 '*' 0.05 '.' 0.1 ' ' 1
```

```
# Permutation test for rda under reduced model
# Permutation: free
# Number of permutations: 99
# 
# Model: rda(formula = imp.sub ~ clim.sub[, vars] + Condition(geo), scale = T)
#          Df Variance      F Pr(>F)   
# Model     6    48067 1.3727   0.01 **
# Residual 86   501894                 
# ---
# Signif. codes:  0 ‘***’ 0.001 ‘**’ 0.01 ‘*’ 0.05 ‘.’ 0.1 ‘ ’ 1

rda.geo_not_clim <- rda(imp.sub ~ geo + Condition(clim.sub[,vars]), scale = T)
RsquareAdj(rda.geo_not_clim) # 0.008529629
```

```
## $r.squared
## [1] 0.02845399
## 
## $adj.r.squared
## [1] 0.008529629
```

```
sig.geo_not_clim <- anova.cca(rda.geo_not_clim, permutations = 99)
sig.geo_not_clim
```

```
## Permutation test for rda under reduced model
## Permutation: free
## Number of permutations: 99
## 
## Model: rda(formula = imp.sub ~ geo + Condition(clim.sub[, vars]), scale = T)
##          Df Variance      F Pr(>F)   
## Model     2    16225 1.3901   0.01 **
## Residual 86   501894                 
## ---
## Signif. codes:  0 '***' 0.001 '**' 0.01 '*' 0.05 '.' 0.1 ' ' 1
```

```
# Permutation test for rda under reduced model
# Permutation: free
# Number of permutations: 99
# 
# Model: rda(formula = imp.sub ~ geo + Condition(clim.sub[, vars]), scale = T)
#          Df Variance      F Pr(>F)   
# Model     2    16225 1.3901   0.01 **
# Residual 86   501894                 
# ---
# Signif. codes:  0 ‘***’ 0.001 ‘**’ 0.01 ‘*’ 0.05 ‘.’ 0.1 ‘ ’ 1

rda.clim <- rda(imp.sub ~ clim.sub[,vars], scale = T)
RsquareAdj(rda.clim) #[1] 0.0294364
```

```
## $r.squared
## [1] 0.09138727
## 
## $adj.r.squared
## [1] 0.0294364
```

```
sig.clim <- anova.cca(rda.clim, permutations = 99)
sig.clim
```

```
## Permutation test for rda under reduced model
## Permutation: free
## Number of permutations: 99
## 
## Model: rda(formula = imp.sub ~ clim.sub[, vars], scale = T)
##          Df Variance      F Pr(>F)   
## Model     6    52112 1.4752   0.01 **
## Residual 88   518119                 
## ---
## Signif. codes:  0 '***' 0.001 '**' 0.01 '*' 0.05 '.' 0.1 ' ' 1
```

```
# Permutation test for rda under reduced model
# Permutation: free
# Number of permutations: 99
# 
# Model: rda(formula = imp.sub ~ clim.sub[, vars], scale = T)
#          Df Variance      F Pr(>F)   
# Model     6    52112 1.4752   0.01 **
# Residual 88   518119                 
# ---
# Signif. codes:  0 ‘***’ 0.001 ‘**’ 0.01 ‘*’ 0.05 ‘.’ 0.1 ‘ ’ 1

rda.geo <- rda(imp.sub ~ geo, scale = T)
RsquareAdj(rda.geo) # 0.01458115
```

```
## $r.squared
## [1] 0.03554751
## 
## $adj.r.squared
## [1] 0.01458115
```

```
sig.geo <- anova.cca(rda.geo, permutations = 99)
sig.geo
```

```
## Permutation test for rda under reduced model
## Permutation: free
## Number of permutations: 99
## 
## Model: rda(formula = imp.sub ~ geo, scale = T)
##          Df Variance      F Pr(>F)   
## Model     2    20270 1.6955   0.01 **
## Residual 92   549961                 
## ---
## Signif. codes:  0 '***' 0.001 '**' 0.01 '*' 0.05 '.' 0.1 ' ' 1
```

```
# Permutation test for rda under reduced model
# Permutation: free
# Number of permutations: 99
# 
# Model: rda(formula = imp.sub ~ geo, scale = T)
#          Df Variance      F Pr(>F)   
# Model     2    20270 1.6955   0.01 **
# Residual 92   549961                 
# ---
# Signif. codes:  0 ‘***’ 0.001 ‘**’ 0.01 ‘*’ 0.05 ‘.’ 0.1 ‘ ’ 1
```

#### 6.3 identify candidate SNPs from RDA outliers

```
# we identify climate SNPs as those that are outliers along the RDA axes
# then determine which climate variable each SNP is most associated with (because this is a multivariate analysis, they will often associate with multiple)

# again basically following Forester et al

# most variation explained by first three axes (~5% out of 9% unadjusted)
screeplot(rda)
```

```
summary(rda)$cont
```

```
## $importance
## Importance of components:
##                            RDA1      RDA2      RDA3      RDA4      RDA5
## Eigenvalue            1.187e+04 9.308e+03 8.808e+03 8.179e+03 7.520e+03
## Proportion Explained  2.082e-02 1.632e-02 1.545e-02 1.434e-02 1.319e-02
## Cumulative Proportion 2.082e-02 3.714e-02 5.258e-02 6.693e-02 8.011e-02
##                            RDA6       PC1       PC2       PC3       PC4
## Eigenvalue            6.428e+03 1.031e+04 9.874e+03 9.491e+03 8.858e+03
## Proportion Explained  1.127e-02 1.808e-02 1.732e-02 1.664e-02 1.553e-02
## Cumulative Proportion 9.139e-02 1.095e-01 1.268e-01 1.434e-01 1.590e-01
##                             PC5       PC6       PC7       PC8       PC9
## Eigenvalue            8.582e+03 8.430e+03 8380.0964 8.363e+03 8.139e+03
## Proportion Explained  1.505e-02 1.478e-02    0.0147 1.467e-02 1.427e-02
## Cumulative Proportion 1.740e-01 1.888e-01    0.2035 2.182e-01 2.324e-01
##                            PC10      PC11      PC12      PC13      PC14
## Eigenvalue            8.032e+03 7.922e+03 7.830e+03 7.727e+03 7.667e+03
## Proportion Explained  1.409e-02 1.389e-02 1.373e-02 1.355e-02 1.345e-02
## Cumulative Proportion 2.465e-01 2.604e-01 2.741e-01 2.877e-01 3.011e-01
##                            PC15      PC16      PC17      PC18      PC19
## Eigenvalue            7.574e+03 7.501e+03 7187.6820 7.049e+03 6957.0585
## Proportion Explained  1.328e-02 1.315e-02    0.0126 1.236e-02    0.0122
## Cumulative Proportion 3.144e-01 3.276e-01    0.3402 3.525e-01    0.3647
##                            PC20      PC21      PC22      PC23      PC24
## Eigenvalue            6.938e+03 6.839e+03 6.768e+03 6.752e+03 6.676e+03
## Proportion Explained  1.217e-02 1.199e-02 1.187e-02 1.184e-02 1.171e-02
## Cumulative Proportion 3.769e-01 3.889e-01 4.008e-01 4.126e-01 4.243e-01
##                            PC25      PC26      PC27      PC28      PC29
## Eigenvalue            6.597e+03 6.591e+03 6.542e+03 6.494e+03 6.471e+03
## Proportion Explained  1.157e-02 1.156e-02 1.147e-02 1.139e-02 1.135e-02
## Cumulative Proportion 4.359e-01 4.474e-01 4.589e-01 4.703e-01 4.817e-01
##                            PC30      PC31      PC32      PC33      PC34
## Eigenvalue            6.417e+03 6.289e+03 6.252e+03 6.228e+03 6.145e+03
## Proportion Explained  1.125e-02 1.103e-02 1.096e-02 1.092e-02 1.078e-02
## Cumulative Proportion 4.929e-01 5.039e-01 5.149e-01 5.258e-01 5.366e-01
##                            PC35      PC36      PC37      PC38      PC39
## Eigenvalue            6.091e+03 6.019e+03 5.957e+03 5.883e+03 5.856e+03
## Proportion Explained  1.068e-02 1.056e-02 1.045e-02 1.032e-02 1.027e-02
## Cumulative Proportion 5.473e-01 5.578e-01 5.683e-01 5.786e-01 5.889e-01
##                            PC40      PC41      PC42      PC43      PC44
## Eigenvalue            5.831e+03 5.784e+03 5.763e+03 5.721e+03 5.692e+03
## Proportion Explained  1.022e-02 1.014e-02 1.011e-02 1.003e-02 9.982e-03
## Cumulative Proportion 5.991e-01 6.092e-01 6.193e-01 6.294e-01 6.394e-01
##                            PC45      PC46      PC47      PC48      PC49
## Eigenvalue            5.580e+03 5.551e+03 5.502e+03 5.453e+03 5.322e+03
## Proportion Explained  9.785e-03 9.736e-03 9.649e-03 9.563e-03 9.332e-03
## Cumulative Proportion 6.491e-01 6.589e-01 6.685e-01 6.781e-01 6.874e-01
##                            PC50      PC51      PC52      PC53      PC54
## Eigenvalue            5.280e+03 5.253e+03 5.185e+03 5.135e+03 5.131e+03
## Proportion Explained  9.259e-03 9.212e-03 9.093e-03 9.004e-03 8.998e-03
## Cumulative Proportion 6.967e-01 7.059e-01 7.150e-01 7.240e-01 7.330e-01
##                            PC55      PC56      PC57      PC58      PC59
## Eigenvalue            5.099e+03 5.078e+03 5.050e+03 5.038e+03 5.028e+03
## Proportion Explained  8.942e-03 8.905e-03 8.856e-03 8.834e-03 8.817e-03
## Cumulative Proportion 7.419e-01 7.508e-01 7.597e-01 7.685e-01 7.773e-01
##                            PC60      PC61      PC62      PC63      PC64
## Eigenvalue            4.966e+03 4961.0483 4.910e+03 4.885e+03 4.879e+03
## Proportion Explained  8.709e-03    0.0087 8.610e-03 8.566e-03 8.556e-03
## Cumulative Proportion 7.860e-01    0.7947 8.034e-01 8.119e-01 8.205e-01
##                            PC65      PC66      PC67      PC68      PC69
## Eigenvalue            4.857e+03 4.805e+03 4.789e+03 4.762e+03 4.736e+03
## Proportion Explained  8.518e-03 8.426e-03 8.398e-03 8.352e-03 8.306e-03
## Cumulative Proportion 8.290e-01 8.374e-01 8.458e-01 8.542e-01 8.625e-01
##                            PC70      PC71      PC72      PC73      PC74
## Eigenvalue            4.723e+03 4.673e+03 4.631e+03 4.561e+03 4.516e+03
## Proportion Explained  8.283e-03 8.194e-03 8.122e-03 7.998e-03 7.920e-03
## Cumulative Proportion 8.708e-01 8.790e-01 8.871e-01 8.951e-01 9.030e-01
##                            PC75      PC76      PC77      PC78      PC79
## Eigenvalue            4.508e+03 4.450e+03 4.423e+03 4390.8010 4.314e+03
## Proportion Explained  7.906e-03 7.803e-03 7.756e-03    0.0077 7.565e-03
## Cumulative Proportion 9.109e-01 9.187e-01 9.265e-01    0.9342 9.417e-01
##                            PC80      PC81      PC82      PC83      PC84
## Eigenvalue            4.172e+03 4.053e+03 3.982e+03 3.964e+03 3.855e+03
## Proportion Explained  7.316e-03 7.108e-03 6.984e-03 6.952e-03 6.760e-03
## Cumulative Proportion 9.490e-01 9.561e-01 9.631e-01 9.701e-01 9.768e-01
##                            PC85      PC86      PC87      PC88
## Eigenvalue            3.736e+03 3.665e+03 3.416e+03 2.387e+03
## Proportion Explained  6.551e-03 6.428e-03 5.990e-03 4.186e-03
## Cumulative Proportion 9.834e-01 9.898e-01 9.958e-01 1.000e+00
```

```
# get SNP loadings for first 3 axes
load.rda <- scores(rda, choices=c(1:3), display="species")

hist(load.rda[,1], main="Loadings on RDA1")
```

```
hist(load.rda[,2], main="Loadings on RDA2")
```

```
hist(load.rda[,3], main="Loadings on RDA3")
```

```
outliers <- function(x,z){
  
  lims <- mean(x) + c(-1, 1) * z * sd(x) # find loadings +/-z sd from mean loading  
  x[x < lims[1] | x > lims[2]] # locus names in these tails
  
}

# start out with a cutoff of 4 std dev because there are a lot of snps
cand1 <- outliers(load.rda[,1],4)
cand2 <- outliers(load.rda[,2],4)
cand3 <- outliers(load.rda[,3],4)

length(cand1)
```

```
## [1] 157
```

```
length(cand2)
```

```
## [1] 85
```

```
length(cand3)
```

```
## [1] 1236
```

```
ncand <- length(cand1) + length(cand2) + length(cand3)
ncand
```

```
## [1] 1478
```

```
# set up results

cand1 <- cbind.data.frame(rep(1,times=length(cand1)), names(cand1), unname(cand1))
cand2 <- cbind.data.frame(rep(2,times=length(cand2)), names(cand2), unname(cand2))
cand3 <- cbind.data.frame(rep(3,times=length(cand3)), names(cand3), unname(cand3))

colnames(cand1) <- colnames(cand2) <- colnames(cand3) <- c("axis","snp","loading")

cand <- rbind(cand1, cand2, cand3)
cand$snp <- as.character(cand$snp)

# get correlations of SNPs with climate
tmp <- matrix(nrow=(ncand), ncol=ncol(clim.sub[,vars]))

colnames(tmp) <- colnames(clim.sub[,vars])

for (i in 1:length(cand$snp)) {
  
  nam <- cand$snp[i] # loop through candidate snp names
  snp.gen <- imp.sub[,nam]
  tmp[i,] <- apply(clim.sub[,vars], 2, function(x) cor(x,snp.gen))
}

cand <- cbind.data.frame(cand, tmp)
head(cand)
```

```
##   axis              snp     loading       bio5       bio6       bio15
## 1    1   1_388897_C_T_T  0.07824970 -0.5434050 -0.4321407 -0.66836185
## 2    1  1_2592491_G_T_T  0.08328580 -0.5262180 -0.6399280 -0.63674768
## 3    1 1_10042008_T_A_A  0.07115121 -0.6590658 -0.2823369 -0.15779921
## 4    1 1_22117242_C_T_C -0.06681278  0.5383435  0.3604462  0.05496143
## 5    1 1_29228382_T_G_G  0.06372350 -0.5202591 -0.1898527 -0.23606424
## 6    1 1_29262201_G_A_A  0.06287149 -0.5446988 -0.1433527 -0.25810341
##        bio18       bio19        elev
## 1  0.7610625  0.01069114 -0.15542737
## 2  0.8281477  0.22898575 -0.02937402
## 3  0.4437467  0.17368390 -0.33422075
## 4 -0.4305284 -0.30376622  0.32066769
## 5  0.4845506  0.08727551 -0.36548007
## 6  0.4660845  0.01364568 -0.37356470
```

```
#  check for duplicate snps - outliers on multiple axes
length(cand$snp[duplicated(cand$snp)]) # 6
```

```
## [1] 6
```

```
foo <- cbind(cand$axis, duplicated(cand$snp)) 
table(foo[foo[,1]==1,2]) # no duplicates on axis 1
```

```
## 
##   0 
## 157
```

```
table(foo[foo[,1]==2,2]) #  6 duplicates on axis 2
```

```
## 
##  0  1 
## 79  6
```

```
table(foo[foo[,1]==3,2]) # no duplicates on axis 3
```

```
## 
##    0 
## 1236
```

```
cand <- cand[!duplicated(cand$snp),] # remove duplicate detections

dim(cand) # 1472 SNPs
```

```
## [1] 1472    9
```

```
# get best climate correlation for each snp
# note: need to modify the columns if different climate variables are used

for (i in 1:length(cand$snp)) {
  
  row <- cand[i,]
  cand[i, 10] <- names(which.max(abs(row[vars]))) # gives the variable
  cand[i, 11] <- max(abs(row[vars]))              # gives the correlation
}

colnames(cand)[10] <- "predictor"
colnames(cand)[11] <- "correlation"

table(cand$predictor)
```

```
## 
## bio15 bio18 bio19  bio5  bio6  elev 
##   108    53    22   915    49   325
```

```
# bio15 bio18 bio19  bio5  bio6  elev 
#   108    53    22   915    49   325 

########
# plot!

# colors
sel <- cand$snp
env <- cand$predictor

colset <- c("#d64b15","#013479","#ffb90f","#22a27c","#b1e3ad","#7d9ceb")
env[env=="bio5"] <- colset[1] # Max Temperature of Warmest Month
env[env=="bio6"] <- colset[2] # Min Temperature of Coldest Month
env[env=="bio15"] <- colset[3] # Precipitation Seasonality (Coefficient of Variation)
env[env=="bio18"] <- colset[4] # Precipitation of Warmest Quarter
env[env=="bio19"] <- colset[5] # Precipitation of Coldest Quarter
env[env=="elev"] <- colset[6]

# color by predictor:
col.pred <- rownames(rda$CCA$v) # pull the SNP names

for (i in 1:length(sel)) {           # color code candidate SNPs
  foo <- match(sel[i],col.pred)
  col.pred[foo] <- env[i]
}

# set color for non-candidate snps
# change if they don't already have a color
col.pred[! col.pred %in% colset] <- '#f1eef6' # non-candidate SNPs
empty <- col.pred
empty[grep("#f1eef6",empty)] <- rgb(0,1,0, alpha=0) # transparent
empty.outline <- ifelse(empty=="#00FF0000","#00FF0000","gray32")
bg <- colset

# now plot

# axes 1 & 2
#png(file = 'rda_plot_SNPs_clim_Qtom107.Qchr17.Qssp3_GEA_noQchrGuadalupe_PC1_PC2.png', width = 10, height = 9, res = 300, units = 'in')
plot(rda, type="n", scaling=3, xlim=c(-1,1), ylim=c(-1,1))
points(rda, display="species", pch=21, cex=1, col="gray32", bg=col.pred, scaling=3)
points(rda, display="species", pch=21, cex=1, col=empty.outline, bg=empty, scaling=3)
text(rda, scaling=3, display="bp", cex=1)
legend("bottomright", legend=vars, bty="n", col="gray32", pch=21, cex=1.2, pt.bg=bg)
```

```
#dev.off()

# axes 1 & 3
png(file = 'rda_plot_SNPs_clim_Qtom107.Qchr17.Qssp3_GEA_noQchrGuadalupe_PC1_PC3.png', width = 10, height = 9, res = 300, units = 'in')
plot(rda, type="n", scaling=3, xlim=c(-1,1), ylim=c(-1,1), choices=c(1,3))
points(rda, display="species", pch=21, cex=1, col="gray32", bg=col.pred, scaling=3, choices=c(1,3))
points(rda, display="species", pch=21, cex=1, col=empty.outline, bg=empty, scaling=3, choices=c(1,3))
text(rda, scaling=3, display="bp", cex=1, choices=c(1,3))
legend("bottomright", legend=vars, bty="n", col="gray32", pch=21, cex=1.2, pt.bg=bg)
dev.off()
```

```
## png 
##   2
```

```
cand.sort <- cand[order(cand$correlation, decreasing = T),]


# colors
cols <- c('black', "#3c4a8b","#009c85","#84bc5f","#edb829","#f57404","#b30000")

# plot just one
# snp <- cand.sort$snp[1]
# cli <- cand.sort$predictor[1]
# plot(jitter(snps.sub[,snp], 0.3), clim.sub[,cli],
#      main = snp,
#      ylab = cli,
#      xlab = 'genotype',
#      bg = cols[info.sub$island],
#      col = 'black',
#      pch = 21,
#      cex = 1)


# plot more
top <- min(96, nrow(cand.sort))

#dev.new()
par(mfrow = c(4,6))
par(mar = c(3,4,2,1))

for(n in 1:top){
  
  snp <- cand.sort$snp[n]
  cli <- cand.sort$predictor[n]
  plot(jitter(snps.sub[,snp], 0.3),
       clim.sub[,cli],
       main = snp,
       ylab = cli,
       xlab = 'genotype',
       bg = cols[info.sub$island],
       col = 'black',
       pch = 21,
       cex = 1)
}
```

```
# to save

# set = 1
# for(n in 1:top){
#   
#   # start a new plot every 24
#   if(n %% 24 == 1){
#     
#     # png(file = paste('RDA_SNP-climate_cors_set_', set, '.png', sep = ''),
#     #     height = 10, width = 16,
#     #     units = 'in',
#     #     res = 300)
#     
#     par(mfrow = c(4,6))
#     par(mar = c(3,4,2,1))
#   }
# 
#   
#   snp <- cand.sort$snp[n]
#   cli <- cand.sort$predictor[n]
#   plot(jitter(snps.sub[,snp], 0.3),
#        clim.sub[,cli],
#        main = snp,
#        ylab = cli,
#        xlab = 'genotype',
#        bg = cols[info.sub$island],
#        col = 'black',
#        pch = 21,
#        cex = 1)
#   
#   if(n %% 24 == 0){
#     #dev.off()
#     set = set+1
#   }
#   
# }
```

```
# write.table(cand.sort, 'results/RDA_candidate_climate_SNP_table_noGuad.csv', row.names = F, quote = F)
# save(cand.sort, file = 'results/RDA_candidate_climate_SNP_table_noGuad.rda')
```

### 7 GEA analysis, only Guadalupe

#### 7.1 RDA of SNPs and climate

```
par(mfrow = c(1,1))

# here we'll use the subset climate variables to do a more detailed GEA

# get subset, only guadalupe
clim.sub <- clim[info$island == 'Guadalupe Island',]
snps.sub <- snps[info$island == 'Guadalupe Island',]
imp.sub <- imp[info$island == 'Guadalupe Island',]
info.sub <- info[info$island == 'Guadalupe Island',]

rda <- rda(imp.sub ~ ., data = as.data.frame(clim.sub[,vars]), scale = T)
screeplot(rda)
```

```
rda
```

```
## Call: rda(formula = imp.sub ~ bio5 + bio6 + bio15 + bio18 + bio19 +
## elev, data = as.data.frame(clim.sub[, vars]), scale = T)
## 
##                 Inertia Proportion Rank
## Total         3.782e+05  1.000e+00     
## Constrained   1.287e+05  3.404e-01    6
## Unconstrained 2.495e+05  6.596e-01   15
## Inertia is correlations 
## 
## Eigenvalues for constrained axes:
##  RDA1  RDA2  RDA3  RDA4  RDA5  RDA6 
## 26907 23979 22696 21655 16947 16532 
## 
## Eigenvalues for unconstrained axes:
##   PC1   PC2   PC3   PC4   PC5   PC6   PC7   PC8   PC9  PC10  PC11  PC12  PC13 
## 19953 18775 18601 17901 17809 17191 16721 16536 16376 16351 15960 15181 14628 
##  PC14  PC15 
## 14497 12988
```

```
rda.info <- summary(rda)
head(rda.info)
```

```
## 
## Call:
## rda(formula = imp.sub ~ bio5 + bio6 + bio15 + bio18 + bio19 +      elev, data = as.data.frame(clim.sub[, vars]), scale = T) 
## 
## Partitioning of correlations:
##               Inertia Proportion
## Total          378183     1.0000
## Constrained    128716     0.3404
## Unconstrained  249467     0.6596
## 
## Eigenvalues, and their contribution to the correlations 
## 
## Importance of components:
##                            RDA1      RDA2      RDA3      RDA4      RDA5
## Eigenvalue            2.691e+04 2.398e+04 2.270e+04 2.165e+04 1.695e+04
## Proportion Explained  7.115e-02 6.341e-02 6.001e-02 5.726e-02 4.481e-02
## Cumulative Proportion 7.115e-02 1.346e-01 1.946e-01 2.518e-01 2.966e-01
##                            RDA6       PC1       PC2       PC3       PC4
## Eigenvalue            1.653e+04 1.995e+04 1.877e+04 1.860e+04 1.790e+04
## Proportion Explained  4.371e-02 5.276e-02 4.964e-02 4.918e-02 4.733e-02
## Cumulative Proportion 3.404e-01 3.931e-01 4.428e-01 4.919e-01 5.393e-01
##                             PC5       PC6       PC7       PC8       PC9
## Eigenvalue            1.781e+04 1.719e+04 1.672e+04 1.654e+04 1.638e+04
## Proportion Explained  4.709e-02 4.546e-02 4.421e-02 4.372e-02 4.330e-02
## Cumulative Proportion 5.864e-01 6.318e-01 6.760e-01 7.198e-01 7.631e-01
##                            PC10      PC11      PC12      PC13      PC14
## Eigenvalue            1.635e+04 1.596e+04 1.518e+04 1.463e+04 1.450e+04
## Proportion Explained  4.324e-02 4.220e-02 4.014e-02 3.868e-02 3.833e-02
## Cumulative Proportion 8.063e-01 8.485e-01 8.886e-01 9.273e-01 9.657e-01
##                            PC15
## Eigenvalue            1.299e+04
## Proportion Explained  3.434e-02
## Cumulative Proportion 1.000e+00
## 
## Accumulated constrained eigenvalues
## Importance of components:
##                            RDA1      RDA2      RDA3      RDA4      RDA5
## Eigenvalue            26906.863 2.398e+04 2.270e+04 2.165e+04 1.695e+04
## Proportion Explained      0.209 1.863e-01 1.763e-01 1.682e-01 1.317e-01
## Cumulative Proportion     0.209 3.953e-01 5.717e-01 7.399e-01 8.716e-01
##                            RDA6
## Eigenvalue            1.653e+04
## Proportion Explained  1.284e-01
## Cumulative Proportion 1.000e+00
## 
## Scaling 2 for species and site scores
## * Species are scaled proportional to eigenvalues
## * Sites are unscaled: weighted dispersion equal on all dimensions
## * General scaling constant of scores:  53.08604 
## 
## 
## Species scores
## 
##                     RDA1       RDA2       RDA3       RDA4       RDA5      RDA6
## 1_39789_A_G_G  2.936e-05 -7.396e-03 -3.894e-02  1.367e-02  1.000e-02 7.793e-04
## 1_52176_G_A_A -3.459e-14  6.308e-14  1.746e-13  4.040e-14 -1.020e-13 1.478e-13
## 1_52195_G_A_A -1.106e-02 -1.303e-03 -4.028e-03 -8.140e-04 -4.954e-03 4.201e-04
## 1_52213_C_T_T  1.494e-02  9.032e-03  3.923e-02  4.119e-02 -2.743e-03 3.055e-04
## 1_52248_T_C_T -1.106e-02 -1.303e-03 -4.028e-03 -8.140e-04 -4.954e-03 4.201e-04
## 1_52331_G_T_T -7.710e-13 -6.696e-13 -1.320e-12 -1.714e-12  4.264e-14 7.828e-14
## ....                                                                          
## 
## 
## Site scores (weighted sums of species scores)
## 
##                RDA1    RDA2   RDA3    RDA4   RDA5    RDA6
## Qtom.I.BC.10 -6.825 -0.5006 -2.113 -0.8155 -2.512 0.88797
## Qtom.I.BC.11 -7.784 -1.1121 -2.935 -0.3909 -4.551 0.25464
## Qtom.I.BC.12 -6.928 -1.1167 -2.527 -0.2301 -3.471 0.04438
## Qtom.I.BC.13 -6.967 -0.9267 -2.610 -0.6376 -3.533 0.49858
## Qtom.I.BC.18 -7.054 -0.6024 -2.154 -0.6163 -3.312 0.28974
## Qtom.I.BC.20  9.154  5.4986 23.462 23.9776 -1.674 0.36477
## ....                                                     
## 
## 
## Site constraints (linear combinations of constraining variables)
## 
##                RDA1    RDA2  RDA3    RDA4   RDA5   RDA6
## Qtom.I.BC.10 -6.644 -0.7828 -2.42 -0.4891 -2.977 0.2524
## Qtom.I.BC.11 -6.644 -0.7828 -2.42 -0.4891 -2.977 0.2524
## Qtom.I.BC.12 -6.644 -0.7828 -2.42 -0.4891 -2.977 0.2524
## Qtom.I.BC.13 -6.644 -0.7828 -2.42 -0.4891 -2.977 0.2524
## Qtom.I.BC.18 -6.644 -0.7828 -2.42 -0.4891 -2.977 0.2524
## Qtom.I.BC.20  8.978  5.4269 23.57 24.7505 -1.648 0.1836
## ....                                                   
## 
## 
## Biplot scores for constraining variables
## 
##            RDA1    RDA2     RDA3     RDA4      RDA5     RDA6
## bio5   0.934178 -0.1987  0.07282 -0.28702 -0.010677 -0.00404
## bio6  -0.573299 -0.7640 -0.20217  0.13751 -0.054810  0.15739
## bio15  0.361631 -0.9005 -0.11533  0.01842 -0.064110  0.20148
## bio18  0.240748  0.9473  0.16560 -0.07198  0.009145 -0.10909
## bio19 -0.578082  0.7720 -0.16613 -0.03666  0.067014  0.19085
## elev  -0.007887  0.9597  0.19139 -0.07418  0.027312 -0.18999
```

```
RsquareAdj(rda)
```

```
## $r.squared
## [1] 0.3403542
## 
## $adj.r.squared
## [1] 0.07649595
```

```
# $r.squared
# [1] 0.3403542
# $adj.r.squared
# [1] 0.07649595

bg <- c('black', "#3c4a8b","#009c85","#84bc5f","#edb829","#f57404","#b30000")

plot_clim(rda, info = info.sub, save = F, file_string = 'Qtom107.Qchr17.Qssp3_GEA_onlyGuadalupe')
```

```
plot_clim(rda, info = info.sub, choices = c(1,3), save = F, file_string = 'Qtom107.Qchr17.Qssp3_onlyGuadalupe')
```

```
plot(rda, type = 'none')
points(rda, display = 'sites', pch = sp[info.sub$sp], bg = bg[info.sub$island], col = 'black', cex = 1.5)
text(rda, display = 'bp')
points(rda, display = 'species', pch = 21, cex = 1, col = 'gray32')
```

##### 7.1.1 identify candidate SNPs from RDA outliers

```
# we identify climate SNPs as those that are outliers along the RDA axes
# then determine which climate variable each SNP is most associated with (because this is a multivariate analysis, they will often associate with multiple)

# again basically following Forester et al

# most variation explained by first three axes (~20% out of 34% unadjusted)
# a lot more variation explained here than in other sample sets
screeplot(rda)
```

```
summary(rda)$cont
```

```
## $importance
## Importance of components:
##                            RDA1      RDA2      RDA3      RDA4      RDA5
## Eigenvalue            2.691e+04 2.398e+04 2.270e+04 2.165e+04 1.695e+04
## Proportion Explained  7.115e-02 6.341e-02 6.001e-02 5.726e-02 4.481e-02
## Cumulative Proportion 7.115e-02 1.346e-01 1.946e-01 2.518e-01 2.966e-01
##                            RDA6       PC1       PC2       PC3       PC4
## Eigenvalue            1.653e+04 1.995e+04 1.877e+04 1.860e+04 1.790e+04
## Proportion Explained  4.371e-02 5.276e-02 4.964e-02 4.918e-02 4.733e-02
## Cumulative Proportion 3.404e-01 3.931e-01 4.428e-01 4.919e-01 5.393e-01
##                             PC5       PC6       PC7       PC8       PC9
## Eigenvalue            1.781e+04 1.719e+04 1.672e+04 1.654e+04 1.638e+04
## Proportion Explained  4.709e-02 4.546e-02 4.421e-02 4.372e-02 4.330e-02
## Cumulative Proportion 5.864e-01 6.318e-01 6.760e-01 7.198e-01 7.631e-01
##                            PC10      PC11      PC12      PC13      PC14
## Eigenvalue            1.635e+04 1.596e+04 1.518e+04 1.463e+04 1.450e+04
## Proportion Explained  4.324e-02 4.220e-02 4.014e-02 3.868e-02 3.833e-02
## Cumulative Proportion 8.063e-01 8.485e-01 8.886e-01 9.273e-01 9.657e-01
##                            PC15
## Eigenvalue            1.299e+04
## Proportion Explained  3.434e-02
## Cumulative Proportion 1.000e+00
```

```
# get SNP loadings for first 3 axes
load.rda <- scores(rda, choices=c(1:3), display="species")

hist(load.rda[,1], main="Loadings on RDA1")
```

```
hist(load.rda[,2], main="Loadings on RDA2")
```

```
hist(load.rda[,3], main="Loadings on RDA3")
```

```
outliers <- function(x,z){
  
  lims <- mean(x) + c(-1, 1) * z * sd(x) # find loadings +/-z sd from mean loading  
  x[x < lims[1] | x > lims[2]] # locus names in these tails
  
}

# start out with a cutoff of 4 std dev because there are a lot of snps
cand1 <- outliers(load.rda[,1],4)
cand2 <- outliers(load.rda[,2],4)
cand3 <- outliers(load.rda[,3],4)

length(cand1)
```

```
## [1] 129
```

```
length(cand2)
```

```
## [1] 11
```

```
length(cand3)
```

```
## [1] 206
```

```
ncand <- length(cand1) + length(cand2) + length(cand3)
ncand # 346
```

```
## [1] 346
```

```
# set up results

cand1 <- cbind.data.frame(rep(1,times=length(cand1)), names(cand1), unname(cand1))
cand2 <- cbind.data.frame(rep(2,times=length(cand2)), names(cand2), unname(cand2))
cand3 <- cbind.data.frame(rep(3,times=length(cand3)), names(cand3), unname(cand3))

colnames(cand1) <- colnames(cand2) <- colnames(cand3) <- c("axis","snp","loading")

cand <- rbind(cand1, cand2, cand3)
cand$snp <- as.character(cand$snp)

# get correlations of SNPs with climate
tmp <- matrix(nrow=(ncand), ncol=ncol(clim.sub[,vars]))

colnames(tmp) <- colnames(clim.sub[,vars])

for (i in 1:length(cand$snp)) {
  
  nam <- cand$snp[i] # loop through candidate snp names
  snp.gen <- imp.sub[,nam]
  tmp[i,] <- apply(clim.sub[,vars], 2, function(x) cor(x,snp.gen))
}

cand <- cbind.data.frame(cand, tmp)
head(cand)
```

```
##   axis              snp    loading      bio5       bio6     bio15     bio18
## 1    1  1_2838327_A_G_G 0.07678457 0.9464671 -0.5649215 0.3054630 0.2464700
## 2    1  1_3685996_A_G_G 0.07678457 0.9464671 -0.5649215 0.3054630 0.2464700
## 3    1  1_4047380_G_T_T 0.07678457 0.9464671 -0.5649215 0.3054630 0.2464700
## 4    1  1_5271048_G_A_G 0.07678457 0.9464671 -0.5649215 0.3054630 0.2464700
## 5    1 1_25808350_A_C_C 0.07678457 0.9464671 -0.5649215 0.3054630 0.2464700
## 6    1 1_29390446_G_A_A 0.07649059 0.6904631 -0.5048597 0.2655399 0.2502808
##       bio19       elev
## 1 -0.480727 0.02697918
## 2 -0.480727 0.02697918
## 3 -0.480727 0.02697918
## 4 -0.480727 0.02697918
## 5 -0.480727 0.02697918
## 6 -0.498975 0.03220206
```

```
#  check for duplicate snps - outliers on multiple axes
length(cand$snp[duplicated(cand$snp)]) # 0
```

```
## [1] 0
```

```
# no duplicates, so skip below

# foo <- cbind(cand$axis, duplicated(cand$snp)) 
# table(foo[foo[,1]==1,2]) 
# table(foo[foo[,1]==2,2])
# table(foo[foo[,1]==3,2])
# 
# cand <- cand[!duplicated(cand$snp),] # remove duplicate detections

dim(cand) # 346 SNPs
```

```
## [1] 346   9
```

```
# get best climate correlation for each snp
# note: need to modify the columns if different climate variables are used

for (i in 1:length(cand$snp)) {
  
  row <- cand[i,]
  cand[i, 10] <- names(which.max(abs(row[vars]))) # gives the variable
  cand[i, 11] <- max(abs(row[vars]))              # gives the correlation
}

colnames(cand)[10] <- "predictor"
colnames(cand)[11] <- "correlation"

table(cand$predictor)
```

```
## 
## bio15 bio19  bio5  bio6  elev 
##     2     2   129   204     9
```

```
# bio15 bio19  bio5  bio6  elev 
#     2     2   129   204     9 

########
# plot!

# colors
sel <- cand$snp
env <- cand$predictor

colset <- c("#d64b15","#013479","#ffb90f","#22a27c","#b1e3ad","#7d9ceb")
env[env=="bio5"] <- colset[1] # Max Temperature of Warmest Month
env[env=="bio6"] <- colset[2] # Min Temperature of Coldest Month
env[env=="bio15"] <- colset[3] # Precipitation Seasonality (Coefficient of Variation)
env[env=="bio18"] <- colset[4] # Precipitation of Warmest Quarter
env[env=="bio19"] <- colset[5] # Precipitation of Coldest Quarter
env[env=="elev"] <- colset[6]

# color by predictor:
col.pred <- rownames(rda$CCA$v) # pull the SNP names

for (i in 1:length(sel)) {           # color code candidate SNPs
  foo <- match(sel[i],col.pred)
  col.pred[foo] <- env[i]
}

# set color for non-candidate snps
# change if they don't already have a color
col.pred[! col.pred %in% colset] <- '#f1eef6' # non-candidate SNPs
empty <- col.pred
empty[grep("#f1eef6",empty)] <- rgb(0,1,0, alpha=0) # transparent
empty.outline <- ifelse(empty=="#00FF0000","#00FF0000","gray32")
bg <- colset

# now plot

# axes 1 & 2
#png(file = 'rda_plot_SNPs_clim_Qtom107.Qchr17.Qssp3_GEA_onlyGuadalupe_PC1_PC2.png', width = 10, height = 9, res = 300, units = 'in')
plot(rda, type="n", scaling=3, xlim=c(-1,1), ylim=c(-1,1))
points(rda, display="species", pch=21, cex=1, col="gray32", bg=col.pred, scaling=3)
points(rda, display="species", pch=21, cex=1, col=empty.outline, bg=empty, scaling=3)
text(rda, scaling=3, display="bp", cex=1)
legend("bottomright", legend=vars, bty="n", col="gray32", pch=21, cex=1.2, pt.bg=bg)
```

```
#dev.off()

# axes 1 & 3
#png(file = 'rda_plot_SNPs_clim_Qtom107.Qchr17.Qssp3_GEA_onlyGuadalupe_PC1_PC3.png', width = 10, height = 9, res = 300, units = 'in')
plot(rda, type="n", scaling=3, xlim=c(-1,1), ylim=c(-1,1), choices=c(1,3))
points(rda, display="species", pch=21, cex=1, col="gray32", bg=col.pred, scaling=3, choices=c(1,3))
points(rda, display="species", pch=21, cex=1, col=empty.outline, bg=empty, scaling=3, choices=c(1,3))
text(rda, scaling=3, display="bp", cex=1, choices=c(1,3))
legend("bottomright", legend=vars, bty="n", col="gray32", pch=21, cex=1.2, pt.bg=bg)
```

```
#dev.off()
```

```
cand.sort <- cand[order(cand$correlation, decreasing = T),]


# colors
cols <- c('black', "#3c4a8b","#009c85","#84bc5f","#edb829","#f57404","#b30000")

# plot just one
# snp <- cand.sort$snp[1]
# cli <- cand.sort$predictor[1]
# plot(jitter(snps.sub[,snp], 0.3), clim.sub[,cli],
#      main = snp,
#      ylab = cli,
#      xlab = 'genotype',
#      bg = cols[info.sub$island],
#      col = 'black',
#      pch = 21,
#      cex = 1)


# plot more
top <- min(96, nrow(cand.sort))

#dev.new()
par(mfrow = c(4,6))
par(mar = c(3,4,2,1))

for(n in 1:top){
  
  snp <- cand.sort$snp[n]
  cli <- cand.sort$predictor[n]
  plot(jitter(snps.sub[,snp], 0.3),
       clim.sub[,cli],
       main = snp,
       ylab = cli,
       xlab = 'genotype',
       bg = cols[info.sub$island],
       col = 'black',
       pch = 21,
       cex = 1)
}
```

```
# to save

# set = 1
# for(n in 1:top){
#   
#   # start a new plot every 24
#   if(n %% 24 == 1){
#     
#     png(file = paste('RDA_SNP-climate_cors_onlyGuadalupe_set_', set, '.png', sep = ''),
#         height = 10, width = 16,
#         units = 'in',
#         res = 300)
#     
#     par(mfrow = c(4,6))
#     par(mar = c(3,4,2,1))
#   }
# 
#   
#   snp <- cand.sort$snp[n]
#   cli <- cand.sort$predictor[n]
#   plot(jitter(snps.sub[,snp], 0.3),
#        clim.sub[,cli],
#        main = snp,
#        ylab = cli,
#        xlab = 'genotype',
#        bg = cols[info.sub$island],
#        col = 'black',
#        pch = 21,
#        cex = 1)
#   
#   if(n %% 24 == 0){
#     dev.off()
#     set = set+1
#   }
#   
# }
```

```
# write.table(cand.sort, 'results/RDA_candidate_climate_SNP_table_onlyGuadalupe.csv', row.names = F, quote = F)
# save(cand.sort, file = 'results/RDA_candidate_climate_SNP_table_onlyGuadalupe.rda')
```

### 8 GEA analysis without Guadalupe, Qchr, or Anacapa

#### 8.1 RDA of SNPs and climate

```
# some of the SNP-climate associations seemed to be strongly affected by Anacapa trees, which are very closely related and at the same site, so they have the same climate variables
# version of RDA without Anacapa samples
# this is what gets used in gradient forest script

# here we'll use the subset climate variables to do a more detailed GEA

# get subset without guadalupe or anacapa
clim.sub <- clim[! info$island %in% c('Guadalupe Island', 'Mainland', 'Anacapa Island'),]
snps.sub <- snps[! info$island %in% c('Guadalupe Island', 'Mainland', 'Anacapa Island'),]
imp.sub <- imp[! info$island %in% c('Guadalupe Island', 'Mainland', 'Anacapa Island'),]
info.sub <- info[! info$island %in% c('Guadalupe Island', 'Mainland', 'Anacapa Island'),]

rda <- rda(imp.sub ~ ., data = as.data.frame(clim.sub[,vars]), scale = T)
screeplot(rda)
```

```
rda
```

```
## Call: rda(formula = imp.sub ~ bio5 + bio6 + bio15 + bio18 + bio19 +
## elev, data = as.data.frame(clim.sub[, vars]), scale = T)
## 
##                 Inertia Proportion Rank
## Total         5.689e+05  1.000e+00     
## Constrained   5.324e+04  9.359e-02    6
## Unconstrained 5.157e+05  9.064e-01   83
## Inertia is correlations 
## 
## Eigenvalues for constrained axes:
##  RDA1  RDA2  RDA3  RDA4  RDA5  RDA6 
## 12273  9745  8788  8150  7530  6759 
## 
## Eigenvalues for unconstrained axes:
##   PC1   PC2   PC3   PC4   PC5   PC6   PC7   PC8 
## 10706 10216  9769  9137  8896  8724  8687  8582 
## (Showing 8 of 83 unconstrained eigenvalues)
```

```
rda.info <- summary(rda)
head(rda.info)
```

```
## 
## Call:
## rda(formula = imp.sub ~ bio5 + bio6 + bio15 + bio18 + bio19 +      elev, data = as.data.frame(clim.sub[, vars]), scale = T) 
## 
## Partitioning of correlations:
##               Inertia Proportion
## Total          568897    1.00000
## Constrained     53245    0.09359
## Unconstrained  515652    0.90641
## 
## Eigenvalues, and their contribution to the correlations 
## 
## Importance of components:
##                            RDA1      RDA2      RDA3      RDA4      RDA5
## Eigenvalue            1.227e+04 9.745e+03 8.788e+03 8.150e+03 7.530e+03
## Proportion Explained  2.157e-02 1.713e-02 1.545e-02 1.433e-02 1.324e-02
## Cumulative Proportion 2.157e-02 3.870e-02 5.415e-02 6.848e-02 8.171e-02
##                            RDA6       PC1       PC2       PC3       PC4
## Eigenvalue            6.759e+03 1.071e+04 1.022e+04 9.769e+03 9.137e+03
## Proportion Explained  1.188e-02 1.882e-02 1.796e-02 1.717e-02 1.606e-02
## Cumulative Proportion 9.359e-02 1.124e-01 1.304e-01 1.475e-01 1.636e-01
##                             PC5       PC6       PC7       PC8       PC9
## Eigenvalue            8.896e+03 8.724e+03 8.687e+03 8.582e+03 8.444e+03
## Proportion Explained  1.564e-02 1.533e-02 1.527e-02 1.509e-02 1.484e-02
## Cumulative Proportion 1.792e-01 1.946e-01 2.098e-01 2.249e-01 2.398e-01
##                            PC10      PC11      PC12      PC13      PC14
## Eigenvalue            8.344e+03 8.165e+03 8.158e+03 8.006e+03 7.923e+03
## Proportion Explained  1.467e-02 1.435e-02 1.434e-02 1.407e-02 1.393e-02
## Cumulative Proportion 2.544e-01 2.688e-01 2.831e-01 2.972e-01 3.111e-01
##                            PC15      PC16      PC17      PC18      PC19
## Eigenvalue            7852.4856 7.524e+03 7.359e+03 7.314e+03 7.260e+03
## Proportion Explained     0.0138 1.323e-02 1.293e-02 1.286e-02 1.276e-02
## Cumulative Proportion    0.3249 3.382e-01 3.511e-01 3.639e-01 3.767e-01
##                            PC20      PC21      PC22      PC23      PC24
## Eigenvalue            7.144e+03 7.078e+03 7055.4846 6.973e+03 6.917e+03
## Proportion Explained  1.256e-02 1.244e-02    0.0124 1.226e-02 1.216e-02
## Cumulative Proportion 3.893e-01 4.017e-01    0.4141 4.264e-01 4.385e-01
##                            PC25      PC26      PC27      PC28      PC29
## Eigenvalue            6884.3751 6.795e+03 6771.1628 6.759e+03 6.692e+03
## Proportion Explained     0.0121 1.194e-02    0.0119 1.188e-02 1.176e-02
## Cumulative Proportion    0.4506 4.626e-01    0.4745 4.864e-01 4.981e-01
##                            PC30      PC31      PC32      PC33      PC34
## Eigenvalue            6.589e+03 6.563e+03 6.513e+03 6.432e+03 6.395e+03
## Proportion Explained  1.158e-02 1.154e-02 1.145e-02 1.131e-02 1.124e-02
## Cumulative Proportion 5.097e-01 5.212e-01 5.327e-01 5.440e-01 5.552e-01
##                            PC35      PC36      PC37      PC38      PC39
## Eigenvalue            6312.6245 6.249e+03 6.246e+03 6.161e+03 6.110e+03
## Proportion Explained     0.0111 1.099e-02 1.098e-02 1.083e-02 1.074e-02
## Cumulative Proportion    0.5663 5.773e-01 5.883e-01 5.991e-01 6.099e-01
##                            PC40      PC41      PC42      PC43      PC44
## Eigenvalue            6.057e+03 6.045e+03 6.002e+03 5.964e+03 5.842e+03
## Proportion Explained  1.065e-02 1.063e-02 1.055e-02 1.048e-02 1.027e-02
## Cumulative Proportion 6.205e-01 6.311e-01 6.417e-01 6.522e-01 6.624e-01
##                            PC45      PC46      PC47      PC48      PC49
## Eigenvalue            5.831e+03 5.783e+03 5.705e+03 5.576e+03 5.540e+03
## Proportion Explained  1.025e-02 1.017e-02 1.003e-02 9.801e-03 9.739e-03
## Cumulative Proportion 6.727e-01 6.829e-01 6.929e-01 7.027e-01 7.124e-01
##                            PC50      PC51      PC52      PC53      PC54
## Eigenvalue            5.494e+03 5.447e+03 5.372e+03 5.357e+03 5.331e+03
## Proportion Explained  9.657e-03 9.576e-03 9.443e-03 9.416e-03 9.370e-03
## Cumulative Proportion 7.221e-01 7.317e-01 7.411e-01 7.505e-01 7.599e-01
##                            PC55      PC56      PC57      PC58      PC59
## Eigenvalue            5.312e+03 5.280e+03 5.270e+03 5.200e+03 5.183e+03
## Proportion Explained  9.337e-03 9.281e-03 9.263e-03 9.141e-03 9.111e-03
## Cumulative Proportion 7.692e-01 7.785e-01 7.878e-01 7.969e-01 8.060e-01
##                            PC60      PC61      PC62      PC63      PC64
## Eigenvalue            5.159e+03 5.134e+03 5.129e+03 5.092e+03 5.057e+03
## Proportion Explained  9.068e-03 9.025e-03 9.016e-03 8.951e-03 8.890e-03
## Cumulative Proportion 8.151e-01 8.241e-01 8.331e-01 8.421e-01 8.510e-01
##                            PC65      PC66      PC67      PC68      PC69
## Eigenvalue            5.037e+03 5.000e+03 4.973e+03 4.952e+03 4.900e+03
## Proportion Explained  8.853e-03 8.788e-03 8.741e-03 8.705e-03 8.612e-03
## Cumulative Proportion 8.598e-01 8.686e-01 8.774e-01 8.861e-01 8.947e-01
##                            PC70      PC71      PC72      PC73      PC74
## Eigenvalue            4.853e+03 4.792e+03 4.728e+03 4.661e+03 4.621e+03
## Proportion Explained  8.531e-03 8.424e-03 8.311e-03 8.193e-03 8.123e-03
## Cumulative Proportion 9.032e-01 9.116e-01 9.199e-01 9.281e-01 9.363e-01
##                            PC75      PC76      PC77      PC78      PC79
## Eigenvalue            4.596e+03 4.578e+03 4.390e+03 4.244e+03 4.140e+03
## Proportion Explained  8.078e-03 8.047e-03 7.716e-03 7.460e-03 7.277e-03
## Cumulative Proportion 9.443e-01 9.524e-01 9.601e-01 9.676e-01 9.748e-01
##                            PC80      PC81      PC82      PC83
## Eigenvalue            4.040e+03 3.915e+03 3.841e+03 2.522e+03
## Proportion Explained  7.102e-03 6.882e-03 6.751e-03 4.434e-03
## Cumulative Proportion 9.819e-01 9.888e-01 9.956e-01 1.000e+00
## 
## Accumulated constrained eigenvalues
## Importance of components:
##                            RDA1      RDA2      RDA3      RDA4      RDA5
## Eigenvalue            1.227e+04 9744.8509 8788.2009 8149.9468 7530.4254
## Proportion Explained  2.305e-01    0.1830    0.1651    0.1531    0.1414
## Cumulative Proportion 2.305e-01    0.4135    0.5786    0.7316    0.8731
##                            RDA6
## Eigenvalue            6758.7848
## Proportion Explained     0.1269
## Cumulative Proportion    1.0000
## 
## Scaling 2 for species and site scores
## * Species are scaled proportional to eigenvalues
## * Sites are unscaled: weighted dispersion equal on all dimensions
## * General scaling constant of scores:  84.35404 
## 
## 
## Species scores
## 
##                    RDA1       RDA2      RDA3       RDA4      RDA5     RDA6
## 1_39789_A_G_G -0.007697 -0.0559688 -0.016291 -0.0230726  0.040357  0.00359
## 1_52176_G_A_A  0.026027 -0.0144205  0.005447  0.0072781  0.007276  0.01406
## 1_52195_G_A_A -0.008104  0.0002707  0.008469 -0.0005330 -0.002084 -0.00491
## 1_52213_C_T_T -0.002082  0.0065792 -0.023509  0.0007379 -0.023285  0.01735
## 1_52248_T_C_T -0.005708 -0.0052330  0.014730 -0.0056368  0.030397  0.03134
## 1_52331_G_T_T -0.020263  0.0076378 -0.002008  0.0143036 -0.003234  0.01111
## ....                                                                      
## 
## 
## Site scores (weighted sums of species scores)
## 
##                  RDA1    RDA2   RDA3    RDA4     RDA5    RDA6
## Qchr.A.LA.215  -6.537  -6.808 -3.068   4.107   0.9685  -2.355
## Qchr.A.LA.222 -11.352  -7.679 -3.086   3.225   1.1973  -3.335
## Qchr.I.LA.20   -7.795  -5.428  2.069  -3.170 -21.3776  -8.264
## Qchr.I.LA.25   -7.581 -10.205  3.595   3.415 -10.1886 -22.172
## Qchr.I.SB.02   -8.761  28.451 18.771 -15.028  -1.0559  -3.317
## Qchr.I.SB.03   -9.257  29.324 19.563 -14.933  -0.7998  -3.055
## ....                                                         
## 
## 
## Site constraints (linear combinations of constraining variables)
## 
##                 RDA1    RDA2   RDA3    RDA4     RDA5    RDA6
## Qchr.A.LA.215 -8.120  -6.650 -3.614   2.693   0.7567  -2.289
## Qchr.A.LA.222 -8.120  -6.650 -3.614   2.693   0.7567  -2.289
## Qchr.I.LA.20  -7.372  -6.183  0.380  -2.281 -20.7349  -7.121
## Qchr.I.LA.25  -7.416 -12.043  1.840   4.475 -10.4983 -20.688
## Qchr.I.SB.02  -5.519  26.811 17.498 -10.851  -0.4440  -1.857
## Qchr.I.SB.03  -5.519  26.811 17.498 -10.851  -0.4440  -1.857
## ....                                                        
## 
## 
## Biplot scores for constraining variables
## 
##          RDA1      RDA2     RDA3     RDA4    RDA5    RDA6
## bio5  -0.9685  0.006987 -0.06329 -0.09913  0.1734  0.1345
## bio6  -0.7264 -0.633823 -0.02849 -0.03410 -0.1981  0.1716
## bio15 -0.4490  0.350588 -0.63984  0.15267 -0.2765  0.4079
## bio18  0.9132 -0.088787  0.07242 -0.24305  0.2566 -0.1677
## bio19  0.3571  0.557238 -0.47380  0.37428  0.4057 -0.1811
## elev  -0.3535  0.421570  0.24546 -0.12583  0.2741 -0.7390
```

```
RsquareAdj(rda)
```

```
## $r.squared
## [1] 0.09359315
## 
## $adj.r.squared
## [1] 0.02806976
```

```
# $r.squared
# [1] 0.09138727
# $adj.r.squared
# [1] 0.0294364

# colors
cols <- c('black', "#3c4a8b","#009c85","#84bc5f","#edb829","#f57404","#b30000")
bg <- c('black', "#3c4a8b","#009c85","#84bc5f","#edb829","#f57404","#b30000")

plot_clim(rda, info = info.sub, save = F, file_string = 'Qtom107.Qchr17.Qssp3_GEA_noQchrGuadalupeAnacapa')
```

```
plot_clim(rda, info = info.sub, choices = c(1,3), save = F, file_string = 'Qtom107.Qchr17.Qssp3_GEA_noQchrGuadalupeAnacapa')
```

```
plot(rda, type = 'none')
points(rda, display = 'sites', pch = sp[info.sub$sp], bg = bg[info.sub$island], col = 'black', cex = 1.5)
text(rda, display = 'bp')
points(rda, display = 'species', pch = 21, cex = 1, col = 'gray32')
```

#### 8.2 identify candidate SNPs from RDA outliers

```
# again basically following Forester et al

# most variation explained by first three axes (~5% out of 9% unadjusted)
screeplot(rda)
```

```
summary(rda)$cont
```

```
## $importance
## Importance of components:
##                            RDA1      RDA2      RDA3      RDA4      RDA5
## Eigenvalue            1.227e+04 9.745e+03 8.788e+03 8.150e+03 7.530e+03
## Proportion Explained  2.157e-02 1.713e-02 1.545e-02 1.433e-02 1.324e-02
## Cumulative Proportion 2.157e-02 3.870e-02 5.415e-02 6.848e-02 8.171e-02
##                            RDA6       PC1       PC2       PC3       PC4
## Eigenvalue            6.759e+03 1.071e+04 1.022e+04 9.769e+03 9.137e+03
## Proportion Explained  1.188e-02 1.882e-02 1.796e-02 1.717e-02 1.606e-02
## Cumulative Proportion 9.359e-02 1.124e-01 1.304e-01 1.475e-01 1.636e-01
##                             PC5       PC6       PC7       PC8       PC9
## Eigenvalue            8.896e+03 8.724e+03 8.687e+03 8.582e+03 8.444e+03
## Proportion Explained  1.564e-02 1.533e-02 1.527e-02 1.509e-02 1.484e-02
## Cumulative Proportion 1.792e-01 1.946e-01 2.098e-01 2.249e-01 2.398e-01
##                            PC10      PC11      PC12      PC13      PC14
## Eigenvalue            8.344e+03 8.165e+03 8.158e+03 8.006e+03 7.923e+03
## Proportion Explained  1.467e-02 1.435e-02 1.434e-02 1.407e-02 1.393e-02
## Cumulative Proportion 2.544e-01 2.688e-01 2.831e-01 2.972e-01 3.111e-01
##                            PC15      PC16      PC17      PC18      PC19
## Eigenvalue            7852.4856 7.524e+03 7.359e+03 7.314e+03 7.260e+03
## Proportion Explained     0.0138 1.323e-02 1.293e-02 1.286e-02 1.276e-02
## Cumulative Proportion    0.3249 3.382e-01 3.511e-01 3.639e-01 3.767e-01
##                            PC20      PC21      PC22      PC23      PC24
## Eigenvalue            7.144e+03 7.078e+03 7055.4846 6.973e+03 6.917e+03
## Proportion Explained  1.256e-02 1.244e-02    0.0124 1.226e-02 1.216e-02
## Cumulative Proportion 3.893e-01 4.017e-01    0.4141 4.264e-01 4.385e-01
##                            PC25      PC26      PC27      PC28      PC29
## Eigenvalue            6884.3751 6.795e+03 6771.1628 6.759e+03 6.692e+03
## Proportion Explained     0.0121 1.194e-02    0.0119 1.188e-02 1.176e-02
## Cumulative Proportion    0.4506 4.626e-01    0.4745 4.864e-01 4.981e-01
##                            PC30      PC31      PC32      PC33      PC34
## Eigenvalue            6.589e+03 6.563e+03 6.513e+03 6.432e+03 6.395e+03
## Proportion Explained  1.158e-02 1.154e-02 1.145e-02 1.131e-02 1.124e-02
## Cumulative Proportion 5.097e-01 5.212e-01 5.327e-01 5.440e-01 5.552e-01
##                            PC35      PC36      PC37      PC38      PC39
## Eigenvalue            6312.6245 6.249e+03 6.246e+03 6.161e+03 6.110e+03
## Proportion Explained     0.0111 1.099e-02 1.098e-02 1.083e-02 1.074e-02
## Cumulative Proportion    0.5663 5.773e-01 5.883e-01 5.991e-01 6.099e-01
##                            PC40      PC41      PC42      PC43      PC44
## Eigenvalue            6.057e+03 6.045e+03 6.002e+03 5.964e+03 5.842e+03
## Proportion Explained  1.065e-02 1.063e-02 1.055e-02 1.048e-02 1.027e-02
## Cumulative Proportion 6.205e-01 6.311e-01 6.417e-01 6.522e-01 6.624e-01
##                            PC45      PC46      PC47      PC48      PC49
## Eigenvalue            5.831e+03 5.783e+03 5.705e+03 5.576e+03 5.540e+03
## Proportion Explained  1.025e-02 1.017e-02 1.003e-02 9.801e-03 9.739e-03
## Cumulative Proportion 6.727e-01 6.829e-01 6.929e-01 7.027e-01 7.124e-01
##                            PC50      PC51      PC52      PC53      PC54
## Eigenvalue            5.494e+03 5.447e+03 5.372e+03 5.357e+03 5.331e+03
## Proportion Explained  9.657e-03 9.576e-03 9.443e-03 9.416e-03 9.370e-03
## Cumulative Proportion 7.221e-01 7.317e-01 7.411e-01 7.505e-01 7.599e-01
##                            PC55      PC56      PC57      PC58      PC59
## Eigenvalue            5.312e+03 5.280e+03 5.270e+03 5.200e+03 5.183e+03
## Proportion Explained  9.337e-03 9.281e-03 9.263e-03 9.141e-03 9.111e-03
## Cumulative Proportion 7.692e-01 7.785e-01 7.878e-01 7.969e-01 8.060e-01
##                            PC60      PC61      PC62      PC63      PC64
## Eigenvalue            5.159e+03 5.134e+03 5.129e+03 5.092e+03 5.057e+03
## Proportion Explained  9.068e-03 9.025e-03 9.016e-03 8.951e-03 8.890e-03
## Cumulative Proportion 8.151e-01 8.241e-01 8.331e-01 8.421e-01 8.510e-01
##                            PC65      PC66      PC67      PC68      PC69
## Eigenvalue            5.037e+03 5.000e+03 4.973e+03 4.952e+03 4.900e+03
## Proportion Explained  8.853e-03 8.788e-03 8.741e-03 8.705e-03 8.612e-03
## Cumulative Proportion 8.598e-01 8.686e-01 8.774e-01 8.861e-01 8.947e-01
##                            PC70      PC71      PC72      PC73      PC74
## Eigenvalue            4.853e+03 4.792e+03 4.728e+03 4.661e+03 4.621e+03
## Proportion Explained  8.531e-03 8.424e-03 8.311e-03 8.193e-03 8.123e-03
## Cumulative Proportion 9.032e-01 9.116e-01 9.199e-01 9.281e-01 9.363e-01
##                            PC75      PC76      PC77      PC78      PC79
## Eigenvalue            4.596e+03 4.578e+03 4.390e+03 4.244e+03 4.140e+03
## Proportion Explained  8.078e-03 8.047e-03 7.716e-03 7.460e-03 7.277e-03
## Cumulative Proportion 9.443e-01 9.524e-01 9.601e-01 9.676e-01 9.748e-01
##                            PC80      PC81      PC82      PC83
## Eigenvalue            4.040e+03 3.915e+03 3.841e+03 2.522e+03
## Proportion Explained  7.102e-03 6.882e-03 6.751e-03 4.434e-03
## Cumulative Proportion 9.819e-01 9.888e-01 9.956e-01 1.000e+00
```

```
# get SNP loadings for first 3 axes
load.rda <- scores(rda, choices=c(1:3), display="species")

hist(load.rda[,1], main="Loadings on RDA1")
```

```
hist(load.rda[,2], main="Loadings on RDA2")
```

```
hist(load.rda[,3], main="Loadings on RDA3")
```

```
outliers <- function(x,z){
  
  lims <- mean(x) + c(-1, 1) * z * sd(x) # find loadings +/-z sd from mean loading  
  x[x < lims[1] | x > lims[2]] # locus names in these tails
  
}

# start out with a cutoff of 4 std dev because there are a lot of snps
cand1 <- outliers(load.rda[,1],4)
cand2 <- outliers(load.rda[,2],4)
cand3 <- outliers(load.rda[,3],4)

length(cand1)
```

```
## [1] 168
```

```
length(cand2)
```

```
## [1] 178
```

```
length(cand3)
```

```
## [1] 214
```

```
ncand <- length(cand1) + length(cand2) + length(cand3)
ncand
```

```
## [1] 560
```

```
# set up results

cand1 <- cbind.data.frame(rep(1,times=length(cand1)), names(cand1), unname(cand1))
cand2 <- cbind.data.frame(rep(2,times=length(cand2)), names(cand2), unname(cand2))
cand3 <- cbind.data.frame(rep(3,times=length(cand3)), names(cand3), unname(cand3))

colnames(cand1) <- colnames(cand2) <- colnames(cand3) <- c("axis","snp","loading")

cand <- rbind(cand1, cand2, cand3)
cand$snp <- as.character(cand$snp)

# get correlations of SNPs with climate
tmp <- matrix(nrow=(ncand), ncol=ncol(clim.sub[,vars]))

colnames(tmp) <- colnames(clim.sub[,vars])

for (i in 1:length(cand$snp)) {
  
  nam <- cand$snp[i] # loop through candidate snp names
  snp.gen <- imp.sub[,nam]
  tmp[i,] <- apply(clim.sub[,vars], 2, function(x) cor(x,snp.gen))
}

cand <- cbind.data.frame(cand, tmp)
head(cand)
```

```
##   axis              snp     loading       bio5       bio6      bio15      bio18
## 1    1   1_388897_C_T_T  0.08359348 -0.7282967 -0.4178589 -0.6847897  0.7597653
## 2    1  1_2592491_G_T_T  0.08932577 -0.7503434 -0.6247846 -0.6291060  0.8223044
## 3    1 1_10042008_T_A_A  0.06895150 -0.6272079 -0.3843603 -0.3165384  0.5508314
## 4    1 1_16403258_G_A_G -0.06771754  0.6123452  0.4337874  0.4145637 -0.5679593
## 5    1 1_16816786_A_G_G  0.06351980 -0.5489538 -0.3426665 -0.5672601  0.5941813
## 6    1 1_23151400_A_G_G  0.06297299 -0.5390203 -0.4242257 -0.4432135  0.5983690
##          bio19        elev
## 1  0.026539728 -0.24822235
## 2  0.257700822 -0.13549685
## 3  0.144876303 -0.23436475
## 4 -0.008567605  0.15750430
## 5  0.019676051 -0.14775036
## 6  0.107858198 -0.06505125
```

```
#  check for duplicate snps - outliers on multiple axes
length(cand$snp[duplicated(cand$snp)]) # 0 - so skip below
```

```
## [1] 0
```

```
# foo <- cbind(cand$axis, duplicated(cand$snp)) 
# table(foo[foo[,1]==1,2]) # no duplicates on axis 1
# table(foo[foo[,1]==2,2]) #  6 duplicates on axis 2
# table(foo[foo[,1]==3,2]) # no duplicates on axis 3
# 
# cand <- cand[!duplicated(cand$snp),] # remove duplicate detections

dim(cand) # 560 SNPs
```

```
## [1] 560   9
```

```
# get best climate correlation for each snp
# note: need to modify the columns if different climate variables are used

for (i in 1:length(cand$snp)) {
  
  row <- cand[i,]
  cand[i, 10] <- names(which.max(abs(row[vars]))) # gives the variable
  cand[i, 11] <- max(abs(row[vars]))              # gives the correlation
}

colnames(cand)[10] <- "predictor"
colnames(cand)[11] <- "correlation"

table(cand$predictor)
```

```
## 
## bio15 bio18 bio19  bio5  bio6  elev 
##    47    73   174    88    23   155
```

```
# bio15 bio18 bio19  bio5  bio6  elev 
#    47    73   174    88    23   155

########
# plot!

# colors
sel <- cand$snp
env <- cand$predictor

colset <- c("#d64b15","#013479","#ffb90f","#22a27c","#b1e3ad","#7d9ceb")
env[env=="bio5"] <- colset[1] # Max Temperature of Warmest Month
env[env=="bio6"] <- colset[2] # Min Temperature of Coldest Month
env[env=="bio15"] <- colset[3] # Precipitation Seasonality (Coefficient of Variation)
env[env=="bio18"] <- colset[4] # Precipitation of Warmest Quarter
env[env=="bio19"] <- colset[5] # Precipitation of Coldest Quarter
env[env=="elev"] <- colset[6]

# color by predictor:
col.pred <- rownames(rda$CCA$v) # pull the SNP names

for (i in 1:length(sel)) {           # color code candidate SNPs
  foo <- match(sel[i],col.pred)
  col.pred[foo] <- env[i]
}

# set color for non-candidate snps
# change if they don't already have a color
col.pred[! col.pred %in% colset] <- '#f1eef6' # non-candidate SNPs
empty <- col.pred
empty[grep("#f1eef6",empty)] <- rgb(0,1,0, alpha=0) # transparent
empty.outline <- ifelse(empty=="#00FF0000","#00FF0000","gray32")
bg <- colset

# now plot

# axes 1 & 2
#png(file = 'rda_plot_SNPs_clim_Qtom107.Qchr17.Qssp3_GEA_noQchrGuadalupeAnacapa_PC1_PC2.png', width = 10, height = 9, res = 300, units = 'in')
plot(rda, type="n", scaling=3, xlim=c(-1,1), ylim=c(-1,1))
points(rda, display="species", pch=21, cex=1, col="gray32", bg=col.pred, scaling=3)
points(rda, display="species", pch=21, cex=1, col=empty.outline, bg=empty, scaling=3)
text(rda, scaling=3, display="bp", cex=1)
legend("bottomright", legend=vars, bty="n", col="gray32", pch=21, cex=1.2, pt.bg=bg)
```

```
#dev.off()

# axes 1 & 3
#png(file = 'rda_plot_SNPs_clim_Qtom107.Qchr17.Qssp3_GEA_noQchrGuadalupeAnacapa_PC1_PC3.png', width = 10, height = 9, res = 300, units = 'in')
plot(rda, type="n", scaling=3, xlim=c(-1,1), ylim=c(-1,1), choices=c(1,3))
points(rda, display="species", pch=21, cex=1, col="gray32", bg=col.pred, scaling=3, choices=c(1,3))
points(rda, display="species", pch=21, cex=1, col=empty.outline, bg=empty, scaling=3, choices=c(1,3))
text(rda, scaling=3, display="bp", cex=1, choices=c(1,3))
legend("bottomright", legend=vars, bty="n", col="gray32", pch=21, cex=1.2, pt.bg=bg)
```

```
#dev.off()


# save

# write.table(cand, 'results/RDA_candidate_climate_SNP_table_noGuadAna.csv', row.names = F, quote = F)
# save(cand, file = 'results/RDA_candidate_climate_SNP_table_noGuadAna.rda')
```

```
cand.sort <- cand[order(cand$correlation, decreasing = T),]


# colors
cols <- c('black', "#3c4a8b","#009c85","#84bc5f","#edb829","#f57404","#b30000")

# plot just one
# snp <- cand.sort$snp[1]
# cli <- cand.sort$predictor[1]
# plot(jitter(snps.sub[,snp], 0.3), clim.sub[,cli],
#      main = snp,
#      ylab = cli,
#      xlab = 'genotype',
#      bg = cols[info.sub$island],
#      col = 'black',
#      pch = 21,
#      cex = 1)


# plot more
top <- min(96, nrow(cand.sort))

#dev.new()
par(mfrow = c(4,6))
par(mar = c(3,4,2,1))

for(n in 1:top){
  
  snp <- cand.sort$snp[n]
  cli <- cand.sort$predictor[n]
  plot(jitter(snps.sub[,snp], 0.3),
       clim.sub[,cli],
       main = snp,
       ylab = cli,
       xlab = 'genotype',
       bg = cols[info.sub$island],
       col = 'black',
       pch = 21,
       cex = 1.2)
}
```

```
# to save

# set = 1
# for(n in 1:top){
#   
#   # start a new plot every 24
#   if(n %% 24 == 1){
#     
#     png(file = paste('RDA_SNP-climate_cors_set_noGuadAna', set, '.png', sep = ''),
#         height = 10, width = 16,
#         units = 'in',
#         res = 300)
#     
#     par(mfrow = c(4,6))
#     par(mar = c(3,4,2,1))
#   }
# 
#   
#   snp <- cand.sort$snp[n]
#   cli <- cand.sort$predictor[n]
#   plot(jitter(snps.sub[,snp], 0.3),
#        clim.sub[,cli],
#        main = snp,
#        ylab = cli,
#        xlab = 'genotype',
#        bg = cols[info.sub$island],
#        col = 'black',
#        pch = 21,
#        cex = 1)
#   
#   if(n %% 24 == 0){
#     dev.off()
#     set = set+1
#   }
#   
# }
```
