## Supplementary Rmarkdown Outputs for "Comparison of conservation strategies for California Channel Island Oak (*Quercus tomentella*) using climate suitability predicted from genomic data": simple_pca_with_vegan.html

PCA of SNP data


### PCA of SNP data

###### Alayna Mead

#### 2024-07-16

- 1 Setup
  - 1.1 packages and data
  - 1.2 setup SNP data
  - 1.3 setup collection data
- 2 impute SNP data
- 3 Plots
  - 3.1 Simple PCA, all samples
  - 3.2 Only Channel Islands, exclude
    Guadalupe Island
  - 3.3 Only Northern Channel
    Islands

Plot PCAs from processed SNP data using vegan

### 1 Setup

#### 1.1 packages and data

```
library('vegan') # redundancy analysis
```

```
## Loading required package: permute
```

```
## Loading required package: lattice
```

```
## This is vegan 2.6-6.1
```

```
library('adegenet') # read.PLINK()
```

```
## Loading required package: ade4
```

```
## 
##    /// adegenet 2.1.10 is loaded ////////////
## 
##    > overview: '?adegenet'
##    > tutorials/doc/questions: 'adegenetWeb()' 
##    > bug reports/feature requests: adegenetIssues()
```

```
library('psych') # pairs.panels()

# Following:
#https://popgen.nescent.org/2018-03-27_RDA_GEA.html

# site data - named dat, needs to be renamed
load('data/clean/CCGP_samples_channel_islands_clean.rda')
info <- dat

# use this to read SNPs!
dat <- read.PLINK('data/raw/Qtom107.Qchr17.Qssp3.20220906.qlob.ef.repeatsOut.renamedChrsVars.biallelicSNPs.meanDP5.genoDP5.MAF0.01.missing0.9.ldPruned.additive.raw', 
                  n.cores = 2, 
                  quiet = F)
```

```
## 
##  Reading PLINK raw format into a genlight object... 
## 
## 
##  Reading loci information... 
## 
##  Reading and converting genotypes... 
## .
##  Building final object... 
## 
## ...done.
```

```
sessionInfo()
```

```
## R version 4.4.0 (2024-04-24)
## Platform: x86_64-pc-linux-gnu
## Running under: Arch Linux
## 
## Matrix products: default
## BLAS:   /usr/lib/libblas.so.3.12.0 
## LAPACK: /usr/lib/liblapack.so.3.12.0
## 
## locale:
##  [1] LC_CTYPE=en_US.UTF-8       LC_NUMERIC=C              
##  [3] LC_TIME=en_US.UTF-8        LC_COLLATE=en_US.UTF-8    
##  [5] LC_MONETARY=en_US.UTF-8    LC_MESSAGES=en_US.UTF-8   
##  [7] LC_PAPER=en_US.UTF-8       LC_NAME=C                 
##  [9] LC_ADDRESS=C               LC_TELEPHONE=C            
## [11] LC_MEASUREMENT=en_US.UTF-8 LC_IDENTIFICATION=C       
## 
## time zone: America/New_York
## tzcode source: system (glibc)
## 
## attached base packages:
## [1] stats     graphics  grDevices utils     datasets  methods   base     
## 
## other attached packages:
## [1] psych_2.4.6.26  adegenet_2.1.10 ade4_1.7-22     vegan_2.6-6.1  
## [5] lattice_0.22-6  permute_0.9-7  
## 
## loaded via a namespace (and not attached):
##  [1] sass_0.4.9        utf8_1.2.4        generics_0.1.3    stringi_1.8.4    
##  [5] digest_0.6.35     magrittr_2.0.3    evaluate_0.23     grid_4.4.0       
##  [9] fastmap_1.2.0     seqinr_4.2-36     plyr_1.8.9        jsonlite_1.8.8   
## [13] Matrix_1.7-0      ape_5.8           promises_1.3.0    mgcv_1.9-1       
## [17] fansi_1.0.6       scales_1.3.0      jquerylib_0.1.4   mnormt_2.1.1     
## [21] cli_3.6.2         shiny_1.8.1.1     rlang_1.1.3       munsell_0.5.1    
## [25] splines_4.4.0     cachem_1.1.0      yaml_2.3.8        tools_4.4.0      
## [29] parallel_4.4.0    reshape2_1.4.4    dplyr_1.1.4       colorspace_2.1-0 
## [33] ggplot2_3.5.1     httpuv_1.6.15     mime_0.12         vctrs_0.6.5      
## [37] R6_2.5.1          lifecycle_1.0.4   stringr_1.5.1     MASS_7.3-60.2    
## [41] cluster_2.1.6     pkgconfig_2.0.3   later_1.3.2       pillar_1.9.0     
## [45] bslib_0.7.0       gtable_0.3.5      glue_1.7.0        Rcpp_1.0.12      
## [49] xfun_0.44         tibble_3.2.1      tidyselect_1.2.1  knitr_1.46       
## [53] xtable_1.8-4      htmltools_0.5.8.1 nlme_3.1-164      igraph_2.0.3     
## [57] rmarkdown_2.27    compiler_4.4.0
```

```
knitr::opts_chunk$set(fig.width = 10, fig.height = 8)
knitr::opts_chunk$set(tidy.opts = list(width.cutoff = 80), tidy = TRUE)
```

#### 1.2 setup SNP data

```
# look at dataset
# it omits the non-SNP info
dat
```

```
##  /// GENLIGHT OBJECT /////////
## 
##  // 127 genotypes,  585,298 binary SNPs, size: 80.1 Mb
##  3736056 (5.03 %) missing data
## 
##  // Basic content
##    @gen: list of 127 SNPbin
##    @ploidy: ploidy of each individual  (range: 2-2)
## 
##  // Optional content
##    @ind.names:  127 individual labels
##    @loc.names:  585298 locus labels
##    @pop: population of each individual (group size range: 1-1)
##    @other: a list containing: sex  phenotype  pat  mat
```

### 3 Plots

#### 3.1 Simple PCA, all samples

Uses vegan rda() function without constraining matrices, which just
runs a PCA

From documentation: “If both matrices Z and Y are missing, the data
matrix is analysed by ordinary correspondence analysis (or principal
components analysis).”

```
# run the redundancy analysis

rda <- rda(imp, scale = T)

rda
```

```
## Call: rda(X = imp, scale = T)
## 
##               Inertia Rank
## Total          585298     
## Unconstrained  585298  126
## Inertia is correlations 
## 
## Eigenvalues for unconstrained axes:
##   PC1   PC2   PC3   PC4   PC5   PC6   PC7   PC8 
## 16078 12277  8152  7876  7338  7186  6959  6772 
## (Showing 8 of 126 unconstrained eigenvalues)
```

```
#summary(rda)
screeplot(rda)
```

```
# save summary
rda.info <- summary(rda)
rda.info$sites
```

```
##                          PC1         PC2           PC3           PC4
## Qchr.A.232       -4.66747445  16.5120682 -19.769688499   1.584348077
## Qchr.A.275       -5.02584116  16.1452589 -18.288636252   2.924312699
## Qchr.A.ED.102    -6.21776673  19.5779428 -16.330827664   3.500171485
## Qchr.A.ELD.53    -6.39686792  20.5905326 -20.365037230   3.448140260
## Qchr.A.KER.16    -4.95166217  14.0570713  -5.342612012   2.880341830
## Qchr.A.LA.02     -6.54332191  17.9849823   0.037087809   0.981202011
## Qchr.A.LA.215     1.82075226   3.7178319   4.374346229   4.903207108
## Qchr.A.LA.222    -1.31903872  10.1899834   3.283659894   3.989005661
## Qchr.A.SIS.309   -5.40741923  17.7251224 -20.794527451   2.814798517
## Qchr.A.PLU.65    -6.11305978  20.7349762 -20.499394363   1.781739421
## Qchr.A.SBD.83    -5.61238365  14.9507283   0.694992004   3.160263241
## Qchr.I.LA.20      0.09600950   5.9551220  -3.976843165   0.504950829
## Qchr.I.LA.25      0.95322607   4.2942300  -0.234010732   1.389324907
## Qchr.I.SB.02     -1.21347198   9.9630605  -9.728397154 -17.682806992
## Qchr.I.SB.03     -1.69680978  11.7438234 -11.284375354 -18.819387289
## Qchr.JJK.SB.13   -1.99251930  11.9901365  -5.450870245  -9.661426863
## Qchr.JR.SB.01     1.67531055   4.3535753  -1.143768700  -7.303276955
## Qspp.I.LA.03     -1.46272453   8.9056059   1.579173774  -5.383552061
## Qspp.I.LA.04      3.73129680   0.5536767   0.090101840  -0.218722846
## Qspp.I.LA.14      1.69356546   3.9194549  -3.928246107   0.133441913
## Qtom.A.LA.169     1.48942581   3.7176936   1.994762139   1.534395840
## Qtom.A.LA.175     3.84275710   1.4456430   5.570157410  -0.441812666
## Qtom.A.LA.176     1.72137798   4.4889043   5.871233737  -0.153831947
## Qtom.A.LA.180     0.70223883   4.9673715   1.125962787   3.329186575
## Qtom.A.LA.181    -4.44640960  14.4345674   2.875792677   0.645400248
## Qtom.A.LA.183    -1.25333222  11.5242761   9.770392343  15.478801200
## Qtom.A.LA.190    -1.31043789   9.9218268  14.361679949   8.606385508
## Qtom.A.LA.191    -1.63923794  13.0241405  10.687452311  15.540396790
## Qtom.A.LA.198    -0.68996466  12.1276340  22.414051931   8.062096550
## Qtom.A.LA.200     3.48630146   4.7010808  18.703048943   3.600456437
## Qtom.A.LA.204     0.44605682  10.9876733  29.430769417  11.166071051
## Qtom.A.LA.207    -0.44506613  12.0703790  29.291487758  10.632135210
## Qtom.A.LA.208     3.67248589   2.1372479   8.363020862  -2.967916760
## Qtom.A.LA.209    -0.08420729  10.5105592  24.257559176   8.894913325
## Qtom.A.LA.218    -1.31006265  10.5317068   3.548437524   3.985361404
## Qtom.A.LA.221     2.54730953   4.4635041  14.631536345   4.452060398
## Qtom.A.SB.324     5.59805154  -4.8902100  -3.393589131   9.418773001
## Qtom.A.SB.325     7.33986506  -5.8132885  -5.976968610  12.165398118
## Qtom.A.SB.327     7.66610758  -6.1712935  -5.884494709  11.561637370
## Qtom.A.SB.329     6.55699527  -5.4501579  -4.418319823   8.843006023
## Qtom.A.SB.330     6.87220862  -5.5410242  -4.030798604   8.279298354
## Qtom.A.SB.331     6.22394616  -5.2761344  -4.635288751  10.163671334
## Qtom.A.SB.332     7.81047946  -6.2550803  -6.318004324  11.946038646
## Qtom.A.SB.333     7.18222098  -5.7745505  -5.951252634  11.305570431
## Qtom.A.SB.335     7.72766331  -6.5124070  -4.749100972   9.086137964
## Qtom.A.SB.336     7.78251214  -6.7952212  -6.160669553  10.167087306
## Qtom.A.SB.337     8.43873821  -7.4402296  -5.481199615   9.946622214
## Qtom.A.SB.338     7.63158421  -6.2735701  -5.894764393  10.647485643
## Qtom.A.SB.339     8.22188164  -6.9149462  -5.705017436   9.840648648
## Qtom.A.SB.341     5.61716505  -5.0312437  -3.360481817   8.144005916
## Qtom.A.SB.342     7.10459585  -5.2506317  -4.472273819   8.077153343
## Qtom.A.SB.343     7.24092703  -5.7503487  -4.712552770   6.918841958
## Qtom.A.SB.344     6.87887236  -5.6559022  -4.103253582   7.447004094
## Qtom.A.SB.345     5.72368993  -4.7510108  -3.078921516   7.276513885
## Qtom.A.SB.346     6.73405834  -5.2579148  -4.101346913   8.760005116
## Qtom.A.SB.347     7.68704404  -6.1385962  -3.263596820   6.512552436
## Qtom.A.SB.348     7.09140727  -5.4350450  -4.557855553   7.442481440
## Qtom.A.SB.352     8.42956602  -8.0245850  -8.165211056  15.912396149
## Qtom.A.SB.353     8.45055568  -8.0769904  -8.162133968  15.739414918
## Qtom.A.SB.355     8.23812245  -7.4616493  -6.604327647  11.773422426
## Qtom.I.BC.10    -14.44608384  -9.7689407  -0.535796325   2.489278926
## Qtom.I.BC.11    -18.49934598 -12.8991779  -0.290784400  -0.597321720
## Qtom.I.BC.12    -17.48610753 -11.1646488  -0.799528214   1.310216311
## Qtom.I.BC.13    -16.46712650 -11.5021281  -0.378576423   0.851472530
## Qtom.I.BC.18    -15.59533123 -10.1810858  -0.798386225   1.710637751
## Qtom.I.BC.20    -11.60303056  -1.4335197   1.032930819  -1.467319730
## Qtom.I.BC.21    -14.74056395  -3.8856760   2.579330681  -3.599691319
## Qtom.I.BC.23    -17.31761158  -9.2006766   2.730051059  -2.713286810
## Qtom.I.BC.24    -20.45871684 -12.5593023   1.604661664  -2.610578245
## Qtom.I.BC.32    -16.44539907 -10.4918771  -0.482276240   3.340725541
## Qtom.I.BC.33    -14.84364558  -9.5204096   0.143499533   2.855719222
## Qtom.I.BC.35    -17.16671341 -11.3070457   0.446899244   1.885045436
## Qtom.I.BC.39    -12.29087585  -7.6177409   1.995592701   0.821973557
## Qtom.I.BC.4     -16.39924770 -11.3846567   0.726654482   1.948709020
## Qtom.I.BC.40    -17.46676898 -12.2474581   0.593006399   2.340593649
## Qtom.I.BC.41    -14.69256831  -9.3014144  -0.413127652   3.432307486
## Qtom.I.BC.42    -10.08980300   0.4407192   0.695425934  -1.014763525
## Qtom.I.BC.43    -17.76420608  -4.1176996   3.265675510  -9.320960825
## Qtom.I.BC.44    -16.08695342  -2.2047375   3.076709022  -7.918351420
## Qtom.I.BC.45    -15.24436515   0.6299695   0.108426676  -4.920355838
## Qtom.I.BC.46    -15.32306225  -8.5452281  -0.005454515   2.269873380
## Qtom.I.BC.48    -14.63946004  -8.1578039  -0.228030777   2.578634824
## Qtom.I.LA.SC.01  -3.84290913  13.6740562   2.370594029  -4.366036524
## Qtom.I.LA.02      2.06053370   3.0935284   5.638786382  -0.342936425
## Qtom.I.LA.05     -0.93211441   7.7416120  -0.911778959  -2.867243679
## Qtom.I.LA.06     -1.45120156  10.7281497   1.675094267  -5.842145184
## Qtom.I.LA.08      4.31833412   0.4517174   2.955399051  -4.413056270
## Qtom.I.LA.10      3.34700371  -1.3773730   0.321964110   1.512320420
## Qtom.I.LA.11      5.64552868  -2.8575536   0.182809911   0.442409349
## Qtom.I.LA.12      3.76987447   0.8860187   0.113676687   0.201610013
## Qtom.I.LA.13      0.84510886   4.9384186   0.365838742   1.595495520
## Qtom.I.LA.21      5.85253852  -3.4969773  -0.288915826   2.494378518
## Qtom.I.LA.22     -0.62706135   5.2369242  -0.298400624   0.210823548
## Qtom.I.LA.23      4.59128226  -0.9805607   0.734290046   0.010408826
## Qtom.I.LA.24      4.11163476  -0.1297478   0.490860691  -0.006346872
## Qtom.I.SB.01      6.37563409  -2.5341654  -5.942118779 -15.747810091
## Qtom.I.SB.04     -1.33915023  10.5438421 -11.101567377 -18.357101871
## Qtom.I.SB.09      6.34463127  -5.1385635   2.762500186  -8.924808842
## Qtom.I.SB.10      8.63816142  -5.0710245   3.617128039 -19.138503418
## Qtom.I.SB.11      7.70188972  -5.0539783   5.545554429 -14.138860209
## Qtom.I.SB.12      7.76505717  -4.9602816   5.672984928 -15.098853389
## Qtom.I.SB.28      5.18188033  -1.3895305  -0.620396330  -8.823755437
## Qtom.I.SB.33      5.86191907  -2.7996224  -2.280408746  -6.180672774
## Qtom.I.SB.35      5.78302708  -4.2236088  -5.225287058  -4.441315218
## Qtom.I.SB.36      6.26453209  -3.0213157  -0.992305089  -7.334776284
## Qtom.I.SB.37      6.18681673  -3.6606168  -3.961749823  -8.120898279
## Qtom.I.SB.38      5.95056102  -3.0138882  -1.642721335  -3.023557766
## Qtom.I.SB.40      4.40870981  -0.9299994  -1.929221401  -3.408222391
## Qtom.I.SB.41      5.55411395  -3.9247050  -3.217996583  -2.371370049
## Qtom.I.SB.42      6.72336026  -4.1904861  -4.155209777  -5.144860157
## Qtom.I.SB.44      6.95118909  -5.0978620  -5.435809532  -5.198585599
## Qtom.I.SB.45      7.21265909  -5.1294210  -5.172624884  -4.573408305
## Qtom.I.SB.73      5.70218908  -3.1120950   2.073826513  -4.958905880
## Qtom.I.SB.74      6.08875094  -2.8252751   3.158647936  -6.301482893
## Qtom.I.VEN.01     8.12681052  -6.1504759   9.160328484 -17.193606134
## Qtom.I.VEN.02     7.82427446  -5.6395637  10.294638819 -19.345819310
## Qtom.I.VEN.03     8.10367193  -5.7467278   7.084328177 -13.356785334
## Qtom.I.VEN.04     7.83774623  -5.3674521   9.062751030 -18.540170917
## Qtom.I.VEN.05     7.46895375  -4.9442021   5.423989008 -11.110468727
## Qtom.JJK.SB.01    5.17668030  -0.7371003   1.333224330 -12.595401117
## Qtom.JJK.SB.03    3.34431593  -1.5301235   1.022080628  -1.418821216
## Qtom.JJK.SB.07    5.08751524  -1.6216405   0.720477069  -9.514924743
## Qtom.JJK.SB.08    4.75852716  -1.0985257   1.003089880  -7.800641379
## Qtom.JJK.SB.09    2.49877126   0.6084852  -0.012964631  -4.040708234
## Qtom.JJK.SB.10    3.26714258   1.6010341  -0.343927465  -7.917208578
## Qtom.JR.SB.01     5.15360223  -0.5572382   0.701601325 -11.069137607
## Qtom.S.LA.186     3.04719808   6.2510928  22.341988658   8.223861857
##                           PC5          PC6
## Qchr.A.232       -3.564844066  10.81088072
## Qchr.A.275       -1.509462590  10.12141151
## Qchr.A.ED.102    -1.230581357   9.92656259
## Qchr.A.ELD.53    -3.281107077  12.11580603
## Qchr.A.KER.16     2.649513447   4.99416790
## Qchr.A.LA.02      0.270005310  -0.83926845
## Qchr.A.LA.215     3.612308637   7.78665553
## Qchr.A.LA.222     2.506451078   7.23919167
## Qchr.A.SIS.309   -2.836264522  12.23811901
## Qchr.A.PLU.65    -3.581678193  11.49694186
## Qchr.A.SBD.83     2.481296551   0.90956514
## Qchr.I.LA.20     -0.634244933  -0.36868671
## Qchr.I.LA.25     -0.246303830   0.74331698
## Qchr.I.SB.02    -11.036801250 -27.08051461
## Qchr.I.SB.03    -14.694496147 -30.64968683
## Qchr.JJK.SB.13    2.080681303  -1.35841826
## Qchr.JR.SB.01     7.312706834   2.47727041
## Qspp.I.LA.03      1.384011134   4.30632875
## Qspp.I.LA.04      0.413970858   1.80409640
## Qspp.I.LA.14     -0.942923245  -0.18014525
## Qtom.A.LA.169     2.061388858   1.92142421
## Qtom.A.LA.175     0.937321402   3.12236542
## Qtom.A.LA.176     1.413135783   3.77594017
## Qtom.A.LA.180     1.132280922   1.52612241
## Qtom.A.LA.181     0.557111803  -0.09914675
## Qtom.A.LA.183    47.902489511 -28.43219799
## Qtom.A.LA.190    -7.836361987  -0.49332260
## Qtom.A.LA.191    50.251207569 -30.66314787
## Qtom.A.LA.198   -13.769165846  -0.97002287
## Qtom.A.LA.200    -9.516032354  -0.54741158
## Qtom.A.LA.204   -23.314903855  -4.14886948
## Qtom.A.LA.207   -24.335013082  -4.92158112
## Qtom.A.LA.208    -4.251551852   0.15913446
## Qtom.A.LA.209   -15.572432145  -0.74131972
## Qtom.A.LA.218     3.025458763   7.18427002
## Qtom.A.LA.221    -1.698252986   0.45425014
## Qtom.A.SB.324     1.538256666   1.91367441
## Qtom.A.SB.325    -1.227650346   0.52614495
## Qtom.A.SB.327    -1.530882319   0.32865431
## Qtom.A.SB.329    -1.245171433   0.83099945
## Qtom.A.SB.330    -0.963800641   0.65235035
## Qtom.A.SB.331     0.457501204   0.98826330
## Qtom.A.SB.332    -1.688057255  -0.33890992
## Qtom.A.SB.333    -1.824533135  -0.60299758
## Qtom.A.SB.335    -2.495162936  -0.07337612
## Qtom.A.SB.336    -2.382557470  -1.41950873
## Qtom.A.SB.337    -2.723389419  -0.85289387
## Qtom.A.SB.338    -2.219558615   0.12725904
## Qtom.A.SB.339    -2.487906364  -0.37922829
## Qtom.A.SB.341     1.318739855   1.46752940
## Qtom.A.SB.342    -0.939191266   0.77963539
## Qtom.A.SB.343    -2.085590873  -0.36059351
## Qtom.A.SB.344    -1.002012860   0.12450880
## Qtom.A.SB.345     0.311426007   1.52361088
## Qtom.A.SB.346    -0.043149782   0.40136030
## Qtom.A.SB.347    -2.642050719   0.04961786
## Qtom.A.SB.348    -1.662939587  -0.25812683
## Qtom.A.SB.352    -3.602220841   2.63015488
## Qtom.A.SB.353    -3.671985447   2.79103687
## Qtom.A.SB.355    -3.115214919   1.71294562
## Qtom.I.BC.10      0.455174316  -3.03163860
## Qtom.I.BC.11     -3.733538261  -7.44223471
## Qtom.I.BC.12     -1.744084351  -4.64547233
## Qtom.I.BC.13     -1.490283071  -4.41408108
## Qtom.I.BC.18     -0.731974424  -4.13339203
## Qtom.I.BC.20      3.769336756   6.26405061
## Qtom.I.BC.21      2.026049659   4.97104154
## Qtom.I.BC.23     -0.967023082   0.63442494
## Qtom.I.BC.24     -3.604877985  -3.09831915
## Qtom.I.BC.32     -0.258311891  -1.44838617
## Qtom.I.BC.33      0.633846140  -0.10412963
## Qtom.I.BC.35     -0.300724567  -1.65960598
## Qtom.I.BC.39      2.735722488   2.50878525
## Qtom.I.BC.4      -1.544958009  -1.87464237
## Qtom.I.BC.40     -1.208342809  -2.58599828
## Qtom.I.BC.41      1.170271038  -0.13518342
## Qtom.I.BC.42      3.977017915   5.57090294
## Qtom.I.BC.43      0.001957359   6.90497918
## Qtom.I.BC.44      1.002424683   8.19256063
## Qtom.I.BC.45      0.135105609   5.49675615
## Qtom.I.BC.46      0.162474841   0.69854200
## Qtom.I.BC.48      1.251045017   1.14288727
## Qtom.I.LA.SC.01   0.450920129   2.59718122
## Qtom.I.LA.02      2.367249716   3.58169391
## Qtom.I.LA.05      3.363868283   4.02651791
## Qtom.I.LA.06     -0.860843507   1.32568886
## Qtom.I.LA.08     -0.703979650   3.61774239
## Qtom.I.LA.10      2.727094675   3.63113405
## Qtom.I.LA.11     -0.650486340   1.71488473
## Qtom.I.LA.12     -1.855366819   0.41977042
## Qtom.I.LA.13     -0.014878624   0.95773590
## Qtom.I.LA.21     -0.349938142   0.90483676
## Qtom.I.LA.22      3.097617942   4.02425054
## Qtom.I.LA.23      1.359059815   3.38804226
## Qtom.I.LA.24     -1.229854421   0.73720953
## Qtom.I.SB.01    -10.907825420 -23.32210018
## Qtom.I.SB.04    -10.451728847 -28.21561983
## Qtom.I.SB.09      2.041940712   1.74984315
## Qtom.I.SB.10     -1.170515477  -2.23821837
## Qtom.I.SB.11      2.690733693   4.72903178
## Qtom.I.SB.12      2.086542874   4.73045282
## Qtom.I.SB.28     -0.900649023  -1.84754043
## Qtom.I.SB.33     -1.711839954  -5.40096735
## Qtom.I.SB.35     -3.915406782  -8.97289452
## Qtom.I.SB.36     -2.771273152  -3.27026644
## Qtom.I.SB.37     -6.520034423 -13.93974120
## Qtom.I.SB.38     -1.493561206  -4.15763975
## Qtom.I.SB.40     -0.275809041  -5.08433728
## Qtom.I.SB.41     -1.641471311  -5.40671077
## Qtom.I.SB.42     -5.766339838 -11.74825856
## Qtom.I.SB.44     -6.951749224 -15.86626659
## Qtom.I.SB.45     -7.100045952 -14.78578829
## Qtom.I.SB.73      3.951487252   3.26535491
## Qtom.I.SB.74      3.796191447   3.54583192
## Qtom.I.VEN.01     8.254038671  16.52833259
## Qtom.I.VEN.02     8.079061139  17.95883021
## Qtom.I.VEN.03     7.261278147  12.81023268
## Qtom.I.VEN.04     7.185126208  15.95400991
## Qtom.I.VEN.05     5.554466613   9.17577805
## Qtom.JJK.SB.01   10.237366461   2.58790819
## Qtom.JJK.SB.03    5.766776543   2.86179806
## Qtom.JJK.SB.07    9.361940538   3.99310041
## Qtom.JJK.SB.08    7.736828303   3.80249204
## Qtom.JJK.SB.09    8.035292394   3.12630525
## Qtom.JJK.SB.10    7.471806289   2.68017145
## Qtom.JR.SB.01     9.700636978   3.32024456
## Qtom.S.LA.186    -7.281850995  -3.48409585
```

```
# plot the individuals ('sites')

# set params
par(cex.axis = 1.2, cex.lab = 1.5)
par(mar = c(5,5,3,1))

# setup colors and points

# color by island
bg <- c('black', "#3c4a8b","#009c85","#84bc5f","#edb829","#f57404","#b30000")

# point shape by species
sp <- c(23, 22, 21)


#png(filename = 'results/pca/PCA_allSamples_axes1-2.png', res = 300, height = 5.5, width = 6, units = 'in')
#pdf(file = 'results/pca/PCA_allSamples_axes1-2.pdf', height = 5.5, width = 6)

par(cex.axis = 1.2, cex.lab = 1.5)
par(mar = c(5,5,3,1))

# PC1 and PC2
choices = c(1,2)

# x and ylab code is ugly, but it's just pulling the variance explained from the RDA summary for each axis
plot(rda, 
     type = 'n', 
     choices = choices,
     xlab = paste('PC', choices[1], ' (', round(rda.info$cont$importance[2, choices[1]], 3)*100, '% variance explained)', sep = ''),
     ylab = paste('PC', choices[2], ' (', round(rda.info$cont$importance[2, choices[2]], 3)*100, '% variance explained)', sep = ''))

points(rda, pch = sp[info$sp], cex = 1.5, display = 'sites', bg = bg[info$island], col = 'black', choices = choices)

legend('topleft', legend = c(levels(info$sp), levels(info$island)), pch = c(sp, rep(15, 8)), col = c(rep('black', 3), bg), cex = 0.8)
```

```
#dev.off()

# PC1 and PC3

#png(filename = 'results/pca/PCA_allSamples_axes1-3.png', res = 300, height = 5.5, width = 6, units = 'in')
#pdf(file = 'results/pca/PCA_allSamples_axes1-3.pdf', height = 5.5, width = 6)

par(cex.axis = 1.2, cex.lab = 1.5)
par(mar = c(5,5,3,1))

choices = c(1,3)
plot(rda, 
     type = 'n', 
     choices = choices,
     xlab = paste('PC', choices[1], ' (', round(rda.info$cont$importance[2, choices[1]], 3)*100, '% variance explained)', sep = ''),
     ylab = paste('PC', choices[2], ' (', round(rda.info$cont$importance[2, choices[2]], 3)*100, '% variance explained)', sep = ''))

points(rda, pch = sp[info$sp], cex = 1.5, display = 'sites', bg = bg[info$island], col = 'black', choices = choices)
```

```
#legend('topleft', legend = c(levels(info$sp), levels(info$island)), pch = c(sp, rep(15, 8)), col = c(rep('black', 3), bg), cex = 1.2)

#dev.off()
```

#### 3.2 Only Channel Islands, exclude Guadalupe Island

```
# plot without guadalupe island

imp.sub <- imp[! info$island %in% c("Guadalupe Island"),]
info.sub <- info[! info$island %in% c("Guadalupe Island"),]

rda <- rda(imp.sub, scale = T)

rda
```

```
## Call: rda(X = imp.sub, scale = T)
## 
##               Inertia Rank
## Total          574832     
## Unconstrained  574832  104
## Inertia is correlations 
## 
## Eigenvalues for unconstrained axes:
##   PC1   PC2   PC3   PC4   PC5   PC6   PC7   PC8 
## 15471  9626  9177  8626  8482  8274  7938  7749 
## (Showing 8 of 104 unconstrained eigenvalues)
```

```
#summary(rda)
screeplot(rda)
```

```
# save summary
rda.info <- summary(rda)
rda.info$sites
```

```
##                         PC1           PC2          PC3          PC4
## Qchr.A.232      -16.4453187 -18.948827643   5.83945933  -4.29747530
## Qchr.A.275      -16.4349914 -17.188430323   6.67467342  -2.43632067
## Qchr.A.ED.102   -19.7544354 -14.577910441   6.71433061  -1.96789449
## Qchr.A.ELD.53   -20.7163312 -18.824831870   7.63891979  -4.34999099
## Qchr.A.KER.16   -14.6786135  -3.887136072   3.92238000   2.38605707
## Qchr.A.LA.02    -18.9649971   1.839731030   0.07485011   1.06870898
## Qchr.A.LA.215    -2.1652544   5.220059200   4.43402523   2.98136240
## Qchr.A.LA.222    -9.1361231   4.756228940   3.48043747   2.46699992
## Qchr.A.SIS.309  -17.8828543 -19.779859891   7.36131775  -3.85136734
## Qchr.A.PLU.65   -20.6068504 -19.097813598   6.01790872  -4.49068418
## Qchr.A.SBD.83   -15.9782079   2.356799732   2.47039977   2.77599097
## Qchr.I.LA.20     -5.3488226  -3.244037836   0.65663803  -0.36272969
## Qchr.I.LA.25     -3.4531235   0.448664043   0.94533366  -0.06290542
## Qchr.I.SB.02     -9.3225346 -10.915517219 -22.38528856  -7.72270425
## Qchr.I.SB.03    -11.0734280 -12.396261110 -24.49121229 -11.51083248
## Qchr.JJK.SB.13  -11.1302794  -5.375275884  -9.54723019   3.35124433
## Qchr.JR.SB.01    -2.9190042  -1.285381302  -6.49157652   7.22764542
## Qspp.I.LA.03     -8.3794562   1.880644876  -5.11242258   2.04201875
## Qspp.I.LA.04      1.1659279   0.299790098  -0.21095263   0.63067267
## Qspp.I.LA.14     -2.7151570  -3.351114550   0.42009599  -0.73695085
## Qtom.A.LA.169    -2.5494192   2.446134013   1.30851590   1.86489615
## Qtom.A.LA.175     0.7364065   5.280205283  -0.84545363   0.73883094
## Qtom.A.LA.176    -2.9202277   5.978098109  -0.72749559   1.27918326
## Qtom.A.LA.180    -3.9697391   1.752844309   2.99561078   0.78450782
## Qtom.A.LA.181   -14.6517384   4.120843959  -0.36682135   1.00246220
## Qtom.A.LA.183    -9.8377309  11.613048703  13.73560256  45.86487525
## Qtom.A.LA.190    -8.7920773  14.902345642   5.64157931  -8.50539849
## Qtom.A.LA.191   -11.2200330  12.633300196  13.77649550  48.62142791
## Qtom.A.LA.198    -9.9033569  22.357391195   3.30465762 -13.47335529
## Qtom.A.LA.200    -1.6723063  17.352900307   0.42927850  -9.04139956
## Qtom.A.LA.204    -8.1639458  28.731139648   4.95930898 -23.01901027
## Qtom.A.LA.207    -9.6225355  29.150865178   4.33799909 -24.39900858
## Qtom.A.LA.208     0.2457142   7.494397306  -3.74869377  -3.64362995
## Qtom.A.LA.209    -8.2565553  23.947365285   3.98343499 -15.45827728
## Qtom.A.LA.218    -9.4343610   5.038881944   3.35643624   3.02004174
## Qtom.A.LA.221    -2.1626534  13.893675755   2.22464744  -2.00325012
## Qtom.A.SB.324     6.3573166  -2.249314884   9.22784825   0.34969695
## Qtom.A.SB.325     7.9484831  -4.364601022  11.78321469  -2.32422738
## Qtom.A.SB.327     8.4326683  -4.351984188  11.24979980  -2.54864659
## Qtom.A.SB.329     7.3028635  -3.365777014   8.76431297  -2.07378860
## Qtom.A.SB.330     7.5599069  -3.039972920   8.09985001  -1.69609087
## Qtom.A.SB.331     6.9331057  -3.338840085   9.94092387  -0.66585284
## Qtom.A.SB.332     8.5333443  -4.709571282  11.48423080  -2.70930117
## Qtom.A.SB.333     7.8445829  -4.437606685  10.93934080  -2.71275048
## Qtom.A.SB.335     8.7406953  -3.676599974   8.83296765  -3.15427358
## Qtom.A.SB.336     8.9468782  -4.784029688   9.82914879  -2.98287515
## Qtom.A.SB.337     9.8172062  -4.210228695   9.56330896  -3.35138010
## Qtom.A.SB.338     8.5028203  -4.525610934  10.41800420  -3.06850847
## Qtom.A.SB.339     9.3565643  -4.428003228   9.48363573  -3.21737196
## Qtom.A.SB.341     6.4641267  -2.348579833   8.11758066   0.34419312
## Qtom.A.SB.342     7.4515510  -3.448732864   7.96550555  -1.65685299
## Qtom.A.SB.343     7.8974913  -3.812926145   6.85254137  -2.61931044
## Qtom.A.SB.344     7.6444535  -3.202499635   7.33403271  -1.61250846
## Qtom.A.SB.345     6.3097240  -2.213696041   7.22998076  -0.50553653
## Qtom.A.SB.346     7.2079606  -2.976130625   8.46996611  -0.90866555
## Qtom.A.SB.347     8.5124115  -2.553385868   6.42914999  -3.01880146
## Qtom.A.SB.348     7.5925271  -3.633740258   7.40418809  -2.17816487
## Qtom.A.SB.352    10.2245946  -6.058259900  15.39708039  -5.01273331
## Qtom.A.SB.353    10.2740975  -6.086436424  15.27981304  -5.07879425
## Qtom.A.SB.355     9.7242561  -5.048424955  11.47102432  -4.06218272
## Qtom.I.LA.SC.01 -13.7150636   3.426287839  -5.15315013   1.39101723
## Qtom.I.LA.02     -1.6328691   5.555862543  -0.85076174   2.14784483
## Qtom.I.LA.05     -7.1065035  -0.593485055  -2.34619406   3.79099278
## Qtom.I.LA.06     -9.7728062   2.006972144  -6.11658302  -0.04637624
## Qtom.I.LA.08      1.8393620   2.410166254  -3.86483183  -0.36278638
## Qtom.I.LA.10      2.4823603   0.429852808   2.01776814   2.16766790
## Qtom.I.LA.11      4.9272029   0.153181117   0.60477674  -0.65513814
## Qtom.I.LA.12      0.9379410   0.310888875   0.05262495  -1.57564891
## Qtom.I.LA.13     -4.0384606   1.118191915   1.15293107   0.29765796
## Qtom.I.LA.21      5.4967956  -0.118460032   2.43940556  -0.44796774
## Qtom.I.LA.22     -5.0285620   0.119679231   0.78417636   2.84337680
## Qtom.I.LA.23      2.8427323   0.791715854   0.15425032   1.14430541
## Qtom.I.LA.24      1.9051855   0.596954030  -0.09767402  -1.00128066
## Qtom.I.SB.01      5.1127621  -7.022204948 -16.57145562  -6.85206594
## Qtom.I.SB.04     -9.9149389 -12.386586324 -23.49987506  -7.29022765
## Qtom.I.SB.09      7.0711302   1.291271238  -8.18078372   2.55522354
## Qtom.I.SB.10      8.4933985   0.988486114 -17.80792997   0.71264374
## Qtom.I.SB.11      8.0597717   3.282335442 -12.32182937   3.31921521
## Qtom.I.SB.12      7.9854292   3.255261419 -13.18336379   2.82382101
## Qtom.I.SB.28      3.6344808  -1.581949180  -7.71044929   0.20414643
## Qtom.I.SB.33      5.0941399  -2.758616111  -5.13796280  -0.48324353
## Qtom.I.SB.35      5.9419266  -5.576285160  -4.36381803  -2.48212292
## Qtom.I.SB.36      5.5490915  -1.766963073  -6.33737582  -1.58957039
## Qtom.I.SB.37      5.7821717  -4.560771908  -8.36405700  -4.02204292
## Qtom.I.SB.38      5.2764356  -1.904870477  -2.56637265  -0.72924981
## Qtom.I.SB.40      2.7172522  -1.883237660  -2.77043855   0.61979889
## Qtom.I.SB.41      5.6329107  -3.472996092  -2.59362167  -0.87157143
## Qtom.I.SB.42      6.4754534  -4.347055728  -4.77831666  -3.65873273
## Qtom.I.SB.44      7.3186588  -5.614068656  -5.10827097  -4.47097158
## Qtom.I.SB.45      7.4341571  -5.244486589  -4.46035143  -4.71156588
## Qtom.I.SB.73      5.2153992   1.358965769  -4.21365883   3.69511551
## Qtom.I.SB.74      5.2622006   2.249338188  -5.54011593   3.67270425
## Qtom.I.VEN.01     8.9475714   5.743359095 -13.89662562   7.39519143
## Qtom.I.VEN.02     8.4876700   6.551333798 -15.78733259   7.35577379
## Qtom.I.VEN.03     8.6052031   4.392726506 -10.69864878   6.46753802
## Qtom.I.VEN.04     8.3346045   5.555134711 -15.10475070   6.72169294
## Qtom.I.VEN.05     7.6348732   3.319024353  -9.13477133   5.16612312
## Qtom.JJK.SB.01    3.1408972   0.349947764 -11.08810787   9.66150649
## Qtom.JJK.SB.03    2.6114515   0.916169976  -0.91502706   5.10893026
## Qtom.JJK.SB.07    3.7516197   0.004341403  -8.11140487   8.60183101
## Qtom.JJK.SB.08    3.1568595   0.462205827  -6.62051879   7.13472849
## Qtom.JJK.SB.09    0.4939780  -0.010943750  -3.46587403   7.25441184
## Qtom.JJK.SB.10    0.1793815  -0.626798227  -7.13879972   7.04265852
## Qtom.JR.SB.01     2.9596603  -0.033966692  -9.63995077   9.00704670
## Qtom.S.LA.186    -3.0481309  21.106087581   3.99048176  -7.36141416
##                          PC5         PC6
## Qchr.A.232       12.81018905 -19.0759732
## Qchr.A.275       11.13593100 -13.9357332
## Qchr.A.ED.102    10.60839451  -7.7531010
## Qchr.A.ELD.53    14.01822427 -15.4056538
## Qchr.A.KER.16     4.31052339   4.9055229
## Qchr.A.LA.02     -3.38583261  18.0821082
## Qchr.A.LA.215     5.19049765  19.5302873
## Qchr.A.LA.222     4.37756024  26.9680922
## Qchr.A.SIS.309   13.58061932 -16.2781065
## Qchr.A.PLU.65    14.25803639 -18.6703172
## Qchr.A.SBD.83    -1.35993411  14.9824020
## Qchr.I.LA.20     -0.95650320   3.6932434
## Qchr.I.LA.25      0.23659891   6.5067649
## Qchr.I.SB.02    -27.42753317   2.7825310
## Qchr.I.SB.03    -30.99119822   2.5260681
## Qchr.JJK.SB.13   -0.24532985   6.7150979
## Qchr.JR.SB.01     3.88051001   5.8562334
## Qspp.I.LA.03      4.26543609   9.0146892
## Qspp.I.LA.04      1.24834490   5.2701782
## Qspp.I.LA.14     -0.88130156   4.0750965
## Qtom.A.LA.169     1.62028085   3.5017264
## Qtom.A.LA.175     3.30185671   2.5881082
## Qtom.A.LA.176     3.73524092   5.0002371
## Qtom.A.LA.180     0.83986441   2.4347183
## Qtom.A.LA.181    -0.59621702   9.1593276
## Qtom.A.LA.183   -23.41112950 -18.9375230
## Qtom.A.LA.190    -1.06568505  -5.9535040
## Qtom.A.LA.191   -25.31085193 -20.9852748
## Qtom.A.LA.198    -0.85144130  -2.4250028
## Qtom.A.LA.200     0.18818091  -6.7925114
## Qtom.A.LA.204    -4.32402884 -14.6756934
## Qtom.A.LA.207    -5.17614399 -15.5022548
## Qtom.A.LA.208     0.89603466  -2.3318725
## Qtom.A.LA.209    -0.96232475  -4.1836936
## Qtom.A.LA.218     4.16736258  27.2924945
## Qtom.A.LA.221     1.32939279  -4.9193392
## Qtom.A.SB.324    -0.83284199   3.1422371
## Qtom.A.SB.325    -3.53815380   3.2487430
## Qtom.A.SB.327    -3.58424802   2.7559829
## Qtom.A.SB.329    -1.87305468   2.0334010
## Qtom.A.SB.330    -2.03758403   2.2493293
## Qtom.A.SB.331    -1.88990351   2.7140335
## Qtom.A.SB.332    -3.90864439   1.8451023
## Qtom.A.SB.333    -3.95006549   1.5778814
## Qtom.A.SB.335    -2.76284541   0.8716063
## Qtom.A.SB.336    -4.20452572   1.3665651
## Qtom.A.SB.337    -3.76987369   1.1136850
## Qtom.A.SB.338    -3.02836745   1.0248358
## Qtom.A.SB.339    -3.55303451   0.9025714
## Qtom.A.SB.341    -0.60547918   2.3555284
## Qtom.A.SB.342    -1.68958933   1.7518200
## Qtom.A.SB.343    -2.53176904   1.0481481
## Qtom.A.SB.344    -2.09350574   1.7331933
## Qtom.A.SB.345    -0.44574252   2.2056275
## Qtom.A.SB.346    -1.95471223   2.3550994
## Qtom.A.SB.347    -2.15477451   0.9900193
## Qtom.A.SB.348    -2.38914977   1.0642645
## Qtom.A.SB.352    -3.58165014   2.7660513
## Qtom.A.SB.353    -3.37642106   2.6886006
## Qtom.A.SB.355    -2.75637958   1.6425743
## Qtom.I.LA.SC.01   1.31147486  15.5454560
## Qtom.I.LA.02      3.80091696   4.1640688
## Qtom.I.LA.05      3.41854871   8.1127053
## Qtom.I.LA.06      0.82076685   8.1183215
## Qtom.I.LA.08      3.43169914   1.9456891
## Qtom.I.LA.10      2.87121696   2.4919649
## Qtom.I.LA.11      1.01346008   2.1628360
## Qtom.I.LA.12      0.00673386   3.4959236
## Qtom.I.LA.13     -0.17793310   8.6667214
## Qtom.I.LA.21      0.22609288   2.2486129
## Qtom.I.LA.22      3.22873312   4.8250794
## Qtom.I.LA.23      3.07143045   3.3508148
## Qtom.I.LA.24      0.39529075   3.0034476
## Qtom.I.SB.01    -16.94076720 -10.0180345
## Qtom.I.SB.04    -28.35191795   3.6665542
## Qtom.I.SB.09      4.86491846  -5.0360972
## Qtom.I.SB.10      2.97570455  -9.4800546
## Qtom.I.SB.11      7.90549667  -5.8501705
## Qtom.I.SB.12      8.41002045  -7.0133168
## Qtom.I.SB.28      0.59676425  -4.3855046
## Qtom.I.SB.33     -2.68366440  -4.5496214
## Qtom.I.SB.35     -6.16727586  -5.3442277
## Qtom.I.SB.36     -1.11354923  -4.0665075
## Qtom.I.SB.37     -9.26975390  -7.4530176
## Qtom.I.SB.38     -2.27295728  -4.0564524
## Qtom.I.SB.40     -2.42069783  -3.2033748
## Qtom.I.SB.41     -3.95499715  -2.7832252
## Qtom.I.SB.42     -7.20344996  -8.2291305
## Qtom.I.SB.44    -10.51798916 -10.1657957
## Qtom.I.SB.45     -9.72255491  -9.1400918
## Qtom.I.SB.73      4.54249234   0.6420494
## Qtom.I.SB.74      4.93941655   0.9559504
## Qtom.I.VEN.01    18.99136321  -8.7004559
## Qtom.I.VEN.02    20.61850700  -9.4138211
## Qtom.I.VEN.03    14.74638224  -6.7112203
## Qtom.I.VEN.04    18.70916084  -8.6616700
## Qtom.I.VEN.05    11.12789451  -4.7497260
## Qtom.JJK.SB.01    5.64645120   5.7321341
## Qtom.JJK.SB.03    3.48463385   2.7222593
## Qtom.JJK.SB.07    5.94160093   6.0547102
## Qtom.JJK.SB.08    5.54626995   4.6251481
## Qtom.JJK.SB.09    3.90068881   6.0387294
## Qtom.JJK.SB.10    4.22807917   6.1717069
## Qtom.JR.SB.01     5.63884123   6.9082681
## Qtom.S.LA.186    -2.15485152 -11.1439091
```

```
# plot the individuals ('sites')

# set params
par(cex.axis = 1.2, cex.lab = 1.5)
par(mar = c(5,5,3,1))

# setup colors and points

# color by island
bg <- c('black', "#3c4a8b","#009c85","#84bc5f","#edb829","#f57404")

# point shape by species
sp <- c(23, 22, 21)


# png(filename = 'results/pca/PCA_California_axes1-2.png', res = 300, height = 5.5, width = 6, units = 'in')
# pdf(file = 'results/pca/PCA_California_axes1-2.pdf', height = 5.5, width = 6)


par(cex.axis = 1.2, cex.lab = 1.5)
par(mar = c(5,5,3,1))

# PC1 and PC2
choices = c(1,2)

# x and ylab code is ugly, but it's just pulling the variance explained from the RDA summary for each axis
plot(rda, 
     type = 'n', 
     choices = choices,
     xlab = paste('PC', choices[1], ' (', round(rda.info$cont$importance[2, choices[1]], 3)*100, '% variance explained)', sep = ''),
     ylab = paste('PC', choices[2], ' (', round(rda.info$cont$importance[2, choices[2]], 3)*100, '% variance explained)', sep = ''))

points(rda, pch = sp[info.sub$sp], cex = 1.5, display = 'sites', bg = bg[info.sub$island], col = 'black', choices = choices)
```

```
#legend('topleft', legend = c(levels(info$sp), levels(info$island)), pch = c(sp, rep(15, 8)), col = c(rep('black', 3), bg), cex = 0.8)

#dev.off()

# PC1 and PC3

# png(filename = 'results/pca/PCA_California_axes1-3.png', res = 300, height = 5.5, width = 6, units = 'in')
# pdf(file = 'results/pca/PCA_California_axes1-3.pdf', height = 5.5, width = 6)

par(cex.axis = 1.2, cex.lab = 1.5)
par(mar = c(5,5,3,1))

choices = c(1,3)
plot(rda, 
     type = 'n', 
     choices = choices,
     xlab = paste('PC', choices[1], ' (', round(rda.info$cont$importance[2, choices[1]], 3)*100, '% variance explained)', sep = ''),
     ylab = paste('PC', choices[2], ' (', round(rda.info$cont$importance[2, choices[2]], 3)*100, '% variance explained)', sep = ''))

points(rda, pch = sp[info.sub$sp], cex = 1.5, display = 'sites', bg = bg[info.sub$island], col = 'black', choices = choices)
```

```
#legend('topleft', legend = c(levels(info$sp), levels(info$island)), pch = c(sp, rep(15, 8)), col = c(rep('black', 3), bg), cex = 1.2)

#dev.off()
```

#### 3.3 Only Northern Channel Islands

```
# plot just SRI and SCI for easier comparison with Qagr data

imp.sub <- imp[info$island %in% c("Mainland", "Santa Rosa Island", "Santa Cruz Island"),]
info.sub <- info[info$island %in% c("Mainland", "Santa Rosa Island", "Santa Cruz Island"),]

rda <- rda(imp.sub, scale = T)

rda
```

```
## Call: rda(X = imp.sub, scale = T)
## 
##               Inertia Rank
## Total          544653     
## Unconstrained  544653   62
## Inertia is correlations 
## 
## Eigenvalues for unconstrained axes:
##   PC1   PC2   PC3   PC4   PC5   PC6   PC7   PC8 
## 21972 14284 13454 12928 12531 12349 12319 12257 
## (Showing 8 of 62 unconstrained eigenvalues)
```

```
#summary(rda)
screeplot(rda)
```

```
# plots

# setup colors and points

# color by island
bg <- c('black', "#3c4a8b","#009c85","#84bc5f","#edb829","#f57404","#b30000")

# point shape by species
sp <- c(23, 22, 21)
#sp <- c(5, 0, 1)

# PC1 and PC2
#png(file = 'results/redundancy_analysis/rda_plot_indivs_clim_Qtom107.Qchr17.Qssp3_SantaRosa_SantaCruz_Mainland_PC1_PC2.png', height = 6, width = 6, res = 300, units = 'in')
choices = c(1,2)
plot(rda, type = 'n', choices = choices)
points(rda, pch = sp[info.sub$sp], cex = 1, display = 'sites', bg = bg[info.sub$island], col = 'black', choices = choices)
#points(rda, pch = sp[info.sub$sp], cex = 2, display = 'sites', col = bg[info.sub$island], choices = choices)
legend('bottomleft', legend = c(levels(info.sub$sp), "Mainland Qchr", "Santa Rosa Island", "Santa Cruz Island"), pch = c(sp, rep(15, 8)), col = c(rep('black', 4), "#3c4a8b","#009c85"), cex = 1)
```

```
#dev.off()


# PC1 and PC3
#png(file = 'results/redundancy_analysis/rda_plot_indivs_clim_Qtom107.Qchr17.Qssp3_SantaRosa_SantaCruz_Mainland_PC1_PC3.png', height = 6, width = 6, res = 300, units = 'in')
choices = c(1,3)
plot(rda, type = 'n', choices = choices)
points(rda, pch = sp[info.sub$sp], cex = 1, display = 'sites', bg = bg[info.sub$island], col = 'black', choices = choices)
```

```
#dev.off()

###########################
# no mainland

imp.sub <- imp[info$island %in% c("Santa Rosa Island", "Santa Cruz Island"),]
info.sub <- info[info$island %in% c("Santa Rosa Island", "Santa Cruz Island"),]

rda <- rda(imp.sub, scale = T)

rda
```

```
## Call: rda(X = imp.sub, scale = T)
## 
##               Inertia Rank
## Total          514282     
## Unconstrained  514282   53
## Inertia is correlations 
## 
## Eigenvalues for unconstrained axes:
##   PC1   PC2   PC3   PC4   PC5   PC6   PC7   PC8 
## 19769 15780 15595 15367 14895 14415 13541 12569 
## (Showing 8 of 53 unconstrained eigenvalues)
```

```
#summary(rda)
screeplot(rda)
```

```
# plots

# setup colors and points

# color by island
bg <- c('black', "#3c4a8b","#009c85","#84bc5f","#edb829","#f57404","#b30000")

# point shape by species
sp <- c(23, 22, 21)
#sp <- c(5, 0, 1)

# PC1 and PC2
#png(file = 'results/redundancy_analysis/rda_plot_indivs_clim_Qtom107.Qchr17.Qssp3_SantaRosa_SantaCruz_PC1_PC2.png', height = 6, width = 6, res = 300, units = 'in')
choices = c(1,2)
plot(rda, type = 'n', choices = choices)
points(rda, pch = sp[info.sub$sp], cex = 1, display = 'sites', bg = bg[info.sub$island], col = 'black', choices = choices)
#points(rda, pch = sp[info.sub$sp], cex = 2, display = 'sites', col = bg[info.sub$island], choices = choices)
legend('topleft', legend = c(levels(info.sub$sp), "Mainland Qchr", "Santa Rosa Island", "Santa Cruz Island"), pch = c(sp, rep(15, 8)), col = c(rep('black', 3), "#3c4a8b","#009c85"), cex = 1)
```

```
#dev.off()


# PC1 and PC3
#png(file = 'results/redundancy_analysis/rda_plot_indivs_clim_Qtom107.Qchr17.Qssp3_SantaRosa_SantaCruz_PC1_PC3.png', height = 6, width = 6, res = 300, units = 'in')
choices = c(1,3)
plot(rda, type = 'n', choices = choices)
points(rda, pch = sp[info.sub$sp], cex = 1, display = 'sites', bg = bg[info.sub$island], col = 'black', choices = choices)
```

```
#dev.off()

# combined fig

#png(file = 'results/redundancy_analysis/rda_plot_indivs_clim_Qtom107.Qchr17.Qssp3_SantaRosa_SantaCruz_PC1-PC3.png', height = 6, width = 12, res = 300, units = 'in')
par(mfrow = c(1,2))
choices = c(1,2)
plot(rda, type = 'n', choices = choices, main = substitute(paste(italic("Q. tomentella"))))
points(rda, pch = sp[info.sub$sp], cex = 1, display = 'sites', bg = bg[info.sub$island], col = 'black', choices = choices)

legend('bottomright', legend = c(levels(info.sub$sp), "Santa Rosa Island", "Santa Cruz Island"), pch = c(sp, rep(15, 8)), col = c(rep('black', 3), "#3c4a8b","#009c85"), cex = 0.8)


# PC1 and PC3
choices = c(1,3)
plot(rda, type = 'n', choices = choices)
points(rda, pch = sp[info.sub$sp], cex = 1, display = 'sites', bg = bg[info.sub$island], col = 'black', choices = choices)
```

```
#dev.off()
```
